# Discovery of New Broad-Spectrum Anti-Infectives for Eukaryotic Pathogens Using Bioorganometallic Chemistry

**DOI:** 10.1101/2023.06.28.546819

**Authors:** Yan Lin, Hyeim Jung, Christina A. Bulman, James Ng, Robin Vinck, Cillian O’Beirne, Matthew S. Moser, Nancy Tricoche, Ricardo Peguero, Robert W. Li, Joseph F. Urban, Patrice Le Pape, Fabrice Pagniez, Marco Moretto, Tobias Weil, Sara Lustigman, Kevin Cariou, Makedonka Mitreva, Judy A. Sakanari, Gilles Gasser

**Affiliations:** Chimie ParisTech, PSL University, CNRS, Institute of Chemistry for Life and Health Sciences, Laboratory for Inorganic Chemical Biology, 75005 Paris, France; Division of Infectious Diseases, Department of Medicine, Washington University School of Medicine, St. Louis, MO 63110, USA; University of California, San Francisco, Department of Pharmaceutical Chemistry, San Francisco, CA 94158, USA; Molecular Parasitology, New York Blood Center, Lindsley F. Kimball Research Institute, New York, NY 10065, USA; Animal Parasitic Diseases Laboratory, United States Department of Agricultural Research Service (USDA-ARS), Beltsville, MD 20705, USA; Diet, Genomics and Immunology Laboratory, United States Department of Agriculture, Beltsville, MD 20705, USA; Nantes Université, CHU de Nantes, Cibles et Médicaments des Infections et de l’Immunité, IICiMed, UR 1155, F-44000 Nantes, France; Fondazione Edmund Mach Via E. Mach 1, Research and Innovation Centre, Via E. Mach 1, 38010 San Michele all’Adige, Italy; McDonnell Genome Institute, Washington University School of Medicine, St. Louis, MO 63108, USA

## Abstract

Drug resistance observed with many anti-infectives clearly highlights the need for new broad-spectrum agents to treat especially neglected tropical diseases (NTDs) caused by eukaryotic parasitic pathogens including fungal infections. Since these diseases target the most vulnerable communities who are disadvantaged by health and socio-economic factors, new agents should be, if possible, easy-to-prepare to allow for commercialization based on their low cost. In this study, we show that simple modification of one of the most well-known antifungal drugs, fluconazole, with organometallic moieties not only improves the activity of the parent drug but also broadens the scope of application of the new derivatives. These compounds were highly effective *in vivo* against pathogenic fungal infections and potent against parasitic worms such as *Brugia,* which causes lymphatic filariasis and *Trichuris,* one of the soil-transmitted helminths that infects millions of people globally. Notably, the identified molecular targets indicate a mechanism of action that differs greatly from the parental antifungal drug, including targets involved in biosynthetic pathways that are absent in humans, offering great potential to expand our armamentarium against drug-resistant fungal infections and NTDs targeted for elimination by 2030. Overall, the discovery of these new compounds with broad-spectrum activity opens new avenues for the development of treatments for several current human infections, either caused by fungi or by parasites, including other NTDs, as well as newly emerging diseases.

**ONE-SENTENCE SUMMARY:** Simple derivatives of the well-known antifungal drug fluconazole were found to be highly effective *in vivo* against fungal infections, and also potent against the parasitic nematode *Brugia,* which causes lymphatic filariasis and against *Trichuris,* one of the soil-transmitted helminths that infects millions of people globally.

## INTRODUCTION

The need for new broad-spectrum anti-infective agents is becoming urgent considering the modest number of compounds available for treatment and the problem of drug resistance observed with many of them. For example, the discovery of new drugs to treat neglected tropical diseases (NTDs) (*1*) and fungal infections (*2–4*) is a top priority as many of these pathogens are developing resistance to the available drugs. Crucially, in order to be accessible to the entire world’s population and not only for the more developed countries, these newly repurposed or derivatized agents should be, if possible, easy-to-prepare to allow for commercialization based on their low cost (*5*).

To tackle this problem for NTDs, our earlier approach used highly annotated nematode genomic information to computationally prioritize genes that are essential for the survival of parasitic helminths and integrated these data with experimental screening studies using repurposed drugs against filarial worms *in vitro* (*6*). By integrating computationally prioritized compounds with the list of experimentally active hits, we identified several repurposed compounds, which are amenable to structural optimization and could be further explored as potentially new lead compounds. One of the major groups of compounds that was identified using this strategy was the azoles, which are five-membered heterocyclic compounds that incorporate a nitrogen atom and at least one other non-carbon atom in its ring and have antifungal activity that targets membrane sterol synthesis (*7*).

In our earlier reports, we have studied azoles, in particular fluconazole (FCZ), a first generation triazole approved for use as a broad-spectrum antifungal agent. FCZ has a clinically favorable pharmacokinetic profile and has been used as a safe and effective treatment for several fungal infections since the 1990s (*8, 9*). A successful strategy for developing low-cost, alternative therapies to improve the potency and therapeutic scope of an organic anti-infective drug is the use of metal complexes (*10, 11*) or the addition of a ferrocenyl moiety (*12*) to the parent drug, as exemplified with the antimalarial ferroquine, (*13, 14*) which successfully completed phase II clinical trials (*15, 16*). Importantly, we demonstrated that the replacement of one of the triazoles of FCZ by an organometallic moiety (e.g., ferrocene and ruthenocene) allowed unveiling of a subset of FCZ analogues (**1a-4a**, Fig. 1A) more effective than the parent FCZ against various fungal strains (*17*). In the present study, we expanded on our previous work and assessed the efficacy of eight newly synthesized FCZ analogs (**5a-12a**; Fig. 1B) that are metabolically more stable than the previous series tested, on a panel of clinical fungal isolates, including highly drug-resistant strains. We then tested all twelve FCZ analogs (**1a-12a**) on two phylogenetically divergent parasitic nematodes, *Brugia pahangi* and *Trichuris muris*. We demonstrate that structural modification of FCZ not only improves the activity of the parent drug but also allows us to broaden the scope of application of these derivatives. Here, we unveil several easy-to-prepare agents that are not only effective *in vivo* against fungal pathogens but also are potent against parasitic worms: *Brugia* that causes lymphatic filariasis and *Trichuris,* one of the soil-transmitted helminths, that infects millions of people globally. This broad-spectrum activity reveals new avenues for the treatment of various infections by eukaryotic organisms that cause major human diseases, including other NTDs and newly emerging diseases.

**Fig. 1.**
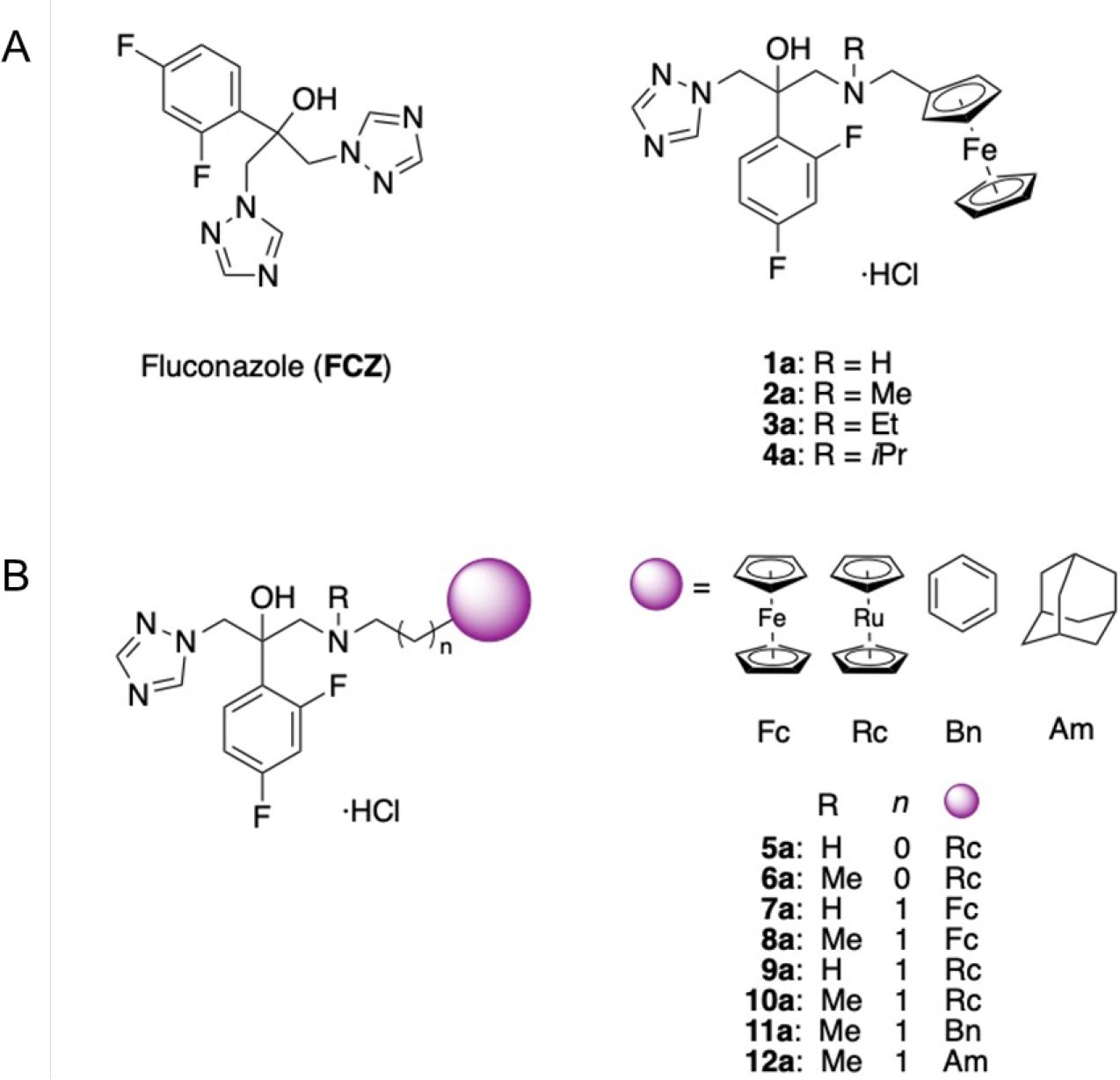
(**A**) The structure of **FCZ** and its derivatives **1a-4a** (*17*) was reported in the previous work. (**B**) The structures of newly developed FCZ derivatives **5a-12a** have been developed in this study.

## RESULTS

### Synthesis and characterization of 8 novel derivatives based on the parent compound FCZ

Eight novel derivatives of FCZ (**5a-12a**, Fig. 1B) were prepared as hydrochloride salts to improve their water solubility and hence their bioavailability. Of note, in order to obtain initial insights into their biological activity, we synthesized these compounds as racemates to avoid an asymmetric synthesis. Details on synthesis and characterization of the complexes can be found in the Supplemental Materials. We then studied the stability of potential drug candidates, including the 4 from previous work (Fig. 1A; **1a-4a**) **1a-12a** in DMSO-d_6_ and D_2_O using ^1^H-NMR spectroscopy and found that the majority of compounds were stable in DMSO-d_6_ and D_2_O over 48 h (except for **4** and **6**, which contain a methyl chain), and slightly decompose in D_2_O after 48 h (Fig. S1‒24). To further assess metabolic stability, two representative compounds (**2a** and **8a**) were incubated with human liver microsomes. The incorporation of a longer linker between the FCZ body and the ferrocenyl moiety in compound **8a** was hypothesized to bring greater metabolic stability over compound **2a**. After 48 h incubation of both compounds with human liver microsomes, samples were analyzed via HPLC. Compound **2a** was shown to undergo 43% metabolism, while compound **8a**, containing the longer linker between ferrocene and the central structure, remained stable throughout (Fig. S25‒35).

### New derivatives have potent activity in *in vitro* antifungal susceptibility screens

The novel compounds were first screened for their antifungal activity against a panel of clinical fungal isolates including FCZ-resistant (MIC_50_ > 100 µM) and mycotoxin-producing fungi (Fig. 2A). A comparison of the novel drug candidates (**5a-12a**) with the previously published compound **2a** (*17*) and the parent drug FCZ revealed that all compounds showed activity against isolates resistant to FCZ. Here, especially the incorporation of either a ferrocenyl (**2a, 7a, 8a**) or a phenyl moiety (**11a**) led to potent antifungal activity, with remarkable activity against all tested isolates highly resistant to FCZ (MIC_50_ > 100 µM). Also, the introduction of a ruthenocenyl (**5a, 6a, 9a, 10a**) or adamantyl moiety (**12a**) conferred high antifungal activity against some FCZ-resistant isolates while showing comparable or weaker (**5a**) antifungal activity compared to FCZ isolates not resistant to FCZ.

**Fig. 2.**
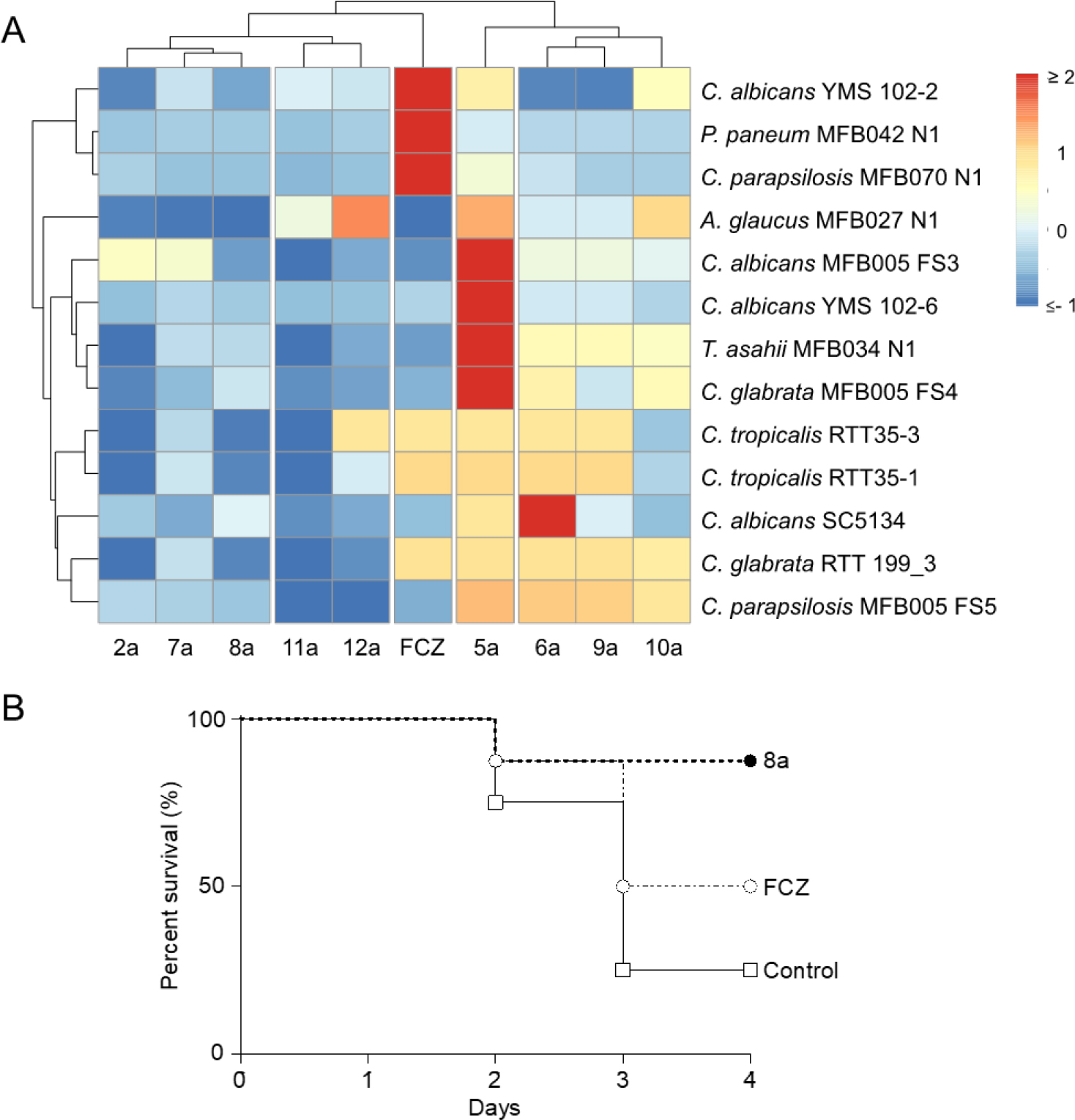
(**A**) Antifungal susceptibility testing on clinical fungal isolates treated with fluconazole (FCZ) and derivatives (**2a**, **5a** – **12a**). MIC50 values (range: 0 µM - 14 µM) have been scaled and centered to a distribution with mean as 0 and standard deviation as 1. (**B**) Mice (8 per group) were infected with FCZ-resistant *C. albicans* CAAL28 and treated immediately with FCZ or **8a** (10 mg/kg) for 3 days *per os*, once daily.

### The 8a compound prolongs mouse survival in an *in vivo* model of murine invasive candidiasis

Since compound **8a** was the most potent analog tested against various clinical fungal isolates *in vitro*, it was also tested in a model of invasive candidiasis in the immunocompromised Balb/c mouse model. After infection with *C. albicans* CAAL28, an FCZ-resistant strain by overexpression of CDR1 and 2 Erg11 mutations (IC_50_ against this strain was > 100 and 13.2 ± 0.8 µM for FCZ and **8a**, respectively), mice were treated for 3 days with 10 mg/kg of FCZ, or compound **8a** *per os*. Survival curves indicated that in contrast to the recognized antifungal drug FCZ (50% survival), compound **8a** significantly (P=0.0247) prolonged mice survival compared to the vehicle control group (survival of 87.5% vs 25%, respectively by day 4) (Fig. 2B).

### FCZ derivatives have potent activity against parasitic nematodes in *in vitro* assays

We next assessed the efficacy of the FCZ and analogs on two groups of phylogenetically diverse species of parasitic nematodes to determine the potential use of these compounds for treatment against a broader range of human pathogens. The lymphatic filarial parasite *Brugia pahangi* and the soil-transmitted gastrointestinal parasite *Trichuris muris,* were selected as representatives of taxonomic groups with very different modes of parasitism (Clade III and I, respectively) (*18*). We hypothesized that any drug with efficacy against both of these nematode species may have broad potential against other parasitic organisms. *B. pahangi* adult females and *T. muris* adults were tested in a whole-worm *in vitro* motility assay (*19–22*) in the presence of the test compounds over 6 days in culture.

Most of the compounds outperformed FCZ in both the *B. pahangi* and *T. muris* assays (Table 1). Particularly, **4a**, **8a**, **10a**, and **12a** were identified as best hits and were then tested to determine the IC_50_s in both species. Overall, the range of activity of the 4 compounds against both species was comparable, and based on IC_50_ values, the compounds were somewhat better against *B. pahangi* (2.2—23.2 µM) compared to *T. muris* (9.0-25.7 µM). The IC_50_ values were lowest for **12a** against the filarial nematode and for **4a** against the whipworm (Table 2).

**Table 1.** *In vitro* whole-worm assays reveal FCZ analogs are potent against both parasite species.

| Compound | % Inhibition of Worm Motility |  |  |  |  |  |  |  |  |  |
| --- | --- | --- | --- | --- | --- | --- | --- | --- | --- | --- |
|  | *Brugia pahangi* females (50 µM) |  |  |  |  | *Trichuris muris* males & females (30 µM) |  |  |  |  |
|  | Day 0 | Day 1 | Day 2 | Day 3 | Day 6 | Day 0 | Day 1 | Day 2 | Day 5 | Day 6 |
| FCZ | 10 | 15 | 7 | 11 | 4 | 6 | 6 | 21 | 20 | 13 |
| 1a | 0 | 0 | 6 | 9 | 78 | 33 | 40 | 44 | 64 | 60 |
| 2a | 0 | 0 | 2 | 0 | 26 | 27 | 16 | 55 | 73 | 75 |
| 3a | 100 | 100 | 3 | 13 | 85 | 16 | 7 | 50 | 66 | 79 |
| 4a | 100 | 99 | 95 | 98 | 100 | 1 | 20 | 32 | 97 | 99 |
| 5a | 11 | 9 | 8 | 4 | 33 | 27 | 43 | 75 | 54 | 61 |
| 6a | 68 | 0 | 0 | 4 | 74 | 23 | 41 | 77 | 82 | 89 |
| 7a | 9 | 11 | 14 | 30 | 100 | 7 | 20 | 22 | 22 | 26 |
| 8a | 100 | 100 | 98 | 99 | 99 | 10 | 77 | 94 | 97 | 79 |
| 9a | 9 | 12 | 9 | 15 | 99 | 16 | 28 | 28 | 0 | 31 |
| 10a | 100 | 99 | 99 | 98 | 99 | 10 | 43 | 73 | 73 | 62 |
| 11a | 32 | 19 | 6 | 8 | 14 | 15 | 15 | 10 | 1 | 0 |
| 12a | 100 | 100 | 100 | 99 | 100 | 10 | 32 | 42 | 98 | 93 |

**Table 2.** IC_50_ of the top 4 hits with *B. pahangi* and *T. muris* on Day 6.

| Compound | *B. pahangi*<br>females<br>IC<sub>50</sub> (μM) | *T. muris*<br>males & females<br>IC<sub>50</sub> (μM) |
| --- | --- | --- |
| **4a** | 23.2 | 9 |
| **8a** | 9 | 23.9 |
| **10a** | 4.6 | 25.7 |
| **12a** | 2.2 | 20.4 |

### *Brugia* female worm fecundity is reduced after treatment with FCZ analogs *in vivo*

With the promising results obtained for *B. pahangi* with **8a**, **10a** and **12a** (IC_50_<10 µM), we next tested these compounds *in vivo*. Male Mongolian jirds infected with *B. pahangi* were given oral doses of **8a**, **10a** or **12a** (10 mg/kg twice a day for 5 days) following reported protocols (*20, 23, 24*). Drug efficacy was evaluated based on: 1) number of adult worms recovered, 2) microfilarial (MF) output to evaluate a female worm’s ability to transmit its progeny, 3) embryogram analyses to determine the effects of the compounds on the developing embryonic stages within the reproductive tract and 4) transmission electron microscopic analyses of the structures in the uteri from recovered female worms. Animals necropsied 5 days after the last dose did not have a significant decrease in worm burden after treatment with **8a**, **10a** nor **12a** (Fig. S36) which suggests that longer-term studies are necessary to observe efficacy of the derivatives to eliminate adult worms. However, female worm fecundity and reproductive output were affected by these derivatives compared to female worms recovered from the vehicle control group (Fig. 3A and Fig. S37). Female worms recovered from animals treated with **8a**, **10a** or **12a** had fewer number of eggs in their reproductive tracts (Fig. 3A and Fig. S37), with **12a** causing significantly lower number of eggs (Fig. S46A) and pre-MF stages (Fig. S37C). Notably, both **10a** and **12a** had significantly higher numbers of deformed embryos (Fig. S37E) and there were significantly fewer numbers of MF released *ex vivo* from female worms removed from animals treated with **12a** compared to the other groups (P < 0.01) (Fig. 3B).

**Fig. 3.**
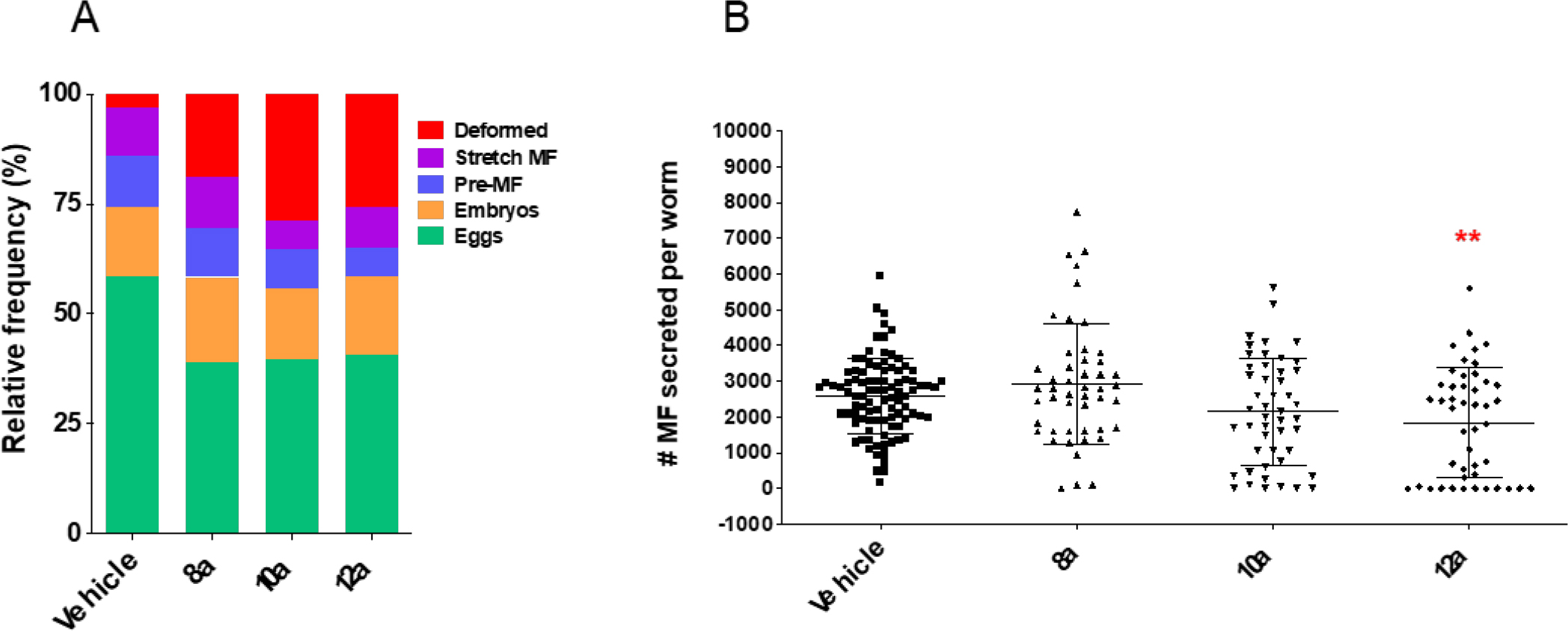
FCZ analogs had a significant effect on the reproductive output and fecundity of female worms recovered from *in vivo* treated infected jirds. (**A**) Embryogram analyses were conducted on female *B. pahangi* worms recovered from control and treated infected jirds (N = 12-36 per group). Bar graphs indicate the relative mean frequency of each of the developmental stages for each treatment group. (**B**) The number of MF released *ex vivo* from the recovered female worms were counted after worms were incubated overnight in media (**P< 0.01).

### Ultrastructural analyses of treated worms support the drug’s effect on the reproductive fitness of *Brugia pahangi* adult female worms recovered from infected jirds after treatment with FCZ analogs

The impact of the FCZ analogs on the embryonic development within the worms’ uteri, from eggs to stretch microfilariae (Fig. 3A), is supported by the ultrastructural analyses of adult female *B. pahangi* recovered from treated infected jirds. The worms collected from animals treated with **8a**, **10a** and **12a** had major tissue damage in their reproductive tracts (Fig. 4). Specifically, female worms from the group treated with **8a** had electron dense, globular material throughout the deformed embryos (Fig. 4G). Notably, disturbances to nuclear ultrastructure were frequently detected in **10a** treated worms (Fig. 4I-L), including several deformed and damaged nuclei within a developing microfilaria (Fig. 4I-L), and an apparent large rupture in the nuclear envelope as well as the appearance of abnormal vacuoles (Fig. 4K). Remarkably, the damage was more profound in worms collected from **12a** treated animals (Fig. 4M-P) as it also showed deformed embryos containing considerable amounts of globular material (Fig. 4M) and punctate, vacuolated spaces adjacent to embryos having deformed and damaged nuclei (Fig. 4N). In addition, many developing embryos also had numerous autophagolysosomes or mature lysosomes (Fig. 4O-P, respectively).

**Fig. 4.**
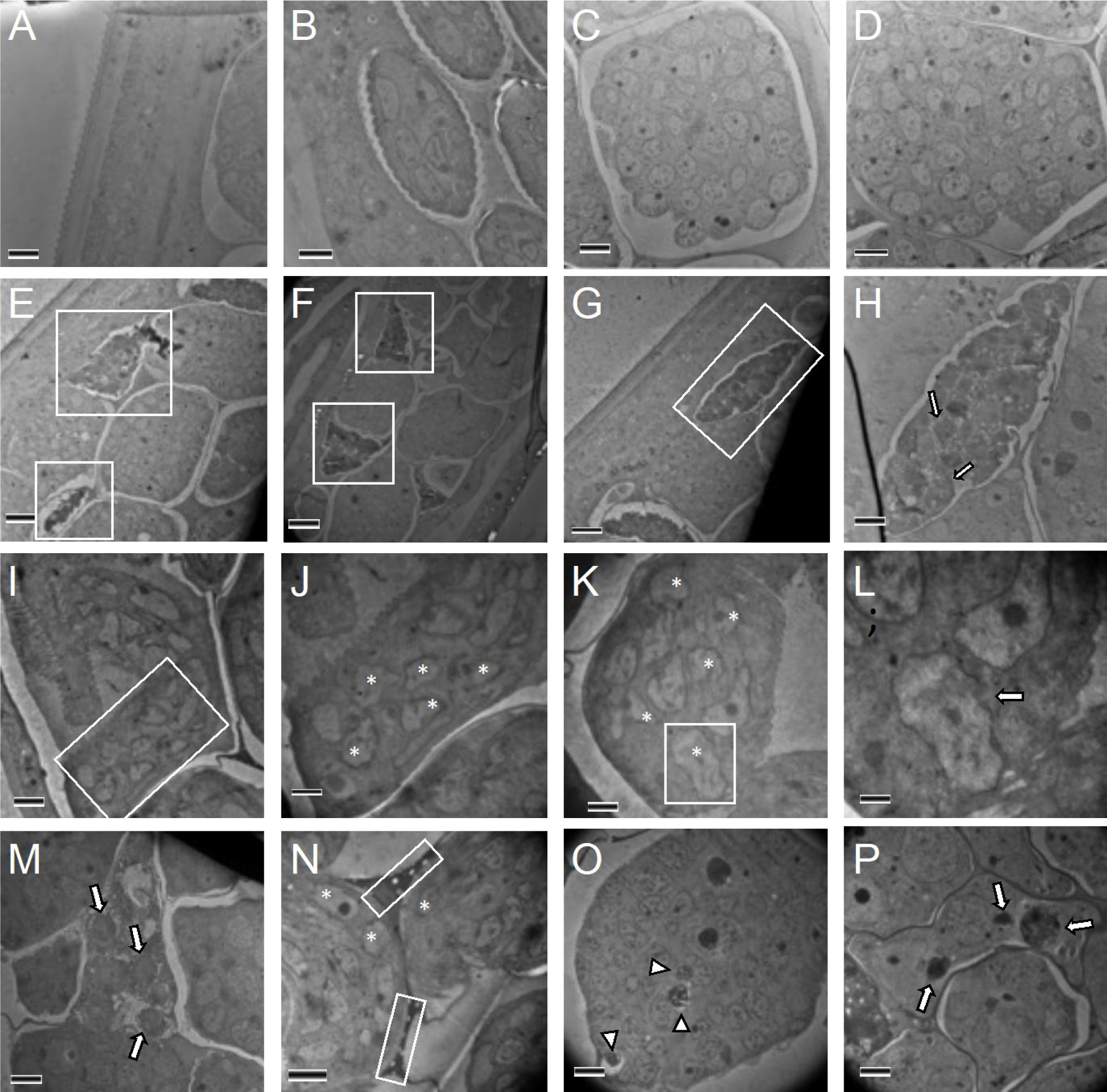
Ultrastructure of embryonic stages within the uteri of female *B. pahangi* worms collected from infected jirds treated with vehicle (A-D), 8a (E-H), 10a (I-L) and 12a (M-P). (**A-D**) Worms from the vehicle treated worms display healthy ultrastructure of the cuticle and varying embryonic stages. (**E** and **F**) Female worms treated with **8a** had both deformed (boxed regions) and healthy embryos. (**G**) A deformed embryo from **8a-**treated worm shown in higher magnification that contains electron dense, globular material (**H**, white arrows) throughout the deformed embryo. (**I-L**) Worms treated with 10a show frequent disturbances to nuclear ultrastructure, including deformed and damaged nuclei (**J-K**, asterisks) within a developing microfilaria. (**L**) A high magnification image of the boxed cell in (**K**) with an apparent large rupture in the nuclear envelope (white arrow). (**M-P**) Diverse effects on the ultrastructure of developing embryos were observed in **12a** treated worms; including (**M**) considerable amounts of globular material (white arrows) within deformed embryos, (**N**) punctate, vacuolated spaces (boxed regions) between developing embryos having deformed and damaged nuclei (asterisks), (**O**) numerous autophagolysosomes (white arrow heads), and (**P**) mature lysosomes (white arrows) within the developing embryos. Scale bars: A-D & H, 2 microns; E-G, 10 microns; I-K & M-P, 3 microns; L, 1 micron.

### Transcriptional profiles of the parasitic nematodes identified pathways and processes affected by treatment

Since FCZ analog **12a** was the most potent compound in the *B. pahangi in vitro* and *in vivo* studies, and second best in the *T. muris in vitro* studies, RNA-seq analyses were conducted on female adult worms treated *in vitro* to further assess the effects of **12a** on both species at the molecular level. To capture early transcriptional responses affected by treatment, we extracted RNA from whole worms exposed at a high concentration (100 µM) for a short period of time (2-3 hours) and then interrogated their transcriptional profiles in comparison to untreated worms. The number of significantly up- and down-regulated genes in **12a** treated *B. pahangi* relative to 1% DMSO controls (Fig. 5A) were 178 (1.2 %) and 96 (0.7 %), respectively. One of the significantly enriched KEGG pathways in the up-regulated *B. pahangi* genes has included chaperones and folding catalysts with 7 out of the 12 enriched genes being ranked in the top 20 most significantly up-regulated genes, and most of them encoded the heat shock protein complex that have orthologs in fungal species (Fig. 5B). The early transcriptomic responses of treated *B. pahangi* genes were less pronounced than those in *T. muris* but they were distinct from those induced in *T. muris*. In **12a** treated *T. muris* worms, 1501 (10.0 %) and 1275 (8.5 %) genes were significantly up- and down-regulated, respectively (Fig. 6A). Interestingly, 9 of the top 10 most significantly up-regulated genes encode proteins involved in mitochondrial respiratory chain complexes and ATP synthesis (Fig. 6A). The functional enrichment analysis indicated that spliceosome, ribosome biogenesis and mitochondria biogenesis pathways were significantly enriched in the up-regulated genes in **12a** treated *T. muris* (Fig. 6C). In addition, we found a significant induction of 7 ABC transporter genes in the **12a** treated worms out of a total of 25 ABC transporters identified in *T. muris* (Fig. 6B and 6D). On the other hand, the expression levels of genes belonging to various signaling pathways (Hippo, mTOR, AMPK, and Wnt), transcription factors, and the fatty acid biosynthesis were significantly decreased (Fig. 6C). Of the 13 genes in the fatty acid biosynthesis pathway, 6 genes including fatty acid synthase, acyl-CoA synthetase, and mitochondrial acyl carrier reductase were significantly decreased in the **12a** treated *T. muris* (Fig. 6E).

**Fig. 5.**
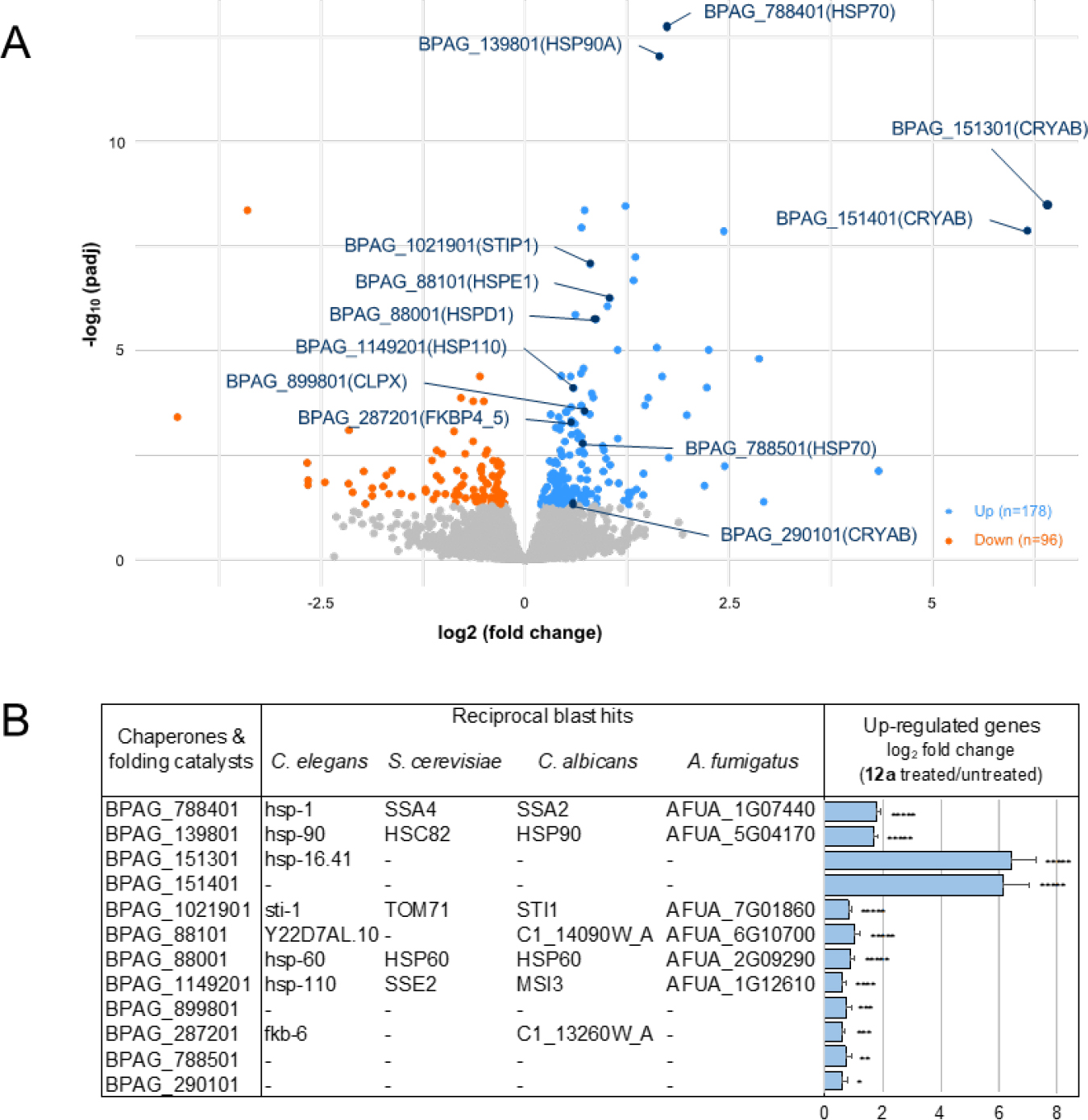
Early transcriptional responses of 12a treated *B. pahangi* adult worms. (**A**) The volcano plot was generated using differentially expressed genes (DEGs) in **12a** treated *B. pahangi* versus DMSO with absolute log2 fold change > 0 and FDR-adjusted p-value (padj) ≤ 0.05 for significance. *B. pahangi* genes related to chaperones and folding catalysts pathway were annotated with KEGG orthology symbol in parentheses. (**B**) 12 *B. pahangi* genes enriched in chaperones and folding catalysts pathway were significantly up-regulated after *in vitro* treatment with **12a**. Sequence-based alignment was performed to identify orthologs of the *B. pahangi* genes to *Caenorhabditis elegans, Saccharomyces cerevisiae, Candida albicans* and *Aspergillus fumigatus* using reciprocal best blast hit search. Log2 fold change values were relative to DMSO control. For significance, FDR-adjusted p-values were represented as defined *padj ≤ 0.05, **padj ≤ 10^-2^, ***padj ≤ 10^-3^, ****padj ≤ 10^-4^, *****padj ≤ 10^-5^.

**Fig. 6.**
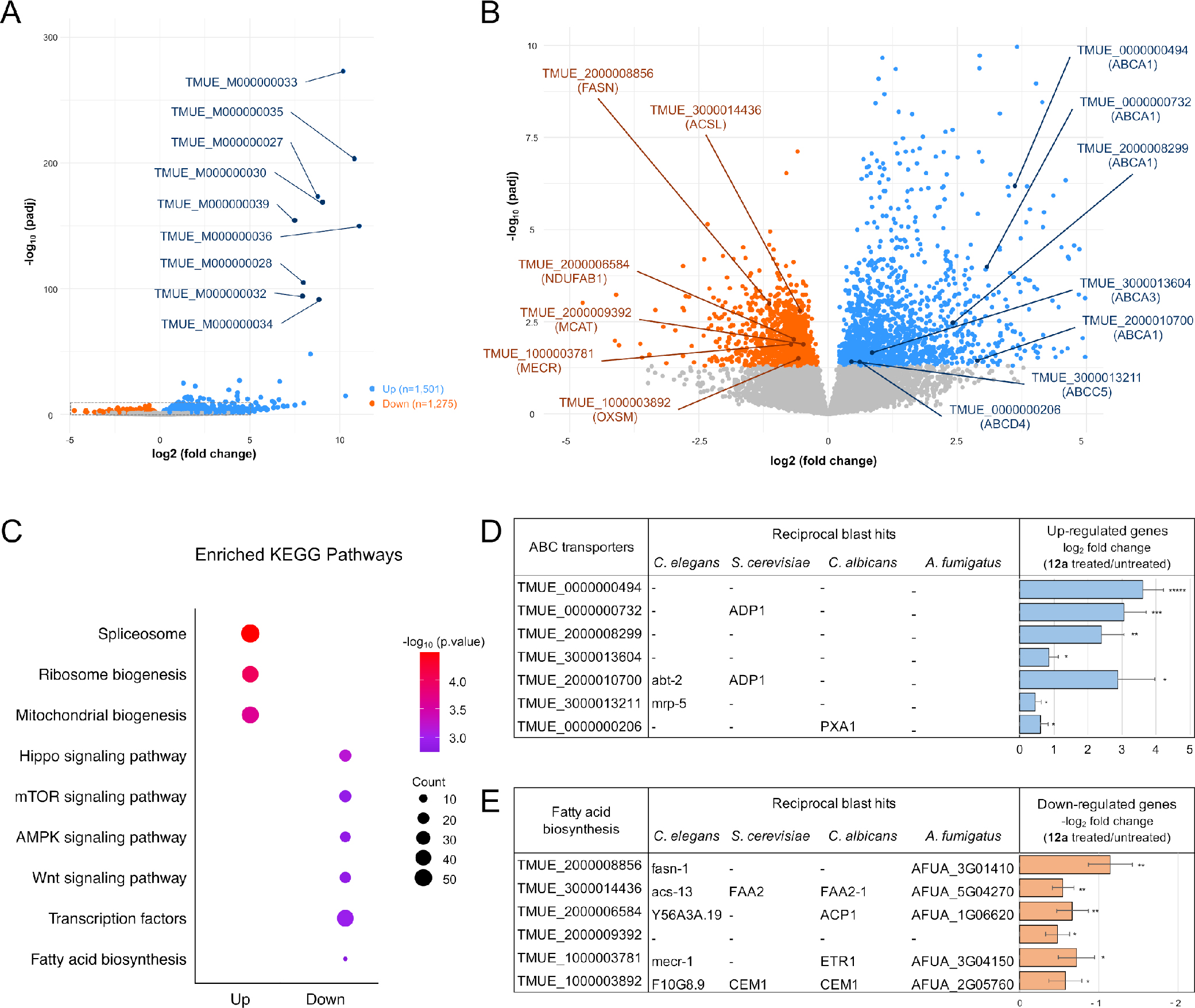
In depth examination of the early transcriptional responses of 12a treated *T. muris* adult worms. (**A**) The volcano plot was generated using DEGs in **12a** treated *T. muris* versus DMSO with absolute log2 fold change > 0 and FDR-adjusted p-value (padj) ≤ 0.05 for significance. The top 9 DEGs associated with mitochondrial biogenesis were highlighted with *T. muris* gene names. (**B**) A dotted box in (**A**) was zoomed in, and *T. muris* DEGs associated with ABC transporters and fatty acid biosynthesis pathways were labeled with KEGG orthology symbols in parentheses. (**C**) Enriched KEGG pathways associated with the various up-regulated and down-regulated genes in **12a** treated *T. muris* worms. The –log10 p-value for each pathway is represented by the color, and the number of significantly DEGs from each pathway is represented by the dot size. (**D**) A summary of the 7 genes annotated as ABC transporters that were significantly up-regulated genes in **12a** treated worms. The orthologous *T. muris* ABC transporter genes in *C. elegans*, *S. cerevisiae*, *C. albicans*, and *A. fumigatus* were determined based on best BLAST reciprocal hits. The Log2 fold change values are relative to DMSO control. For significance, FDR-adjusted p-values were represented as defined *padj ≤ 0.05, **padj ≤ 10^-2^, ***padj ≤ 10^-3^, ****padj ≤ 10^-4^, *****padj ≤ 10^-5^. (**E**) Same as (**D**) but for gens associated with the fatty acid biosynthesis pathway.

### Chemogenomic studies identify new drug targets

Analogues of FCZ that showed elevated activity against fungi and parasitic nematodes (**8a**, **10a**, **11a** and **12a**) were selected to investigate their mechanism of action using a chemogenomic assay. Most interestingly, the genome wide screen revealed that the main targets of the new analogs are different from the main target of FCZ, which is the lanosterol 14α-demethylase encoded by the gene *ERG11*. While the previously synthesized compound **2a**, (*17*) *ERG11* was identified as the main target, the elongation of the linker between the FCZ body and the moiety in the new compounds led surprisingly to a series of new drug targets. These include top targets involved in transcription and translation, such as an acetyl-CoA synthetase with function in histone acetylation (*ACS2*), a snoRNA binding protein (*ENP1*) and a translation initiation factor (*HYP2*), as well as subunits of the RSC-type chromatin remodeling complex (*RSC9*), the RNA polymerase II mediator complex (*SRB2*), the RNA polymerase (*RPC19*) and the small subunit processome complex (*UTP14*) (Table 3 and Fig. S38). Furthermore, identified deletion strains that are highly sensitive to the new compounds were heterozygous for genes involved in calcium homeostasis (*PMC1*), cytoskeleton organization, cell wall deposition and endocytosis (*INP52*), the biosynthesis of branched-chain amino acids (*ILV6*) and in activation of the plasma membrane and segregation of phosphatidylserines (*PMP1*) (Table 3 and Fig. S38).

**Table 3.** The 5 *S. cerevisiae* heterozygous deletion mutants showing highest sensitivity to the compounds **8a**, **10a**, **11a** and **12a** based on the chemogenomic screen. Values are log 2 ratios (control intensity/treatment intensity) calculated as a function of gene (P<0.01).

| Compound | **8a** |  | **10a** |  | **11a** |  | **12a** |  |
| --- | --- | --- | --- | --- | --- | --- | --- | --- |
| Sensitive mutants | *ACS2* | -5.94 | *YGR115C* | -6.34 | *UTP14* | -5.53 | *PMP1* | -6.54 |
|  | *ILV6* | -5.42 | *PMP1* | -6.23 | *YDL144C* | -4.93 | *RSC9* | -6.48 |
|  | *INP52* | -5.32 | *PMC1* | -5.30 | *YPR136C* | -3.12 | *PMC1* | -4.67 |
|  | *SRB2* | -4.42 | *UTP14* | -4.04 | *HYP2* | -2.45 | *YGR115C* | -4.65 |
|  | *YGR122W* | -3.84 | *YDL199C* | -3.83 | *ENP1* | -2.17 | *RPC19* | -4.18 |

We then compared the results for down-regulated genes of the chemogenomic screen of **12a**-treated *S. cerevisiae* with those obtained by RNA-seq for the nematode species *B. pahangi and T. muris* treated by **12a** to determine which pathways and associated functions are affected and identify the ones that are shared among them. Membrane trafficking was the most abundant KEGG category shared by the 3 species, and several of the KEGG categories were related to protein biosynthesis (transcription machinery, transcription factors, spliceosome, and mRNA, tRNA and ribosome biogenesis), modification (chaperones, glycosyltransferases and protein phosphatases) and degradation (ubiquitin system, peptidases, proteasome) (Fig. 7). Together with the KEGG terms DNA repair, chromosome, cytoskeleton and exosome, we can assume a severe impact of **12a** on core biological processes suggesting abnormal cell division and proliferation similar to cancer cells.

**Fig. 7.**
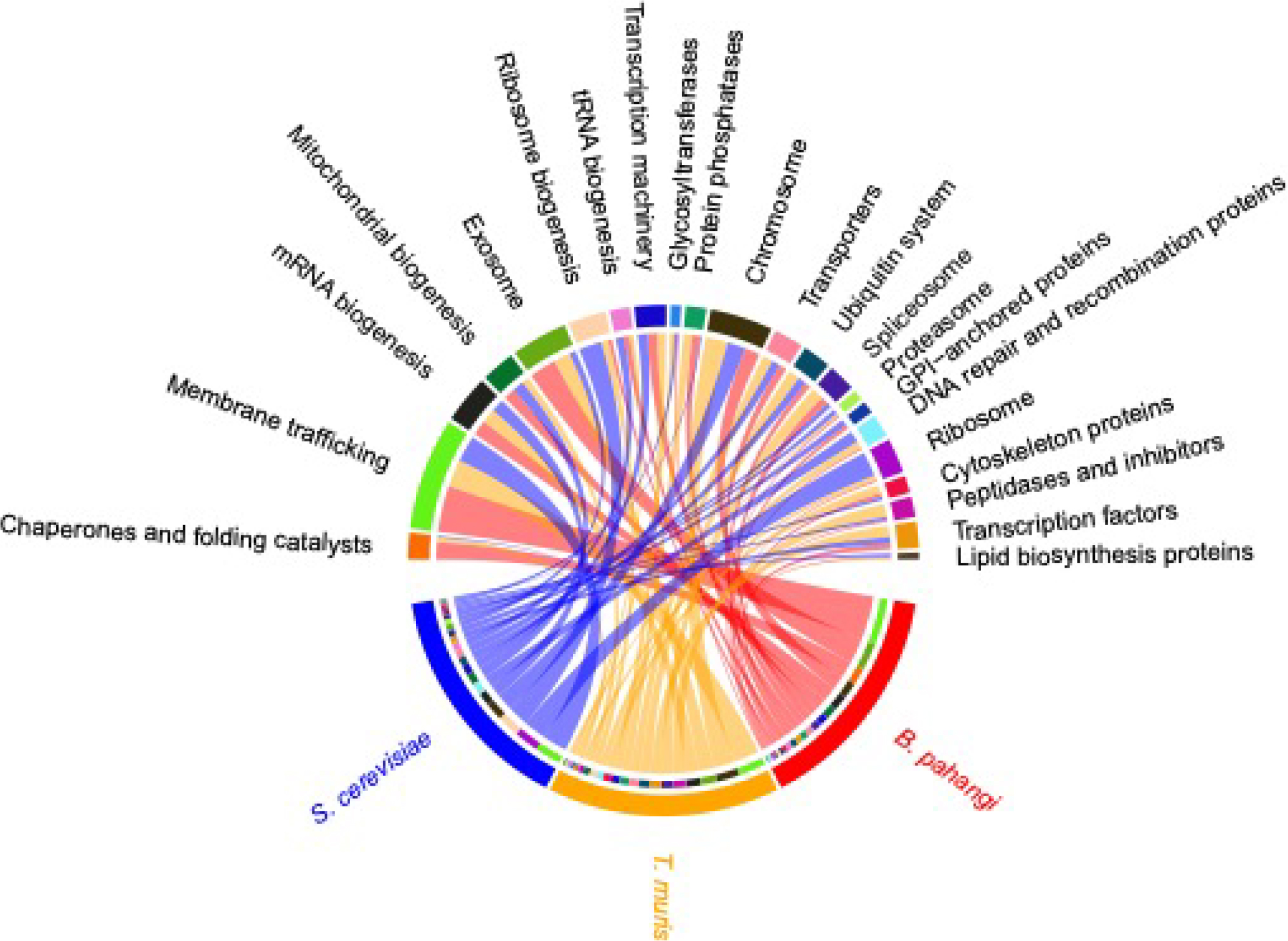
FCZ analog 12a affects similar KEGG pathways among fungal and parasitic nematode species. The relative abundance of the down-regulated genes in three species (*S. cerevisiae*, *B. pahangi* and *T. muris*) are represented in each KEGG category. In order to reduce the total number of KEGG categories, only the ones shared among all three species are depicted.

## DISCUSSION

We report the discovery of novel FCZ derivatives, including organometallic-containing ones, that are uniquely active against not only a panel of clinical fungal isolates, including highly drug-resistant strains, but also on two phylogenetically distant species of parasitic nematodes, *Brugia* (Clade III) and *Trichuris* (Clade I). Significantly, the most potent analog of our *in vitro* antifungal screen (e.g., **8a**) was found to be extremely effective in a model of invasive candidiasis in the immunocompromised Balb/c mouse (i.e., an infection with *C. albicans* CAAL28, an FCZ-resistant strain by overexpression of CDR1 and 2 ERG11 mutations). More surprisingly, chemogenomic studies showed that **2a** and **8a**, which have only a CH_2_ difference in their structures, had different targets. While ERG11 is the main target of **2a**, **8a** has other top targets such as the acetyl-CoA synthetase *ACS2,* required for histone acetylation and the regulatory subunit of the acetolactate synthase *ILV6*. In yeast *ACS2p* is notable because it produces acetyl-coA required for histone acetylation, consequently impacting global gene expression (*25*). Acetolactate synthase (ALS), on the other hand, catalyzes the first steps of branched-chain amino acid biosynthesis (isoleucine, leucine, and valine). Since mammals cannot synthesize branched-chain amino acids, this pathway, and especially the ALS, represents an interesting target for antifungal drugs and other inhibitors such as herbicides (*26*). Furthermore*, C. albicans* ALS mutants showed significant reduced virulence and *in vivo* viability (*27*) which might explain the prolonged survival of mice infected with FCZ-resistant *C. albicans* under treatment with **8a**. The NH**-**(CH_2_-Ferrocene) bond in **2a** was not completely stable, contrary to the NH**-**(CH_2_-CH_2_-Ferrocene) bond in **8a** which may explain this difference in their respective targets.

Resistance to azoles is mainly due to increased drug efflux and alteration of the drug target. Our genome wide screen revealed that the analogues **10a**, **11a** and **12a** also seem to act on cellular mechanisms that are distinct from the parental antifungal, desirable features that may help to decrease tolerance and resistance against FCZ. For example, both blocking Ca^2+^ sequestration accompanied by accumulation of intracellular Ca^2+^ (*PMC1*) and epigenetic alteration of gene expression via chromatin regulation (*RSC9*), can potentiate antifungal activity (*28*). Ribosome biogenesis (*UTP14*) and translation initiation and elongation (*HYP2*), dependent or independent of TOR (target of rapamycin) signalling, are thought to be required for adaptation to azoles (*4, 29, 30*). Moreover, phosphatidylserine secreting membrane proteins (*PMP1*) were discussed as an interesting drug target as phosphatidylserines are among others involved in virulence, mitochondrial function and cell wall thickness of *C. albicans* (*31, 32*). Interestingly, it was shown that the symmetry in the membrane and the exposure of phosphatidylserines as well as their impact on extracellular vesicle formation do not only play an important role in virulence and pathogenicity of pathogenic fungi but also for parasite*s* (*31*).

Since the new FCZ analogs were highly potent against several clinical fungal isolates, we next evaluated the broader application of these compounds on two parasitic helminths, *B. pahangi* and *T. muris*. Both the ferrocenyl analogs **8a** and **10a** were potent against both species of parasitic worms *in vitro* as was the *N*-methyl-β-adamantylethylamine analogue **12a**. In particular, analog **12a** had the greatest effect on the overall reproductive capacity of female *B. pahangi in vivo*, i.e., female worms produced more deformed embryonic stages and released significantly fewer progeny (microfilariae). The structural cellular abnormalities seen in the developing embryos and multifocal damage in the various embryonic stages within the worm’s uterus support the embryogram analyses that revealed reduced fitness of fecundity in these treated worms. Taken together, the *in vivo* data show that the FCZ analogs were highly effective in reducing female worm fecundity and reproductive output. This is an important finding since transmission of the parasites depends upon the female worms’ ability to produce microfilariae that are transmitted to the insect vector, thus perpetuating the life cycle of the parasite. Further *in vivo* studies, however, are needed to optimize the potency and increase the efficacy of these new leads. For example, treatment of infected people with antibiotics that target *Wolbachia*, a mutualistic symbiont of filarial worms, affects worm burden only after 18 months or more post-treatment (*33*). Importantly, the Target Drug Profile for candidate macrofilaricidal drugs includes evidence of impacting adult worms, killing adults or impeding fecundity (*34*). Our study showed that the ferrocenyl and adamantyl analogs are potential new leads for further development not only as antifungals but also as anthelmintics to treat NTDs that can impact the transmission and prevalence of new infections. The FCZ analogs also show promise as a low-cost treatment for developing countries whose communities suffer from years of disabilities from parasitic infections.

By analyzing transcriptional profiles of **12a**-treated *B. pahangi* and *T. muris*, we were able to identify DEGs and molecular pathways disturbed by the *in vitro* treatment. Although the number of DEGs in *B. pahangi* was relatively lower than the ones in *T. muris*, one of the significantly enriched pathways in the up-regulated genes was the chaperones and folding catalysts, and 7 of top 20 up-regulated genes were associated with heat shock protein (HSP) functions. A total 12 *hsp* genes were significantly induced in **12a** treated *B. pahangi* and most of them were highly conserved in other species, from nematodes to yeast: for example, BPAG_139801 (orthologous to HSP90 in *C. albicans*), BPAG_1021901 (STI1), and BPAG_1149201 (MSI3/HSP70). In addition to important functional roles in protein refolding and maturation, the Hsp90-Hsp70 complex interacts with client proteins such as calcineurin via another co-chaperone, a stress-inducible phosphoprotein 1 (STI1) which regulates various cellular signaling pathways (*35–40*). In *C. albicans*, the calcineurin signaling pathway has been shown to confer tolerance to antifungal azoles including fluconazole (*35, 36*). It is possible that the immediate transcriptional induction of chaperones and co-chaperones in *B. pahangi* could be part of an adaptative mechanism responding to stress induced by the FCZ derivative **12a** as well.

In *T. muris*, 9 of the top 10 DEGs that encode proteins involved in mitochondrial respiratory chain complexes and ATP synthesis were significantly up regulated. Mitochondrial functions are directly or indirectly related to efflux pump-mediated azole resistance in various fungal species (*41*). Mitochondria are the major cellular source of ATP, and increased activity of the mitochondrial respiratory metabolism can produce and transport more ATP to drug efflux pumps, enhancing the efflux of intracellular azole drugs (*32*). In addition, defects in mitochondrial functions can affect cytoplasmic Ca^2+^ homeostasis via the calcineurin signaling pathway followed by activation of transcription factors which regulate expression of drug efflux transporter genes (*42–44*). Remarkably, the significant induction of 7 ABC transporter genes in the **12a**-treated *T. muris* in addition to the upregulated genes associated with mitochondrial biogenesis, suggest that although phylogenetically distant from each other, *T. muris* seemed to display similar responses to azole drugs that have been observed in fungal species. On the other hand, fatty acid biosynthesis was one of the significantly enriched pathways that were down-regulated in the **12a**-treated *T. muris*, including fatty acid synthase (TMUE_2000008856). Fatty acids whose synthesis is mediated by fatty acid synthase provide energy during oxidation in mitochondria, peroxisomes and the ER, and are precursors of the building blocks such as sterols, phospholipids, and sphingolipids (*45, 46*). Therefore, the alteration observed may broadly impact important metabolic processes, membrane fluidity/permeability and signaling cascades.

In the *in vitro* motility screening on *B. pahangi* and *T. muris*, FCZ showed no direct effect but the FCZ analogs such as **4a**, **8a**, **10a**, and **12a** effectively inhibited worm motility, which eventually resulted in worm death. Considering the loss of some genes involved in *de novo* cholesterol synthesis in nematode species (unlike fungal species which have Erg11, the target of FCZ), the modifications on the FCZ analogs likely altered other molecular target(s), consequently improving their potencies against the parasitic nematodes as well as FCZ-resistant fungal strains. In consideration of these observations, we interfaced the chemogenomic result from *S. cerevisiae* and the transcriptomic profiles from the two parasitic nematode species to examine affected targets and/or pathways shared among them.

The results suggest that similar pathways are down-regulated when fungal and parasitic nematode species are treated with our new FCZ analogs. In particular, we can assume a severe impact of **12a** on core biological processes suggesting abnormal cell division and proliferation similar to cancer cells; especially because, next to the KEGG categories DNA repair and cytoskeleton, the vast majority of KEGG categories shared by the 3 species are related to protein biosynthesis, modification and degradation. Moreover, membrane trafficking was the most enriched of the shared KEGG categories. Membrane trafficking is a key process for maintaining vital cellular activities by transporting micro- and macromolecules via endo- and exocytosis into, within and out of the cell, including trafficking of pharmacological agents and their intracellular delivery, and is therefore of great interest for drug development (*47*). In this context, GPI-anchors are crucial for the transport of proteins to the plasma membrane and their function, as well as for the endocytosis of cell surface proteins that are recycled in the endosome. GPI-anchored proteins, another shared KEGG category, are often associated with lipid rafts of the membrane that are rich in sterols and sphingolipids (*48, 49*). They are also involved in modifications of the cell wall and have been discussed not only for their role in antifungal susceptibility (*29, 50*) but also for germline development and eggshell formation in nematodes which may explain the increase in number of deformed embryos and ultrastructural damage observed in the developing stages of microfilariae in treated *Brugia* (*48*). Hence, the manifold interactions between lipid biosynthesis, vesicle production, cell wall homeostasis, mitochondrial function and immune activation represent a fascinating area for future studies. Overall, these novel FCZ analogs showed broad-spectrum efficacy across two phylogenetically distinct groups of eukaryotic organisms and may also show promise against other pathogens of medical and veterinary importance in future studies.

## CONCLUSIONS

The search for new anti-infective drugs, notably for the treatment of helminth and fungal infections is a race against the clock, as clearly highlighted by the World Health Organization, that published in October 2022 its first-ever list of health-threatening fungi and its overreaching global targets for control or elimination, including filarial infections, by 2030 (*51*). In this work, we demonstrated that the facile structural modulation of a well-established antifungal drug, fluconazole (FCZ), by some organic but also organometallic moieties, led to the identification of new compounds with not only high activity *in vivo* against a broad range of resistant fungi but also, more surprisingly, the phylogenetically distant parasitic worms such as *Brugia* and *Trichuris,* which cause NTDs that infect millions of the poorest people around the globe. As demonstrated in this work, the newly identified targets for our derivatives are distinctive from the ones of the parent drug FCZ, a vital attribute to circumvent resistance. The discovery of targets that are unique to pathogens, as described herein, is of high value and we are confident that our new family of easy-to-make compounds with broad-spectrum activity offers new hope to treat multiple eukaryotic infectious agents as well as other major human diseases, including newly emerging diseases. The discovery of these advanced hit compounds and the characterization of the mode of action provide the groundwork for highly promising lead optimization in either antifungal or antiparasitic anti-infective development.

## MATERIALS AND METHODS

### Study design

A new series of (metal-containing) FCZ analogs or azoles were prepared and characterized by ^1^H, ^13^CNMR, ^19^F, HRMS, IR and their purity were confirmed by microanalysis. The (metabolic) stability of the compounds were then assessed. This series of compounds was then screened to identify compounds that were active *in vitro* against a panel of *Candida* strains, *B. pahangi* and *T. muris.* After the identification of several promising hits, these compounds were tested on infected mice (invasive candidiasis) and Mongolian jirds (*B. pahangi*). In order to ascertain tissue damage in the reproductive tracts of worms collected, ultrastructural analyses of *B. pahangi* adult females was performed using TEM microscopy. To further assess the effects of the compounds at the molecular level, chemogenomic studies and RNA-seq analyses of *B. pahangi* and *T. muris* adult worms were conducted.

### Chemistry

Compounds **1a-4a** were prepared as previously described (*17*). The synthesis and characterization of compounds **5a-12a** is described in-depth in the supplementary materials.

### *In vitro* antifungal susceptibility assay

Clinical isolates of *Candida* strains (*C. albicans* MFB005FS3, *C. albicans* YMS 102_2, *C. albicans* YMS 102-6, *C. parapsilosis* MFB005FS5, *C. parapsilosis* MFB070 N1, *C. glabrata* MFB005FS4, *C. glabrata* RTT199_3 and *C. tropicalis* RTT35-1, *C. tropicalis* RTT35-3, *Penicillium paneum* MFB042 N1, *Aspergillus glaucus* MFB027 N1, *Trichosporon asahii* MFB034 N1 and the *C. albicans* reference strain SC5134 were tested for their susceptibility to FCZ and the novel compounds. Fungal isolates were grown on Sabouraud agar medium (Sigma Aldrich, St. Louis, MO, USA) for 48 h at 30°C and resuspended in distilled water at a concentration of 1-5 × 105 CFU/ml before testing. Minimal inhibitory concentration (MIC) values for FCZ and derivates were determined following the European Committee for Antimicrobial Susceptibility Testing protocol (EUCAST Definitive Document EDef 7.3.2 Revision, 2020). Briefly, cells were grown in RPMI-1640 medium supplied with 2.0% glucose, counted and inoculated at a concentration of 1-5 × 105 CFU/ml. MIC_50_ values were detected using a spectrophotometer (at 530 nm) after 48 h of incubation, as the lowest concentration of the drug that resulted in a ≥ 50% inhibition of growth, relative to the control.

### *In vivo* studies of murine invasive candidiasis

The compound was tested in a model of invasive candidiasis in the immunocompromised Balb/c mouse (*52*). Mice were immunosuppressed by subcutaneous injection of 15 mg/kg dexamethasone (Dexamedium^®^) one day before challenge. On day 0, mice, 8 per group, were infected intravenously (100 µL) with a suspension of blastopores (3. 10^6^/mL) of *C. albicans* strain, CAAL28, FCZ-resistant by overexpression of CDR1 and 2 Erg11 mutations. One hour after infection, mice were treated *per os* for 3 days once daily with 10 mg/kg of compound **8a** or fluconazole solubilized in distillated water/5% DMSO. The control group received 100 μL of distillated water/5% DMSO. Survival differences in cohorts were analyzed by the log-rank test. A P value of <0.05 was considered statistically significant. Swiss mice (Janvier Labs, Le Genest-Saint-Isle, France) with a body weight of ∼25 g were obtained and allowed to acclimate for 7 days prior to use. Environmental controls for the animal room were set to maintain a temperature of 16 to 22°C, a relative humidity of 30 to 70%, and a 12:12 hourly light-dark cycle. All the experimental protocols were prior approved by the Ethic Committee on Animal Testing *(comité d’éthique en expérimentation animale N°006)* and was authorized by the French Ministry of Higher Education, Research and Innovation (APAFIS#29252-2021011815391956 v4).

### *In vitro* motility assays with adult female *Brugia pahangi* and adult *Trichuris muris*

The efficacy of the FCZ derivatives were tested on adult female *B. pahangi* and *T. muris* in *in vitro* inhibition assays (*19–22*). Briefly, each well in a 24-well plate contained one adult female *B. pahangi* (4 replicate wells per compound) and 500 µL of complete media (RPMI-1640 with 25 mM HEPES, 2.0 g/L NaHCO_3_, 5% heat inactivated FBS, and 1X antibiotic/antimycotic solution), and the plates were incubated for 6 days at 37°C with 5% CO_2_. The motility of the worms was measured using the Worminator instrument on days 0, 1, 2, 3, and 6 (*19*). Each plate of worms was recorded for 60 s to determine the number of pixels displaced by the worms per second. The percentage of inhibition of motility was calculated by dividing the mean movement units (MMUs) of treated worms by the average MMUs of the control worms (treated with 1% DMSO), then subtracting that value from 1. The values were floored to zero and multiplied by 100%. IC_50_ assays were conducted for **4a**, **8a**, **10a** and **12a** using 6-point dilutions and were calculated using Prism v6.0. Adult *T. muris* were assayed in the same manner as *B. pahangi* except each well contained 2 worms in culture media (4 replicates wells per compound).

### Animal infections and dosing for *Brugia pahangi/jird* model of infection

Animal studies were approved under the New York Blood Center (NYBC) Institutional Animal Care and Use Committee Protocol 369 and adhered to the guidelines set forth in the NIH Guide for the Care and Use of Laboratory Animals and the USDA Animal Care Policies. Mongolian jirds (*Meriones unguiculatus*), 5-7 week old male weighing 50–60 grams, were injected intraperitoneally with 200 *B. pahangi* third-stage larvae and then shipped to the NYBC from the NIH/NIAID Filariasis Research Reagent Resource Center, Athens, GA. Five months post-infection, when larvae have matured into fertile adult worms, animals were given oral doses of 10 mg/kg of **8a**, **10a** and **12a** twice a day for 5 days. Compounds were formulated with 5% glucose and the vehicle group received 5% glucose only. Five days after the last dose, animals (6 animals per treatment group and 9 animals in the vehicle group) were euthanized following NIH Animal Guidelines (NYBC) and necropsies were performed.

### *B. pahangi* microfilarial output and embryogram analyses

Individual adult *B. pahangi* female worms (n = 12–24 females per group) from each group were collected during necropsies and maintained overnight in 24-well plates with 2 mL of RPMI-1640 supplemented with 1X GlutaMAX, 1X Antibiotic/Antimycotic (Life Technologies, Grand Island, NY), and 10% heat inactivated-FBS (R&D Systems, Flowery Branch, Georgia).. The MF released *ex vivo* from individual females after 18 hours in culture were counted using a dissecting microscope. For the embryogram analyses, individual adult female worms were finely chopped in 1 mL of PBS (Life Technologies, Grand Island, NY) to disrupt the cuticle and release the reproductive content. The various developing embryonic stages released from the two gonads (eggs, embryos, pre-MF, and stretched MF, deformed embryos) were assessed in a blinded fashion for their developmental stages using an inverted microscope and hemocytometer. A minimum of 200 developmental stages were counted from each female and the relative proportions of each stage as well as the % of each developmental stage per female worm in 6– 9 worms per treatment group were calculated (*24, 53*). Release of MF *ex vivo* were analyzed using a Kruskal–Wallis test followed by Dunn’s multiple comparison test. For the embryogram analyses, a two-way ANOVA was conducted with a Dunnett’s multiple comparisons test where significance levels were determined based on comparisons with worms from vehicle treated animals. All statistical analyses were determined using Prism 8 version 8.2.0.

### Transmission electron microscopy of female *Brugia*

*B. pahangi* adult females from necropsy were fixed in 2.5% glutaraldehyde and 2% paraformaldehyde buffered with 0.1 M sodium cacodylate, worms were cut into small pieces while in the fixative and stored at 4°C until processed as described (*24*). Samples were washed with 0.1 M sodium cacodylate, and a secondary fixation was done with 1% osmium tetroxide for 1 hour. Following another wash with 0.1 M sodium cacodylate, samples were dehydrated in increasing ethanol concentrations (30-100%) and washed two times with propylene oxide for complete desiccation. Dehydrated samples were embedded in BEEM capsules using Spur low viscosity embedding kit (Electron Microscopy Sciences). Blocks were polymerized at 70°C for 8 hours. Ultrathin sections were collected on nickel formvar/carbon coated 100 mesh grids and contrasted with Uranyless (Electron Microscopy Sciences) and lead citrate. Sections were imaged on a Tecnai G2 Spirit TEM equipped with an AMT camera.

### RNA-seq of *Brugia pahangi* and *Trichuris muris* adult worms treated *in vitro* with 12a

Adult female *B. pahangi* and adult female *T. muris* were treated with 100 µM of 12a and 1% DMSO (vehicle control) for 2-3 hours *in vitro* and stored at – 80 °C. Four replicates of *B. pahangi* (2-3 individual worms per sample) and *T. muris* (1 worm per sample) were then subjected to RNA extraction. Worms were homogenized using 1.5 mm zirconium beads (Benchmark scientific, D1032-15) and BeadBug6 (Benchmark scientific, D1036, Sayre-ville, NJ, USA) and total RNA was isolated using Trizol (Invitrogen, Waltham, MA, USA). cDNA libraries were prepared from RNA samples with poly(A) enrichment and the processed cDNA was sequenced on the Illumina NovaSeq S4 platform (paired-end 150bp reads).

### RNA-seq analysis and functional enrichment tests

Sequencing produced an average of 42 million reads per sample. An average of 93.1 % and 94.9 % of reads were mapped to the *B. pahangi* genome assembly (PRJEB497, WBPS13) and *T. muris* (PRJEB126, WBPS16) respectively using STAR and quantified per gene using featureCounts. Relative gene expression was calculated using “fragments per kilobase per million reads” (FPKM) with normalization per gene per sample. Fold changes and differential expression significance values (relative to DMSO controls) were calculated using DESeq2 (version 1.34.0; default settings, FDR ≤ 0.05). Functional annotations for all *B. pahangi* and *T. muris* genes were assigned using annotations from InterProScan v5.42 to identify gene ontology classifications and InterPro functional domains, and GhostKOALA v2.2 to identify KEGG annotations. Enriched KEGG pathways among differentially expressed gene (DEG) sets were identified using WebGestaltR (minimum 2 genes per pathway, FDR ≤ 0.05 threshold for significance).

### Protein sequence-based comparison to identify orthologous genes

Protein sequence data were downloaded from WormBase Parasite: *B. pahangi* (PRJEB497), *T. muris* (PRJEB126) and *Caenorhabditis elegans* (WBcel235). In addition, for protein sequence comparison, yeast genomes were also downloaded from Ensembl release 55: *S. cerevisiae* (R64-1-1), *C. albicans* (GCA000182965v3) and *A. fumigatus* (Af293). Best bi-directional hits were identified based on protein sequence alignment of the longest isoforms per gene of parasitic nematodes (*B. pahangi* and *T. muris*, each) against 4 comparative species including the 3 yeast species (*S. cerevisiae*, *C. albicans*, and *A. fumigatus*) and *C. elegans* using Diamond blastp (v2.0.6.144). This alignment was performed twice for each pairwise combination of the two parasitic nematodes and the 4 comparative species. Once with the parasitic nematode being the query and the comparative species being the subject, and again with the comparative species being the query and the parasitic nematode being the subject. Cases in which a pair of proteins were each other’s top scoring hit were flagged as being reciprocal hits. Gene symbols and descriptions were harvested from the BioMart instance hosted at WormBase Parasite and were used to annotate the results.

### Chemogenomic Studies

Frozen aliquots of Yeast Deletion Heterozygous Diploid Pools (Cat. no.95401.H4Pool) were recovered for 10 generations and logarithmically growing cells were diluted to a final OD600 of 0.06 (=10^5 cells/ml) in YPD containing 1% DMSO or compound. The compound was applied at a dose of 1 µM corresponding to 10%–20% wild-type growth inhibition. Cells were harvested after 20 generations of growth and frozen at -20°C for subsequent preparation of DNA. Genomic DNA was purified according to the Qiagen supplementary protocol ‘Purification of total DNA from yeast using the DNeasy® Blood &Tissue Kit’ (Qiagen). DNA quality was assessed via agarose gel electrophoresis and UV-Vis spectroscopy. For each compound three biological replicates were generated.

A two-step PCR protocol for efficient multiplexing of Bar-seq libraries was applied as previously described with the following modifications. In a first step, UPTAGs and DNTAGs from a single sample were amplified using the primers Illumina UPTAG Index and Illumina UPkanMX and Illumina DNTAG Index and Illumina DNkanMX in separate PCR reactions. Illumina UPTAG and Illumina DNTAG primers contain a 5-bp sequence that uniquely identifies the sample. A complete list of primer sequences is provided in Table Sxx. Genomic DNA was normalized to 20ng/ml and 100ng were used as template for amplification of barcodes using KAPA Hifi HotStart Ready Mix (Roche), applying the following PCR program: 4 min at 95°C followed by 25 cycles of 10 sec at 95°C, 10 sec at 50°C, 10 sec at 72°C, and a final extension step of 2 min at 72°C. PCR products were confirmed on 2% agarose gels and purified using the NucleoSpin Gel and PCR clean-up kit (Macherey-Nagel). Purified PCR products were quantified using the Quant-iT dsDNA kit (Invitrogen) and 60ng from each of the 15 different UPTAG libraries and, in a separate tube, 60 ng from each of the 15 different DNTAG libraries were combined. The multiplexed UPTAG libraries were then amplified using the primers P5 and Illumina UPkanMX, and the combined DNTAG libraries were amplified using the P5 and IlluminaDNkanMX primers (Table Sx) and the PCR program: 4 min at 95°C followed by 20 cycles of 10 sec at 95°C, 10 sec at 50°C, 10 sec at 72°C, and a final extension step of 4 min at 72°C. The 140-bp UPTAG and DNTAG libraries were purified using the NucleoSpin Gel and PCR clean-up kit (Macherey-Nagel), quantified using the Qubit dsDNA HS kit (ThermoFisher Scientific) and the libraries were sequenced separately on an Illumina NextSeq 2000 using standard methods, including the use of the standard Illumina sequencing primer (5’-ACA CTC TTT CCC TAC ACG ACG CTC TTC CGA TCT-3’). The qseq files are available from the European Nucleotide Archive (ENA) with the accession number PRJEB57349. Read matching was performed using an in-house developed Python script (https://github.com/marcomoretto/Bar-seq) as previously described (*54*). The statistical analysis was performed with the voom (*55*) transformation package that estimates the mean–variance relationship of log counts, generating a precision weight for each observation that is fed into the limma (*56*) empirical Bayes analysis pipeline. A double threshold based on both p-value (≤0.01) and expression log_2_ fold change (≥1.5) was imposed to identify barcodes differentially abundant through pairwise comparison. The Fig. 7 has been obtained using the R language and the package circlize 0.4.15. DEGs from the three species (*S. cerevisiae*, *T. muris* and *B. pahangi*) obtained as described above, have been annotated using KEGG BRITE database (https://www.genome.jp/kegg/brite.html). The relative abundance of the species DEGs are represented in each KEGG category, for example, for all the genes in the three species annotated as membrane trafficking ∼30% are coming from *S. cerevisiae*, ∼30% from *T. muris* and ∼40% from *B. pahangi*. In order to reduce the total number of KEGG categories, only the ones shared among all three species have been retained.

## Supplementary Materials

This PDF file includes:

Fig. S1-S38.

Cytotoxicity results at 25,50 and 100µM on RPE-1 cells (Figure S39)

List of primer sequences (Table S1)

Complete synthetic route (Figure S40), procedures and characterization of the compounds, copies of the IR (Figure S41‒S48) and NMR (S49‒S121) spectra.

## Acknowledgments

We thank Dr. Andy Moorhead and Katie Greenway and the NIH/NIAID Filariasis Research Reagent Resource Center (www.filariasiscenter.org) for providing us with live adult female *Brugia pahangi* worms and the infected gerbils.

## Funding

This work was financially supported by: the Swiss National Science Foundation to G.G. (Grant Sinergia CRSII5_173718); an ERC Consolidator Grant PhotoMedMet to G.G. (GA 681679) received support under the program *Investissements d’Avenir* launched by the French Government and implemented by the ANR with the reference ANR-10-IDEX-0001-02 PSL (to G.G.) and NR-l 7-CONV-0005 (bourse Qlife, to C.O. and G.G.); the University of California, San Francisco Quantitative Biosciences Institute (QBI)-Curie/PSL for Quantitative Approaches for Studying Complex Biological Phenomena Project to J.A.S.; NIH grants AI59450 (to M.M.), EY033195 (to M.M. and S.L.) and AI153649 (to S.L.). This research was also supported by the Autonomous Province of Trento (Accordo di Programma P2211003I & P1611051I to T.W. and M.M), and the China Scholarship Council’s financial support to Y.L.

## Author Contributions

T.W., S.L., K.C, M.Mi., J.A.S. and G.G. conceived the project. Y.L., J.N. and C.O. prepared and characterized the compounds. T.W. and M.Mo. performed chemogenomics studies. C.A.B. and H.J. performed *in vitro* experiments on *T. muris* and *B. pahangi*. R.V. performed cytotoxic studies on non-cancerous and cancerous cells. N.T., R.P., M.S.M., J.A.S, and S.L. performed *in vivo* studies on *B. pahangi.* T.W. performed *in vitro* studies on fungi. F.P. and P.L.P. performed *in vivo* studies on fungi. R.L. and J.F.U. maintained the *T. muris* life cycle and provided worms. T.W., S.L., K.C, M.Mi., J.A.S. and G.G. wrote the manuscript. All authors edited the manuscript

## Competing Interests

The authors declare that they have no competing interests.

## Data and materials Availability

All data associated with this study are present in the paper or the Supplementary Materials.

## Notes

### Competing Interest Statement

The authors have declared no competing interest.

