## Supplementary Information for "Discovery of New Broad-Spectrum Anti-Infectives for Eukaryotic Pathogens Using Bioorganometallic Chemistry"

### 1. $^{1}$H NMR spectra for stability

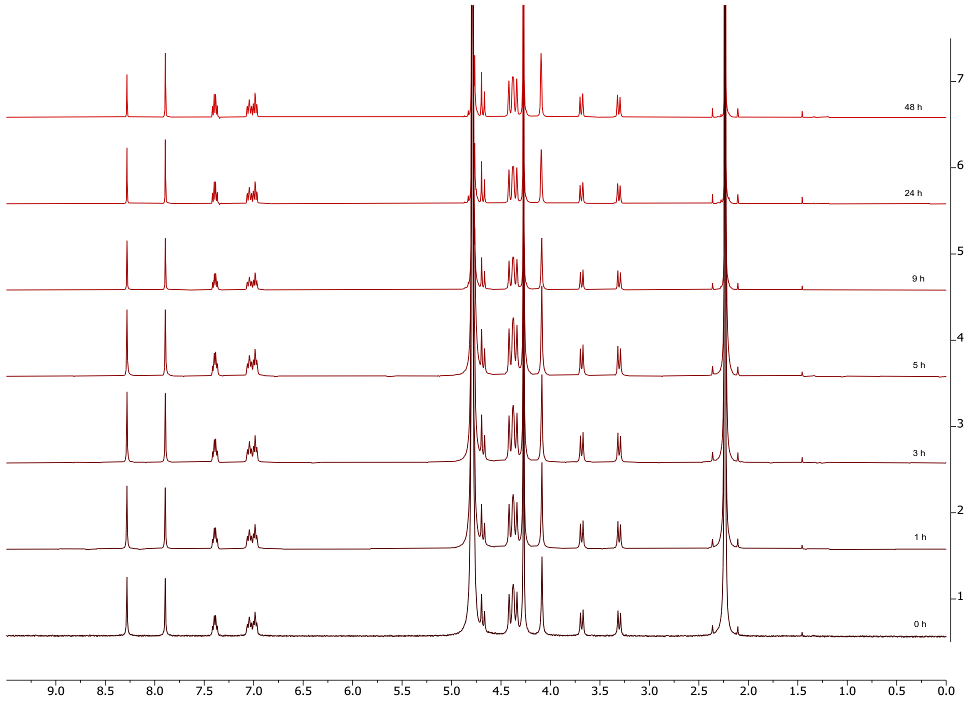

**Figure S1.** Stability of **1a** in $D_2O$ up to two days.

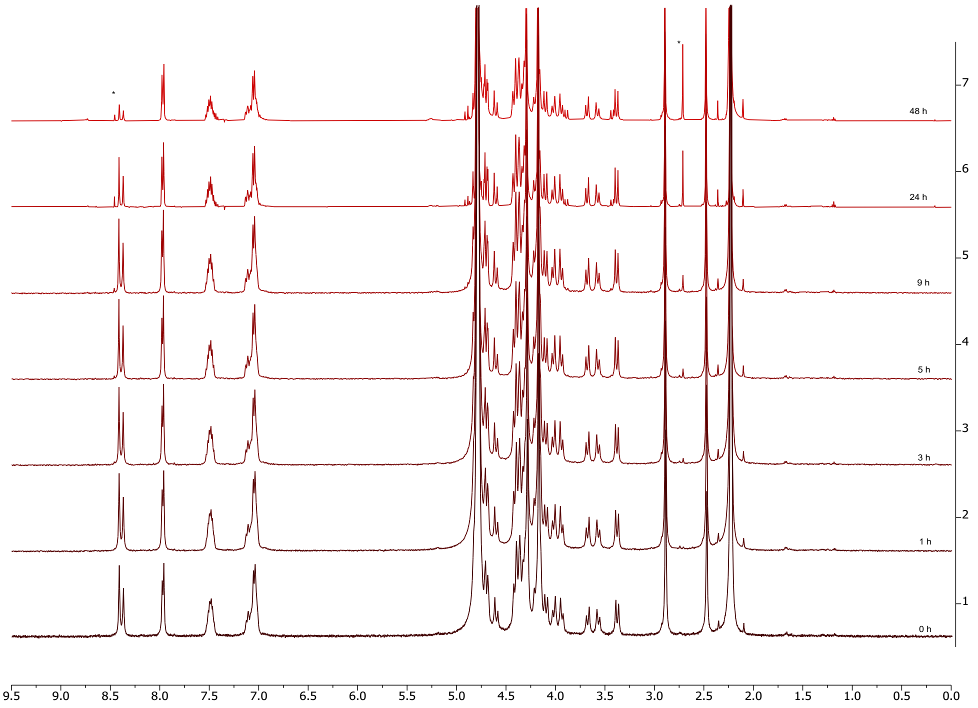

**Figure S2.** Stability of **2a** in $D_2O$ up to two days.

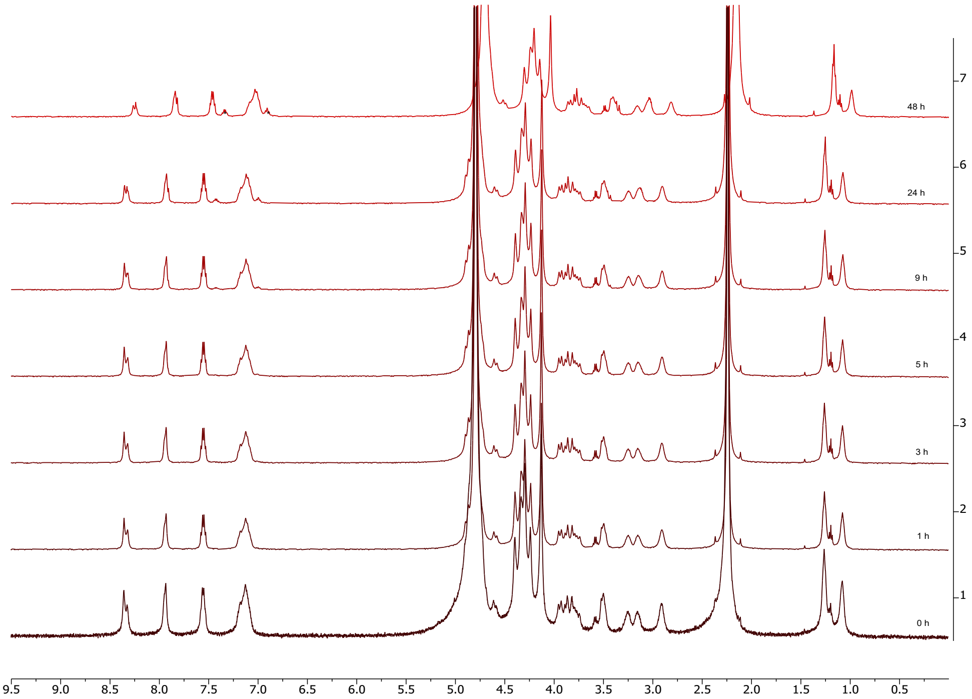

**Figure S3.** Stability of **3a** in $D_2O$ up to two days.

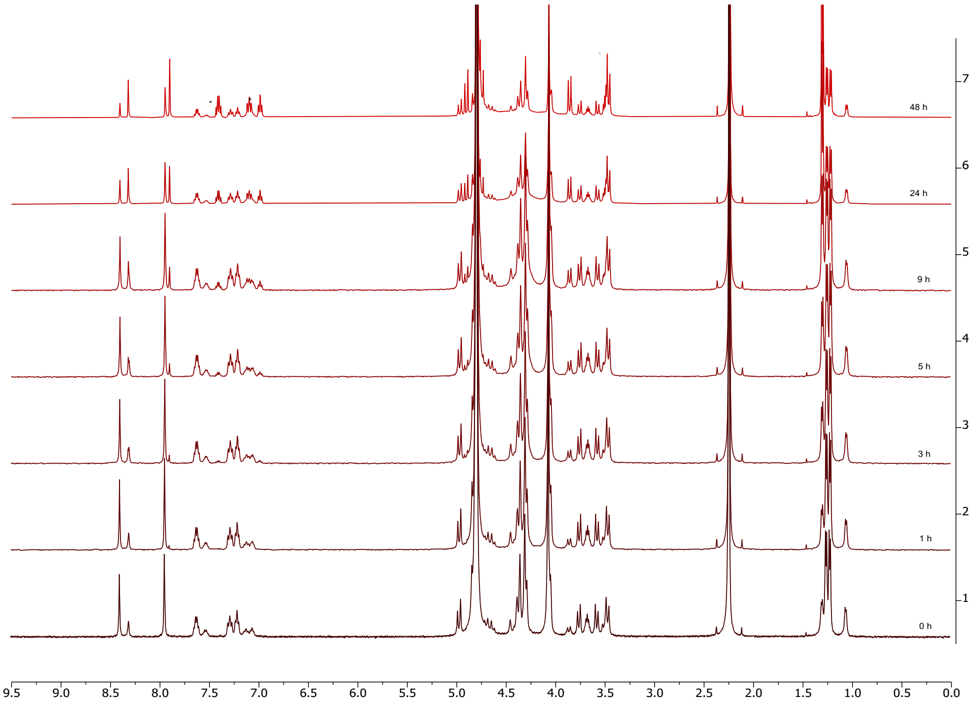

**Figure S4.** Stability of **4a** in $D_2O$ up to two days.

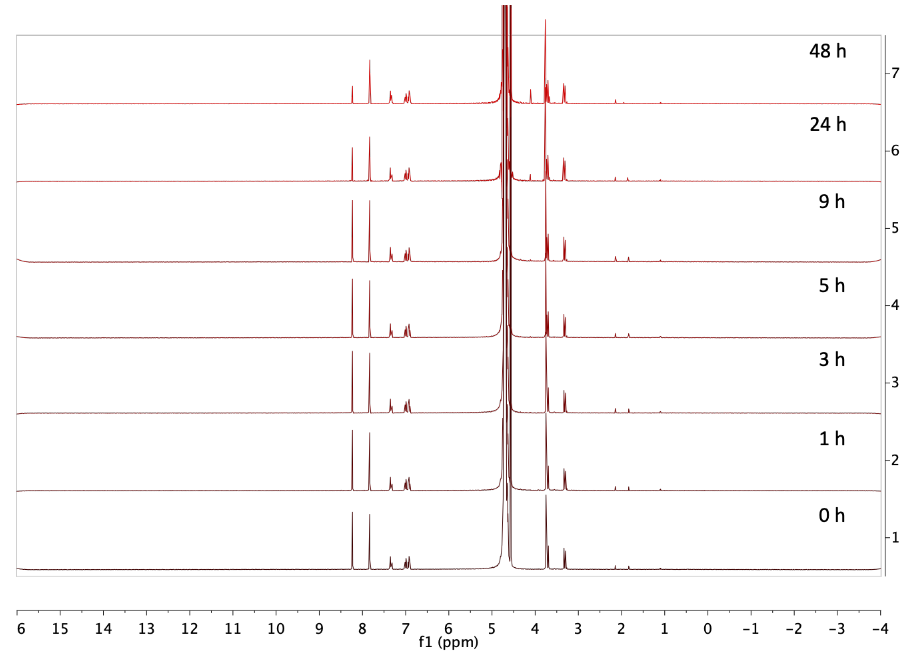

**Figure S5.** Stability of **5a** in $D_2O$ up to two days.

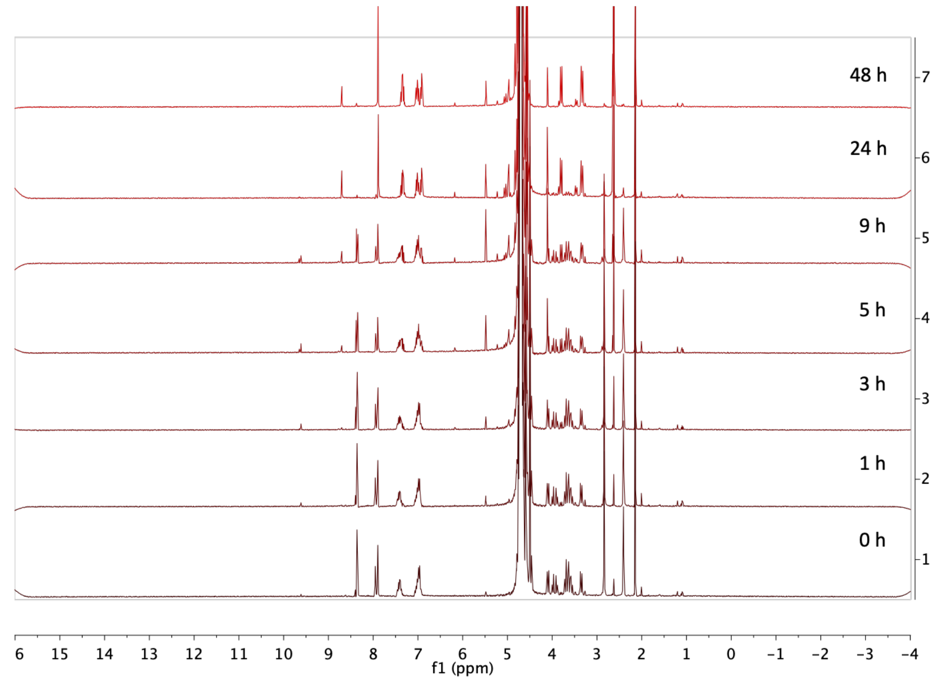

**Figure S6.** Stability of **6a** in $D_2O$ up to two days.

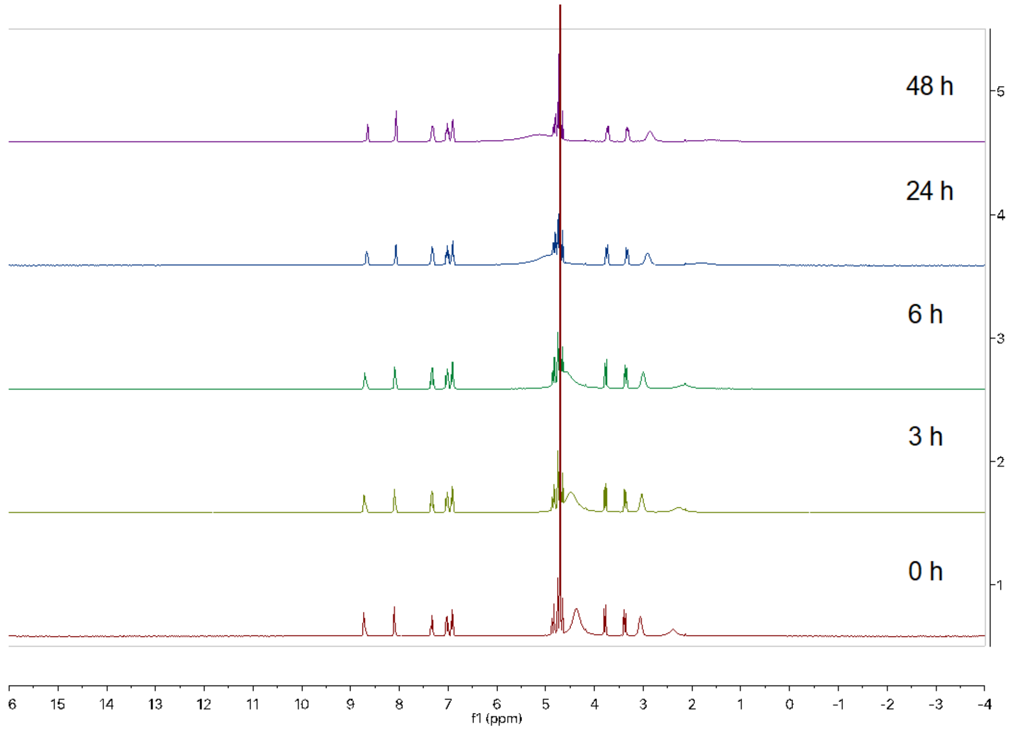

**Figure S7.** Stability of **7a** in $D_2O$ up to two days.

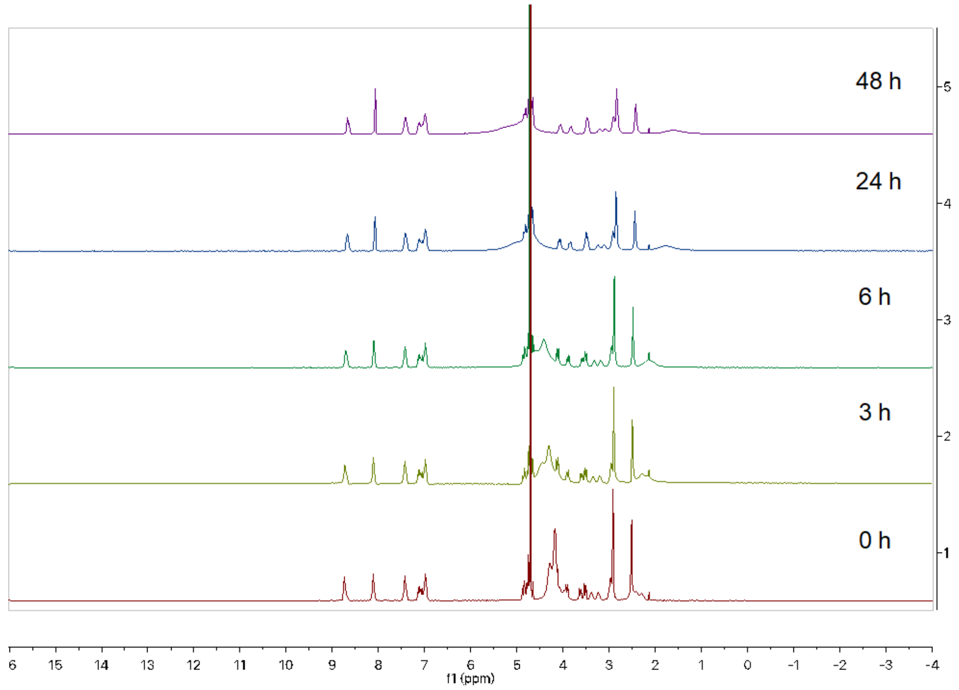

**Figure S8.** Stability of **8a** in $D_2O$ up to two days.

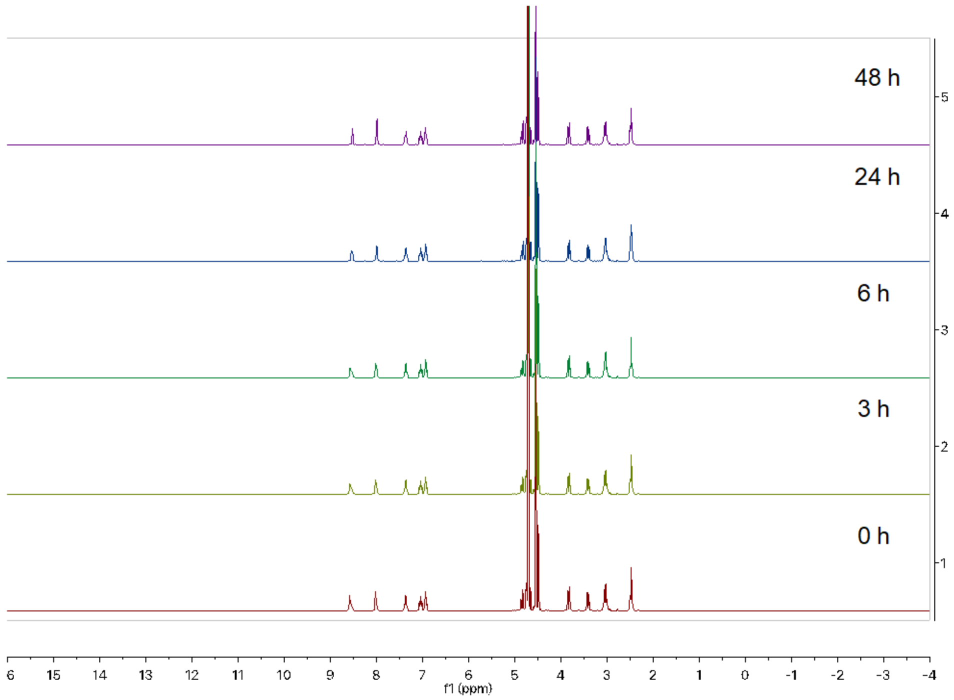

**Figure S9.** Stability of **9a** in $D_2O$ up to two days.

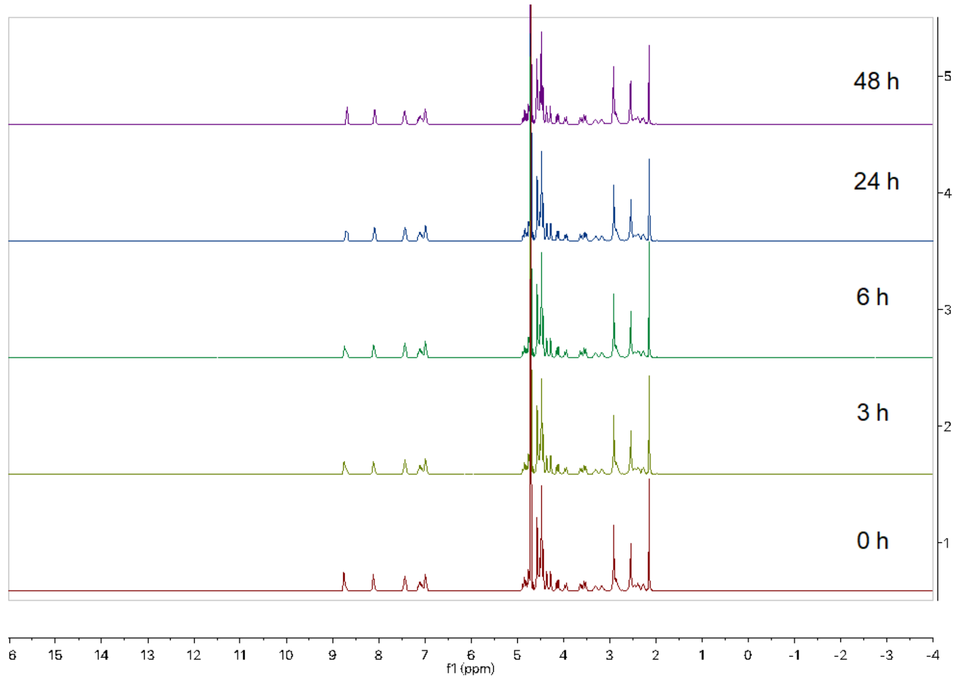

**Figure S10.** Stability of **10a** in $D_2O$ up to two days.

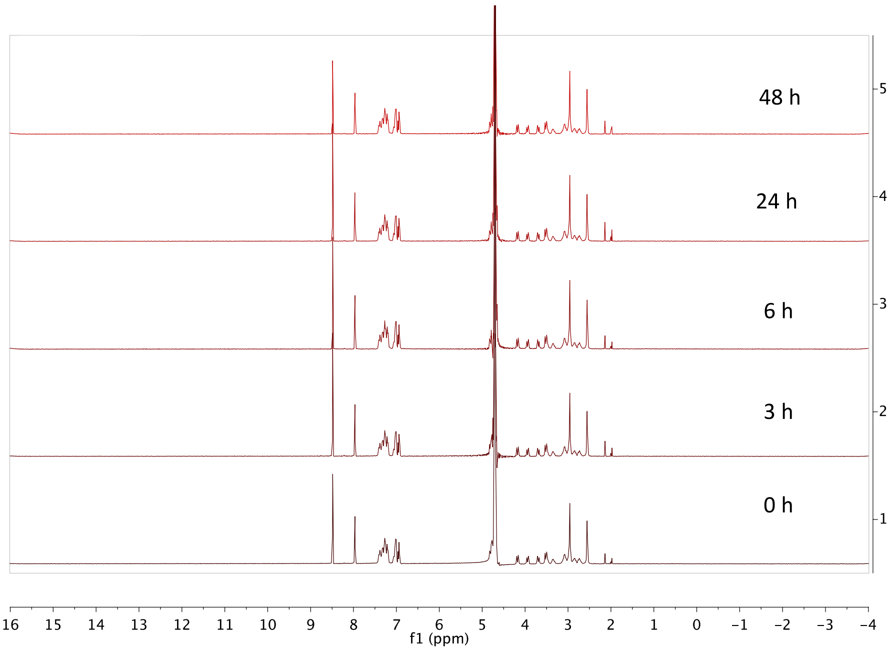

**Figure S11.** Stability of **11a** in $D_2O$ up to two days.

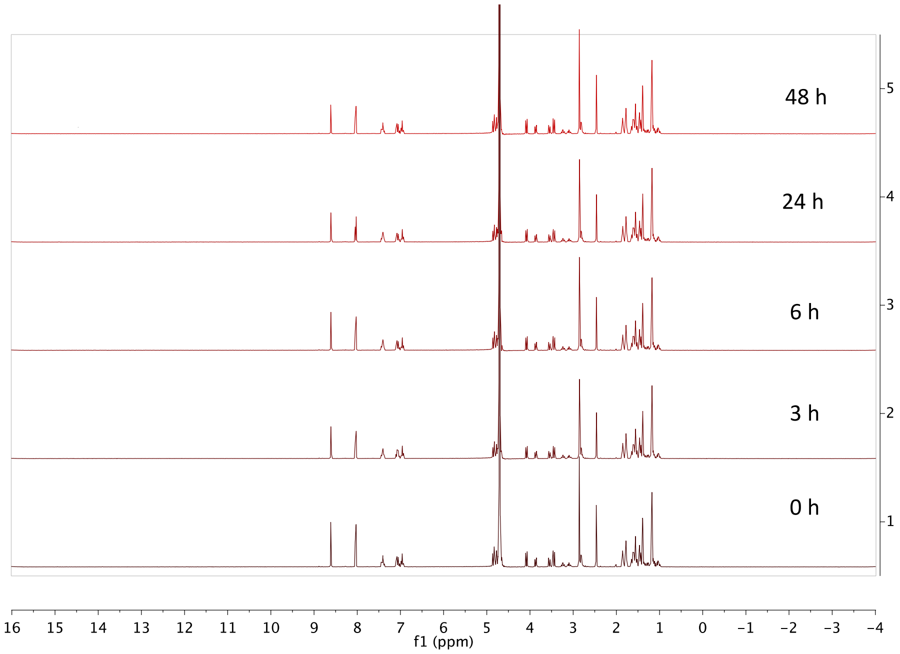

**Figure S12.** Stability of **12a** in $D_2O$ up to two days.

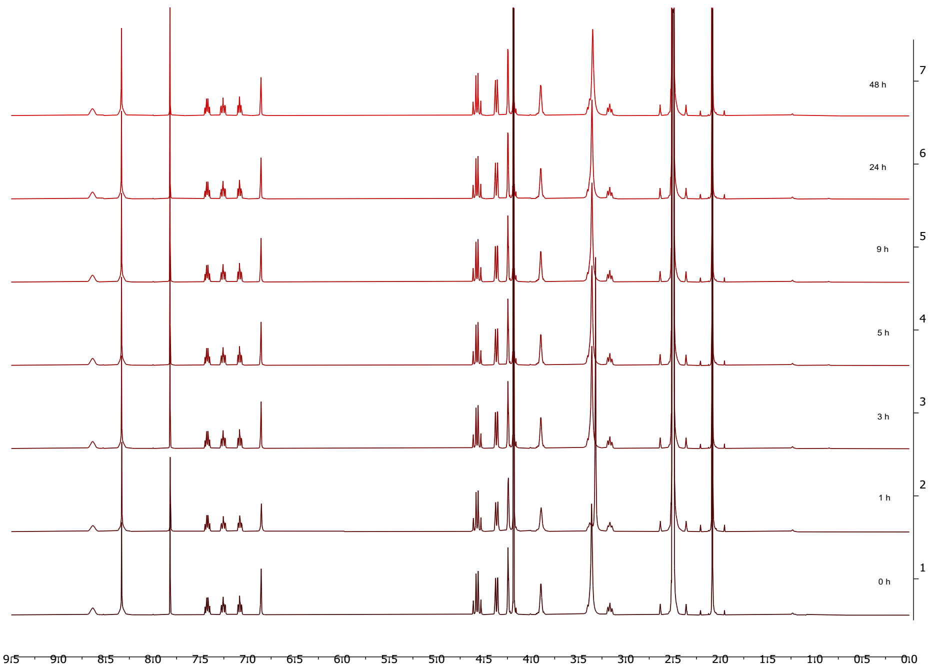

**Figure S13.** Stability of **1a** in DMSO up to two days.

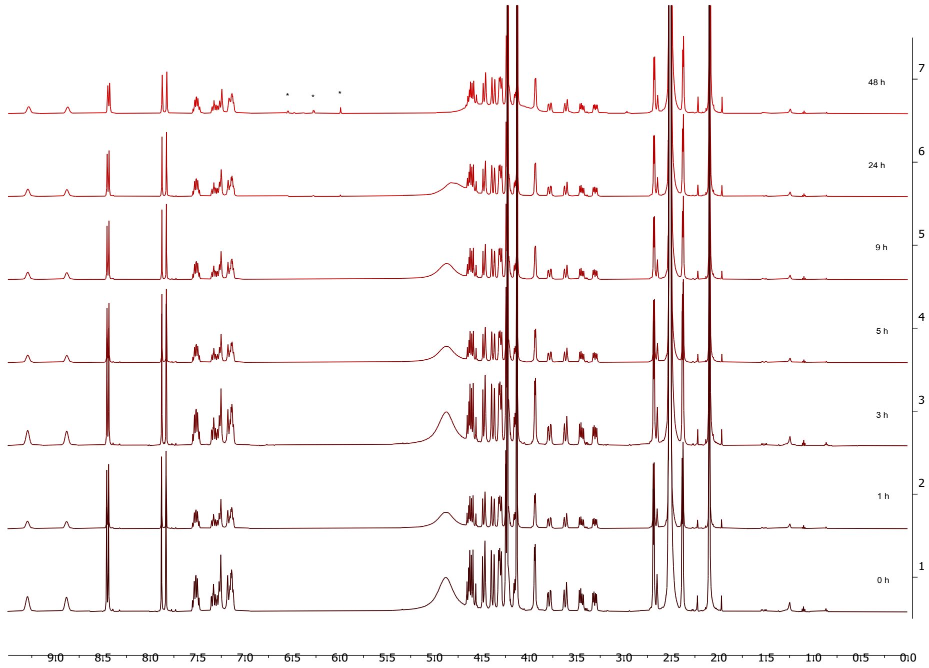

**Figure S14.** Stability of **2a** in DMSO up to two days.

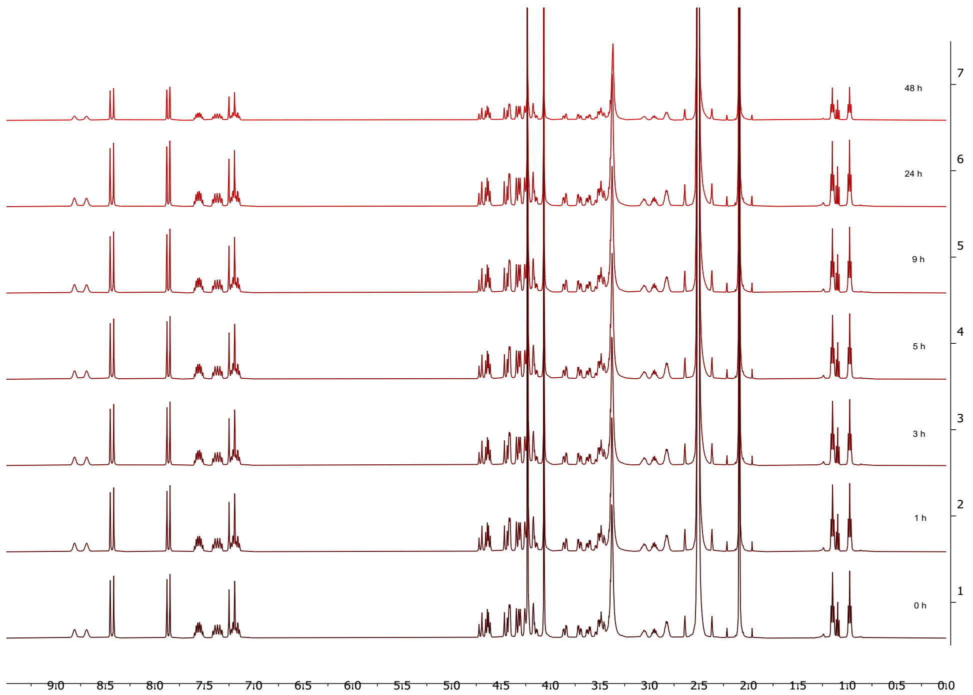

**Figure S15.** Stability of **3a** in DMSO up to two days.

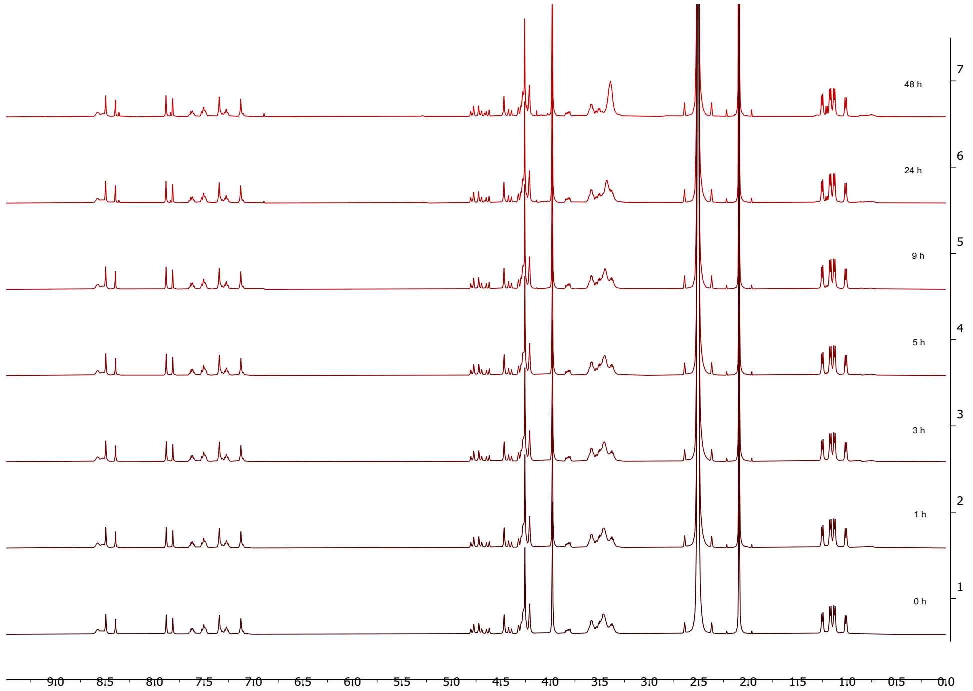

**Figure S16.** Stability of **4a** in DMSO up to two days.

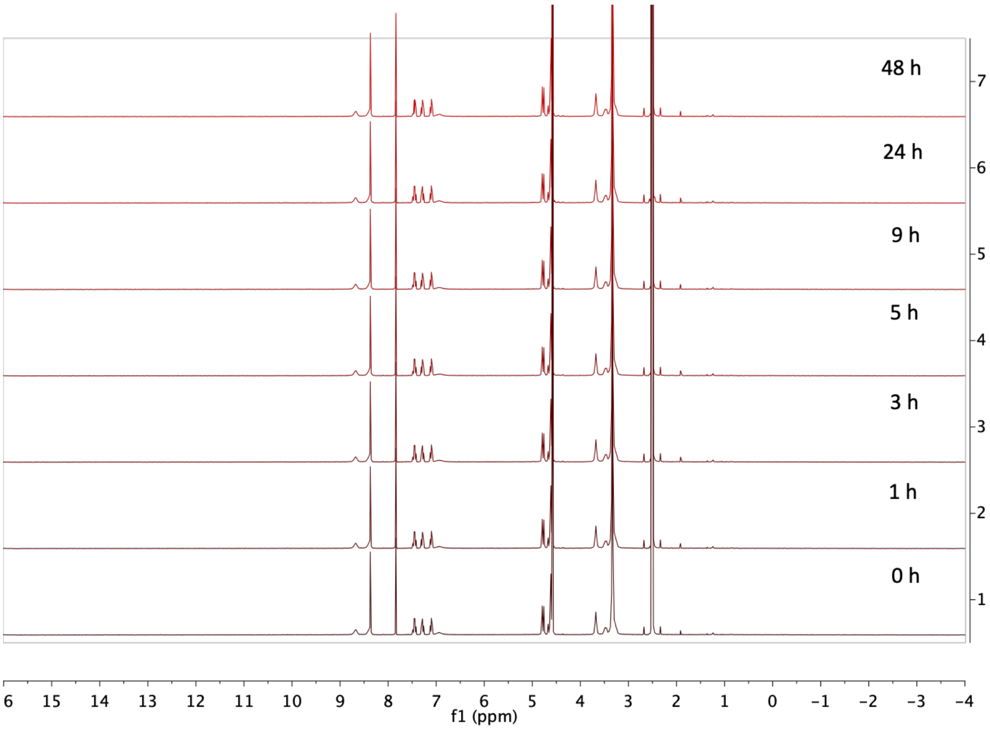

**Figure S17.** Stability of **5a** in DMSO up to two days.

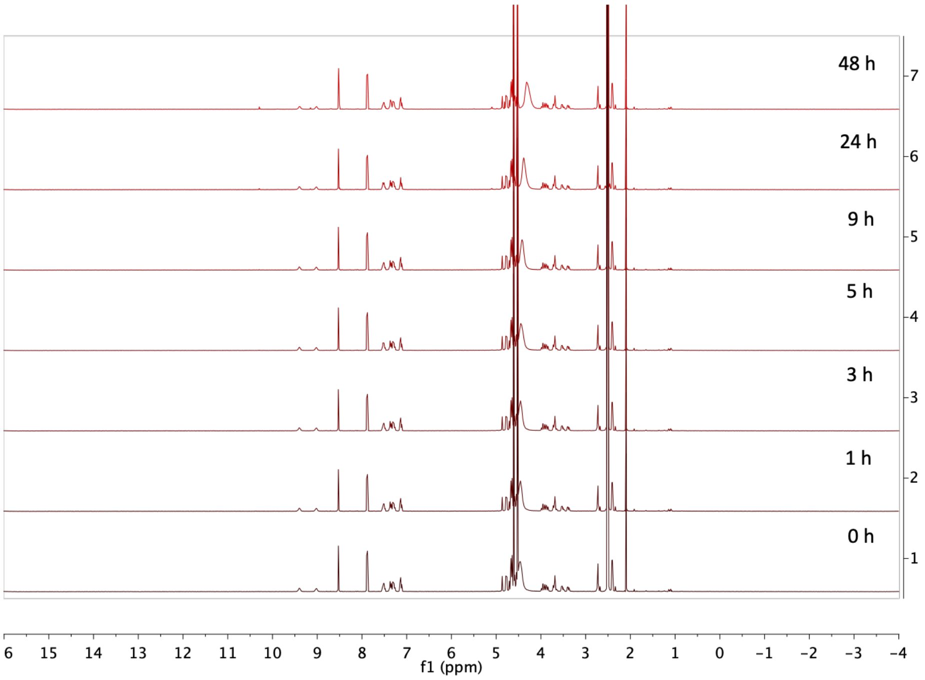

**Figure S18.** Stability of **6a** in DMSO up to two days.

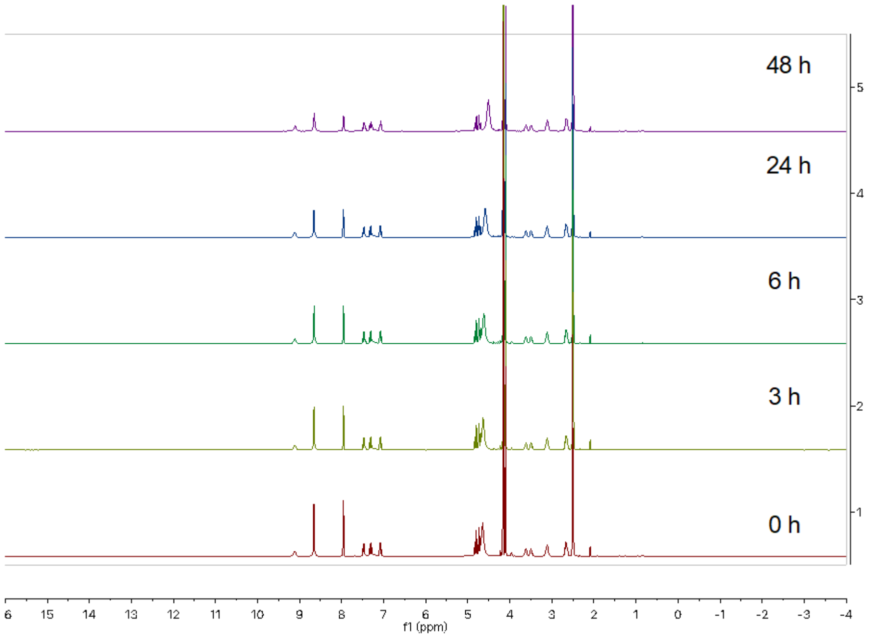

**Figure S19.** Stability of **7a** in DMSO up to two days.

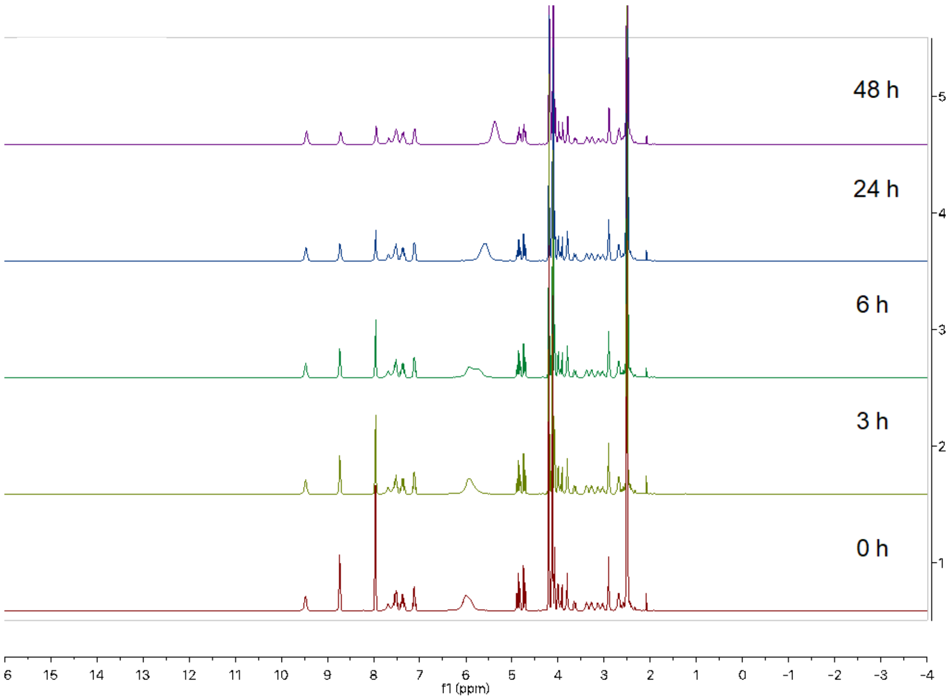

**Figure S20.** Stability of **8a** in DMSO up to two days.

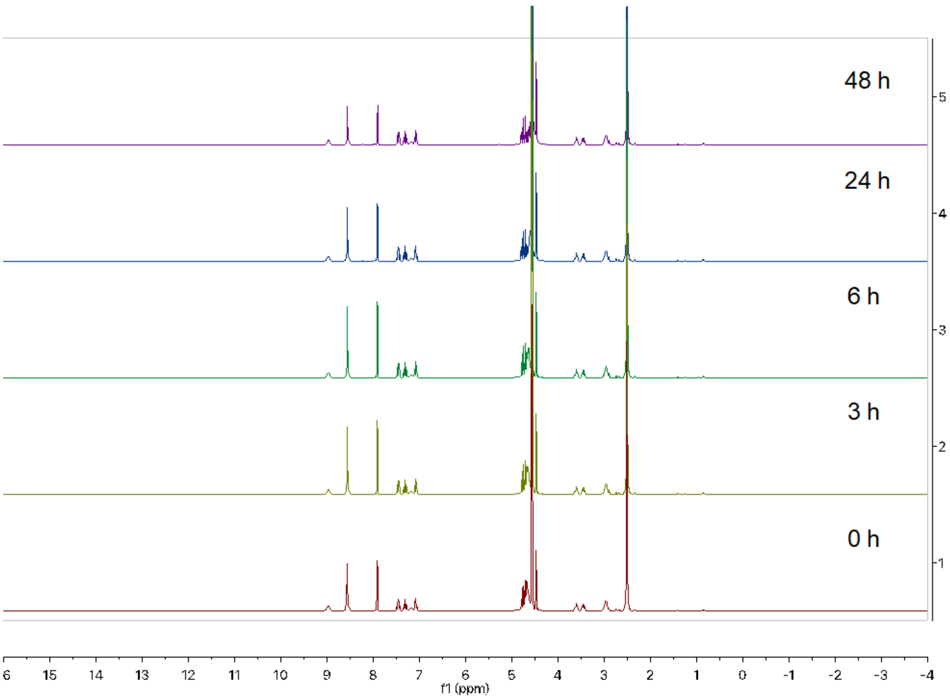

**Figure S21.** Stability of **9a** in DMSO up to two days.

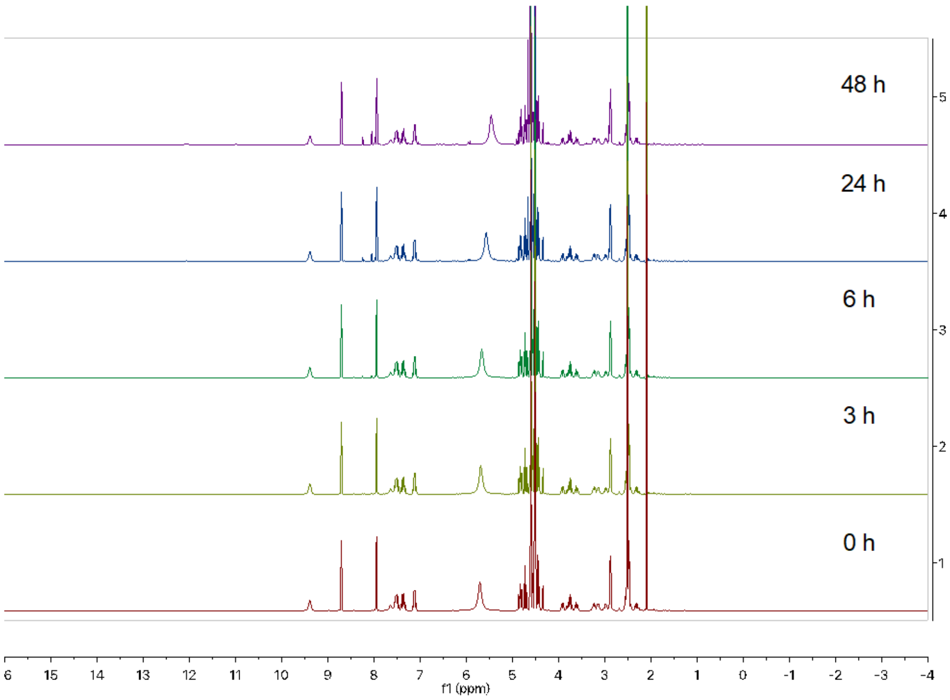

**Figure S22.** Stability of **10a** in DMSO up to two days.

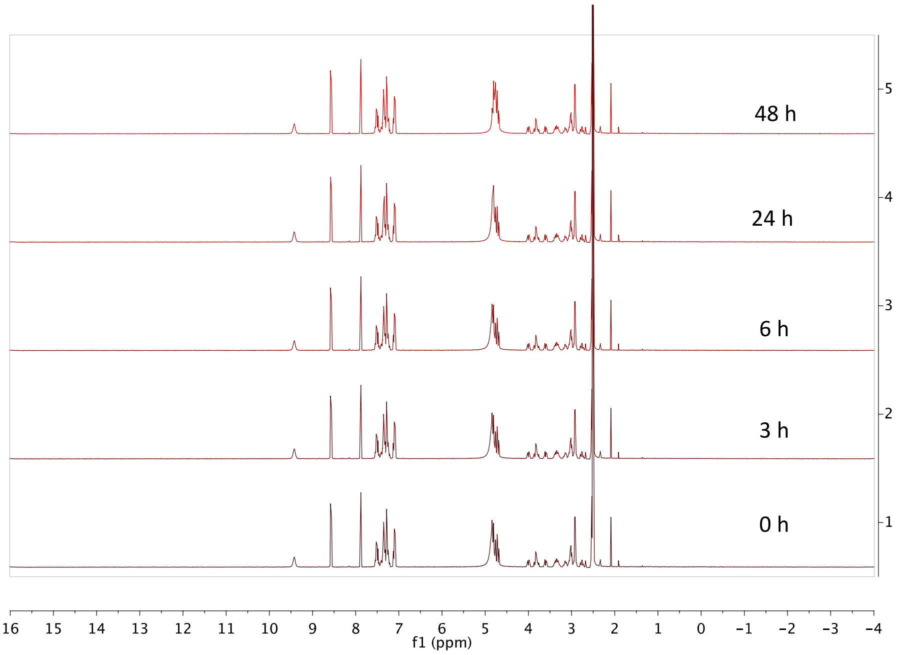

**Figure S23.** Stability of **11a** in DMSO up to two days.

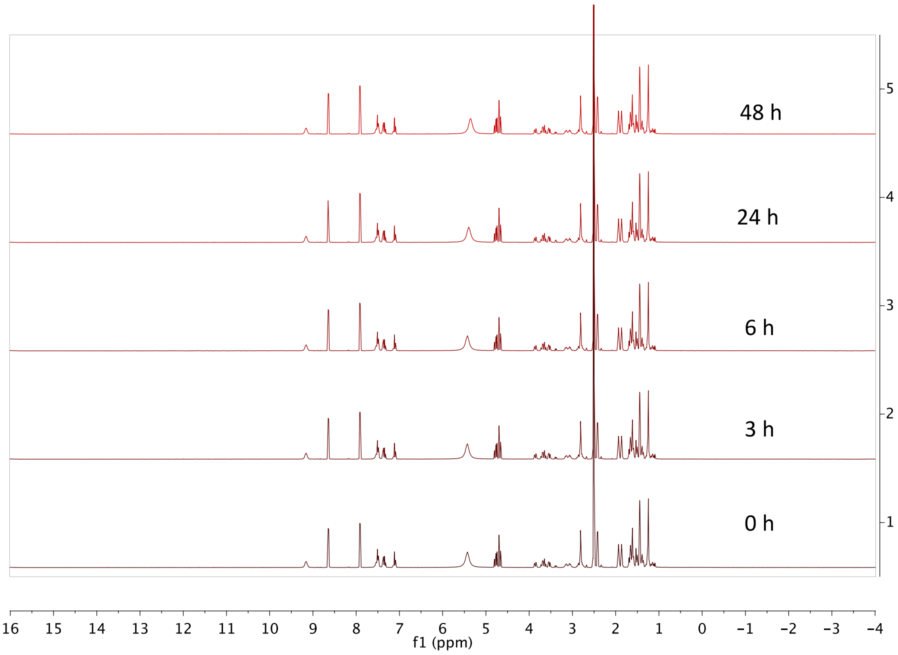

**Figure S24.** Stability of **12a** in DMSO up to two days.

#### 2. HPLC spectra for microsome stability testing

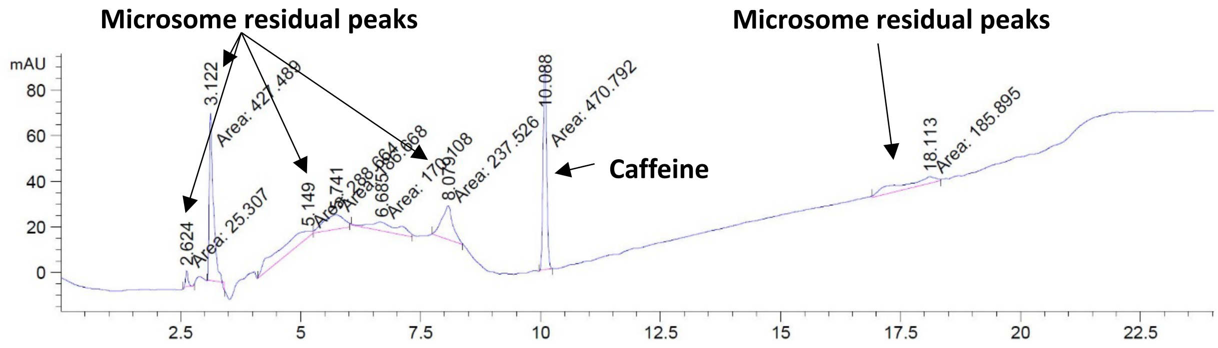

**Figure S25.** Microsome control at 0 h.

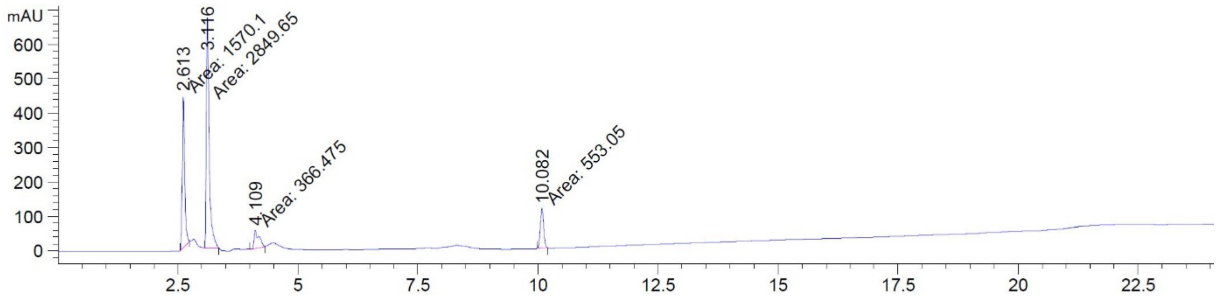

**Figure S26.** Microsome control at 48 h.

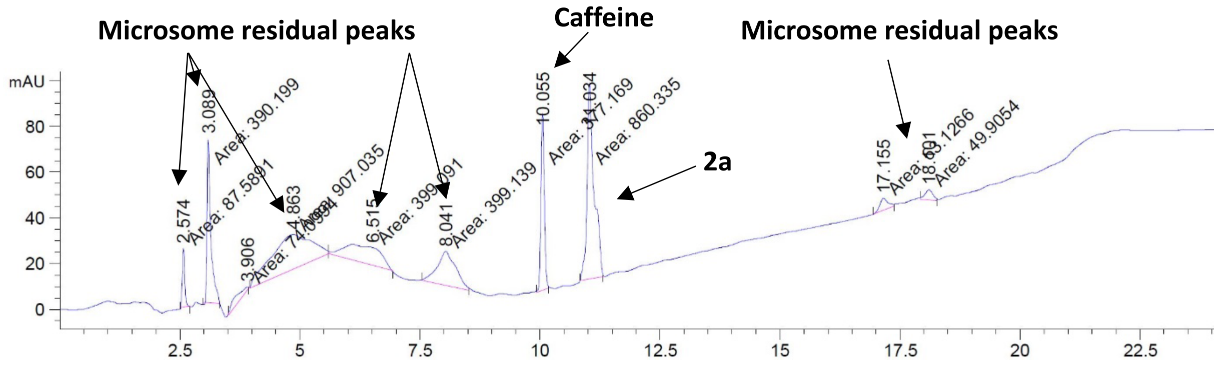

**Figure S27.** Compound **2a** after 0 h incubation**.**

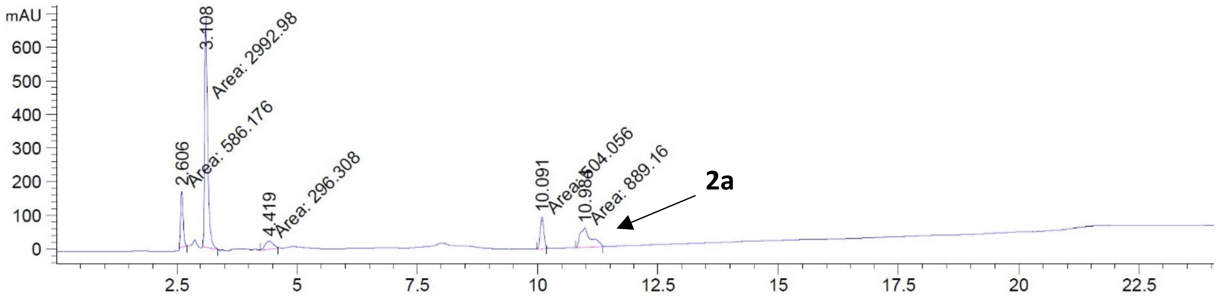

**Figure S28.** Compound **2a** after 6 h incubation.

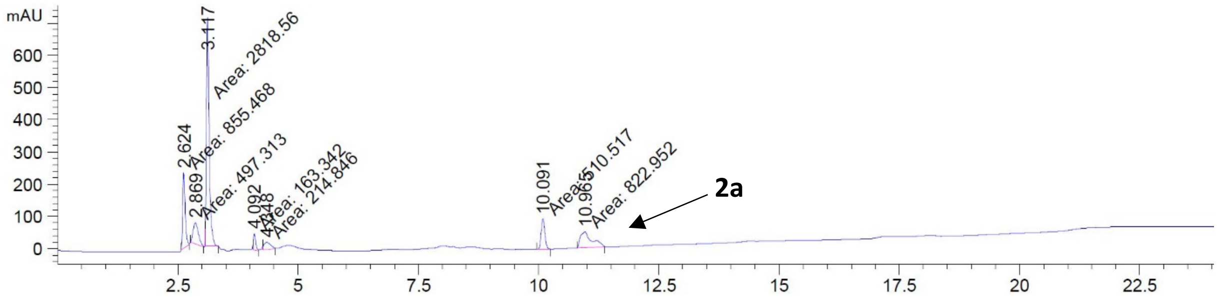

**Figure S29.** Compound **2a** after 24 h incubation**.**

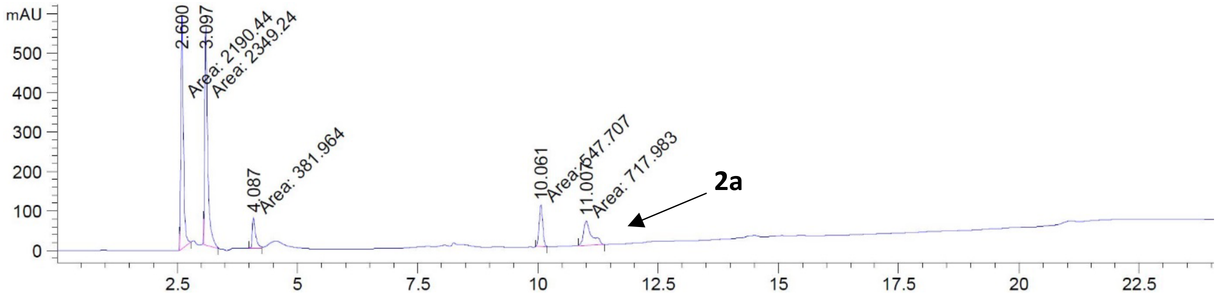

**Figure S30.** Compound **2a** after 48 h incubation.

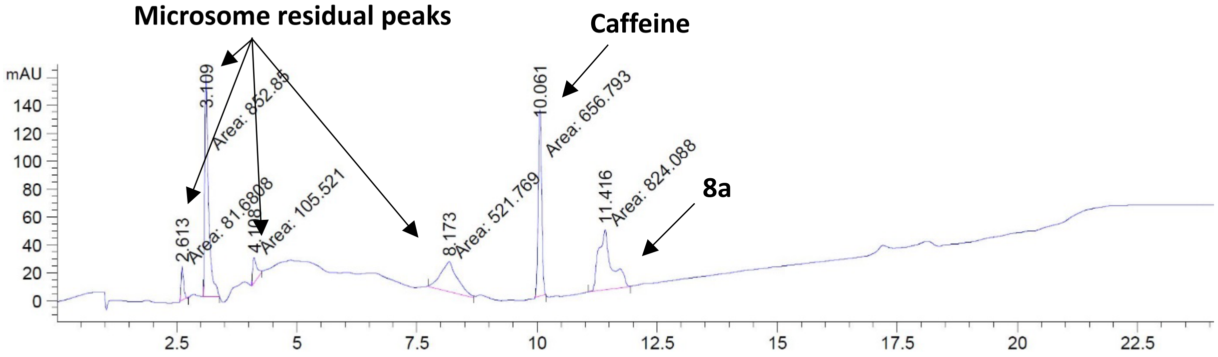

**Figure S31.** Compound **8a** after 0 h incubation.

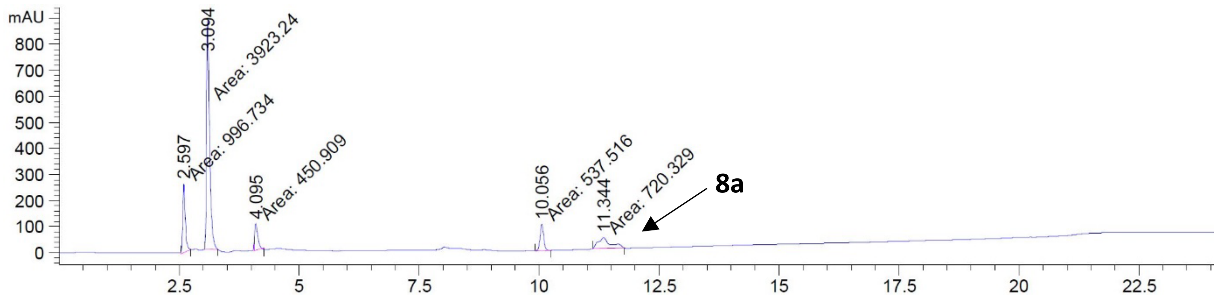

**Figure S32.** Compound **8a** after 6 h incubation.

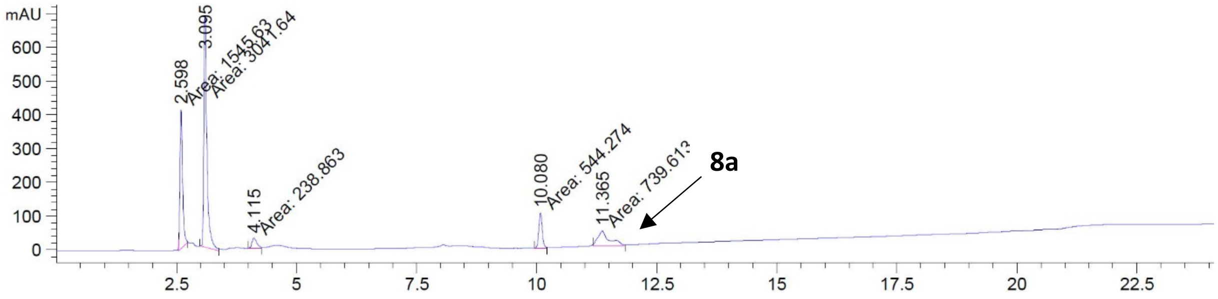

**Figure S33.** Compound **8a** after 24 h incubation.

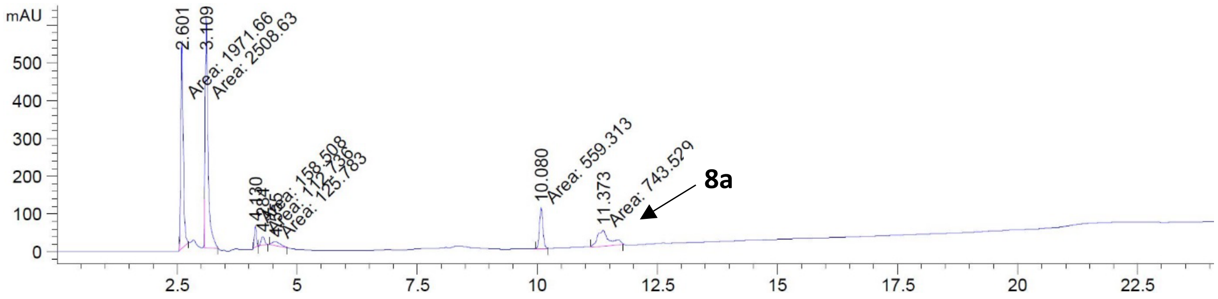

**Figure S34.** Compound **8a** after 48 h incubation.

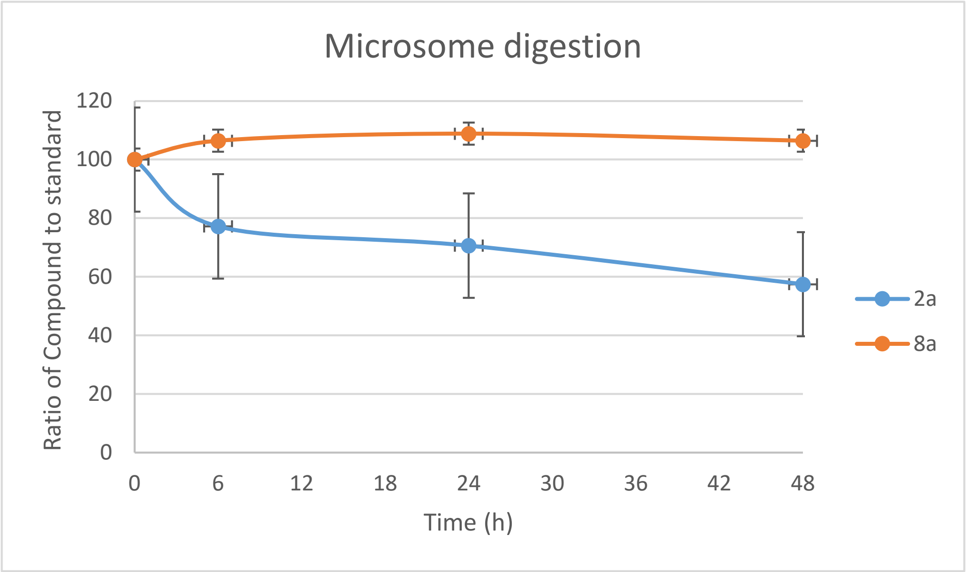

**Figure S35.** Metabolic stability of compounds **2a** and **8a.**

#### 3. Worm burden

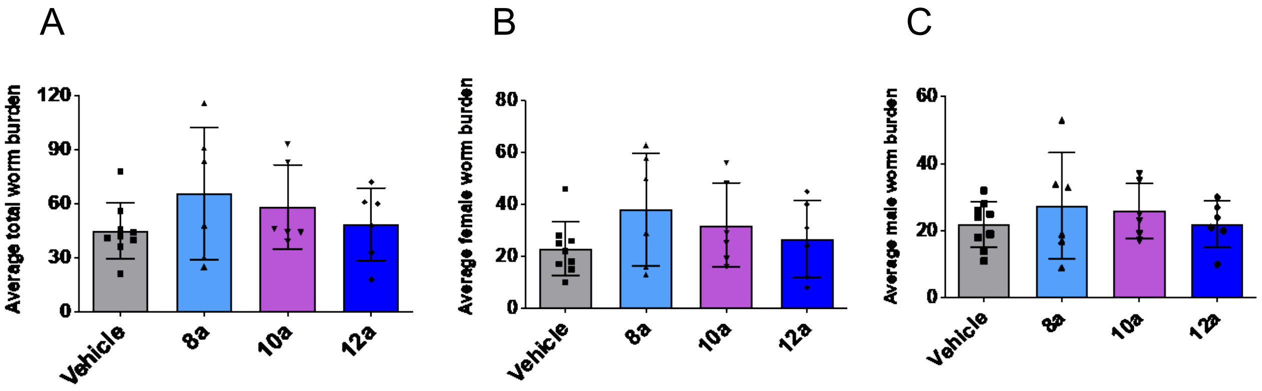

**Figure S36.** Adult *B. pahangi* worms were removed and counted from each jird at necropsy. There was no significant difference in (**A)** the total number of worms recovered from each group, (**B**) the number of female worms and (**C**) the number of male worms compared to the vehicle control group. Each dot represents the total number of worms from each animal; horizontal bars represent the mean ± SD for each group. There was no significant difference in the number of female vs male worms (sex ratio) within each group (data not shown).

#### 4. Embryogram results

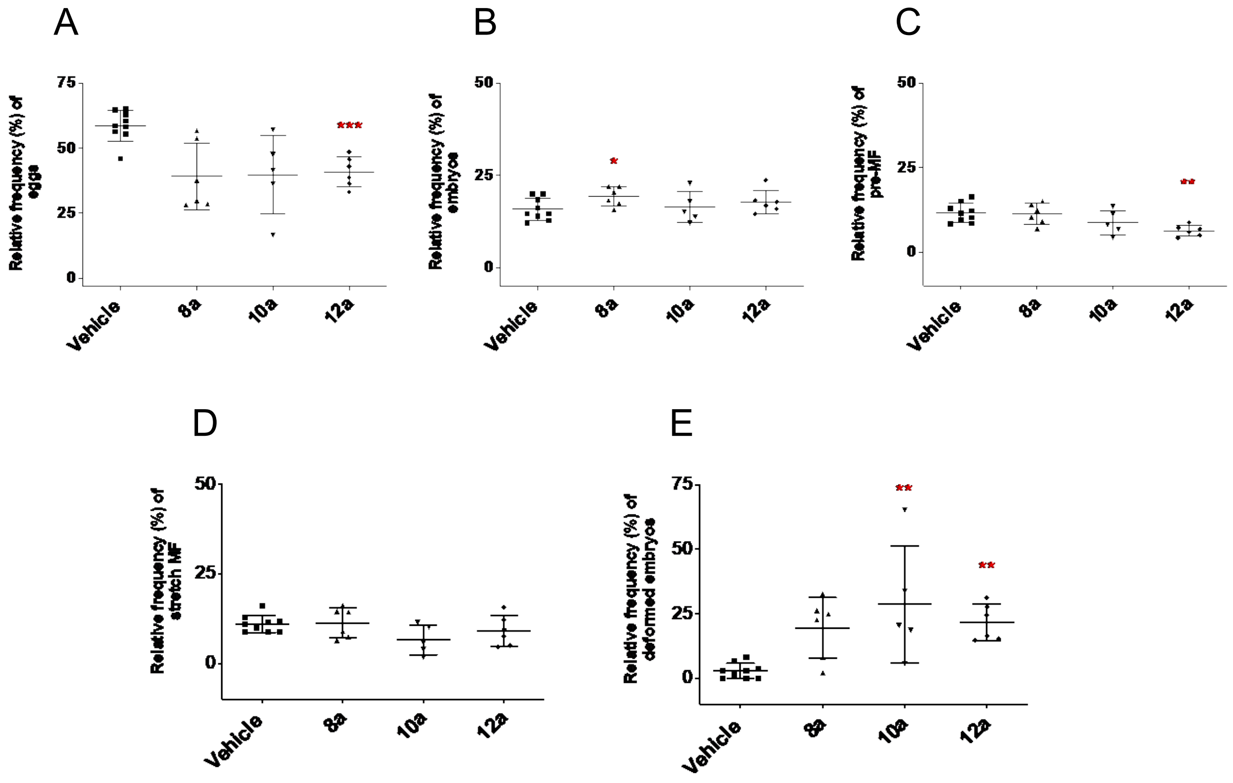

**Figure S37.** Embryogram analyses were conducted on 12-36 female *B. pahangi* worms per group to determine the effects of the compounds on the developing stages within the reproductive tract. Each graph shows the means and SD for each of the developmental stages: (**A**) eggs, (**B**) embryos, (**C**) pre-MF, (**D**) stretch MF and (**E**) deformed embryos. *P<0.05; **P<0.01; ***P<0.001.

#### 5. Chemogenomic screen

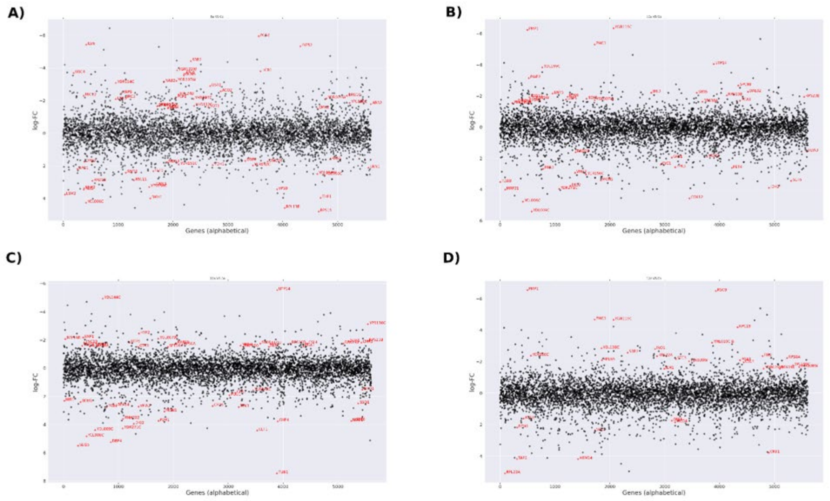

**Figure S38.** To identify potential drug targets, the pool of tagged 5936 *S. cerevisiae* heterozygous deletion mutants was grown competitively for 20 generations in the presence and absence of the compounds (**A**) **8a**, (**B**) **10a**, (**C**) **11a** and (**D**) **12a**. Strain fitness was determined via high-throughput barcode sequencing and normalization to the untreated control. Log 2 ratio (control intensity/treatment intensity) was calculated and plotted as a function of gene (red dots: logFC=1.5 and P value=0.01).

#### 6. Cytotoxicity results

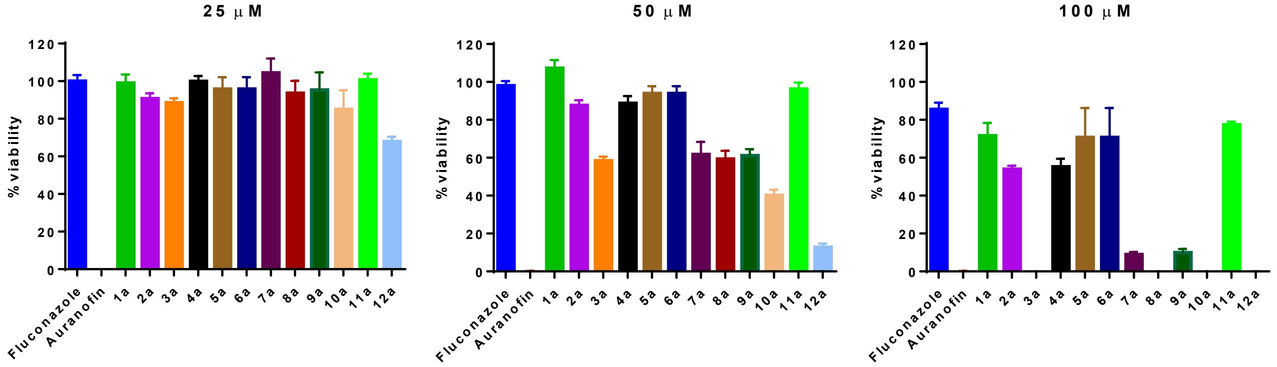

**Figure S39.** RPE-1 cells viability following 48 h of incubation with 25, 50 or 100 µM of FCZ, auranofin or compounds **1a-12a.**

#### 7. Primer sequences

| oligoname | Sequence 5' - 3' |
| --- | --- |
| UPTAG Index 1 | ACG CTC TTC CGA TCT ATACC GTC CAC GAG GTC TCT |
| UPTAG Index 2 | ACG CTC TTC CGA TCT TCTAG GTC CAC GAG GTC TCT |
| UPTAG Index 3 | ACG CTC TTC CGA TCT GCAGC GTC CAC GAG GTC TCT |
| UPTAG Index 4 | ACG CTC TTC CGA TCT CCGAG GTC CAC GAG GTC TCT |
| UPTAG Index 5 | ACG CTC TTC CGA TCT AAGAT GTC CAC GAG GTC TCT |
| UPTAG Index 6 | ACG CTC TTC CGA TCT TAGTA GTC CAC GAG GTC TCT |
| UPTAG Index 7 | ACG CTC TTC CGA TCT GAACT GTC CAC GAG GTC TCT |
| UPTAG Index 8 | ACG CTC TTC CGA TCT CAATG GTC CAC GAG GTC TCT |
| UPTAG Index 9 | ACG CTC TTC CGA TCT AGACA GTC CAC GAG GTC TCT |
| UPTAG Index 10 | ACG CTC TTC CGA TCT TGTTC GTC CAC GAG GTC TCT |
| UPTAG Index 11 | ACG CTC TTC CGA TCT GCCGG GTC CAC GAG GTC TCT |
| UPTAG Index 12 | ACG CTC TTC CGA TCT CGTAG GTC CAC GAG GTC TCT |
| UPTAG Index 13 | ACG CTC TTC CGA TCT ATACC GTC CAC GAG GTC TCT |
| UPTAG Index 14 | ACG CTC TTC CGA TCT TGTAA GTC CAC GAG GTC TCT |
| UPTAG Index 15 | ACG CTC TTC CGA TCT GTATT GTC CAC GAG GTC TCT |
| UPTAG Index 16 | ACG CTC TTC CGA TCT CTCTA GTC CAC GAG GTC TCT |
| DNTAG Index 1 | ACG CTC TTC CGA TCT ATACC GTG TCG GTC TCG TAG |
| DNTAG Index 2 | ACG CTC TTC CGA TCT TCTAG GTG TCG GTC TCG TAG |
| DNTAG Index 3 | ACG CTC TTC CGA TCT GCAGC GTG TCG GTC TCG TAG |
| DNTAG Index 4 | ACG CTC TTC CGA TCT CCGAG GTG TCG GTC TCG TAG |
| DNTAG Index 5 | ACG CTC TTC CGA TCT AAGAT GTG TCG GTC TCG TAG |
| DNTAG Index 6 | ACG CTC TTC CGA TCT TAGTA GTG TCG GTC TCG TAG |
| DNTAG Index 7 | ACG CTC TTC CGA TCT GAACT GTG TCG GTC TCG TAG |
| DNTAG Index 8 | ACG CTC TTC CGA TCT CAATG GTG TCG GTC TCG TAG |
| DNTAG Index 9 | ACG CTC TTC CGA TCT AGACA GTG TCG GTC TCG TAG |
| DNTAG Index 10 | ACG CTC TTC CGA TCT TGTTC GTG TCG GTC TCG TAG |
| DNTAG Index 11 | ACG CTC TTC CGA TCT GCCGG GTG TCG GTC TCG TAG |
| DNTAG Index 12 | ACG CTC TTC CGA TCT CGTAG GTG TCG GTC TCG TAG |
| DNTAG Index 13 | ACG CTC TTC CGA TCT ACACG GTG TCG GTC TCG TAG |
| DNTAG Index 14 | ACG CTC TTC CGA TCT TGTAA GTG TCG GTC TCG TAG |
| DNTAG Index 15 | ACG CTC TTC CGA TCT GTATT GTG TCG GTC TCG TAG |
| DNTAG Index 16 | ACG CTC TTC CGA TCT CTCTA GTG TCG GTC TCG TAG |
| UPkanMX | CAA GCA GAA GAC GGC ATA CGA GAT GTC GAC CTG CAG CGT ACG |
| DNkanMX | CAA GCA GAA GAC GGC ATA CGA GAT ACG AGC TCG AAT TCA TCG |
| P5 | A ATG ATA CGG CGA CCA CCG AGA TCT ACA CTC TTT CCC TAC ACG ACG CTC TTC CGA TCT |

**Table S1.** Complete list of primer sequences

### 8. Synthesis and characterization

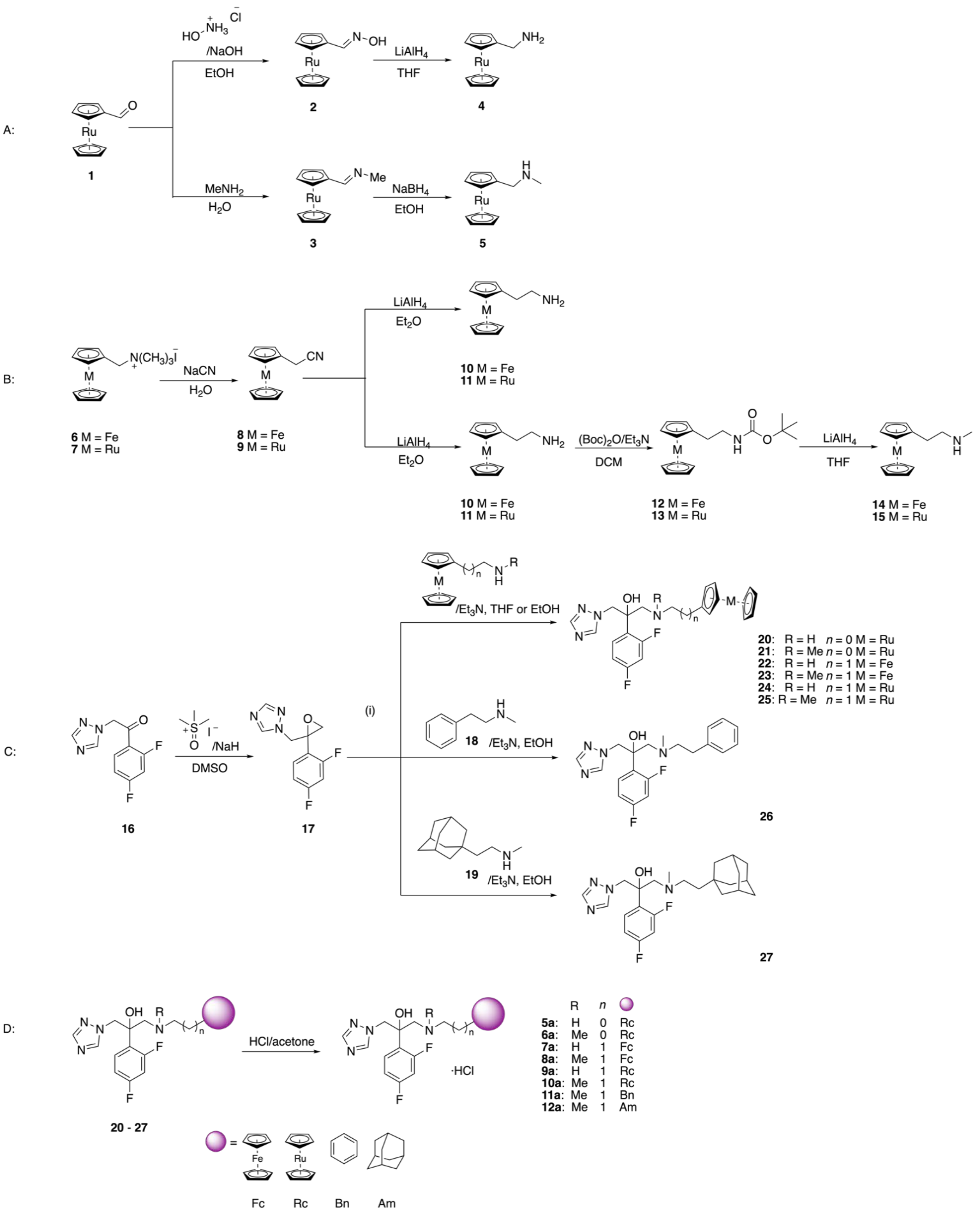

**Scheme S40**. Synthetic route to eight newly developed analogues **5a-12a**.

#### 9. Preparation of starting material

**Materials**. All chemicals were of reagent grade quality, obtained from commercial suppliers unless otherwise stated and used without further purification. Solvents were dried over molecular sieves. All preparations were carried out using standard Schlenk techniques, unless otherwise stated.

**Instrumentation and Methods**. Evaporation of the solvents in vacuo was done with a rotary evaporator at 40°C. Thin-layer chromatography (TLC) was performed using silica gel 60 F-254 (Merck) plates with detection of spots being achieved by exposure to UV light. 1H and 13C NMR spectra were recorded in deuterated solvents on Bruker AV-400 (1H, 500 MHz; 19F, 376 MHz) and AV-500, AV-501 (1H, 500 MHz; 13C, 126 MHz) spectrometers at room temperature. The chemical shifts, δ, are reported in ppm (parts per million). The residual solvent peaks have been used as an internal reference. The abbreviations for the peak multiplicities are as follows: s (singlet), d (doublet), t (triplet), m (multiplet). Elemental microanalyses were performed on a LECO TruSpec Micro elemental analyser. The UPLC-MS spectra were measured on an Acquity from Waters system equipped with a PDA detector and an auto sampler using an Agilent Zorbax 300SB-C18 analytical column (3.5 µm particle size, 300 Å pore size, 150 x 4.6 mm). This UPLC was coupled to an Esquire HCT apparatus from Bruker for the MS measurements. The UPLC run (flow rate 0.3 mL min-1) was performed with a linear gradient of A (distilled water containing 0.1% v/v formic acid) and B (acetonitrile, Sigma-Aldrich HPLC grade): t = 0 min, 0% B; t = 1 min, 0% B; t = 20 min, 66% B. Infrared spectra were recorded on a Bruker Vertex 70 FTIR spectrometer.

**Ruthenocene carboxaldehyde** (**1**) was prepared following several adapted literature procedures.*(1–3)* A solution of ruthenium trichloride hydrate ($RuCl_3 \times H_2O$, 31.4 g, 0.12 mol, 1 equiv.) in 200 mL of absolute ethanol was placed in an ice bath cooled to 0 °C. Subsequently, cyclopentadiene (100 mL, 79.5 g, 1.20 mol, 10 equiv.) was added to the solution. Zinc dust (78 g, 1.20 mol, 10 equiv.) was added to the reaction mixture over 1 h in 10 portions, while the temperature was kept between 0 - 10 °C during the addition. The reaction mixture was stirred at 0 °C for 30 min, and then warmed to room temperature (23 °C) with continued stirring for 3 h. The suspension was filtered, and the grey metallic residue was washed with hot toluene (90 °C). The filtrate was concentrated in vacuo, the residue was dissolved in toluene at room temperature. The solution was passed through a plug of silica gel with toluene as eluent. The toluene was removed in vacuo to obtain ruthenocene as an off-white solid (9.02 g, 0.039 mol, 33%). **$^1$H NMR** (400 MHz, $CDCl_3$): δ 4.56 (s, 10H).

Ruthenocene (1.00 g, 4.3 mmol, 1 equiv.), and KO*t*Bu (56 mg, 0.5 mmol, 0.12 equiv) were dissolved in dry THF (100 mL) and cooled to -78 ℃. At -78 ℃, *t*BuLi in pentane (4.6 mL, 8.6 mmol, 1.9 M, 2 equiv.) was added dropwise to the solution over a period of 30 min. After the addition of *t*BuLi, dry dimethylformamide (0.8 mL, 0.8 g, 10.9 mmol, 2.5 equiv.) was added dropwise into the reaction mixture, and stirred for another 10 min. The reaction mixture was then warmed to -40℃, and stirred for a further 10 min, before deionized water (50 mL) was added. The THF was removed in vacuo and the aqueous solution was extracted with DCM (50 mL × 3). The DCM layers were combined, washed with water (50 mL × 2), dried with $MgSO_4$, filtered, and the DCM removed in vacuo. The residue was purified by flash chromatography using DCM as eluent (Rf = 0.61), to obtain **1** as a bright yellow solid (0.90 g, 3.49 mmol, 82%). **$^1$H NMR** (400 MHz, Chloroform-d) δ 9.72 (s, 1H), 5.10 – 5.05 (m, 2H), 4.87 – 4.83 (m, 2H), 4.64 (s, 5H). The spectral data corresponds to previously reported data.*(4)*

**Ruthenocenylmethylamine** (**4**) was prepared following an adapted published literature procedure.*(5)* A solution of RcCHO (389 mg, 1.50 mmol, 1 equiv.) in EtOH (15 mL) was added $(NH_3OH)Cl$ (417 mg, 6.00 mmol, 4 equiv.) and NaOH (360 mg, 9.00 mmol, 6 equiv.). The resultant mixture was stirred under reflux for 2.5 h, and water (35 mL) was added. The mixture was extracted with DCM (50 mL × 4), and the organic layers combined, dried with $MgSO_4$, filtered, and the solvent was removed in vacuo to obtain ruthenocenecarbaldehyde oxime as a yellow solid (408 mg, 1.48 mmol, 95%). **$^1$H NMR** (400 MHz, CDCl3): δ 7.82 (s, 1H), 4.90 – 4.87 (m, 2H), 4.68 – 4.65 (m, 2H), 4.59 (m, 5H).

The ruthenocenecarbaldehyde oxime (**2**) obtained (408 mg, 1.48 mmol, 1 equiv.) was dissolved in dry THF (7 mL) and a solution of $LiAlH_4$ (314.8 mg, 7.4 mmol, 5 equiv.) in dry THF (8 mL) was added. The resultant grey suspension was stirred under reflux for 3 h, cooled to room temperature, and quenched with 25 mL of saturated aqueous $Na_2SO_4$ solution. The mixture was filtered with Celite (packed with pentane), and the filtrate extracted with DCM (50 mL × 3). The organic layers were combined, dried with $MgSO_4$, filtered, and the DCM was removed in vacuo to obtain ruthenocenylmethylamine **4** as a dark green solid (272 mg, 1.04 mmol, 71%). **$^1$H NMR** (400 MHz, CDCl3): δ 4.64 – 4.62 (m, 2H), 4.58 (s, 5H), 4.53 – 4.48 (m, 2H), 3.38 (s, 2H). The spectral data corresponds to previously reported data.*(4)*

**1-ruthenocenyl-N-methylmethanamine** (**5**) To a solution of RcCHO (100 mg, 0.386 mmol, 1 equiv.) in toluene (2 mL) was added aqueous $MeNH_2$ (40 wt. % in $H_2O$, 1.85 mL) and the biphasic mixture was stirred at 70°C for 2 h. The toluene was removed under reduced pressure, and $H_2O$ was added to the residue. The solution was extracted twice with $Et_2O$, dried with $MgSO_4$, filtered, and evaporated under reduced pressure to obtain 1-ruthenocenyl-N-methylmethanimine (175 mg, 0.643 mmol). The imine was then dissolved in EtOH (6 mL), and $NaBH_4$ (243 mg, 6.43 mmol, 10 equiv.) was added. The white solution with grey suspension was stirred at room temperature for 2 h, the reaction was quenched by adding $H_2O$ (20 mL), and the mixture was extracted twice with $Et_2O$. The organic layers were combined, dried with $MgSO_4$, filtered, and the $Et_2O$ was removed under reduced pressure to obtain 1-ruthenocenyl-N-methylmethanamine **5** as a white oil (80 mg, 0.29 mmol, 76%). **$^1$H NMR** (400 MHz, CDCl3): δ 4.65 – 4.63 (m, 2H), 4.54 (s, 5H), 4.50 – 4.48 (m, 2H), 3.29 (s, 2H), 2.48 (s, 3H). The spectral data corresponds to previously reported data.*(4)*

**Trimethylammoniomethylruthenocene iodide (7)** was synthesized following an adapted literature procedure.*(6)* The spectroscopic data matched those previously reported. **$^1$H NMR** (400 MHz, Chloroform-d) δ 4.91 (t, *J* = 1.7 Hz, 2H), 4.71 – 4.70 (m, 2H), 4.67 (m, 5H), 4.56 (s, 2H), 3.38 (s, 9H).

**Ferroceneacetonitrile (8)** was synthesized following an adapted literature procedure.*(7)* Ferrocenylmethyltrimethylammonium iodide (2.3 g, 6.1 mmol) was added to 30 mL of a solution of NaCN (3.0 g, 61.8 mmol) in water. The reaction mixture was stirred at reflux overnight and then cooled to rt. The solid product was separated by filtration and dissolved in diethyl ether, while the filtrate was extracted with diethyl ether (3 x 100 mL). The organic layers were combined wash with water and dry over $Na_2SO_4$. Evaporation of solvent under reduced pressure give compound as an orange solid. Yield: 74% (1.03 g, 4.5 mmol). The spectroscopic data matched those previously reported.*(7)* **$^1$H NMR** (400 MHz, Chloroform-d) δ 4.24 (bs, 7H), 4.17 (s, 2H), 3.42 (s, 2H).

**Ruthenoceneacetonitrile (9).** Trimethylammoniomethylrutheneocene iodide (4.3 g, 10 mmol) was added to 100 mL of a solution of NaCN (4.9 g, 100 mmol) in water. The reaction mixture was stirred at reflux overnight and then cooled to rt. The solid product was separated by filtration and dissolved in diethyl ether, while the filtrate was extracted with diethyl ether (3 x 100 mL). The organic layers were combined and washed with water and dry over $Na_2SO_4$. Evaporation of solvent under reduced pressure give compound as a pale grey solid. Yield: 79%

(2.13 g, 7.9 mmol). **$^1$H NMR** (400 MHz, Chloroform-*d*) δ 4.69 (t, *J* = 1.7 Hz, 2H), 4.61 (s, 5H), 4.54 – 4.53 (m, 2H), 3.37 (s, 2H). **$^{13}$C NMR** (101 MHz, Chloroform-*d*) δ 118.1, 80.8, 71.6, 70.8, 70.6, 19.4. **ESI-MS**: m/z calcd. for $C_{12}H_{11}NRu^+$, $[M]^+$ 270.9930, found 270.9931.

**2-(Ferrocenyl)ethylamine (10)** was synthesized following an adapted literature procedure.*(8)* $FcCH_2CN$ (3.4 g, 15 mmol) in dry diethyl ether (50 mL) was added dropwise to a stirred suspension of $LiAlH_4$ (1.14 g, 30 mmol) in 70 mL of dry diethyl ether and heated at 45°C. After being stirred for 4 h under reflux, the reaction mixture was externally cooled with an ice bath, and cold water was slowly added dropwise until $H_2$ release stopped. Once decanted, the solution was acidified by addition of 6 M $H_2SO_4$. The solid formed was separated by filtration and treated with a 6 M NaOH aqueous solution to adjust the PH ~ 11. The dark orange mixture was extracted with diethyl ether, the organic layers were combined, dried over $Na_2CO_3$, filtered and the solvent was removed under vacuum to give compound **10** as a brown oil. Yield: 74% (2.53 g, 11 mmol). The spectroscopic data matched those previously reported.*(8)* **$^1$H NMR** (400 MHz, Chloroform-d) δ 4.10 (s, 5H), 4.07 (d, *J* = 1.8 Hz, 2H), 4.06 (d, *J* = 1.7 Hz, 2H), 2.80 (t, *J* = 7.0 Hz, 2H), 2.47 (t, *J* = 7.0 Hz, 2H).

**2-(Ruthenocenyl)ethylamine (11).** Ruthenoceneacetonitrile **9** (2.10 g, 7.8 mmol) in dry diethyl ether (50 mL) was added dropwise to a stirred suspension of $LiAlH_4$ (0.6 g, 16 mmol) in 50 mL of dry diethyl ether and heated at reflux. After being stirred for 4h under reflux, the reaction mixture was externally cooled with an ice bath, and cold water was slowly added dropwise until the $H_2$ release stopped. Once decanted, the solution was acidified by addition of 6 M $H_2SO_4$. The solid formed was separated by filtration and treated with a 6 M NaOH aqueous solution to adjust the PH ~ 11. The dark orange mixture was extracted with diethyl ether, the organic phases were combined, dried over $Na_2CO_3$, filtered and the solvent was removed under vacuum to give compound **11** as a yellow oil. Yield: 70% (1.51 g, 5.5 mmol). **$^1$H NMR** (400 MHz, Chloroform-*d*) δ 4.51 (s, 7H), 4.45 – 4.44 (m, 2H), 2.77 (t, *J* = 7.0 Hz, 2H), 2.32 (t, *J* = 7.0 Hz, 2H). **$^{13}$C NMR** (101 MHz, Chloroform-d) δ 90.2, 71.0, 70.5, 69.5, 44.2, 33.3**. ESI-MS**: m/z calcd. for $C_{12}H_{16}NRu^+$, $[M+H]^+$ 276.0321, found 276.0324.

***N*-Methyl-$\beta$-ferrocenylethylamine (14)** was synthesized in two steps following an adapted literature procedure.*(9)* $\beta$-ferrocenylethylamine (0.23 g, 1 mmol) and triethylamine (0.2 mL, 1.5 mmol) were dissolved in methylene chloride (5 mL). Di-*tert*-butyl dicarbonate (0.22 g, 1 mmol) was added and the mixture was stirred at room temperature for 10 h. Evaporation of solvent under reduced pressure and purification by chromatography on silica with hexanes :

ethyl ecetate (8 : 1) as the eluent ($R_f$ = 0.52, hexane : ethyl acetate (5 : 1)) afforded the *N*-(*tert*-butyloxycarbonyl) derivative of $\beta$-ferrocenylethylamine **12** as a yellow oil: 0.20 g (60%). The spectroscopic data matched those previously reported.*(9)* **$^1$H NMR** (400 MHz, Chloroform-d) δ 4.68 (br, 1H), 4.12 (s, 5H), 4.08 (s, 4H), 3.25 (q, *J* = 6.9 Hz, 2H), 2.52 (t, *J* = 6.9 Hz, 2H), 1.45 (s, 9H).

A mixture of the the *N*-Boc derivative obtained in the previous step (0.16 g, 0.48 mmol) and $LiAlH_4$ (0.07 g, 1.92 mmol) in anhydrous THF (5 mL) was heated under reflux for 12 h. The reaction mixture was cooled in an ice bath and then diluted with methylene chloride, followed by the dropwise addition of a saturated aqueous solution of $Na_2SO_4$ until no further effervescence was observed. The mixture was stirred for 10 min, after which the clear solution was decanted from the white precipitate. The precipitate was washed with DCM and the combined organic layer was concentrated *in vacuo* to give compound **14** as an oil: 0.10 g (90%). The spectroscopic data matched those previously reported.*(9)* **$^1$H NMR** (400 MHz, Chloroform-d) δ 4.10 (s, 5H), 4.10 – 4.07 (m, 2H), 4.07 – 4.05 (m, 2H), 2.70 (td, *J* = 7.0, 0.8 Hz, 2H), 2.54 (t, *J* = 7.0 Hz, 2H), 2.42 (s, 3H).

***N*-Methyl-$\beta$-ruthenocenylethylamine (15).** $\beta$-ruthenocenylethylamine (2.27 g, 8.28 mmol) and triethylamine (2.1 mL, 15 mmol) were dissolved in methylene chloride (50 mL). Di-*tert*-butyl dicarbonate (2.2 g, 10 mmol) was added, and the mixture was stirred at room temperature for 10 h. Evaporation of the solvent under reduced pressure and purification by chromatography on silica with hexanes : ethyl ecetate (6 : 1) as the eluent ($R_f$ = 0.22, hexane : ethyl acetate (10 : 1)) afforded the *N*-(*tert*-butyloxycarbonyl) derivative of $\beta$-ruthenocenylethylamine **13** as a white solid: 2.59 g (83%). **$^1$H NMR** (400 MHz, Chloroform-d) δ 5.16 (s, 1H), 4.56 (s, 5H), 4.51 (t, *J* = 1.7 Hz, 2H), 4.47 (t, *J* = 1.7 Hz, 2H), 3.24 (t, *J* = 6.7 Hz, 2H), 2.40 (t, *J* = 6.7 Hz, 2H), 1.46 (bs, 9H). **$^{13}$C NMR** (101 MHz, Chloroform-d) δ 155.8, 89.1, 79.1, 70.8, 70.6, 69.6, 41.1, 28.6, 28.5. **ESI-MS**: m/z calcd. for $C_{17}H_{24}NO_2Ru^+$, $[M+H]^+$ 376.0845, found 376.0848.

A mixture of the the *N*-Boc derivative obtained in the previous step (2.6 g, 6.9 mmol) and $LiAlH_4$ (1 g, 27.6 mmol) in anhydrous THF (80 mL) was heated under reflux for 12h. The reaction mixture was cooled in an ice bath and then diluted with methylene chloride, followed by the dropwise addition of a saturated aqueous solution of $Na_2SO_4$ until no further effervescence was observed. The mixture was stirred for 10 min, after which the clear solution was decanted from the white precipitate. The precipitate was washed with DCM and the combined organic solution was concentrated *in vacuo* to give compound **15** as a yellow oil: 1.6 g (78%). **$^1$H NMR** (400 MHz, Chloroform-d) δ 4.53 – 4.51 (m, 7H), 4.45 – 4.44 (m, 2H), 2.68 – 2.65 (m, 2H), 2.44 (s, 3H), 2.39 (t, *J* = 7.1 Hz, 2H). **$^{13}$C NMR** (101 MHz, Chloroform-d) δ 90.3, 70.9, 70.5, 69.5, 53.7, 36.4, 29.2. **ESI-MS**: m/z calcd. for $C_{13}H_{18}NRu^+$, $[M+H]^+$ 290.0477, found 290.0480.

**1-((2-(2,4-difluorophenyl)oxiran-2-yl)methyl)-1H-1,2,4-triazole** (**17**) was prepared following an adapted literature procedure.*(10)* A suspension of NaH (96 mg, 2.4 mmol, 1.2 equiv.) in dry DMSO (16 mL) and trimethylsulfoxonium iodide (528 mg, 2.4 mmol, 1.2 equiv.) was stirred at r.t. until the solution became clear. 1-(2,4-difluorophenyl)-2-(1H-1,2,4-triazol-1-yl)ethan-1-one (446 mg, 2.0 mmol, 1 equiv.) in dry DMSO (32 mL) was added to the previous solution and the reaction mixture was heated to 85°C for 5 h. After cooling the solution to room temperature, the reaction mixture was poured into cold $H_2O$ (50 mL, 4°C). The product was extracted with EtOAc (2 × 50 mL) and the combined organic phases were washed with $H_2O$ (50 mL), brine (50 mL), dried over MgSO4, filtered and evaporated. The residue was purified by flash chromatography on silica with EtOAc:cyclohexane (3:2) as the eluent system (Rf = 0.26, EtOAc:cyclohexane (3:2)) to obtain 1-((2-(2,4-difluorophenyl)oxiran-2-yl)methyl)-1H-1,2,4-triazole 9 an orange oil (313 mg, 1.32 mmol, 66%). **$^1$H NMR** (400 MHz, CDCl3): δ 8.02 (s, 1H), 7.80 (s, 1H), 7.15 – 7.08 (m, 1H), 6.80 – 6.71 (m, 2H), 4.78 (d, J = 14.9 Hz, 1H), 4.45 (d, J = 14.9 Hz, 1H), 2.95 (d, J = 4.7 Hz, 1H), 2.89 (d, J = 4.7 Hz, 1H). The spectral data corresponds to previously reported data.

#### General Procedure for addition to the epoxide 17

Epoxide **17** (1.0 eq.), primary or secondary amine (1.2 eq.) and $Et_3N$ (2-4 eq.) were dissolved in dry EtOH/THF (0.1 M) under nitrogen atmosphere. The reaction mixture was stirred at room temperature for 1 h and then at reflux overnight. The solvent was evaporated under vacuum and the residue purified by column chromatography on silica.

**1-((ruthenocenylmethyl)amino)-2-(2,4-difluorophenyl)-3-(1H-1,2,4-triazol-1-yl)propan-2-ol** (**20**) was prepared with **17**, **4**, and 2.0 equiv. of $Et_3N$, THF as solvent, and purified by flash chromatography with EtOAc as eluent (Rf = 0.21) to obtain expected compound as a brown oil with 10% yield.**$^1$H NMR** (400 MHz, CDCl3): δ 8.13 (s, 1H), 7.82 (s, 1H), 7.58 (td, J = 8.9, 6.6 Hz, 1H), 6.89 – 6.78 (m, 2H), 4.63 (d, J = 14.3 Hz, 1H), 4.53 (d, J = 14.3 Hz, 1H), 4.52 – 4.49 (m, 2H), 4.45 (t, J = 1.7 Hz, 2H), 4.37 (s, 5H), 3.29 (dd, J = 12.3, 1.6 Hz, 1H), 3.21 (d, J = 12.9 Hz, 1H), 3.10 (d, J = 12.9 Hz, 1H), 2.83 (dd, J = 12.3, 0.8 Hz, 1H). **$^{13}$C NMR** (101 MHz, CDCl3): δ 163.0 (dd, J = 250.4, 12.1 Hz), 159.1 (dd, J = 247.1, 11.8 Hz), 151.43, 144.86, 130.18 (dd, J = 9.1, 6.1 Hz), 125.1

(dd, J = 13.2, 3.4 Hz), 111.8 (dd, J = 20.6, 2.8 Hz), 104.5 (t, J = 26.4 Hz), 91.3, 73.6 (d, J = 5.6 Hz), 71.0 (d, J = 12.8 Hz), 70.45 (d, J = 6.3 Hz), 70.1 (d, J = 11.6 Hz), 56.0 (d, J = 4.8 Hz), 54.3 (d, J = 3.8 Hz), 48.1, 29.8.**$^{19}$F NMR** (376 MHz, CDCl3) δ -108.8 (q, J = 9.5 Hz), -110.2 (p, J = 7.9 Hz). **ESI-MS**: m/z calcd. for $C_{22}H_{22}F_2N_4ORuH^+$, $[M-Cl]^+$ 499.0878, found 499.0887.

**1-((ruthenocenylmethyl)(methyl)amino)-2-(2,4-difluorophenyl)-3-(1H-1,2,4-triazol-1-yl)propan-2-ol** (**21**) was prepared with **17**, **5**, and 3.5 eq. of $Et_3N$, EtOH as solvent, and purified by flash chromatography with an eluent system of EtOAc:DCM (3:2, Rf = 0.66) to obtain expected compound as a brown oil with 15% yield. **$^1$H NMR** (400 MHz, CDCl3): δ 8.11 (s, 1H), 7.76 (s, 1H), 7.60 – 7.47 (m, 1H), 6.79 (m, 2H), 4.49 – 4.38 (m, 11H), 3.05 (d, J = 13.4 Hz, 1H), 2.98 (dd, J = 13.5, 1.7 Hz, 2H), 2.96 (d, J = 13.4 Hz, 2H), 2.71 (d, J = 13.6 Hz, 1H), 2.00 (s, 3H). **$^{13}$C NMR** (101 MHz, CDCl3): δ 162.8 (dd, J = 249.6, 12.1 Hz), 159.0 (dd, J = 246.9, 11.8 Hz) 151.0, 144.8, 129.7 (dd, J = 9.1, 6.1 Hz), 126.5 (dd, J = 12.9, 3.4 Hz), 111.5 (dd, J = 20.5, 2.7 Hz), 104.2 (t, J = 26.3 Hz), 86.1, 72.6, 72.4, 71.9, 70.7, 70.5, 70.4, 60.4 (d, J = 3.3 Hz), 58.2, 56.4 (d, J = 4.7 Hz), 43.5. **$^{19}$F NMR** (376 MHz, CDCl3): δ -108.5 (dd, J = 18.2, 9.0 Hz), -110.9 (p, J = 7.5 Hz). **ESI-MS**: m/z calcd. for $C_{23}H_{24}F_2N_4ORuH^+$, $[M-Cl]^+$ 513.1034, found 513.1039.

**1-((Ferrocenylethyl)amino)-2-(2,4-difluorophenyl)-3-(1H-1,2,4-triazol-1-yl)propan-2-ol (22)** was prepared with **17**, **10**, 2 eq. of $Et_3N$, THF as solvent, and purified by column chromatography on silica using EtOAc : hexane (2 : 1→4 : 1→EA) as the eluent (Rf = 0.23, EtOAc) to obtain expected compound as an orange oil. Yield: 65 % (2.0 g, 4.3 mmol). **$^1$H NMR** (400 MHz, Chloroform-d) δ 8.11 (s, 1H), 7.80 (s, 1H), 7.56 – 7.50 (m, 1H), 6.84 – 6.75 (m, 2H), 4.59 (d, *J* = 14.2 Hz, 1H), 4.48 (d, *J* = 14.2 Hz, 1H), 4.05 (s, 5H), 4.03 (t, *J* = 1.8 Hz, 2H), 3.95 (t, *J* = 1.8 Hz, 2H), 3.13 (dd, *J* = 12.5, 1.5 Hz, 1H), 2.82 (dd, *J* = 12.6, 1.4 Hz, 1H), 2.63 – 2.52 (m, 2H), 2.40 - 2.36 (m, 2H). **$^{13}$C NMR** (101 MHz, Chloroform-d) δ 162.7 (dd, J = 249.6, 12.1 Hz), 159.0 (dd, J = 247.1, 11.6 Hz), 151.3, 144.7, 129.9 (dd, J = 9.5, 6.1 Hz), 125.2 (dd, J = 13.3, 3.6 Hz), 111.5 (dd, J = 20.5, 3.3 Hz), 104.3 (t, J = 26.4 Hz), 85.7, 73.1 (d, J = 5.4 Hz), 68.6, 68.3 (d, J = 11.3 Hz), 67.5 (d, J = 2.6 Hz), 56.0 (d, J = 5.1 Hz), 54.1 (d, J = 4.2 Hz), 51.2, 30.2. **$^{19}$F NMR** (376 MHz, Chloroform-d) δ -108.9 (d, *J* = 7.8 Hz), -110.7 (d, *J* = 7.9 Hz). **ESI-MS**: m/z calcd. for $C_{23}H_{25}F_2FeN_4O^+$, $[M+H]^+$ 467.1340, found 467.1344.

**1-((Ferrocenylethyl)(methyl)amino)-2-(2,4-difluorophenyl)-3-(1H-1,2,4-triazol-1-yl)propan-2-ol (23)** was prepared with **17**, **14**, 3.5 eq. of $Et_3N$, EtOH as solvent, and purified by column chromatography on silica using EtOAc : hexane (1 : 1) as the eluent (Rf = 0.23, EtOAc : hexane = 1 : 1) to obtain expected compound as an orange oil. Yield: 70% (1.2 g, 2.5 mmol). **$^1$H NMR**

(500 MHz, Chloroform-d) δ 8.13 (s, 1H), 7.77 (s, 1H), 7.47 – 7.42 (m, 1H), 6.81 – 6.76 (m, 2H), 4.46 (d, *J* = 14.2 Hz, 1H), 4.41 (d, *J* = 14.2 Hz, 1H), 4.06 (bs, 7H), 3.95 – 3.91 (m, 2H), 3.01 (dd, *J* = 13.6, 1.8 Hz, 1H), 2.69 (d, *J* = 13.6 Hz, 1H), 2.42 – 2.34 (m, 2H), 2.31 - 2.28 (m, 2H), 2.06 (s, 3H). **$^{13}$C NMR** (101 MHz, Chloroform-d) δ 162.7 (dd, J = 249.3, 12.0 Hz), 158.9 (dd, J = 246.6, 11.9 Hz), 150.9, 144.8, 129.6 (dd, J = 9.4, 6.0 Hz), 126.3 (d, J = 14.4 Hz), 111.4 (dd, J = 20.7, 3.3 Hz), 104.1 (t, J = 26.3 Hz), 85.7, 71.8 (d, J = 5.7 Hz), 68.5, 68.2 (d, J = 9.3 Hz), 67.5 (d, J = 11.4 Hz), 62.1 (d, J = 4.0 Hz), 60.3, 56.3 (d, J = 5.2 Hz), 43.5, 27.7. **$^{19}$F NMR** (471 MHz, Chloroform-d) δ -108.7 (d, *J* = 7.6 Hz), -111.0 (d, *J* = 7.9 Hz). **ESI-MS**: m/z calcd. for $C_{24}H_{27}F_2FeN_4O^+$, $[M+H]^+$ 481.1497, found 481.1497.

**1-((Ruthenocenylethyl)amino)-2-(2,4-difluorophenyl)-3-(1H-1,2,4-triazol-1-yl)propan-2-ol (24)** was prepared with **17**, **11**, 3 eq. of $Et_3N$, THF as solvent, and purified by column chromatography on silica using EtOAc : hexane (2 : 1→4 : 1→EA) as the eluent (Rf = 0.2, EtOAc) to obtain expected compound as a white solid . Yield: 40 % (0.25 g, 0.5 mmol). **$^1$H NMR** (400 MHz, Chloroform-d) δ 8.12 (s, 1H), 7.81 (s, 1H), 7.59 – 7.52 (m, 1H), 6.86 – 6.77 (m, 2H), 4.61 (d, *J* = 14.2 Hz, 1H), 4.51 (d, *J* = 14.2 Hz, 1H), 4.44 (s, 5H), 4.41 – 4.39 (m, 4H), 3.17 (dd, *J* = 12.5, 1.5 Hz, 1H), 2.82 (dd, *J* = 12.6, 1.4 Hz, 1H), 2.57 – 2.53 (m, 2H), 2.26 – 2.22 (m, 2H). **$^{13}$C NMR** (101 MHz, Chloroform-d) δ 162.8 (dd, J = 249.5, 12.1 Hz), 159.0 (dd, J = 247.2, 11.8 Hz), 151.3, 144.7, 129.9 (dd, J = 9.1, 6.3 Hz), 125.2 (d, J = 16.6 Hz), 111.5 (dd, J = 20.5, 2.8 Hz), 104.3 (t, J = 26.5 Hz), 89.6, 73.1 (d, J = 5.4 Hz), 70.9 (d, J = 10.3 Hz), 70.5, 69.6, 56.0 (d, J = 5.3 Hz), 54.1 (d, J = 4.1 Hz), 51.9, 29.4. **$^{19}$F NMR** (376 MHz, Chloroform-d) δ -108.9 – -109.0 (m), -110.6 – 110.7 (m). **ESI-MS**: m/z calcd. for $C_{23}H_{25}F_2N_4ORu^+$, $[M+H]^+$ 513.1034, found 513.1039.

**1-((Ruthenocenylethyl)(methyl)amino)-2-(2,4-difluorophenyl)-3-(1H-1,2,4-triazol-1-yl)propan-2-ol (25)** was prepared with **17**, **15**, 3 eq. of $Et_3N$, EtOH as solvent, and purified by column chromatography on silica using EtOAc : hexane (3 : 2) as the eluent (Rf = 0.32, EtOAc : hexane = 3 : 2) to expected compound as a white solid . Yield: 85 % (2.23 g, 4.3 mmol). **$^1$H NMR** (400 MHz, Chloroform-d) δ 8.16 (s, 1H), 7.78 (s, 1H), 7.55 – 7.49 (m, 1H), 6.84 – 6.77 (m, 2H), 5.24 (br, 1H), 4.53 – 4.42 (m, 9H), 4.40 – 4.39 (m, 1H), 4.35 – 4.34 (m, 1H), 3.04 (dd, J = 13.6, 1.8 Hz, 1H), 2.70 (d, J = 13.6 Hz, 1H), 2.39 – 2.35 (m, 2H), 2.18 – 2.14 (m, 2H), 2.06 (s, 3H). **$^{13}$C NMR** (101 MHz, Chloroform-d) δ 162.7 (dd, J = 249.4, 12.1 Hz), 158.9 (dd, J = 246.8, 11.8 Hz), 150.9, 144.8, 129.6 (dd, J = 9.4, 6.0 Hz), 126.4 (dd, J = 13.0, 3.8 Hz), 111.5 (dd, J = 20.4, 3.3 Hz), 104.2 (t, J = 26.3 Hz), 89.3, 71.8 (d, J = 5.6 Hz), 70.8, 70.5, 69.6 (d, J = 8.7 Hz), 62.0 (d, J = 3.9 Hz), 61.0, 56.3 (d, J = 5.2 Hz), 43.5, 27.0. **$^{19}$F NMR** (376 MHz, Chloroform-d) δ -108.6 – -108.7

(m), -110.8 – -110.9 (m). **ESI-MS**: m/z calcd. for $C_{24}H_{27}F_2N_4ORu^+$, $[M+H]^+$ 527.1191, found 527.1194.

**1-((Phenethylethyl)(methyl)amino)-2-(2,4-difluorophenyl)-3-(1H-1,2,4-triazol-1-yl)propan-2-ol (26)** was prepared with **17**, **18**, 3 eq. of $Et_3N$, EtOH as solvent, and purified by column chromatography on silica using EtOAc : hexane (3 : 2) as the eluent (Rf = 0.46, EtOAc : hexane = 3 : 2) to obtain expected compound as a white solid . Yield: 70 % (0.56 g, 1.5 mmol). **$^1$H NMR** (500 MHz, Methanol-d4) δ 8.27 (s, 1H), 7.74 (s, 1H), 7.36 (td, J = 9.0, 6.7 Hz, 1H), 7.26 – 7.23 (m, 2H), 7.18 – 7.14 (m, 1H), 7.07 – 7.05 (m, 2H), 6.93 – 6.89 (m, 1H), 6.85 – 6.81 (m, 1H), 4.58 (dd, J = 14.3, 0.9 Hz, 1H), 4.47 (d, J = 14.3 Hz, 1H), 3.05 (dd, J = 13.8, 1.7 Hz, 1H), 2.84 (d, J = 13.8 Hz, 1H), 2.64 – 2.60 (m, 2H), 2.59 – 2.54 (m, 2H), 2.22 (s, 3H). **$^{13}$C NMR** (126 MHz, Methanol-d4) δ 164.2 (dd, J = 247.5, 12.1 Hz), 160.7 (dd, J = 246.2, 11.9 Hz), 151.1, 146.2, 141.3, 131.0 (dd, J = 9.7, 6.0 Hz), 129.8, 129.4, 127.5 (dd, J = 12.9, 3.9 Hz), 127.1, 112.1 (dd, J = 21.1, 3.5 Hz), 104.9 (t, J = 26.8 Hz), 74.2 (d, J = 5.6 Hz), 64.0 (d, J = 3.8 Hz), 62.0, 57.4 (d, J = 4.9 Hz), 44.1, 34.6. **$^{19}$F NMR** (471 MHz, Methanol-d4) δ -109.41 (d, J = 7.8 Hz), -113.52 (d, J = 7.8 Hz). **ESI-MS**: m/z calcd. for $C_{20}H_{23}F_2N_4O^+$, $[M+H]^+$ 373.1834, found 373.1832

**1-((Adamantylethyl)(methyl)amino)-2-(2,4-difluorophenyl)-3-(1H-1,2,4-triazol-1-yl)propan-2-ol (27)** was prepared with **17**, **19**, 4 eq. of $Et_3N$, EtOH as solvent, and purified by column chromatography on silica using DCM : MeOH (95 : 5) as the eluent (Rf = 0.27) to obtain expected compound as a white solid . Yield: 75 % (0.75 g, 1.8 mmol). **$^1$H NMR** (500 MHz, Methanol-d4) δ 8.37 (s, 1H), 7.77 (s, 1H), 7.53 (td, J = 8.9, 6.6 Hz, 1H), 6.96 – 6.92 (m, 1H), 6.89 – 6.85 (m, 1H), 4.62 (d, J = 14.4 Hz, 1H), 4.55 (d, J = 14.2 Hz, 1H), 3.01 (dd, J = 13.7, 1.8 Hz, 1H), 2.76 (d, J = 13.7 Hz, 1H), 2.31 – 2.27 (m, 2H), 2.12 (s, 3H), 1.88 – 1.87 (m, 3H), 1.72 – 1.69 (m, 3H), 1.63 – 1.60 (m, 3H), 1.37 – 1.36 (m, 6H), 1.11 – 0.98 (m, 2H). **$^{13}$C NMR** (126 MHz, Methanol-d4) δ 164.3 (dd, J = 247.6, 12.2 Hz), 160.7 (dd, J = 246.3, 11.9 Hz), 151.1, 146.3, 131.0 (dd, J = 9.6, 6.0 Hz), 127.8 (dd, J = 12.9, 3.7 Hz), 112.1 (dd, J = 20.3, 3.3 Hz), 105.0 (t, J = 26.8 Hz), 73.8 (d, J = 5.8 Hz), 63.6 (d, J = 3.7 Hz), 57.6 (d, J = 5.3 Hz), 54.2, 44.6, 43.5, 42.4, 38.2, 32.6, 30.1. **$^{19}$F NMR** (376 MHz, Methanol-d4) δ -109.3 – -109.4 (m), -113.5 – -113.6 (m). **ESI-MS**: m/z calcd. for $C_{24}H_{33}F_2N_4O^+$, $[M+H]^+$ 431.2617, found 431.2614.

#### General Procedure for hydrochloride salt preparation

Compounds **20** – **27** were dissolved in deoxygenated acetone (0.2 M), and aq. HCl (37% in $H_2O$, 1 equiv.) was added. The reaction was stirred at r.t. (25°C) for 1 h, and the precipitate was filtered. The residue was washed with acetone and $Et_2O$, and dried *in vacuo* to obtain **5a** – **12a** as powder.

**1-((Ruthenocenylmethyl)amino)-2-(2,4-difluorophenyl)-3-(1H-1,2,4-triazol-1-yl)propan-2-ol hydrochloride salt (5a)** as white amorphous solid. Yield: 65 % (0.52 g, 0.97 mmol). **IR** ($cm^{-1}$): 3107, 1616, 1500, 1394, 1269, 1139, 968, 853, 674, 655. **$^1$H NMR** (500 MHz, Methanol-d4) δ 8.40 (s, 1H), 7.95 (s, 1H), 7.56 (td, J = 9.0, 6.3 Hz, 1H), 7.11 – 6.97 (m, 2H), 4.82 – 4.69 (m, 4H), 4.69 – 4.64 (m, 2H), 4.60 (s, 5H), 3.85 – 3.75 (m, 2H), 3.67 (d, J = 13.1 Hz, 1H), 3.43 (d, J = 13.1 Hz, 1H). **$^{13}$C NMR** (126 MHz, Methanol-d4) δ 165.2 (dd, J = 249.9, 12.3 Hz), 161.0 (dd, J = 247.3, 12.4 Hz), 151.7, 146.4, 131.4 (dd, J = 9.9, 5.0 Hz), 123.0 (d, J = 12.5 Hz), 113.0 (d, J = 21.1 Hz), 105.9 (t, J = 26.9 Hz), 80.2, 73.7, 73.6 (d, J = 5.6 Hz), 72.7, 72.3, 56.7 (d, J = 5.5 Hz), 52.5 (d, J = 5.6 Hz), 49.0 (overlapping with solvent residual peak identified in the HSQC spectrum). **$^{19}$F NMR** (376 MHz, Methanol-d4) δ -109.0 – -109.1 (m), -110.7– -110.8 (m). **ESI-MS**: m/z calcd. for $C_{22}H_{22}F_2N_4ORuH^+$, $[M-Cl]^+$ 499.0878, found 499.0887. **Elemental Analysis**: calcd. for $C_{22}H_{22}ClF_2N_4ORu \cdot 0.5H_2O$= C, 48.71; H, 4.37; N, 10.33. Found = C, 48.45; H, 4.25; N,10.14.

**1-((Ruthenocenylmethyl)(methyl)amino)-2-(2,4-difluorophenyl)-3-(1H-1,2,4-triazol-1-yl)propan-2-ol hydrochloride salt (6a)** as a white amorphous solid. Yield: 60 % (0.31 g, 0.60 mmol). **IR** ($cm^{-1}$): 3024, 1615, 1499, 1392, 1273, 1116, 966, 855, 817, 724. **$^1$H NMR** (500 MHz, Methanol-d4) δ 9.62 – 9.54 (m, 1H), 8.65 – 8.62 (m, 1H), 7.62 – 7.55 (m, 1H), 7.18 – 7.04 (m, 2H), 5.00 – 4.92 (m, 2H), 4.82 – 4.68 (m, 3H), 4.63 (s, 2.5H), 4.62 – 4.58 (m, 1H), 4.54 (s, 2.5H), 4.13 (dd, J = 13.7, 4.5 Hz, 1H), 4.04 – 3.91 (m, 1H), 3.80 (s, 1H), 3.69 – 3.59 (m, 1H), 2.95 (s, 1.5H), 2.52 (s, 1.5H). **$^{13}$C NMR** (126 MHz, Methanol-d4) δ 165.3 (dd, J = 252.4, 10.4 Hz), 160.8 (dd, J = 247.2, 11.9 Hz), 146.4 – 145.9 (m), 145.3 – 144.7 (m), 131.5 – 130.9 (m), 123.4 – 121.9 (m), 113.6 – 113.1 (m), 106.6 – 105.8 (m), 78.3 – 78.1 (m), 74.8 – 73.7 (m), 73.4 – 72.9 (m), 72.6 – 72.3 (m), 71.6 – 71.5 (m), 60.5 – 60.4 (m), 59.7 – 59.5 (m), 58.5 – 58.2 (m), 43j.8 (d, J = 185.9 Hz). **$^{19}$F NMR** (471 MHz, Methanol-d4) δ -107.3 – -109.5 (m), -109.7 – -110.6 (m). **ESI-MS**: m/z calcd. for $C_{23}H_{24}F_2N_4ORuH^+$, $[M-Cl]^+$ 513.1034, found 513.1039. **Elemental Analysis**: calcd. for $C_{23}H_{24}ClF_2N_4ORu \cdot H_2O$= C, 48.81; H, 4.81; N, 9.90. Found = C, 48.55; H, 4.66; N,9.51.

**1-((Ferrocenylethyl)amino)-2-(2,4-difluorophenyl)-3-(1H-1,2,4-triazol-1-yl)propan-2-ol hydrochloride salt (7a)** as an orange amorphous solid. Yield: 63 % (1.31 g, 2.6 mmol). **IR** ($cm^-$

[1]): 3096, 1617, 1501, 1422, 1268, 1140, 968, 817, 778, 676. **$^1$H NMR** (400 MHz, Methanol-d4) δ 9.54 (s, 1H), 8.52 (s, 1H), 7.47 – 7.41 (m, 1H), 7.06 – 7.00 (m, 1H), 6.94 – 6.89 (m, 1H), 4.94 (d, J = 14.5 Hz, 1H), 4.87 (d, J = 14.5 Hz, 1H), 4.06 (bs, 9H), 3.73 (d, J = 13.2 Hz, 1H), 3.55 (d, J = 13.2 Hz, 1H), 3.14 – 3.06 (m, 2H), 2.66 – 2.62 (m, 2H). **$^{13}$C NMR** (101 MHz, Methanol-d4) δ 163.9 (dd, *J =* 250.2, 12.3 Hz), 159.7 (dd, *J =* 247.4, 12.1 Hz), 144.4, 143.3, 130.1 – 121.9 (m), 120.7 (d, *J =* 13.5 Hz), 111.7 (d, *J =* 20.5 Hz), 104.5 (t, *J =* 26.8 Hz), 82.8, 71.9 (d, *J =* 4.2 Hz), 68.5, 67.8, 67.7, 56.9 (d, *J =* 5.7 Hz), 52.6 (d, *J =* 5.6 Hz), 49.2, 25.2. **$^{19}$F NMR** (376 MHz, Methanol-d4) δ -108.8 – -108.9 (m), -110.3 – -110.4 (m). **ESI-MS**: m/z calcd. for $C_{23}H_{25}F_2FeN_4O^+$, $[M-Cl]^+$ 467.1340, found 467.1338. **Elemental Analysis**: calcd. for $C_{23}H_{25}ClF_2FeN_4O \cdot H_2O$= C, 53.05; H, 5.23; N, 10.76. Found = C, 52.71; H, 5.20; N,10.34.

**1-((Ferrocenylethyl)(methyl)amino)-2-(2,4-difluorophenyl)-3-(1H-1,2,4-triazol-1-yl)propan-2-ol hydrochloride salt (8a)** as an orange amorphous solid. Yield: 70 % (0.90 g, 1.76 mmol). **IR** ($cm^{-1}$): 3025, 1616, 1499, 1428, 1293, 1119, 966, 854, 779, 663. **$^1$H NMR** (500 MHz, Methanol-d4) δ 9.53 (d, J = 6.1 Hz, 1H), 8.57 (d, J = 7.7 Hz, 1H), 7.63 – 7.55 (m, 1H), 7.24 – 7.15 (m, 1H), 7.11 – 7.04 (m, 1H), 5.06 – 4.93 (m, 2H), 4.29 – 4.02 (m, 9H), 3.97 – 3.76 (m, 2H), 3.45 – 3.31 (m, 1H), 3.17 – 3.00 (m, 2.5H), 2.73 – 2.46 (m, 3.5H). **$^{13}$C NMR** (101 MHz, DMSO-d6) δ 164.4 – 161.7 (m), 160.8 – 158.1 (m), 149.9, 145.6, 130.9 – 130.6 (m), 123.4 (dt, J = 18.1, 7.1 Hz), 112.2 (d, J = 20.9 Hz), 105.1 (t, J = 26.5 Hz), 83.5 (d, J = 33.3 Hz), 72.5 (d, J = 48.7 Hz), 68.9 (d, J = 7.9 Hz), 68.1 (d, J = 5.2 Hz), 67.8, 60.4 (d, J = 23.4 Hz), 57.9 (d, J = 115.4 Hz), 55.9 (d, J = 11.7 Hz), 43.2 (d, J = 84.6 Hz), 23.4 (d, J = 15.4 Hz). **$^{19}$F NMR** (376 MHz, DMSO-d6) δ -107.3 – -107.4 (m), -110.0– -110.1 (m). **ESI-MS**: m/z calcd. for $C_{24}H_{27}F_2FeN_4O^+$, $[M-Cl]^+$ 481.1497, found 481.1493. **Elemental Analysis**: calcd. for $C_{24}H_{27}ClF_2FeN_4O \cdot 2H_2O$= C, 52.14; H, 5.65; N, 10.13. Found = C, 52.14; H, 5.37; N,10.13.

**1-((Ruthenocenylethyl)amino)-2-(2,4-difluorophenyl)-3-(1H-1,2,4-triazol-1-yl)propan-2-ol hydrochloride salt (9a)** as a white amorphous solid. Yield: 73 % (1.26 g, 2.3 mmol). **IR** ($cm^{-1}$): 3077, 1617, 1501, 1420, 1274, 1100, 967, 804, 678. **$^1$H NMR** (500 MHz, Methanol-d4) δ 9.39 (s, 1H), 8.49 (s, 1H), 7.57 – 7.52 (m, 1H), 7.15 – 7.11 (m, 1H), 7.04 – 7.00 (m, 1H), 4.97 (d, J = 14.6 Hz, 1H), 4.93 (d, J = 14.6 Hz, 1H), 4.56 – 4.55 (m, 7H), 4.50 (dd, J = 1.9, 1.4 Hz, 2H), 3.83 (d, J = 13.2 Hz, 1H), 3.62 (d, J = 13.2 Hz, 1H), 3.17 – 3.07 (m, 2H), 2.57 (t, J = 8.5 Hz, 2H). **$^{13}$C NMR** (126 MHz, Methanol-d4) δ 165.3 (dd, J = 250.4, 12.5 Hz), 161.1 (dd, J = 247.3, 12.4 Hz), 147.0, 145.0, 131.4 (dd, J = 9.5, 4.7 Hz), 122.2 (d, J = 12.6 Hz), 113.1 (d, J = 20.8 Hz), 106.0 (t, J = 26.8 Hz), 87.7, 73.4 (d, J = 4.3 Hz), 71.7, 71.5 (d, J = 5.3 Hz), 70.8, 58.0 (d, J = 5.8 Hz), 54.0 (d,

J = 5.6 Hz), 51.6, 25.7. **$^{19}$F NMR** (376 MHz, Methanol-d4) δ -108.9 (d, J = 8.8 Hz), -110.4 (d, J = 9.1 Hz). **ESI-MS**: m/z calcd. for $C_{23}H_{25}F_2N_4ORu^+$, $[M-Cl]^+$ 513.1034, found 513.1038. **Elemental Analysis**: calcd. for $C_{23}H_{25}ClF_2N_4ORu \cdot H_2O$= C, 48.81; H, 4.81; N, 9.90. Found = C, 48.75; H, 4.39; N,9.27.

**1-((Ruthenocenylethyl)(methyl)amino)-2-(2,4-difluorophenyl)-3-(1H-1,2,4-triazol-1-yl)propan-2-ol hydrochloride salt (10a)** as a white amorphous solid. Yield: 65 % (1.4 g, 2.5 mmol). **IR** (cm$^{-1}$): 3036, 1704, 1618, 1503, 1420, 1275, 1109, 971, 851, 739. **$^1$H NMR** (500 MHz, Methanol-d4) δ 9.67 (d, J = 2.7 Hz, 1H), 8.64 (d, J = 6.9 Hz, 1H), 7.64 – 7.55 (m, 1H), 7.24 – 7.15 (m, 1H), 7.11 – 7.04 (m, 1H), 5.09 – 4.93 (m, 2H), 4.62 – 4.51 (m, 7H), 4.47 – 4.38 (m, 2H), 4.16 – 3.77 (m, 2H), 3.44 – 3.37 (m, 1H, overlapping with solvent residual peak identified in the HSQC spectrum), 3.13 – 3.06 (m, 0.5H), 3.04 (s, 1.5H), 3.01 – 2.95 (m, 0.5H), 2.68 – 2.62 (m, 1H), 2.60 (s, 1.5H), 2.55 – 2.34 (m, 1H). **$^{13}$C NMR** (126 MHz, Methanol-d4) δ 165.4 (dd, J = 254.2, 11.2 Hz), 160.9 (dd, J = 247.2, 12.0 Hz), 145.8 (d, J = 10.9 Hz), 144.8 (d, J = 9.8 Hz), 131.5 – 131.1 (m), 122.6 – 122.4 (m), 113.5 (t, J = 19.8 Hz), 106.4 – 106.0 (m), 87.1 (d, J = 61.6 Hz), 73.4 (dd, J = 52.2, 4.2 Hz), 71.7 (d, J = 8.2 Hz), 71.5 (d, J = 17.0 Hz), 70.9 (d, J = 4.9 Hz), 61.9, 61.3 (d, J = 149.0 Hz), 58.5 (dd, J = 22.1, 5.4 Hz), 44.2 (d, J = 140.2 Hz), 23.9 (d, J = 7.3 Hz). **$^{19}$F NMR** (376 MHz, Methanol-d4) δ -108.6 – -108.7 (m), -109.4 – -109.7 (m). **ESI-MS**: m/z calcd. for $C_{24}H_{27}F_2N_4ORu^+$, $[M-Cl]^+$ 527.1191, found 527.1191. **Elemental Analysis**: calcd. for $C_{24}H_{27}ClF_2N_4ORu \cdot 2.5\ H_2O$= C, 47.49; H, 5.31; N, 9.23. Found = C, 47.76; H, 4.93; N, 8.95.

**1-((Phenethyl)(methyl)amino)-2-(2,4-difluorophenyl)-3-(1H-1,2,4-triazol-1-yl)propan-2-ol hydrochloride salt (11a)** as a white amorphous solid. Yield: 46 % (0.4 g, 0.86 mmol). **IR** (cm$^{-1}$): 2968, 1618, 1503, 1418, 1274, 1106, 968, 846, 757. **$^1$H NMR** (500 MHz, Methanol-d4) δ 9.60 (d, J = 2.4 Hz, 1H), 8.60 (d, J = 4.9 Hz, 1H), 7.63 – 7.57 (m, 1H), 7.36 – 7.22 (m, 4H), 7.20 – 7.14 (m, 1H), 7.11 – 7.04 (m, 2H), 5.09 – 4.95 (m, 2H), 4.24 – 3.79 (m, 2H), 3.55 – 3.40 (m, 1H), 3.24 – 3.05 (m, 3.5 H), 2.98 – 2.79 (m, 1H), 2.65 (s, 1.5 H). **$^{13}$C NMR** (126 MHz, Methanol-d4) δ 165.4 (dd, J = 249.3, 8.4 Hz), 160.9 (dd, J = 247.4, 12.3 Hz), 146.2 (d, J = 14.2 Hz), 144.9 (d, J = 9.3 Hz), 137.1 (d, J = 47.7 Hz), 131.3 (d, J = 26.8 Hz), 130.0 (d, J = 4.9 Hz), 129.7, 128.4, 122.8 – 122.3 (m), 113.7 – 113.4 (m), 106.4 – 105.9 (m), 73.4 (d, J = 54.4 Hz), 61.9 (dd, J = 15.1, 5.6 Hz), 60.8 (d, J = 148.1 Hz), 58.5 (dd, J = 22.7, 5.2 Hz), 44.2 (d, J = 127.0 Hz), 30.9 (d, J = 26.7 Hz). **$^{19}$F NMR** (471 MHz, Methanol-d4) δ -108.7 – -108.8 (m), -109.6 – -109.7 (m). **ESI-MS**: m/z calcd. for $C_{20}H_{23}F_2N_4O^+$, $[M+H]^+$ 373.1834, found 373.1832 **Elemental Analysis**: calcd. for $C_{20}H_{23}ClF_2N_4O \cdot H_2O \cdot HCl$= C, 51.84; H, 5.66; N, 12.09; Found = C, 51.35; H, 5.58; N, 11.91.

**1-((Adamantylethyl)(methyl)amino)-2-(2,4-difluorophenyl)-3-(1H-1,2,4-triazol-1-yl)propan-2-ol hydrochloride salt (12a)** as a white amorphous solid. Yield: 44 % (0.62 g, 1.23 mmol). **IR** ($cm^{-1}$): 2900, 1500, 1416, 1270, 1221, 1130, 947, 846, 800. **$^1$H NMR** (500 MHz, Methanol-d4) δ 9.66 (d, J = 13.2 Hz, 1H), 8.64 (d, J = 5.1 Hz, 1H), 7.62 –7.55 (m, 1H), 7.23 – 7.14 (m, 1H), 7.09 – 7.03 (m, 1H), 5.06 – 4.89 (m, 2H), 4.11 – 3.71 (m, 2H), 3.26 – 3.20 (m, 1H, overlapping with solvent residual peak identified in the HSQC spectrum), 2.99 (s, 1.5H), 2.96 – 2.87 (m, 1H), 2.54 (s, 1.5H), 1.94 (d, J = 35.8 Hz, 3H), 1.79 – 1.58 (m, 9H), 1.55 – 1.48 (m, 1H), 1.36 (d, J = 2.8 Hz, 3H), 1.34 – 1.16 (m, 1H). **$^{13}$C NMR** (126 MHz, Methanol-d4) δ 165.4 (dd, J = 250.9, 12.8 Hz), 160.9 (dd, J = 247.1, 12.2 Hz), 145.8 (d, J = 23.2 Hz), 144.8 (d, J = 16.1 Hz), 131.5 – 131.1 (m), 122.8 – 122.5 (m), 113.6 – 113.4 (m), 106.4 – 106.0 (m), 73.3 (dd, J = 51.7, 4.2 Hz), 61.9 (dd, J = 34.0, 5.5 Hz), 58.5 (dd, J = 19.4, 5.7 Hz), 55.7 (d, J = 185.3 Hz), 44.2 (d, J = 142.2 Hz), 43.0 (d, J = 17.7 Hz), 38.2 (d, J = 7.2 Hz), 37.8 (d, J = 13.3 Hz), 32.8 (d, J = 32.9 Hz), 29.9 (d, J = 15.1 Hz). **$^{19}$F NMR** (471 MHz, Methanol-d4) δ -108.6 – -108.7 (m), -109.7 – -109.8 (m). **ESI-MS**: m/z calcd. for $C_{24}H_{33}F_2N_4O^+$, $[M+H]^+$ 431.2617, found 431.2614. **Elemental Analysis**: calcd. for $C_{24}H_{33}ClF_2N_4O \cdot HCl$= C, 57.26; H, 6.81; N, 11.13; Found = C, 57.38; H, 6.71; N, 11.11.

#### 10.Infrared spectra

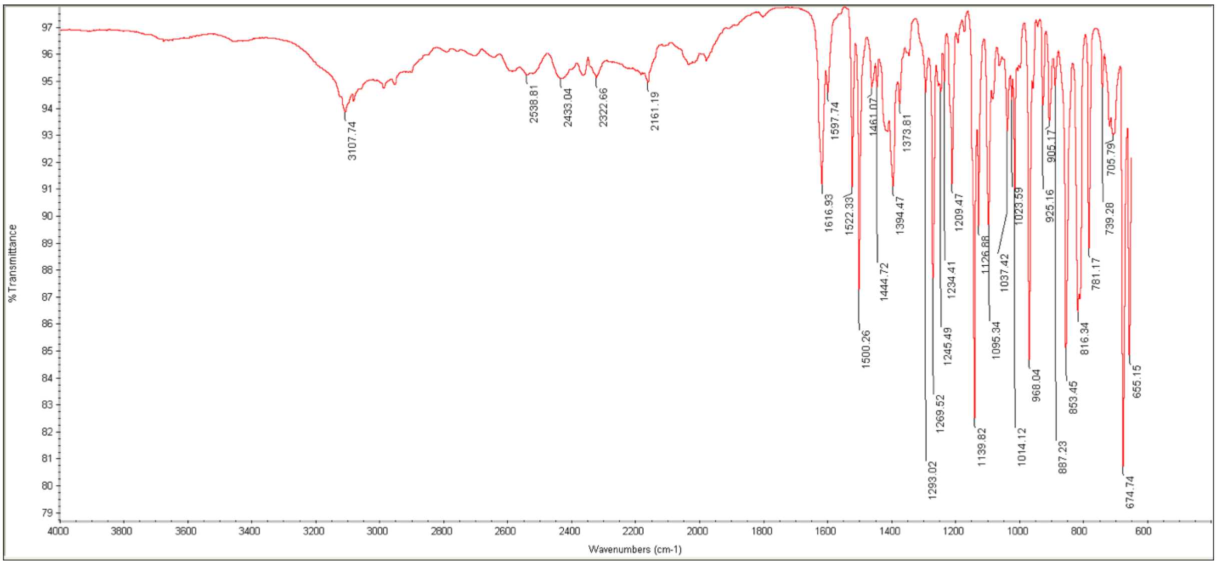

**Figure S41.** Infrared spectra of **5a**.

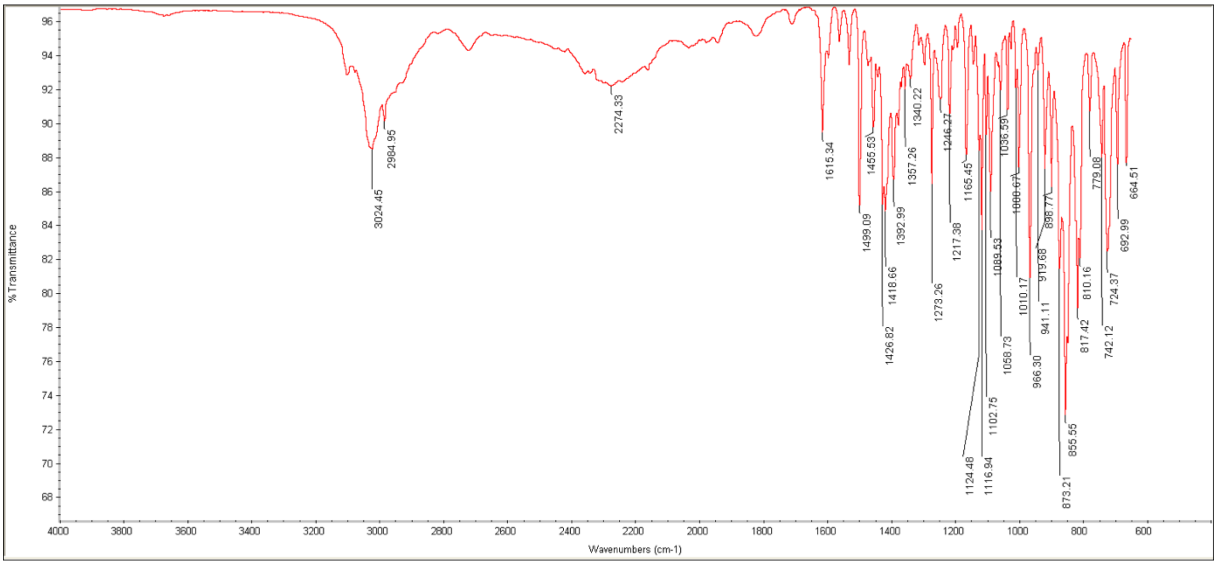

**Figure S42.** Infrared spectra of **6a**.

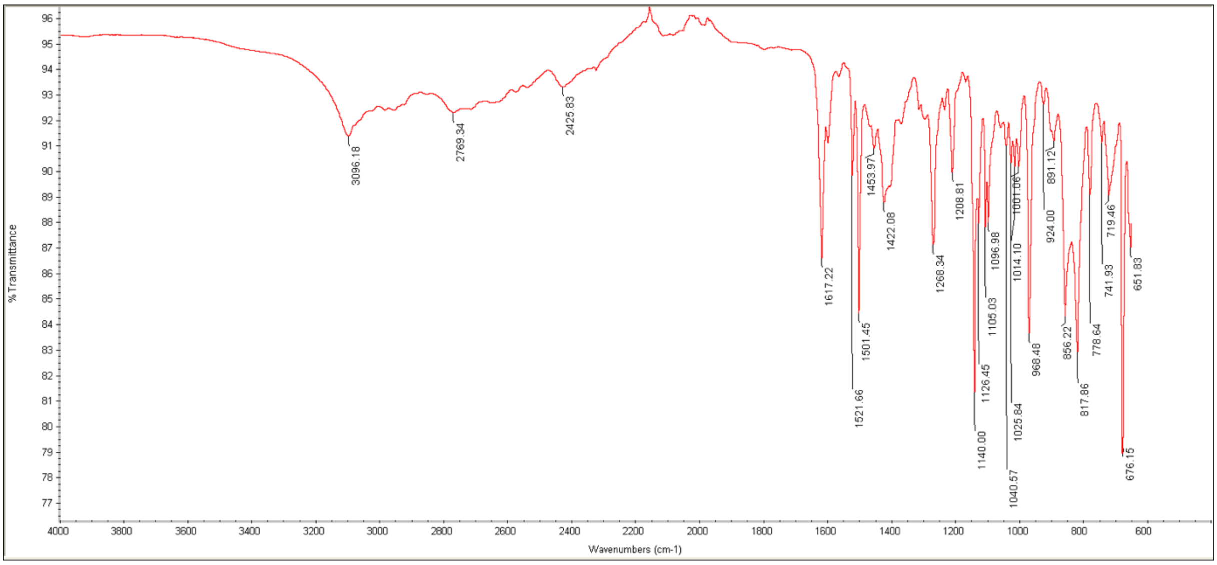

**Figure S43.** Infrared spectra of **7a**.

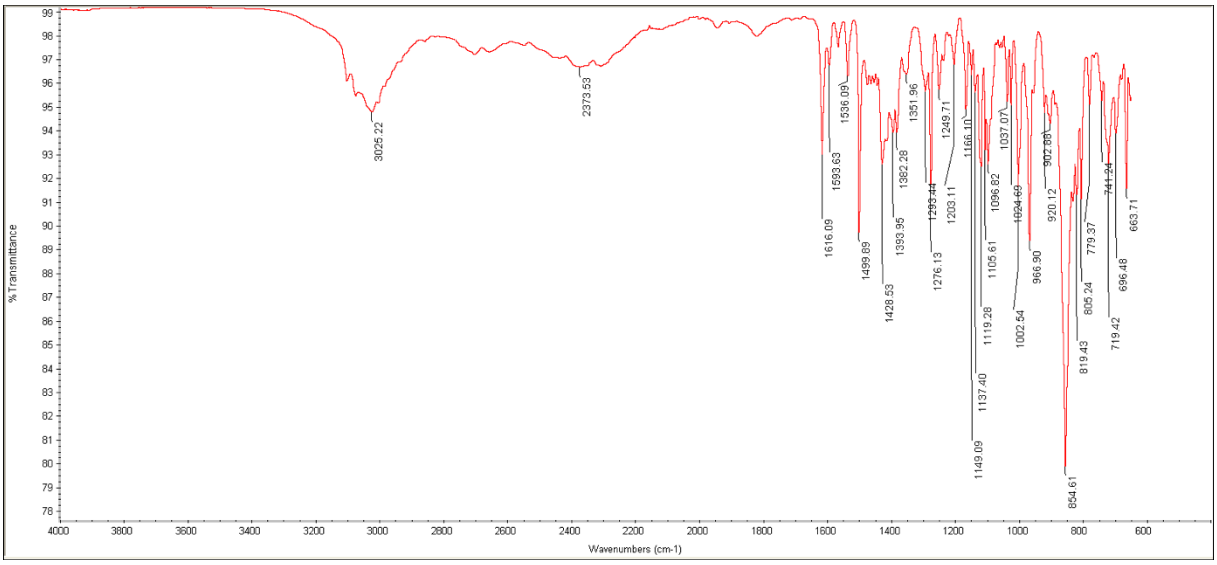

**Figure S44.** Infrared spectra of **8a**.

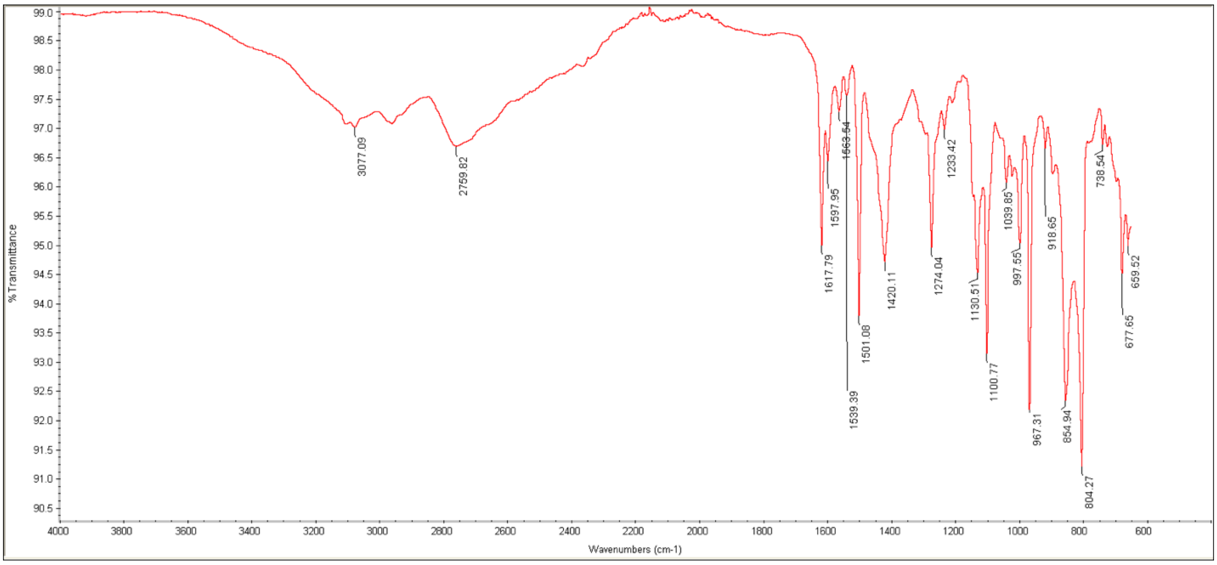

**Figure S45.** Infrared spectra of **9a**.

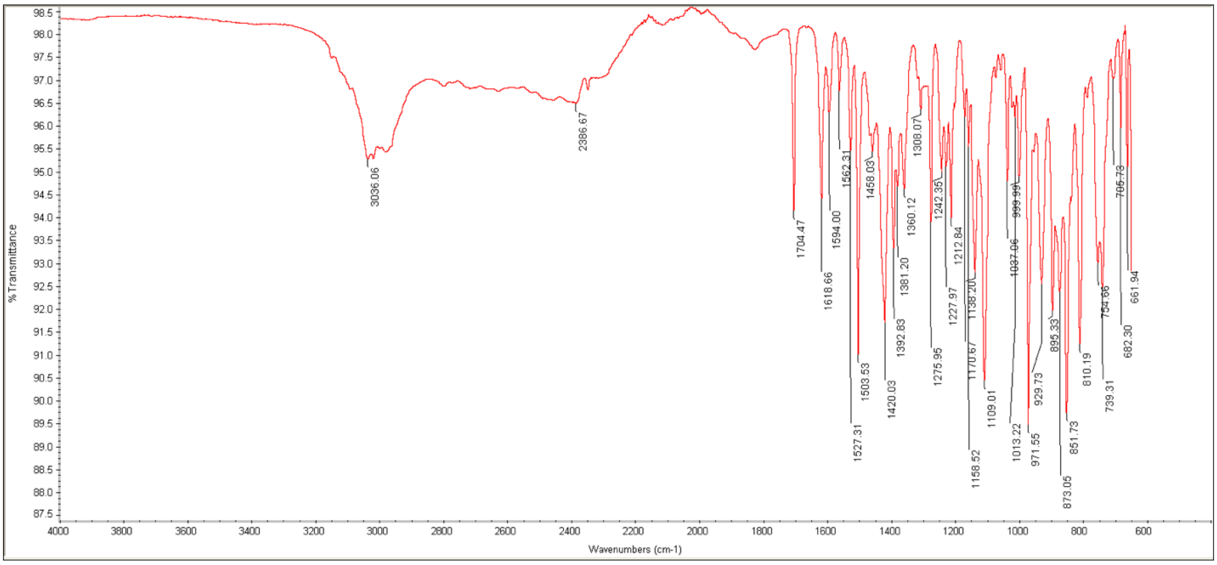

**Figure S46.** Infrared spectra of **10a**.

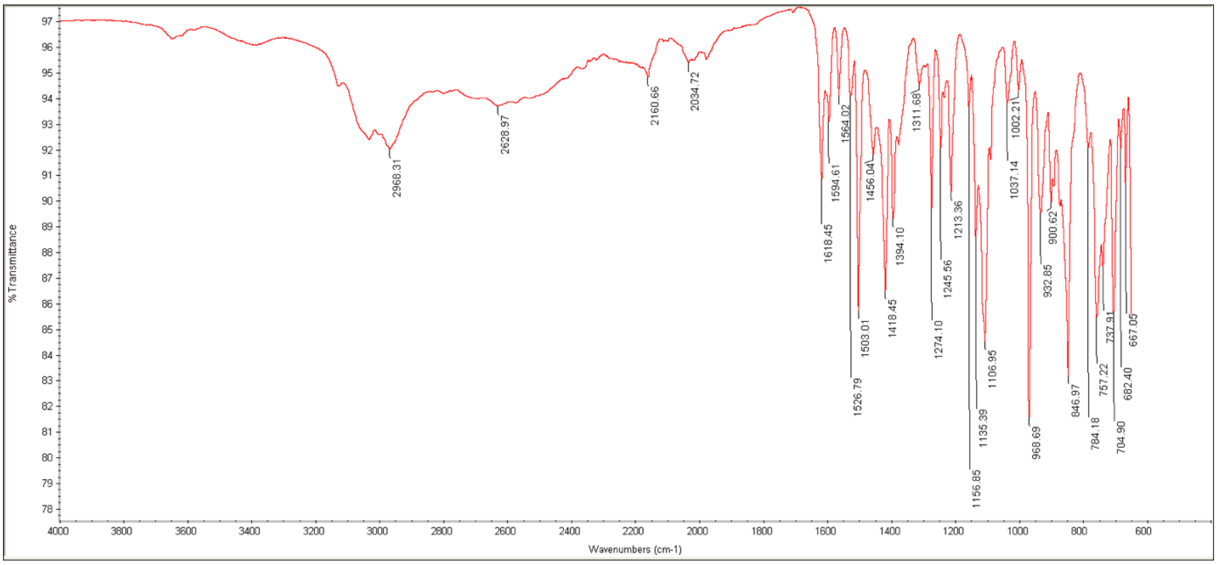

**Figure S47.** Infrared spectra of **11a**.

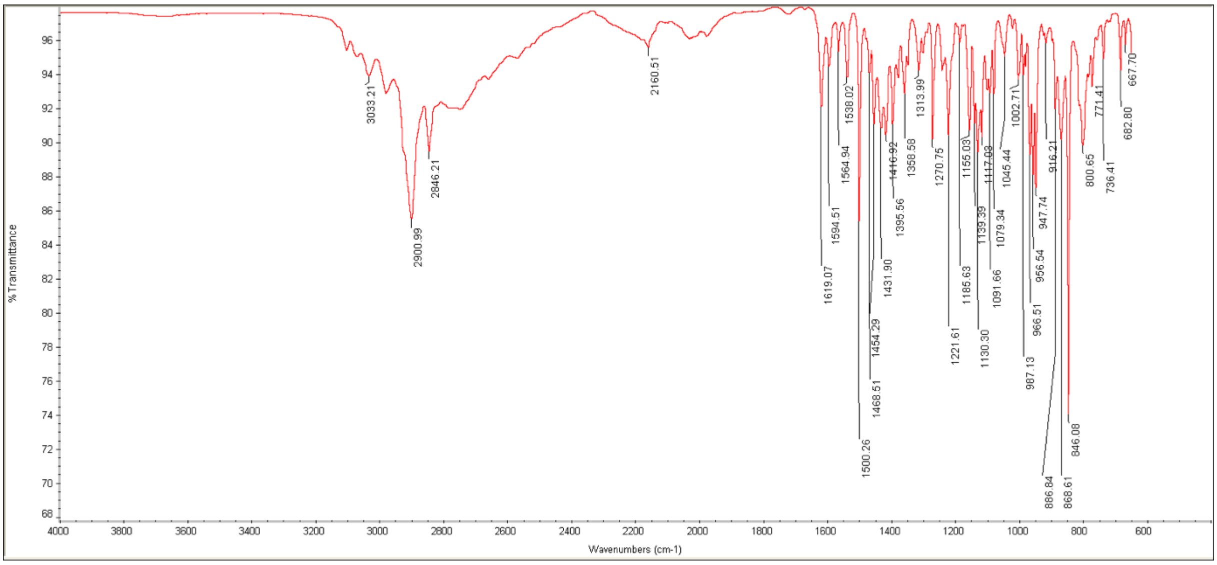

**Figure S48.** Infrared spectra of **12a**.

#### 11.NMR spectra

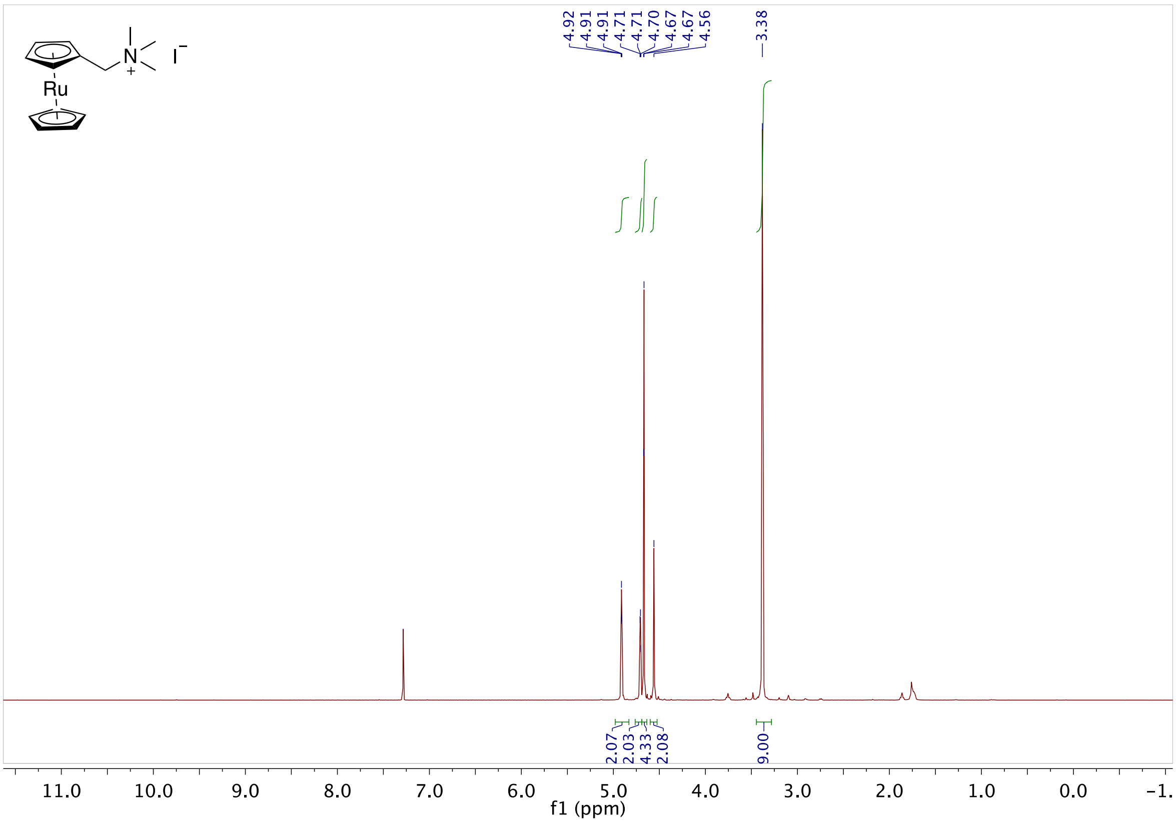

**Fig S48.** 1H NMR of **7** in $CDCl_3$

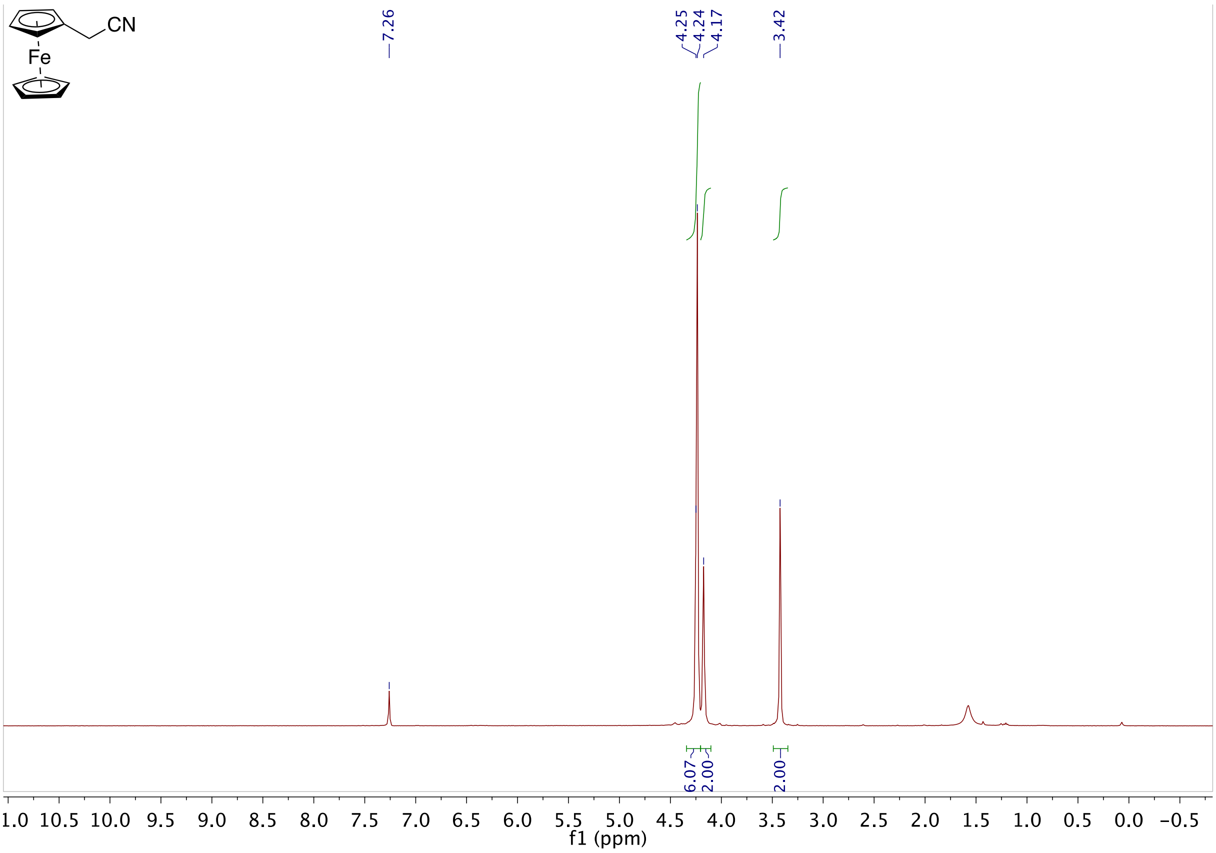

**Fig S49.** 1H NMR of **8** in $CDCl_3$

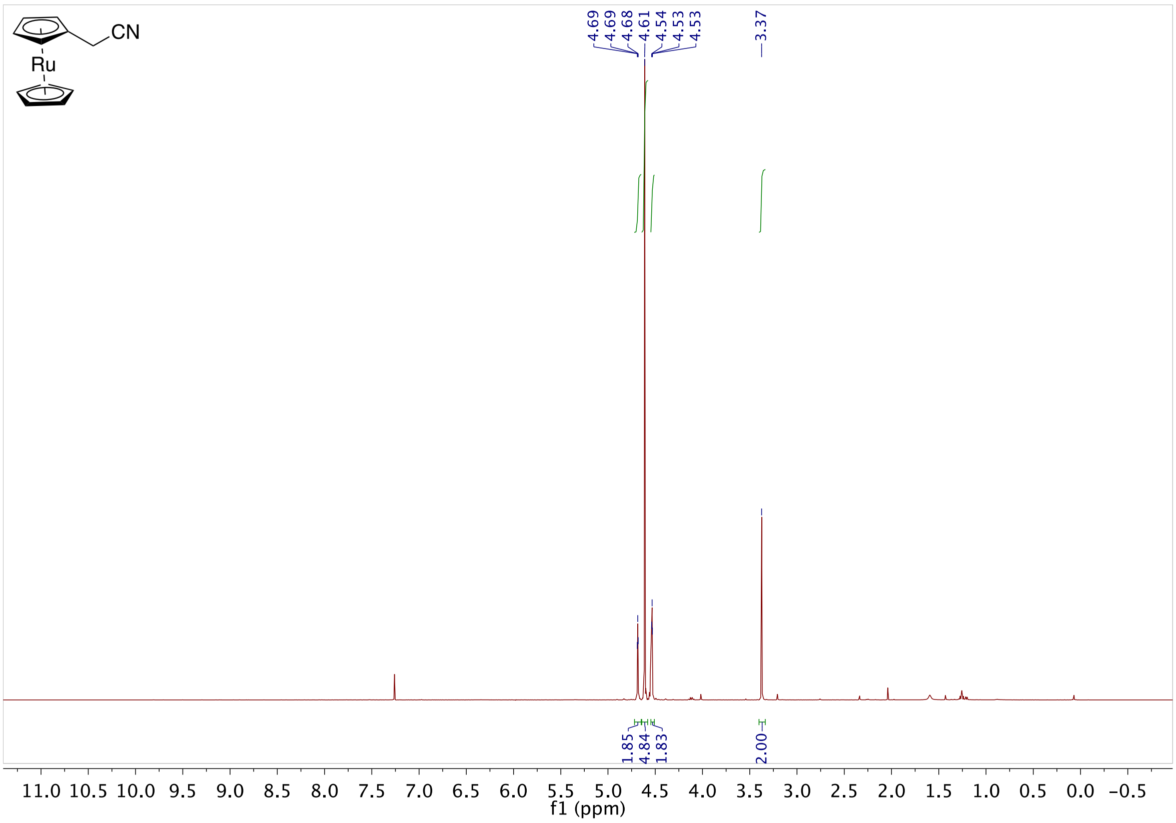

**Fig S50.** 1H NMR of **9** in $CDCl_3$

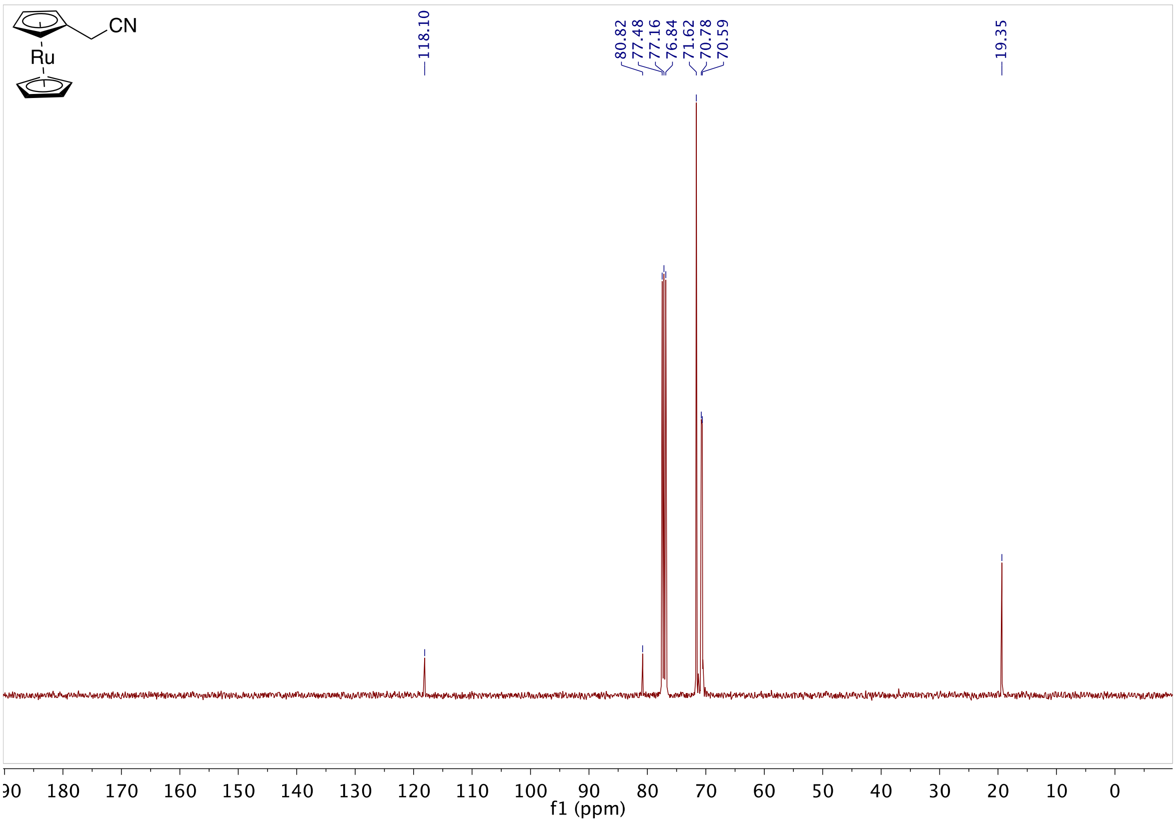

**Fig S51.** 13C NMR of **9** in $CDCl_3$

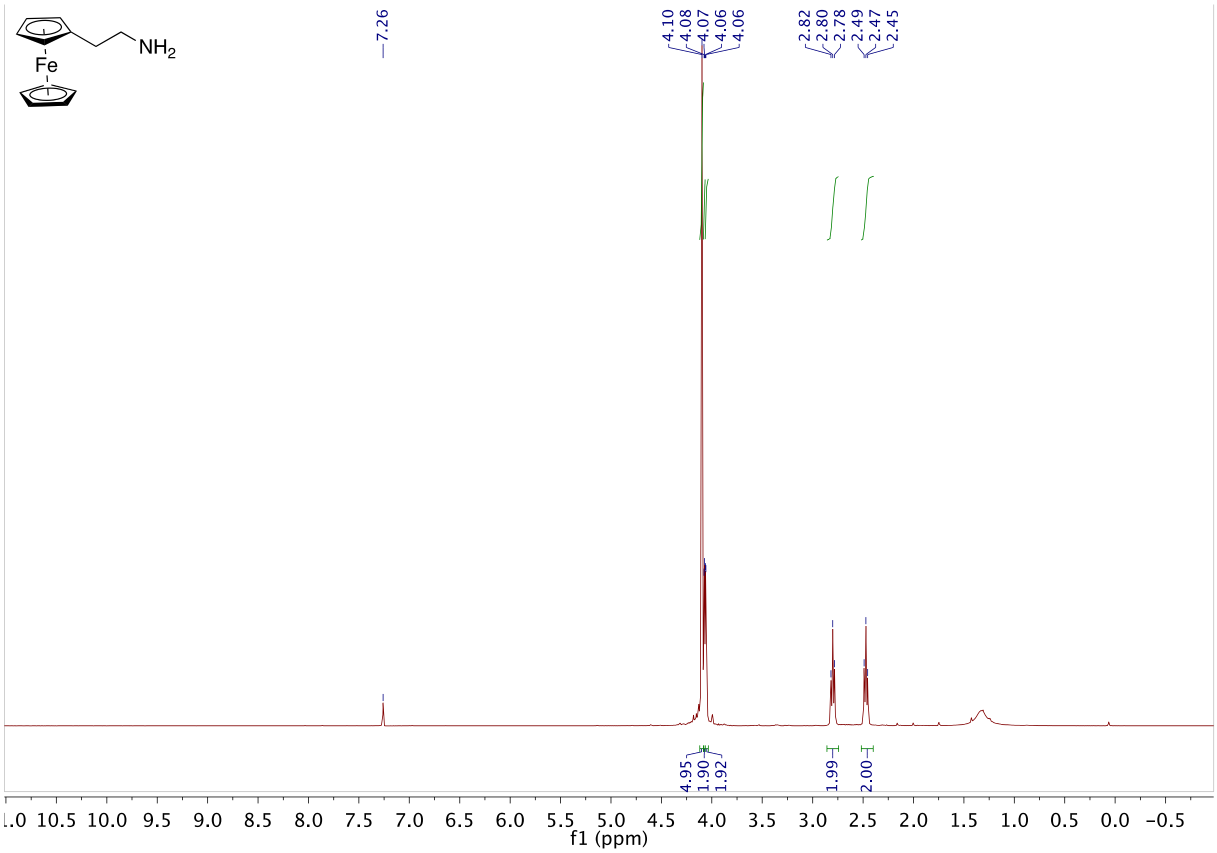

**Fig S52.** 1H NMR of **10** in $CDCl_3$

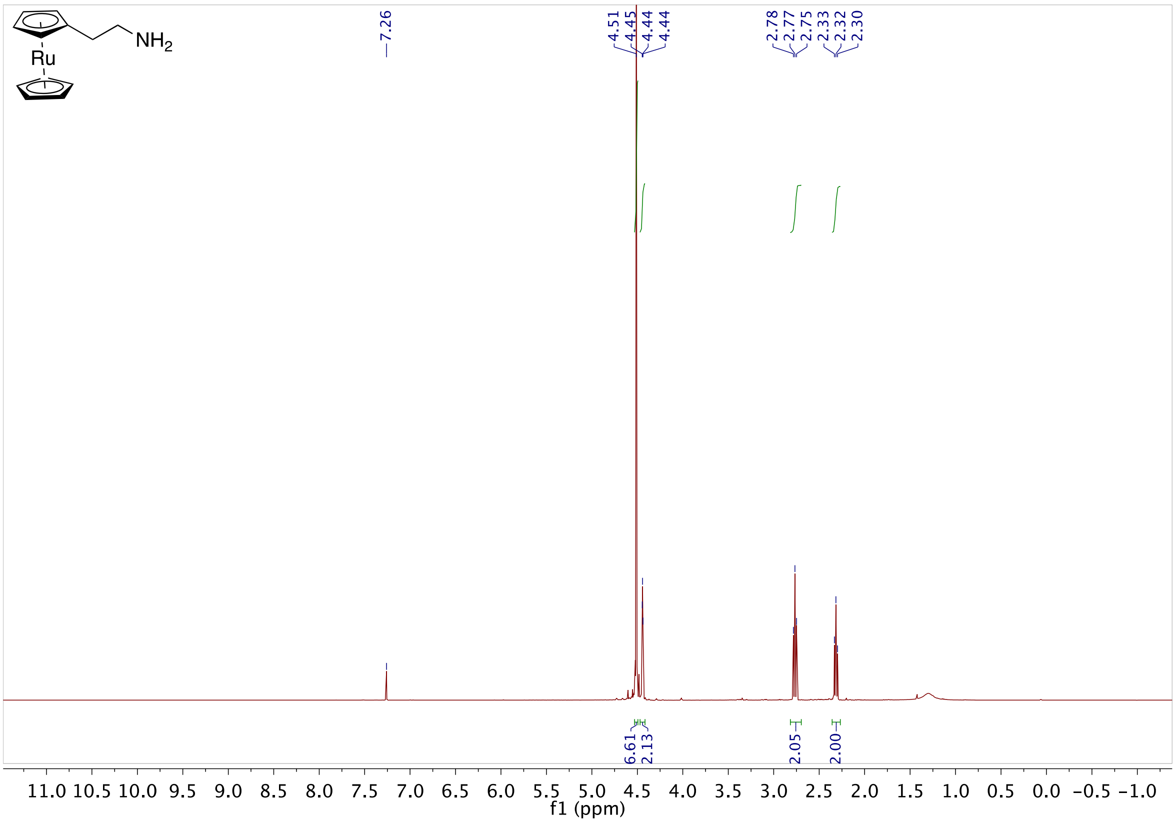

**Fig S53.** 1H NMR of **11** in CDCl₃

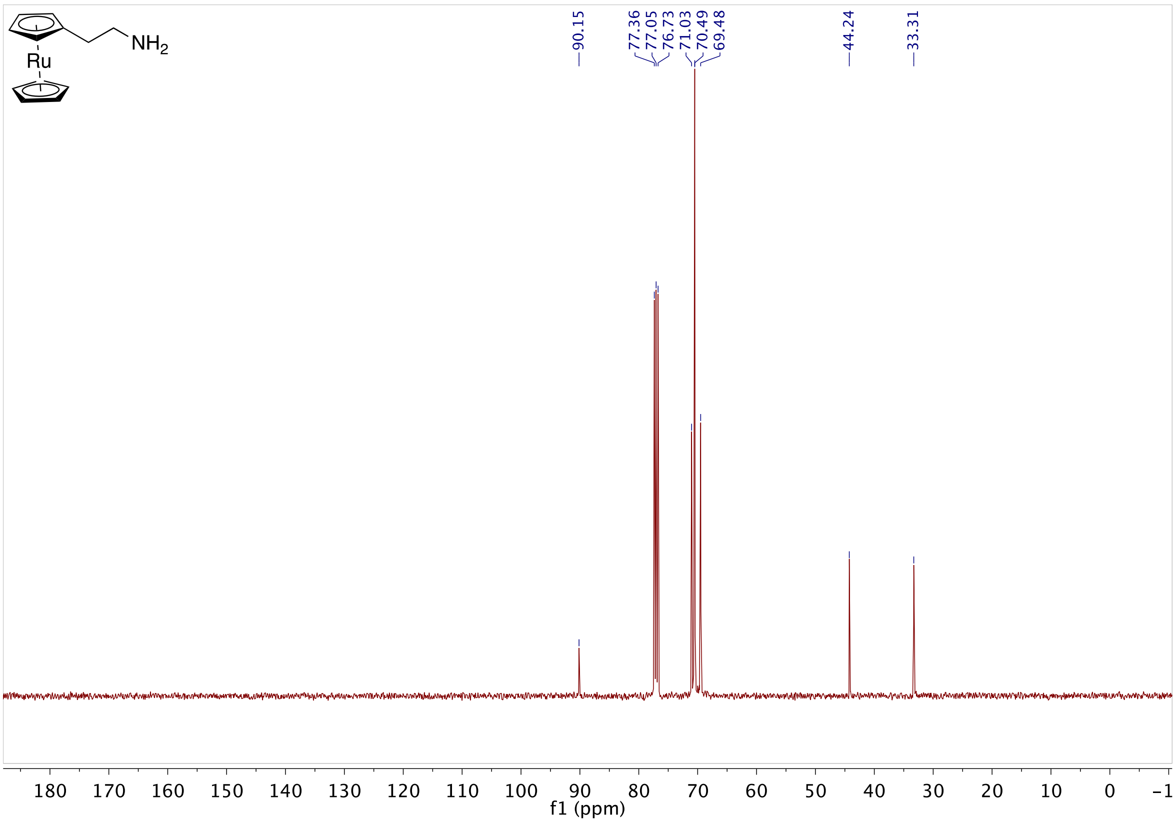

**Fig S54.** 13C NMR of **11** in CDCl₃

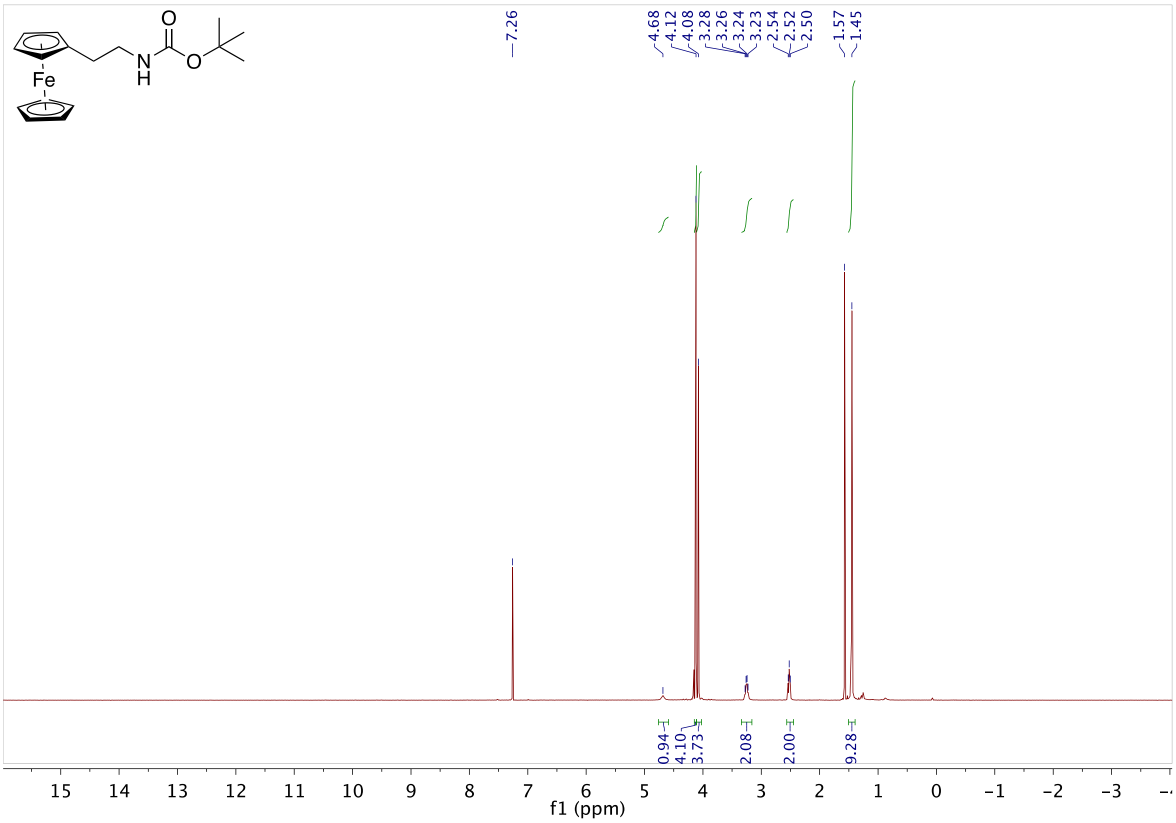

**Fig S55.** 1H NMR of **12** in $CDCl_3$

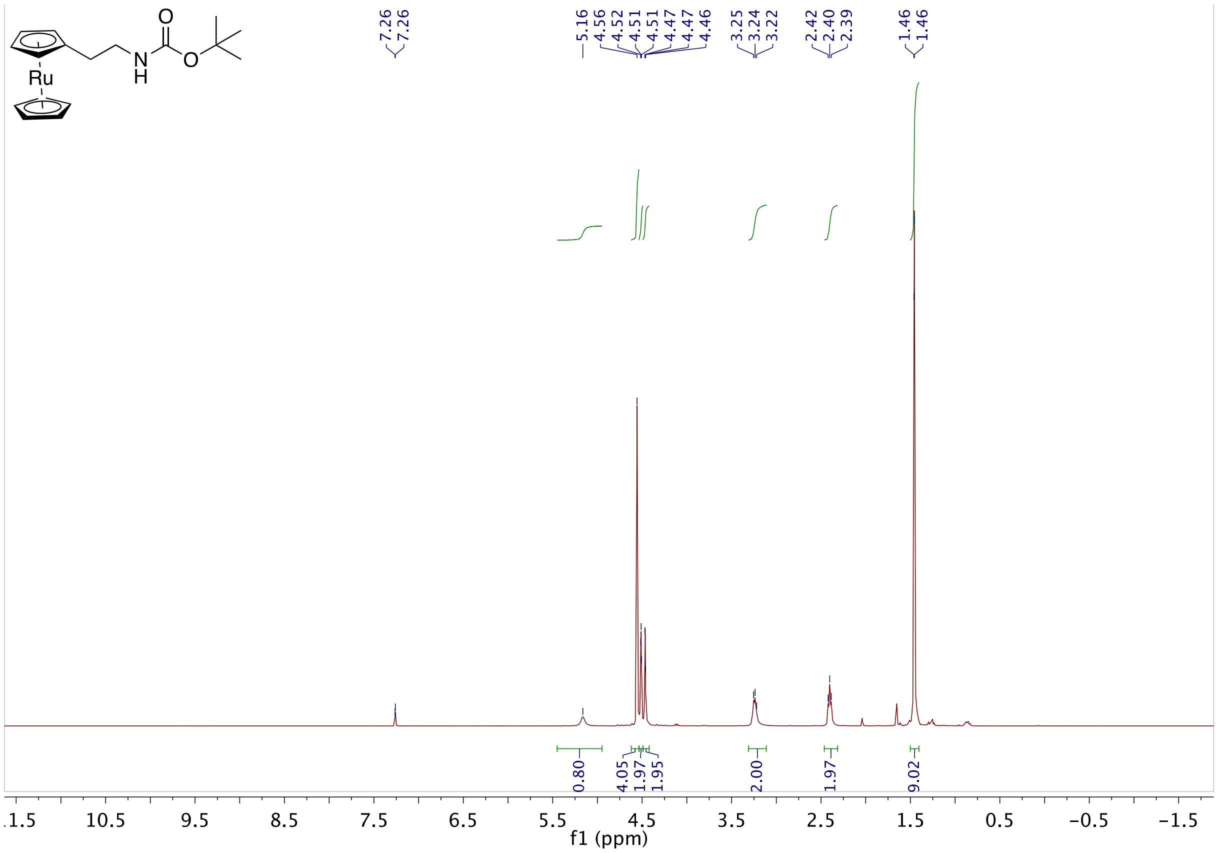

**Fig S56.** 1H NMR of **13** in $CDCl_3$

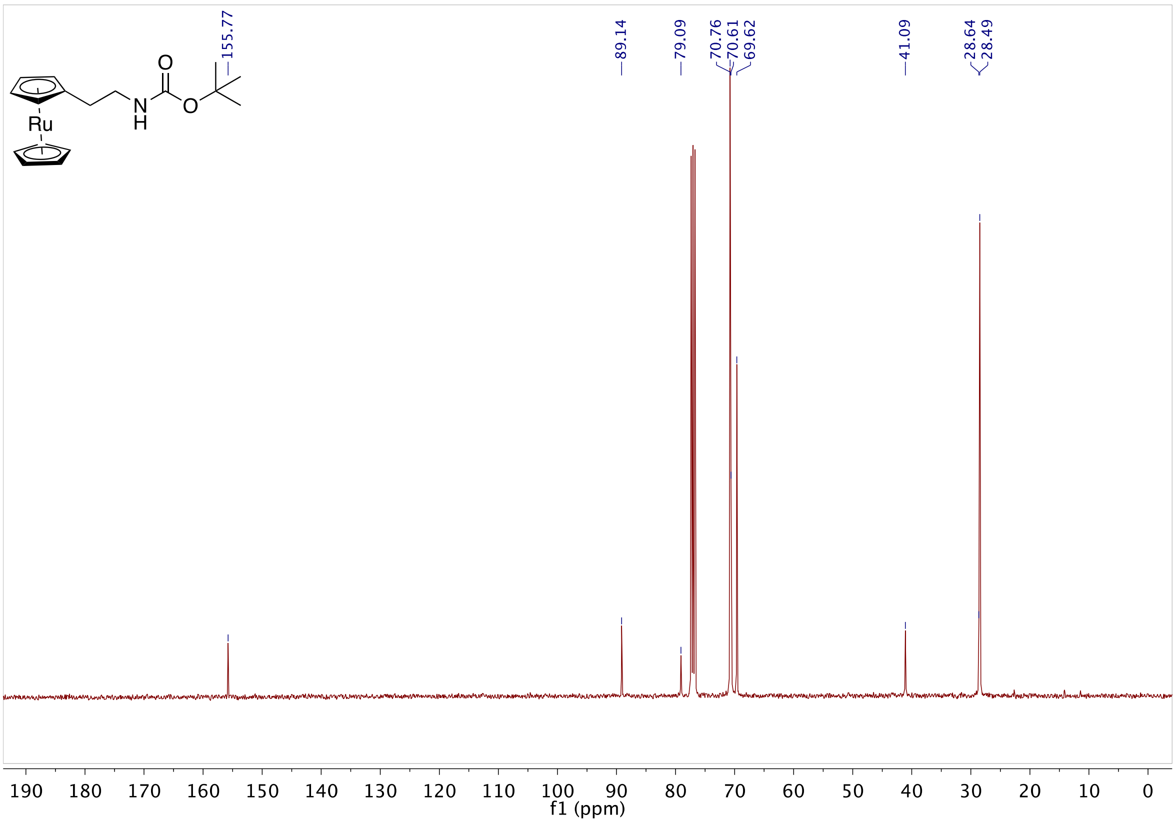

**Fig S57.** 13C NMR of **13** in CDCl₃

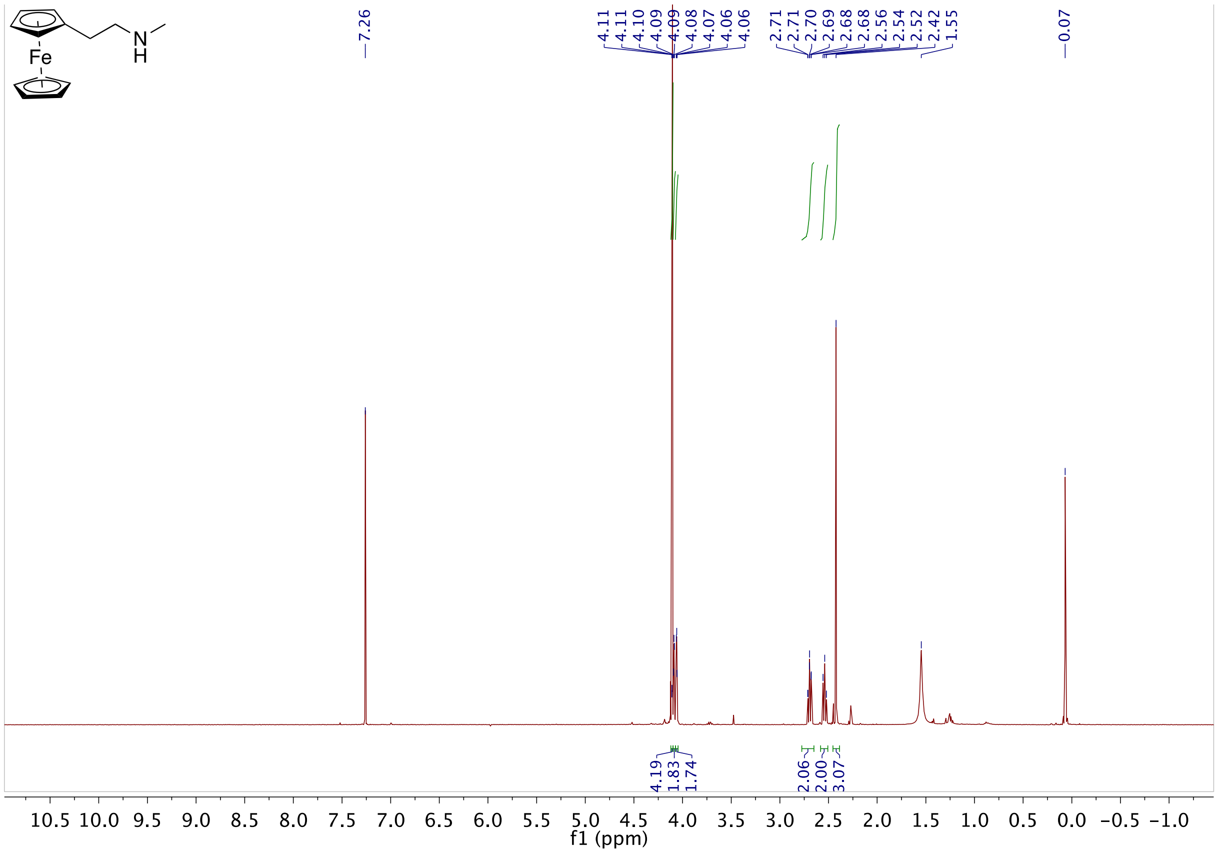

**Fig S58.** 1H NMR of **14** in CDCl₃

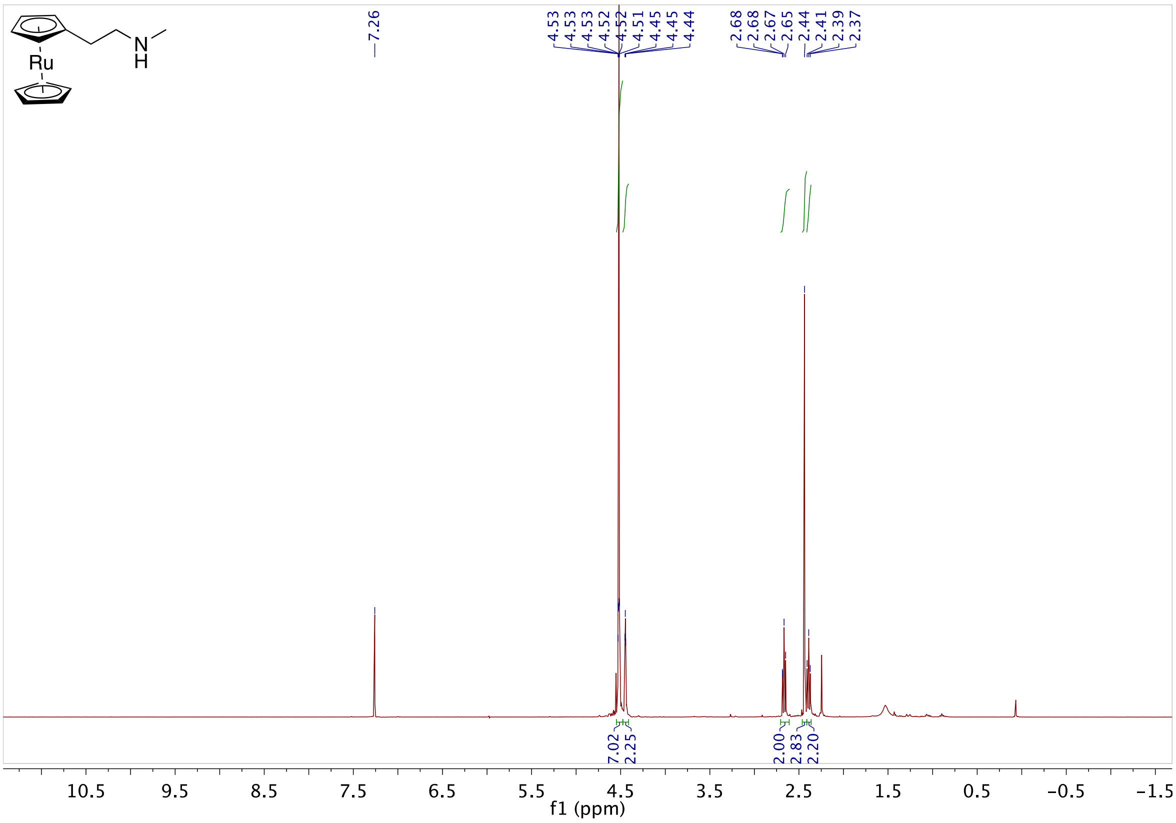

**Fig S59.** 1H NMR of **15** in $CDCl_3$

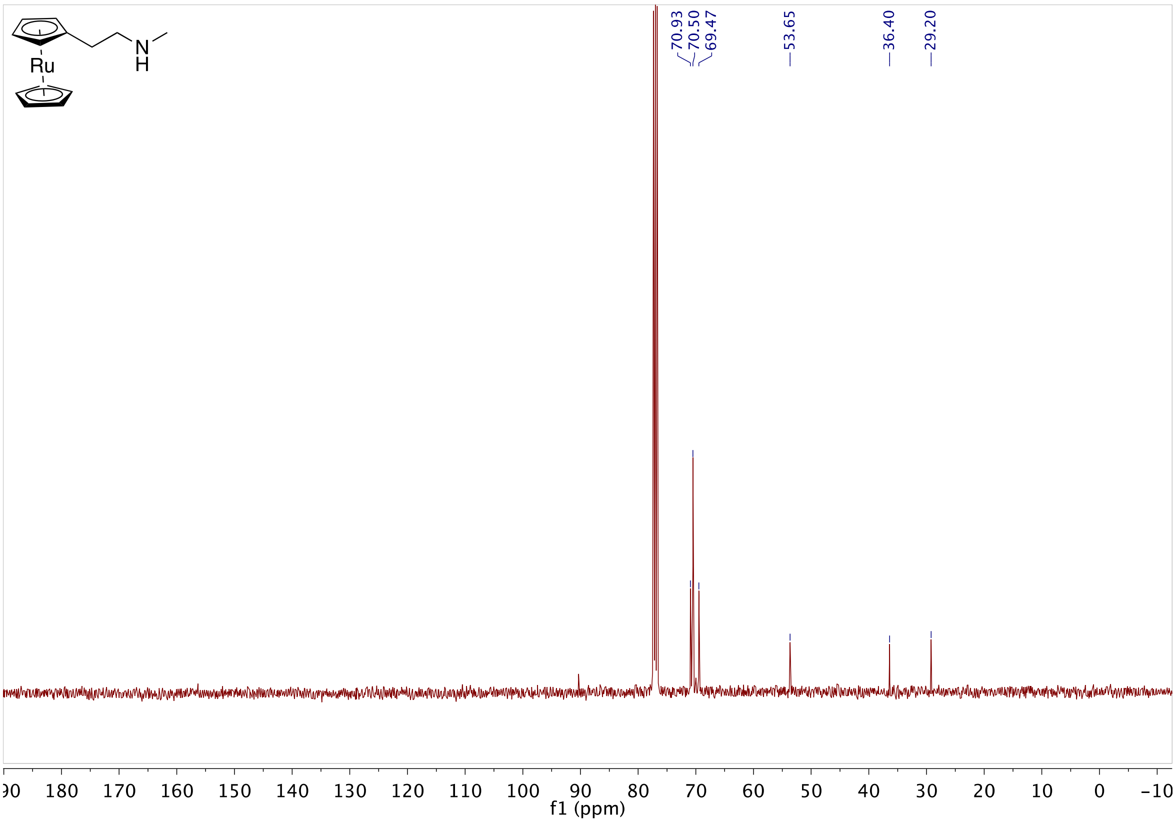

**Fig S60.** 13C NMR of **15** in $CDCl_3$

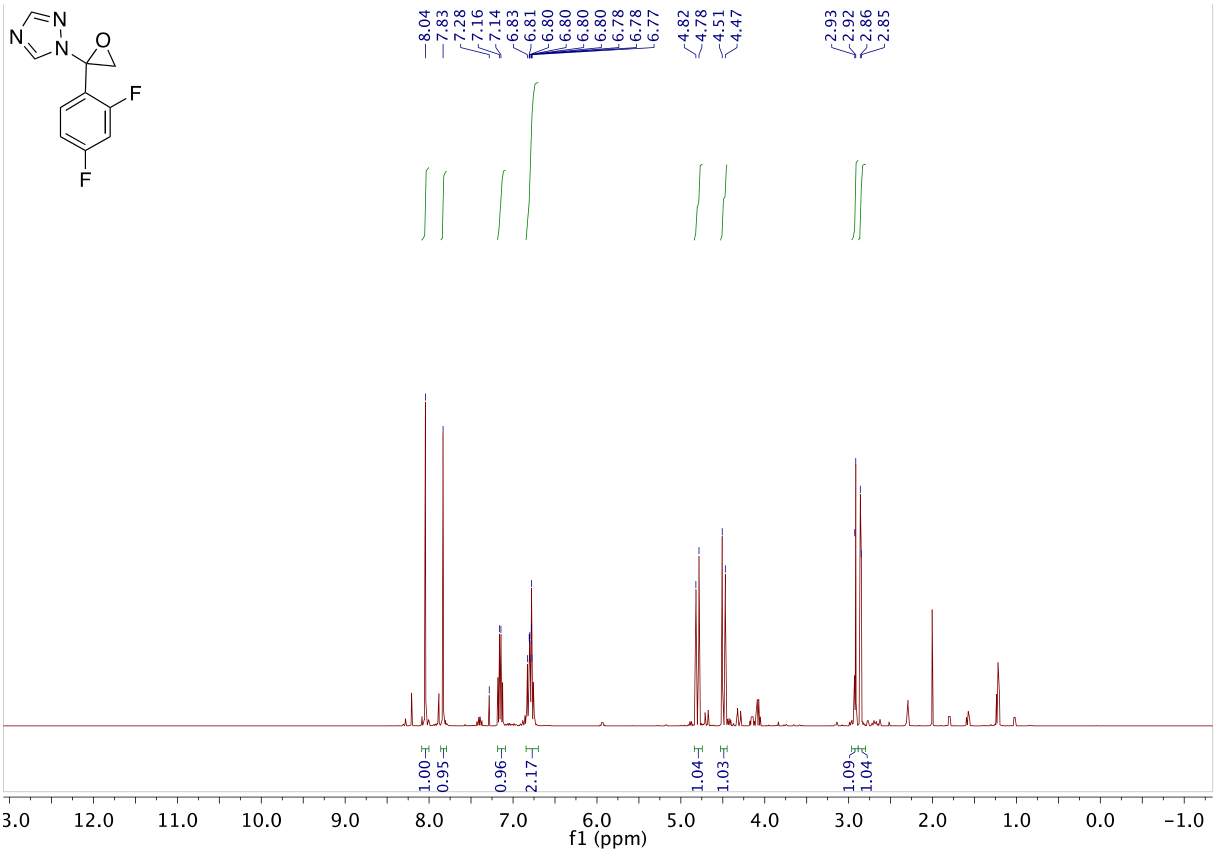

**Fig S61.** 1H NMR of **17** in $CDCl_3$

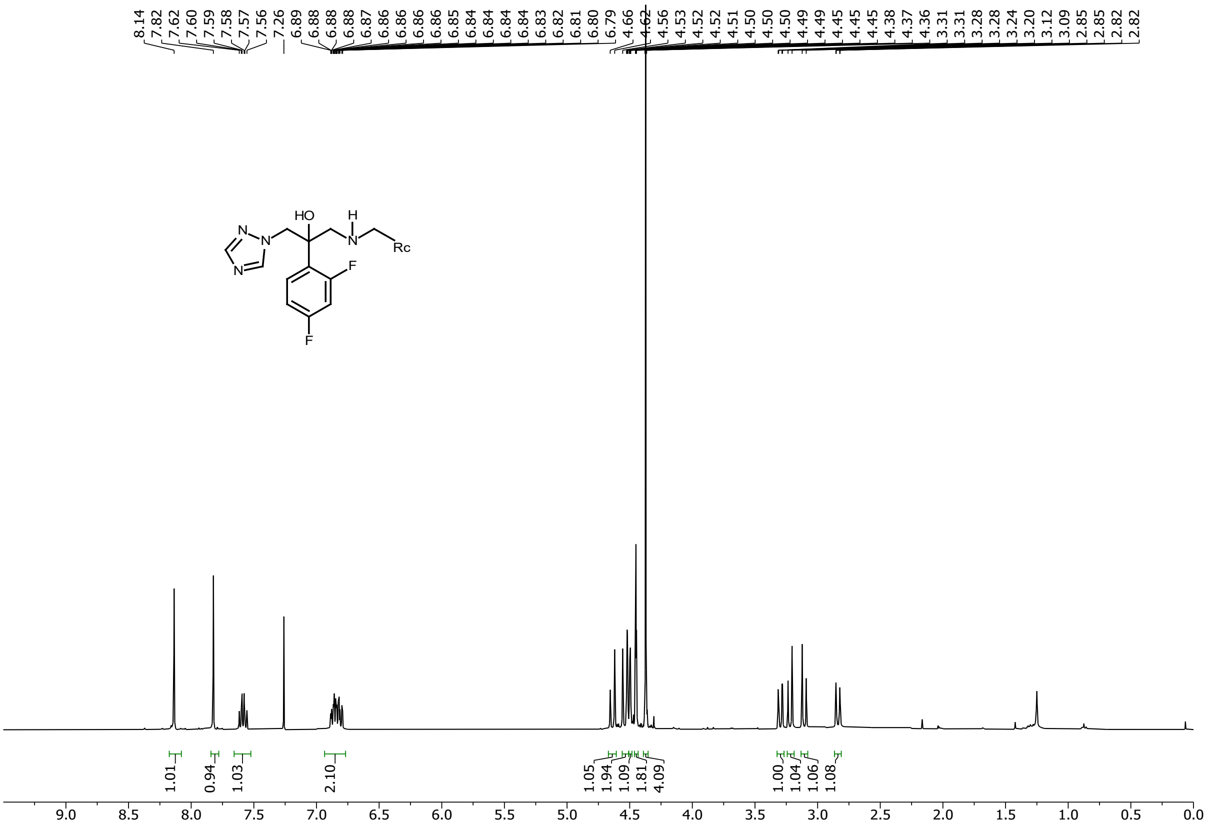

**Fig S62.** 1H NMR of **20** in $CDCl_3$

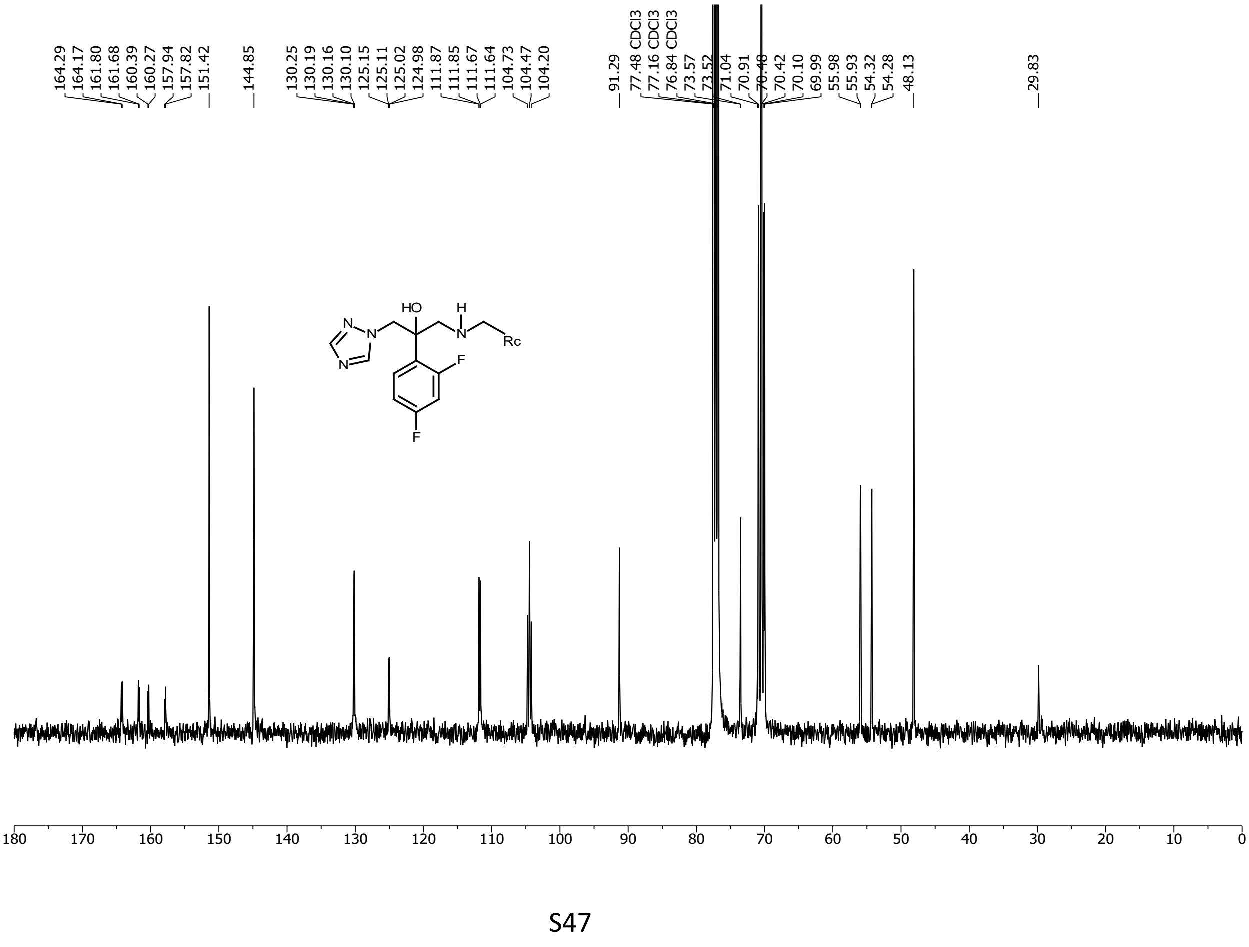

**Fig S63.** 13C NMR of **20** in $CDCl_3$

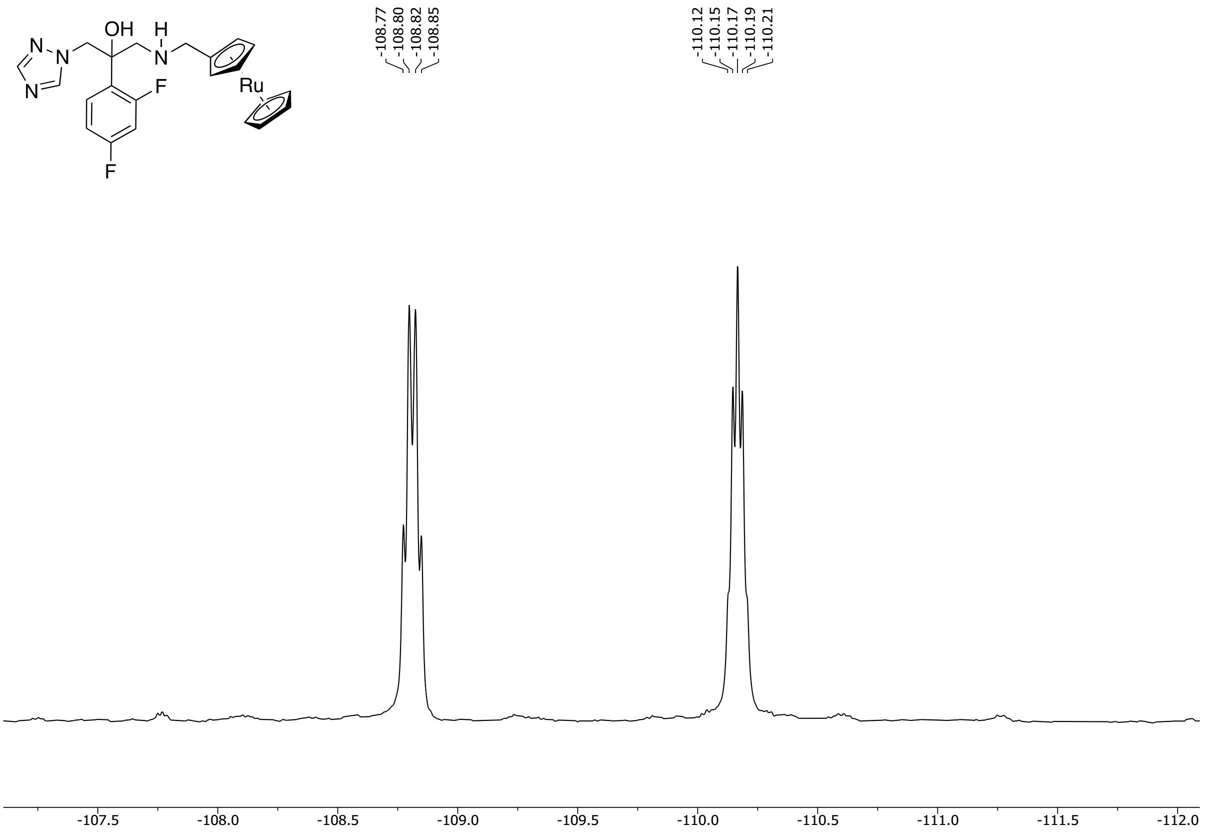

**Fig S64.** 19F NMR of **20** in $CDCl_3$

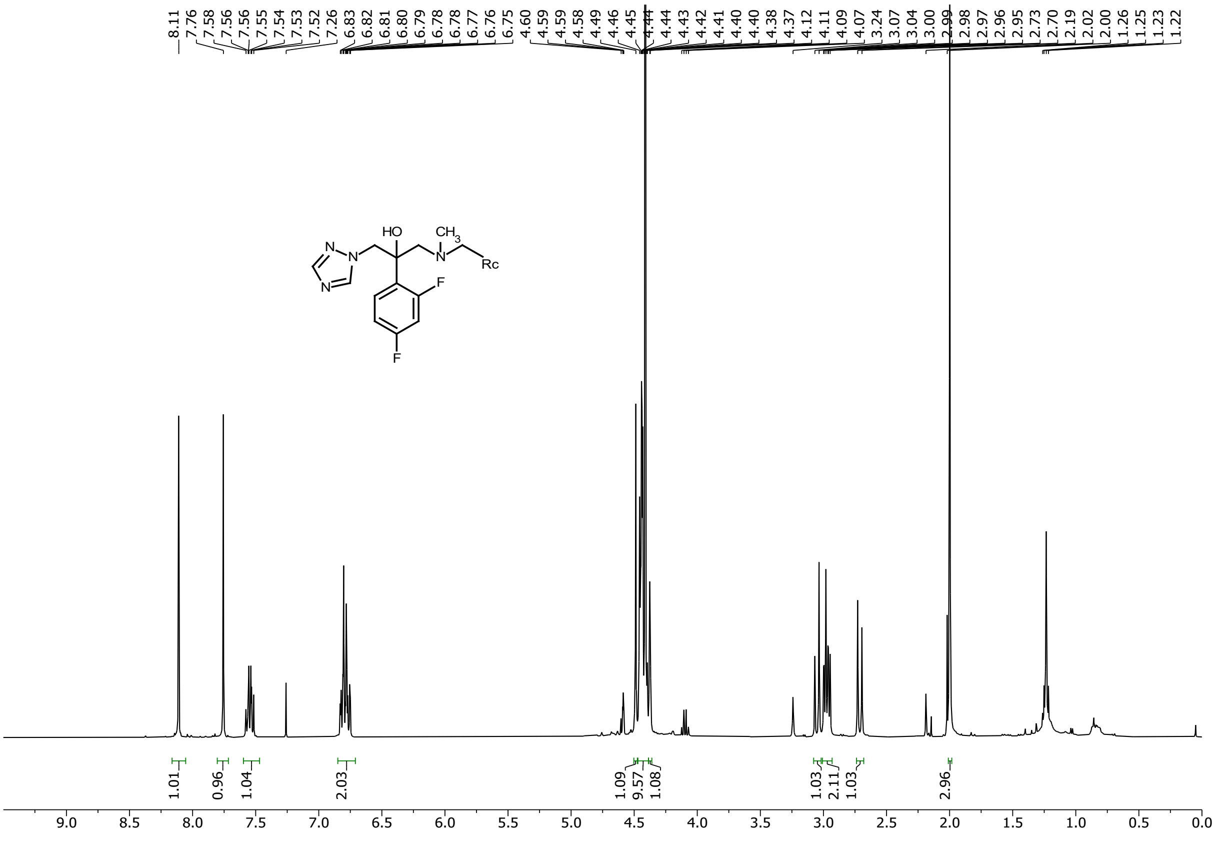

**Fig S65.** 1H NMR of **21** in $CDCl_3$

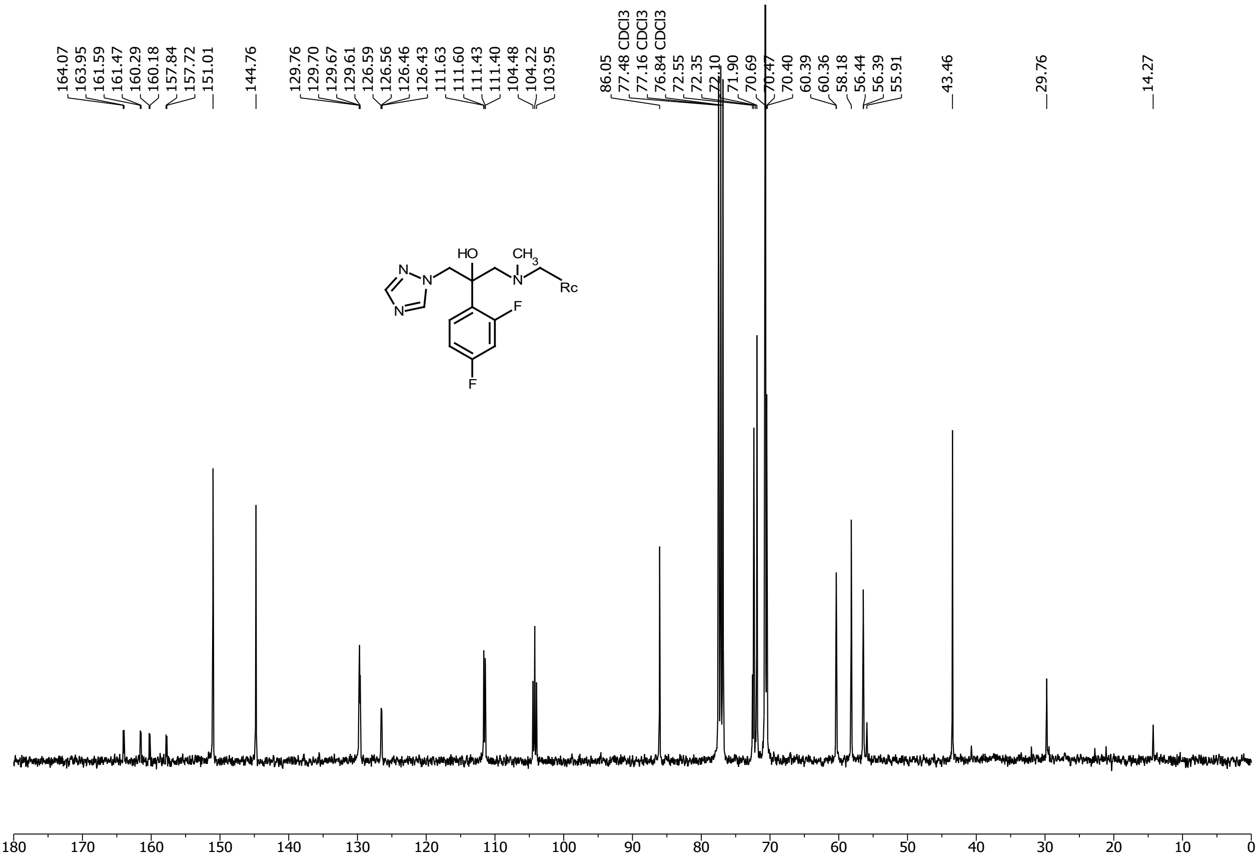

**Fig S66.** 13C NMR of **21** in $CDCl_3$

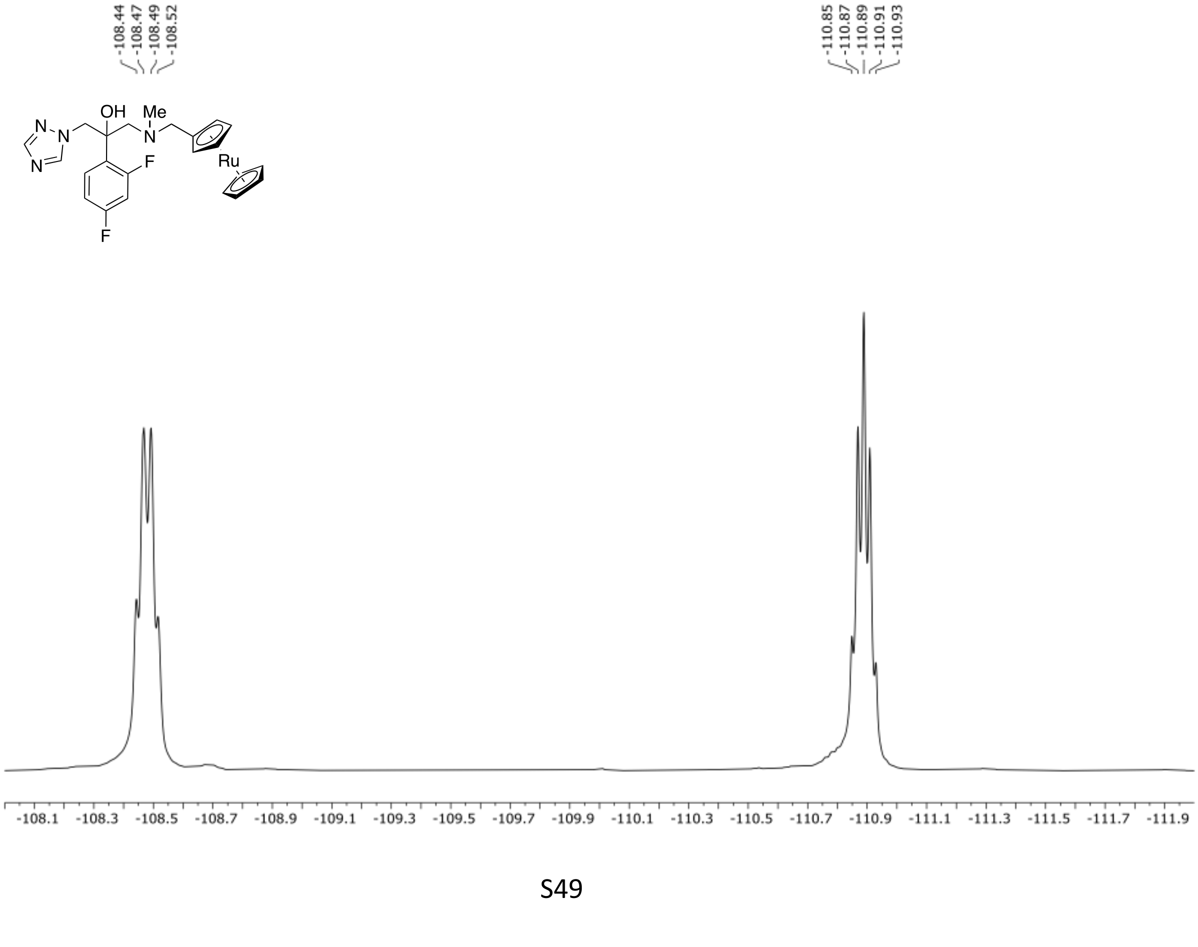

**Fig S67.** 19F NMR of **21** in $CDCl_3$

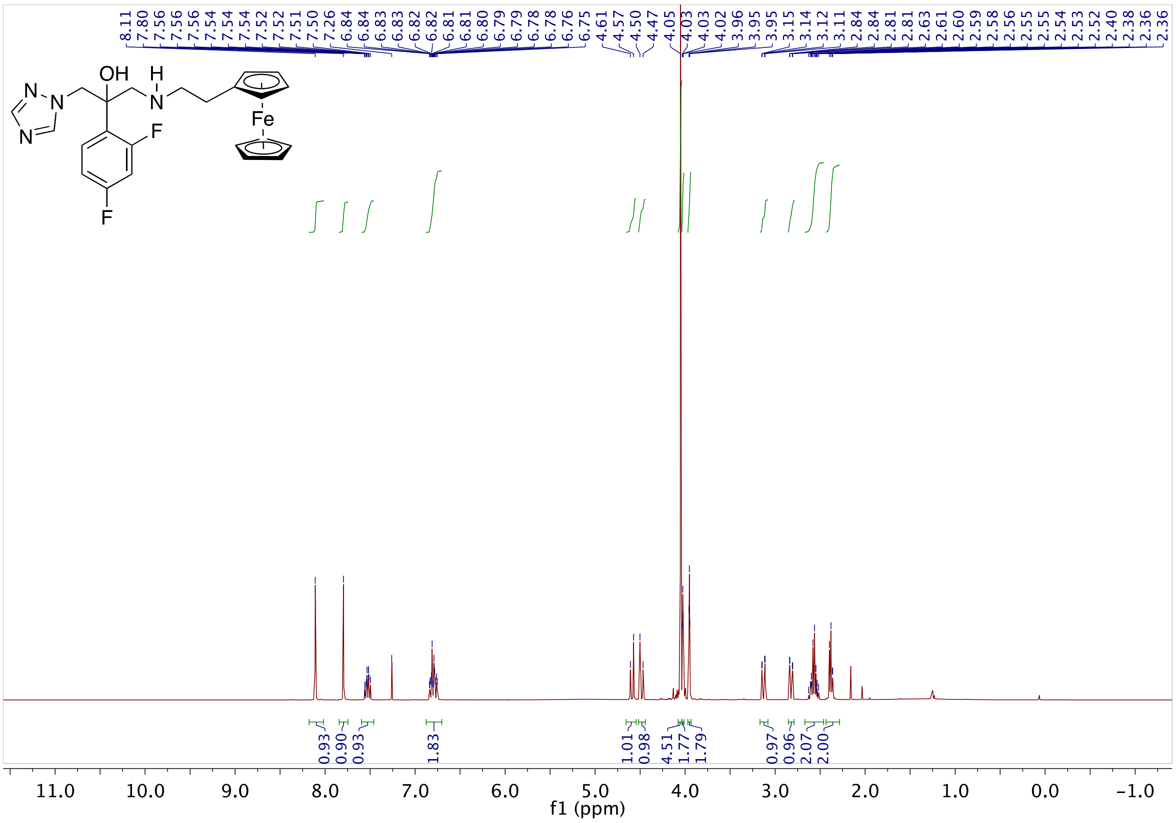

**Fig S68.** 1H NMR of **22** in $CDCl_3$

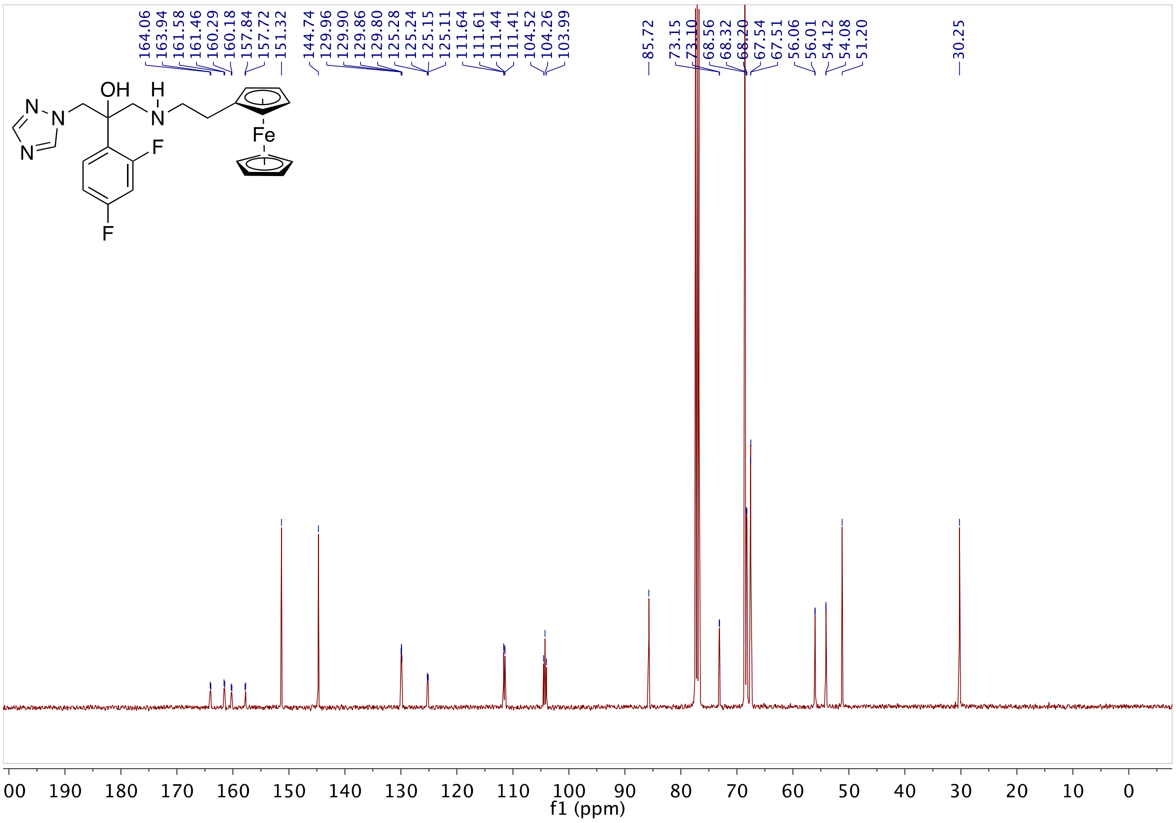

**Fig S69.** 13C NMR of **22** in $CDCl_3$

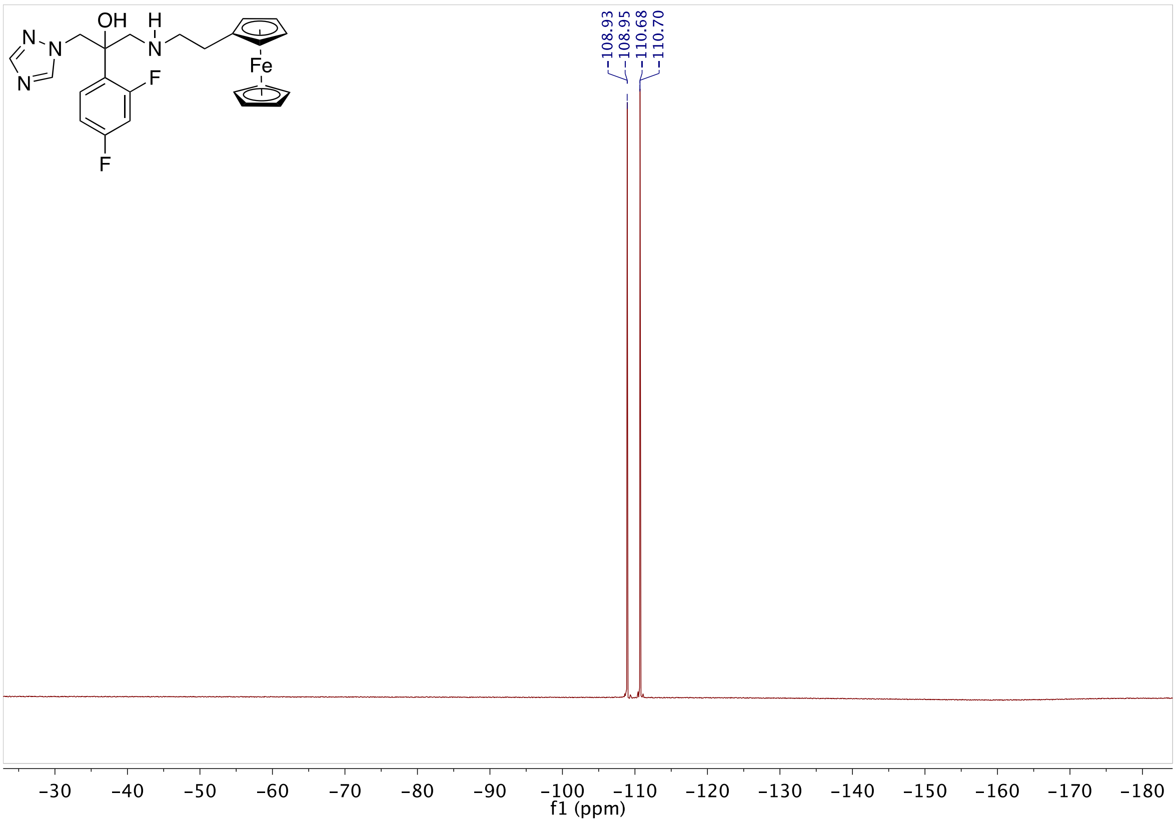

**Fig S70.** 19F NMR of **22** in $CDCl_3$

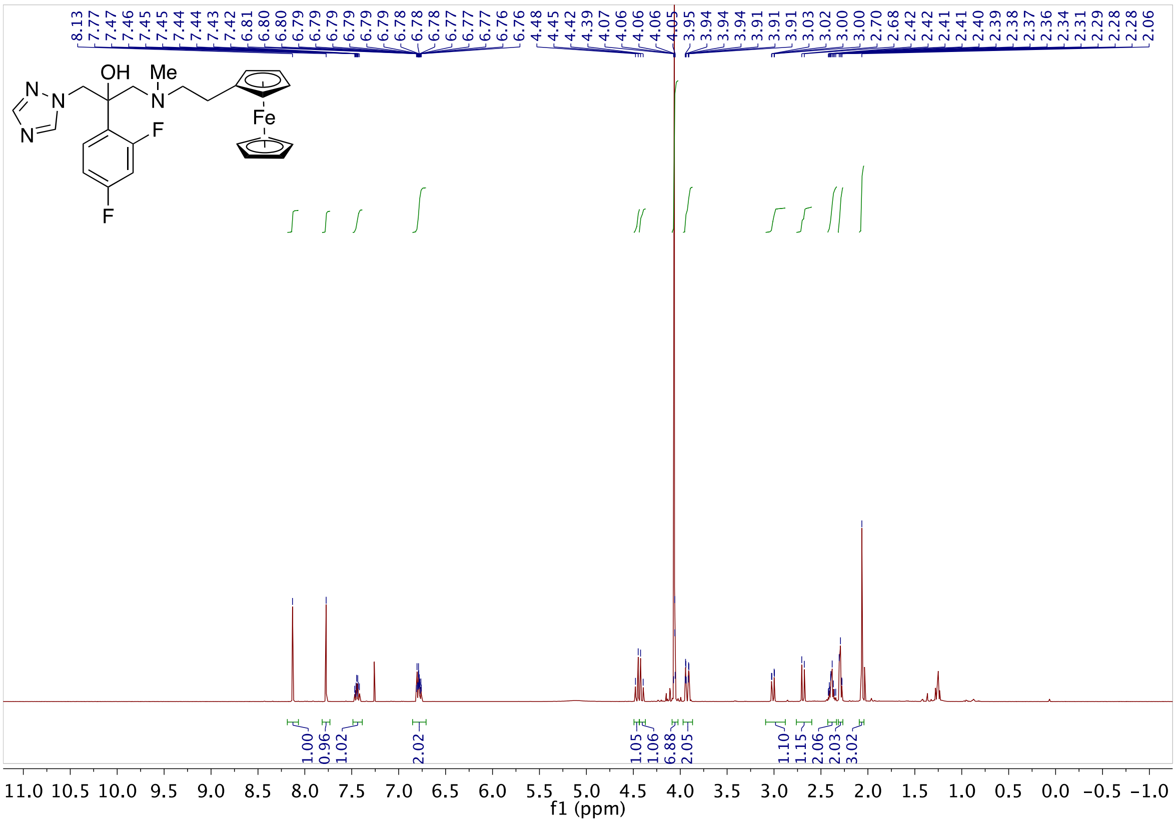

**Fig S71.** 1H NMR of **23** in CDCl₃

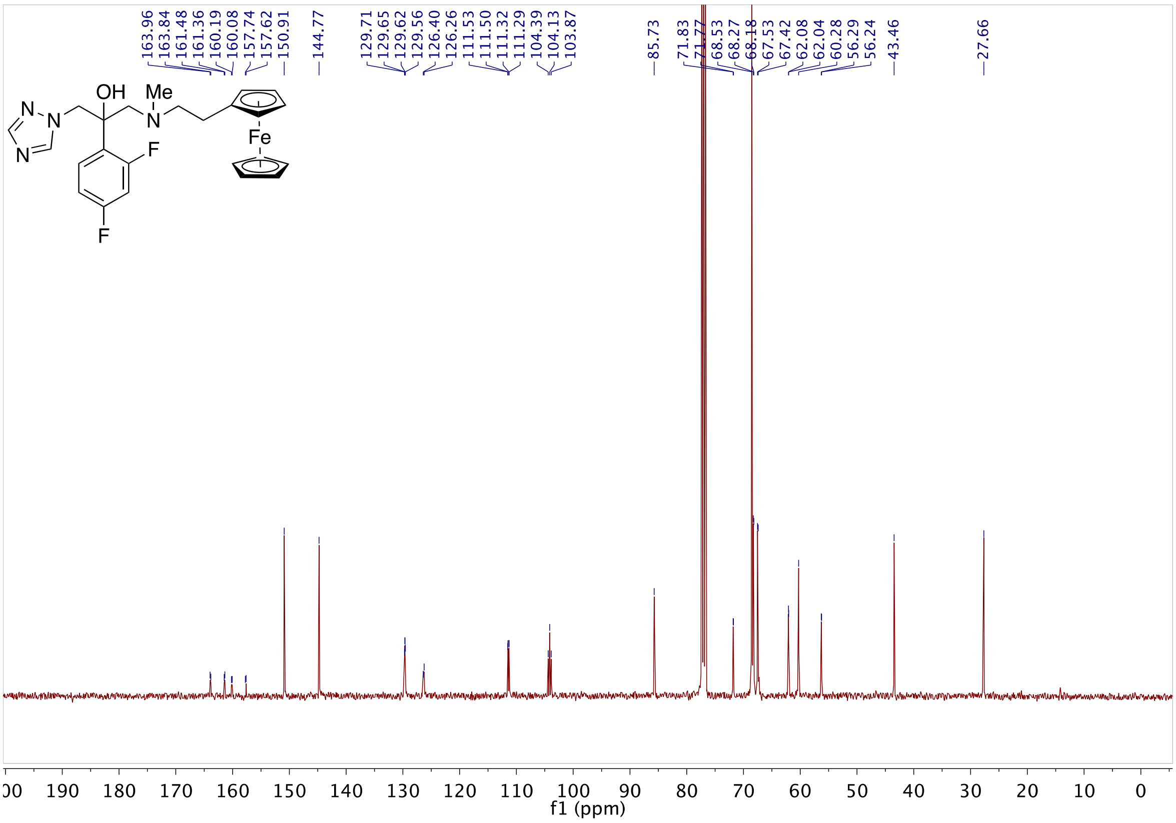

**Fig S72.** 13C NMR of **23** in CDCl₃

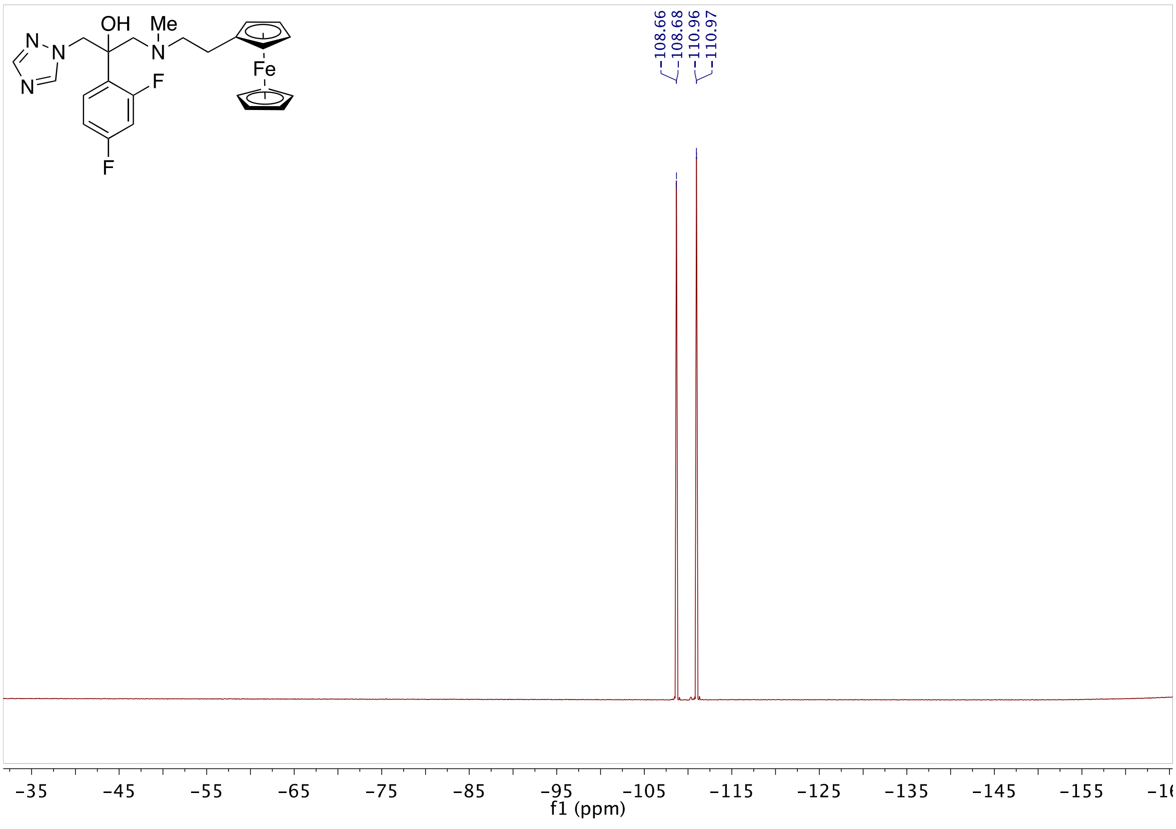

**Fig S73.** 19F NMR of **23** in $CDCl_3$

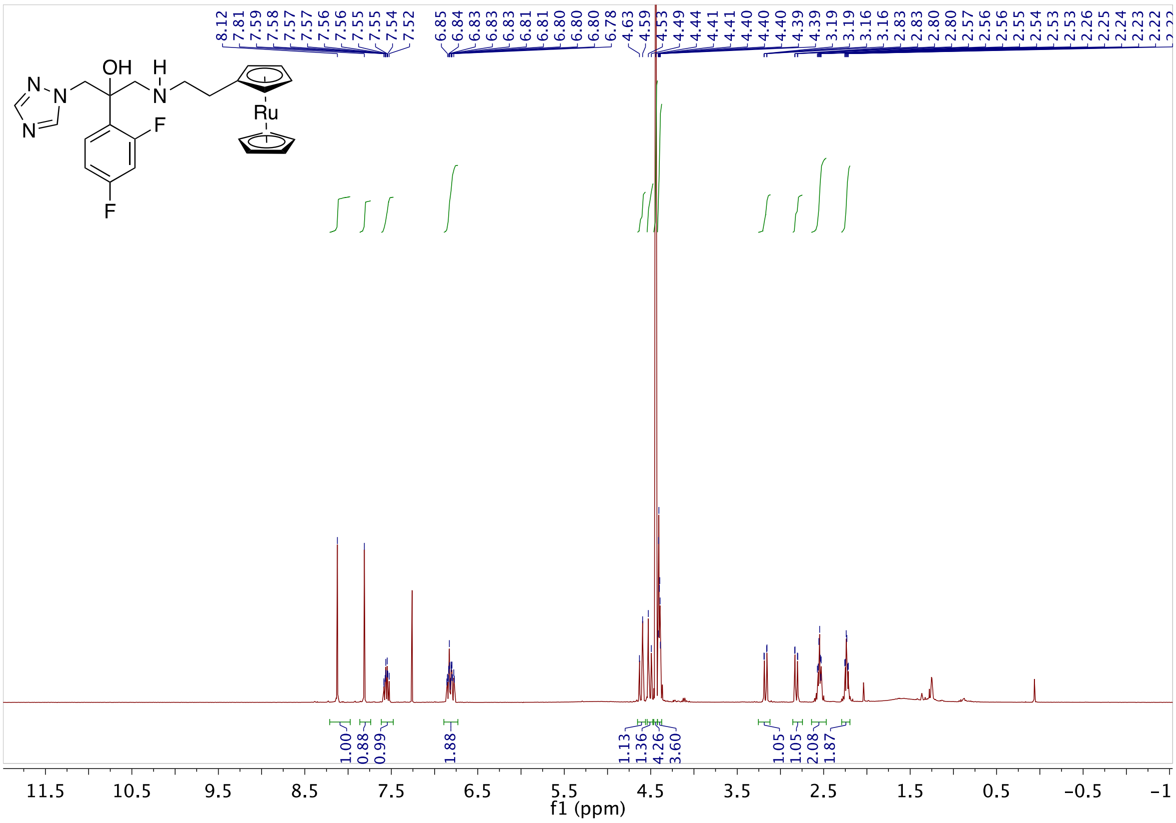

**Fig S74.** 1H NMR of **24** in $CDCl_3$

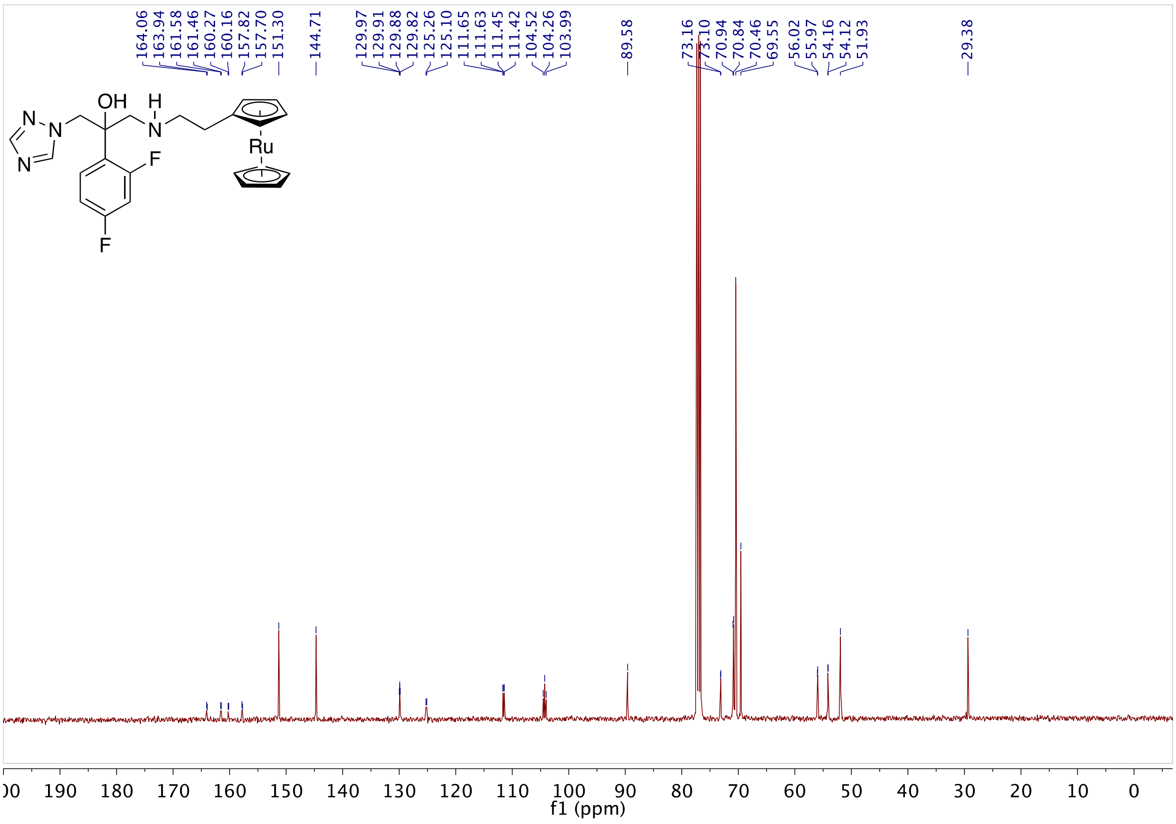

**Fig S75.** 13C NMR of **24** in $CDCl_3$

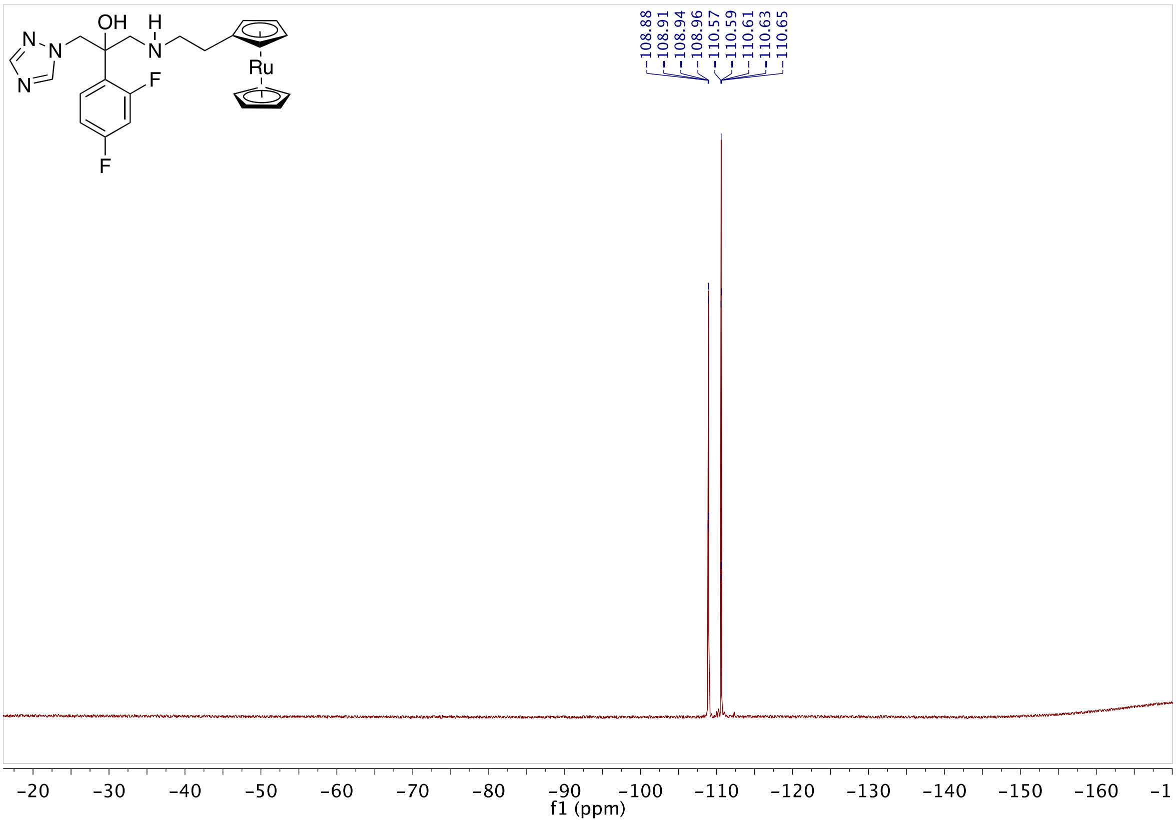

**Fig S76.** 19F NMR of **24** in $CDCl_3$

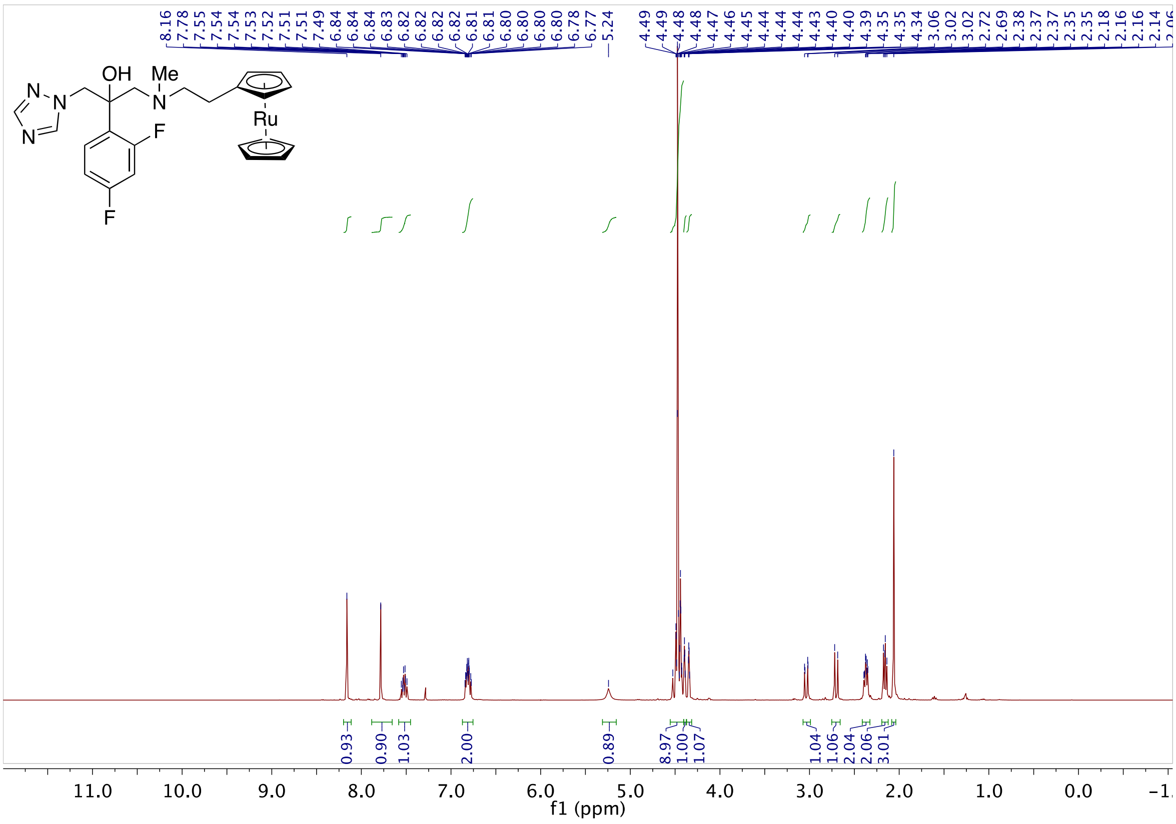

**Fig S77.** 1H NMR of **25** in $CDCl_3$

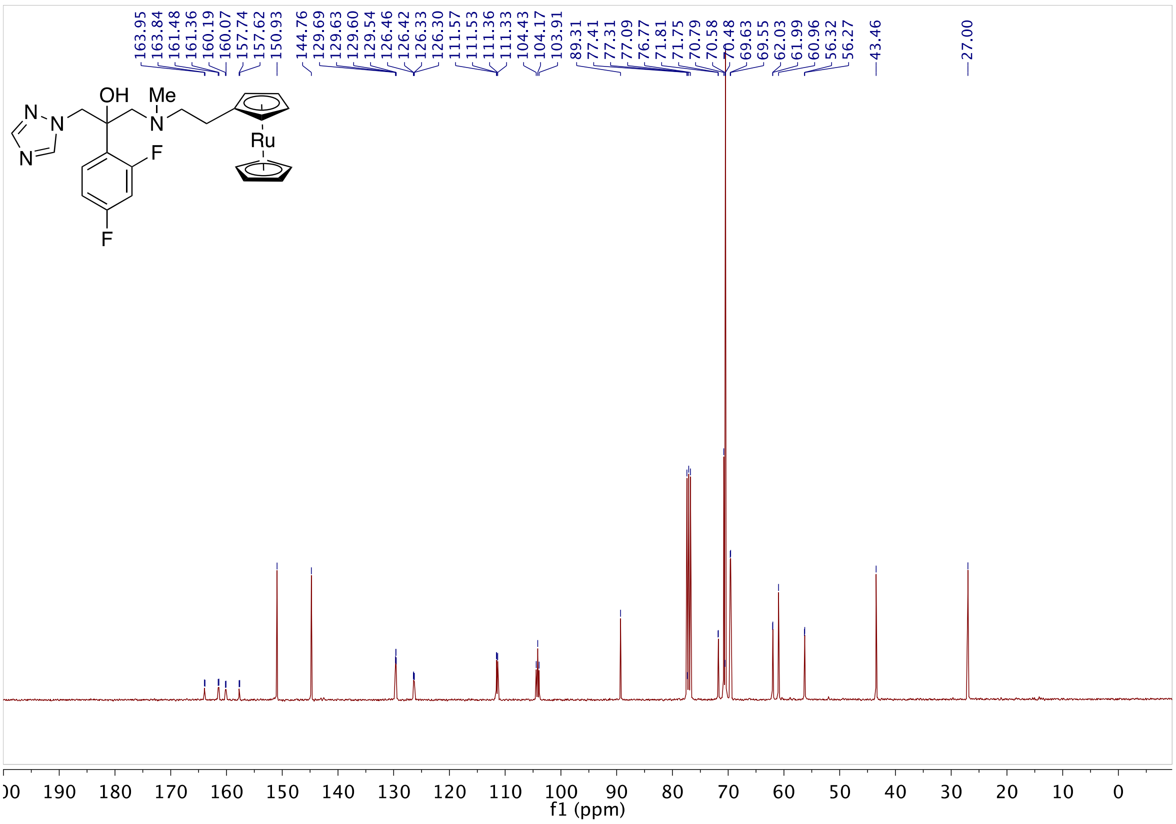

**Fig S78.** 13C NMR of **25** in $CDCl_3$

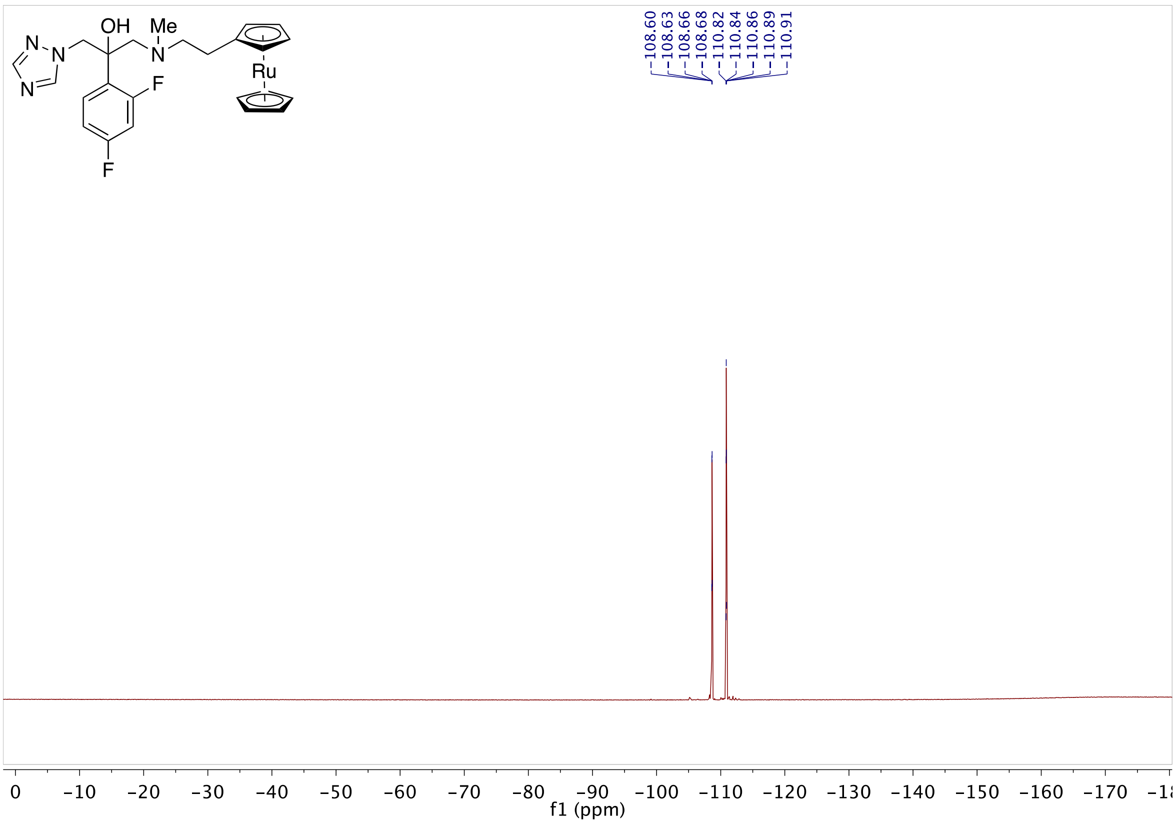

**Fig S79.** 19F NMR of **25** in $CDCl_3$

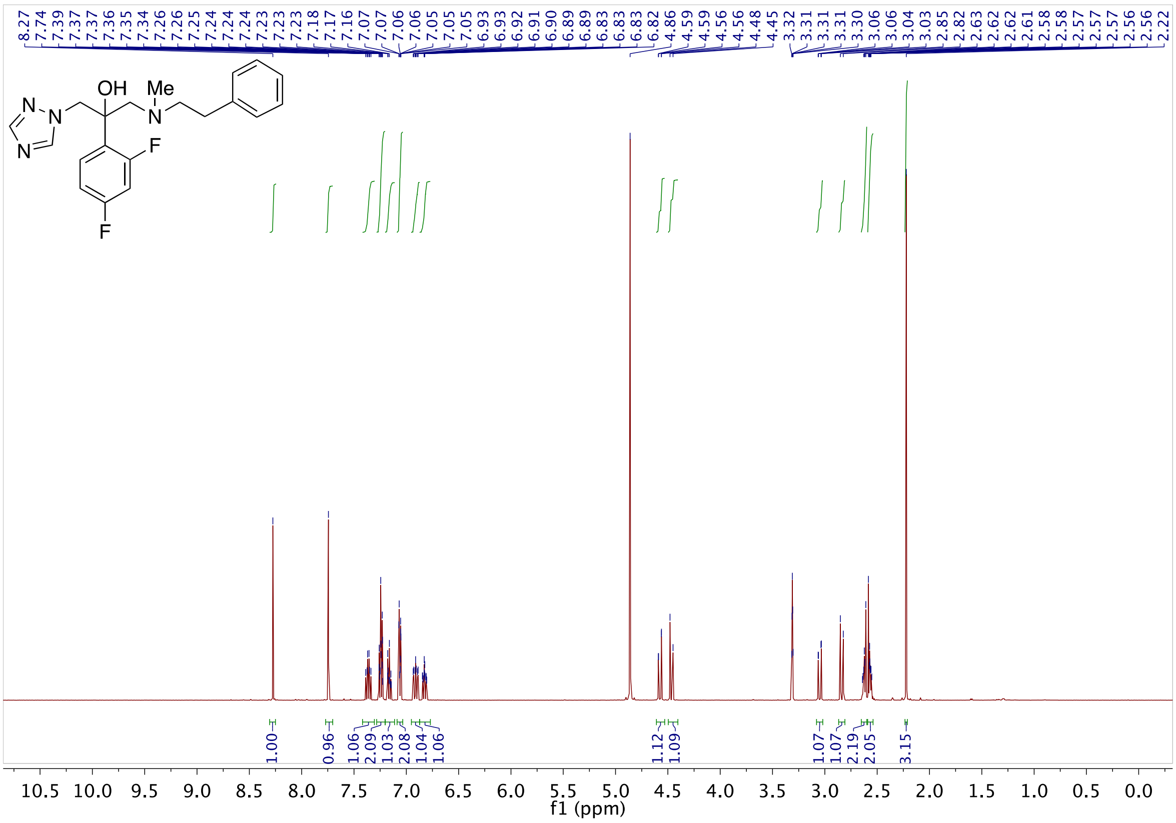

**Fig S80.** 1H NMR of **26** in CD3OD

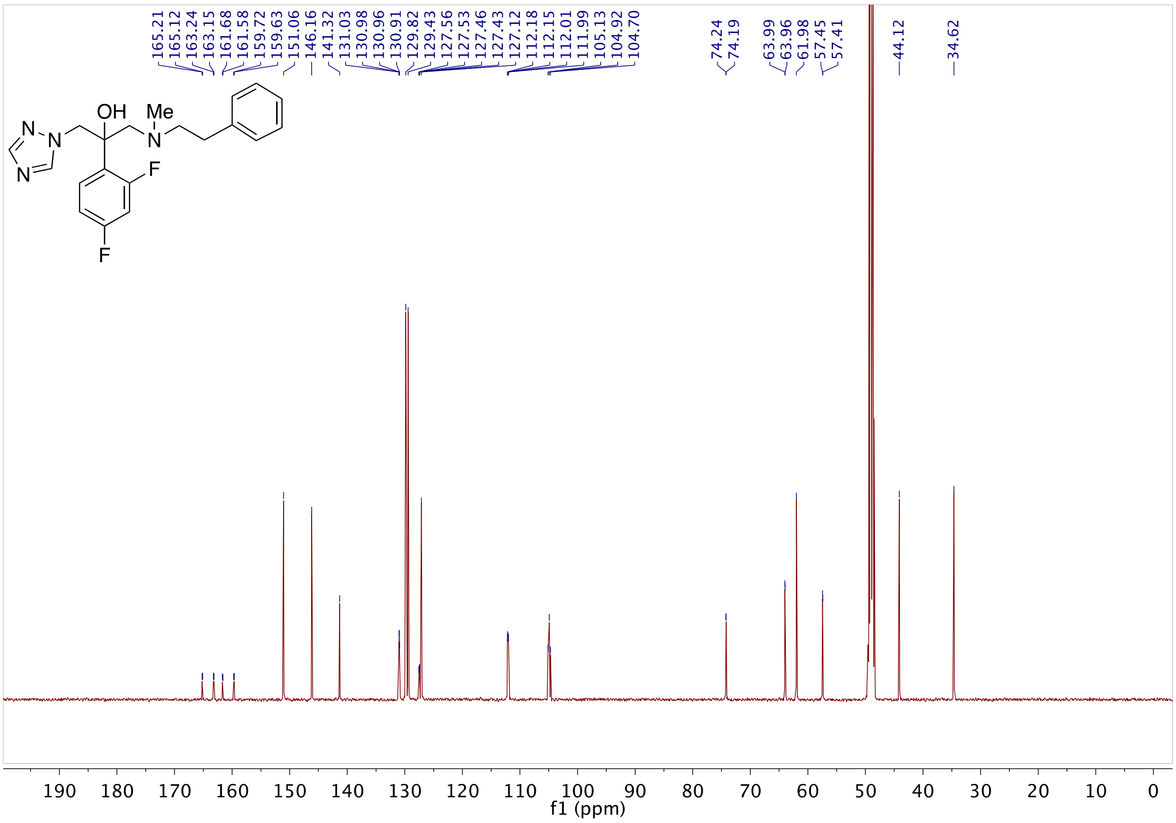

**Fig S81.** 13C NMR of **26** in CD3OD

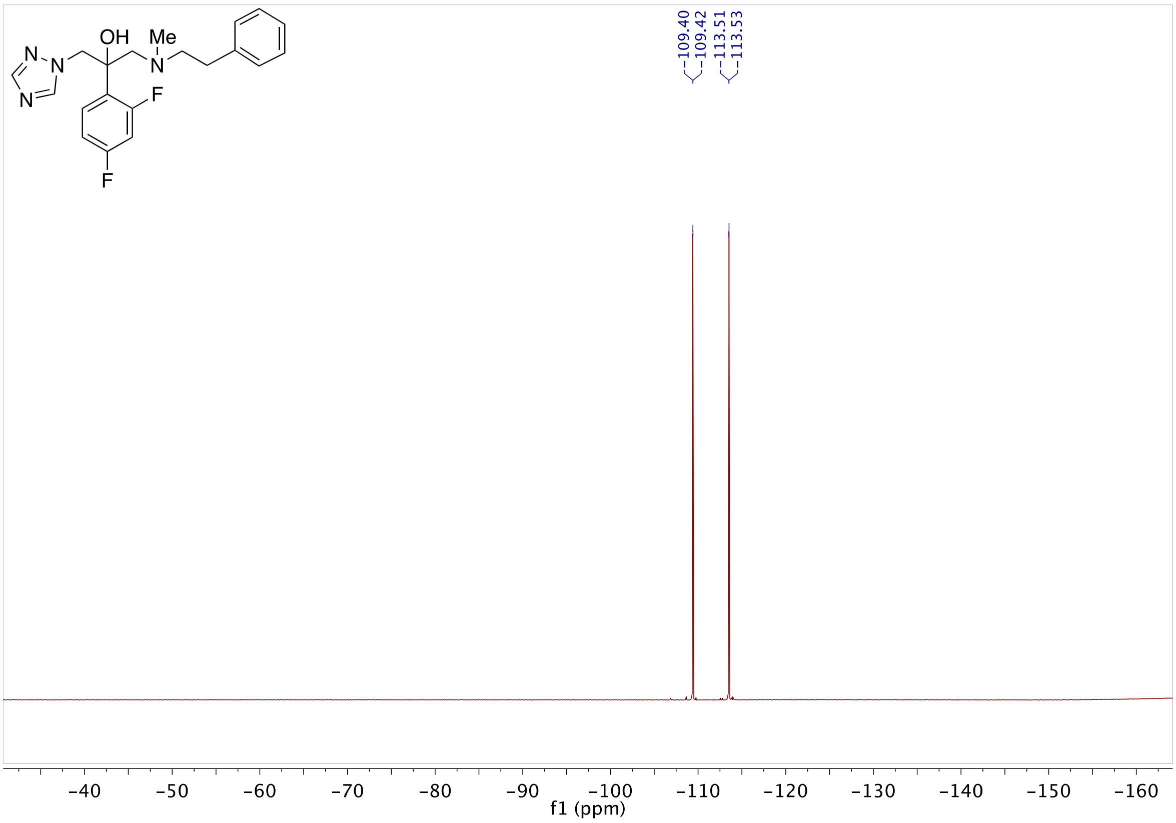

**Fig S82.** 19F NMR of **26** in CD3OD

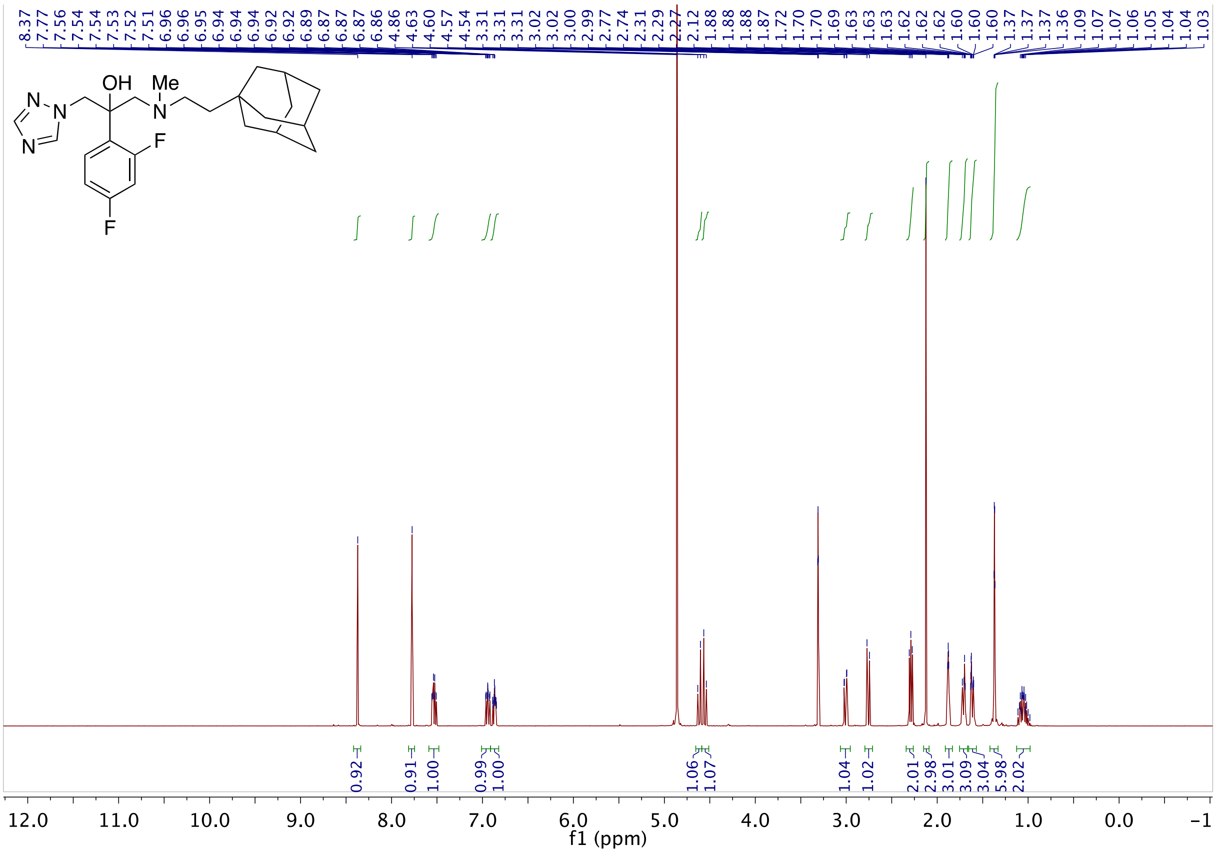

**Fig S83.** 1H NMR of **27** in CD3OD

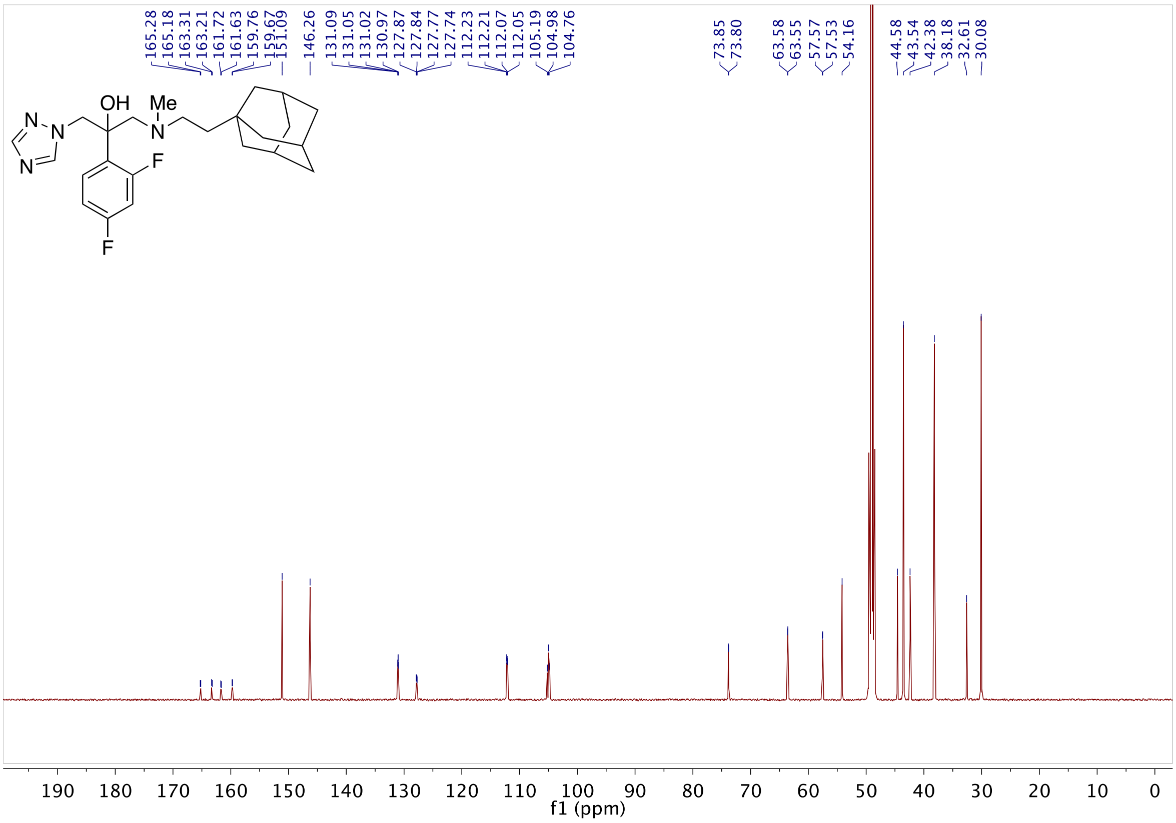

**Fig S84.** 13C NMR of **27** in CD3OD

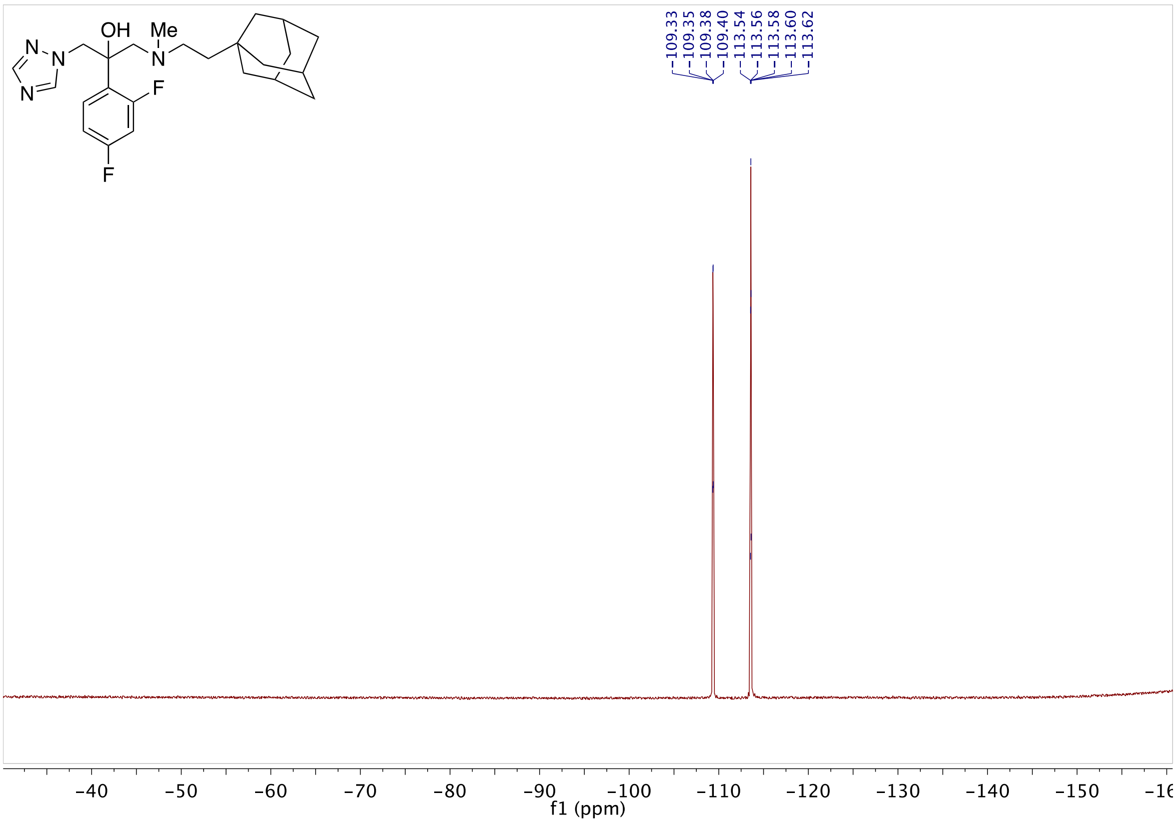

**Fig S85.** 19F NMR of **27** in CD3OD

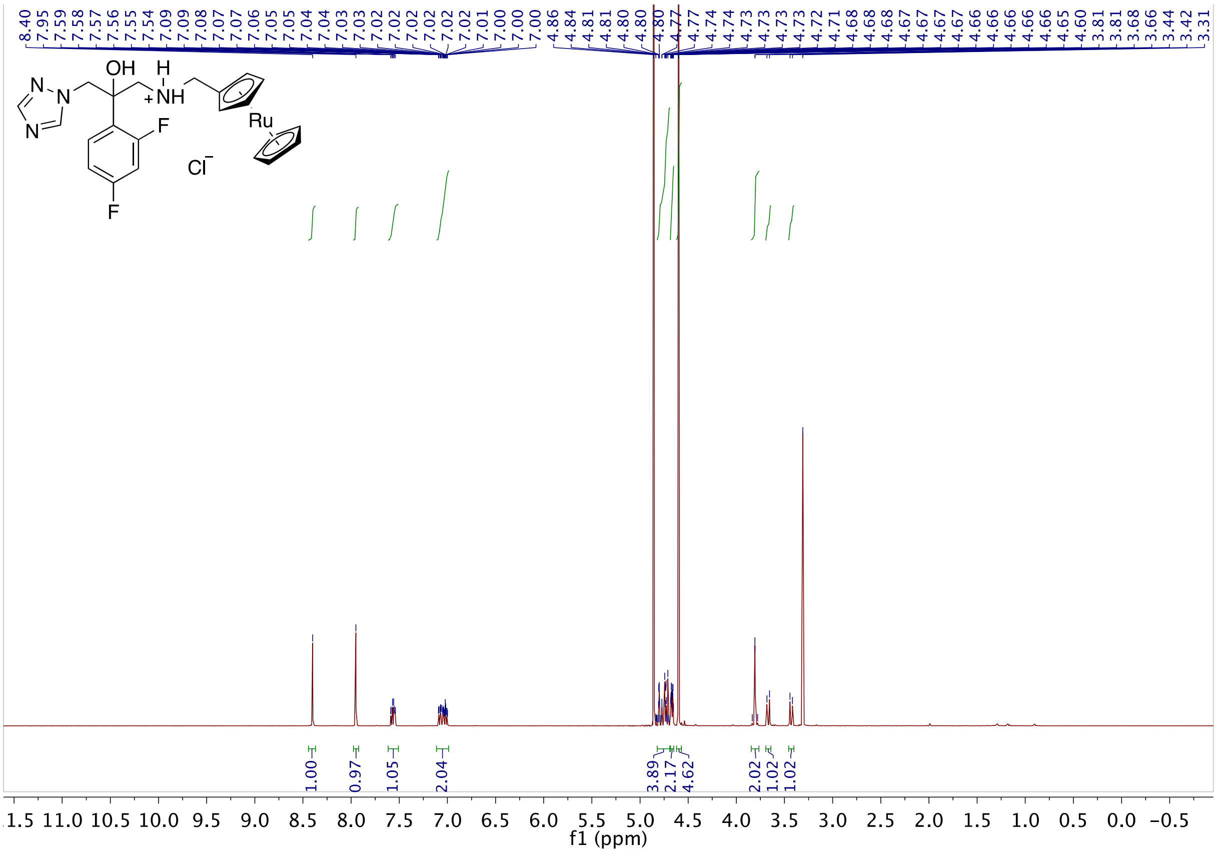

**Fig S86.** 1H NMR of **5a** in CD3OD

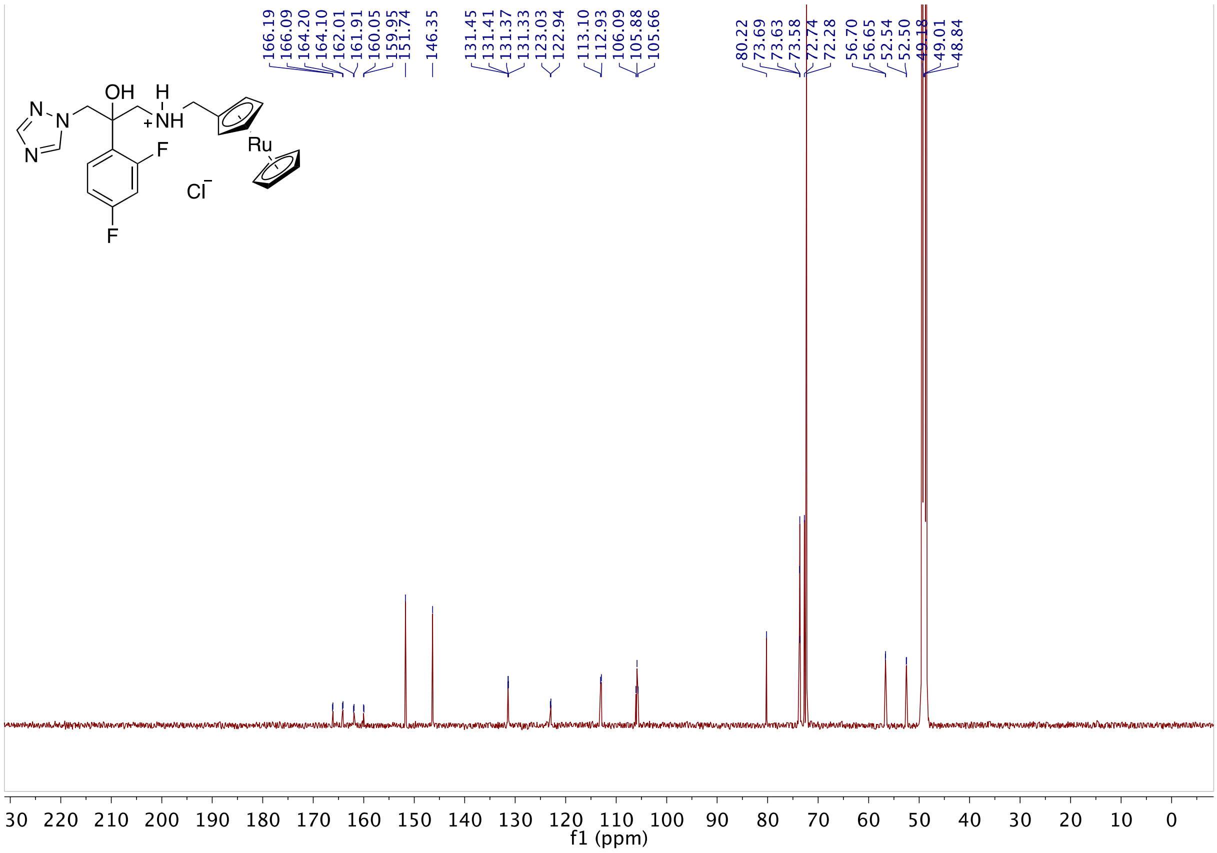

**Fig S87.** 13C NMR of **5a** in CD3OD

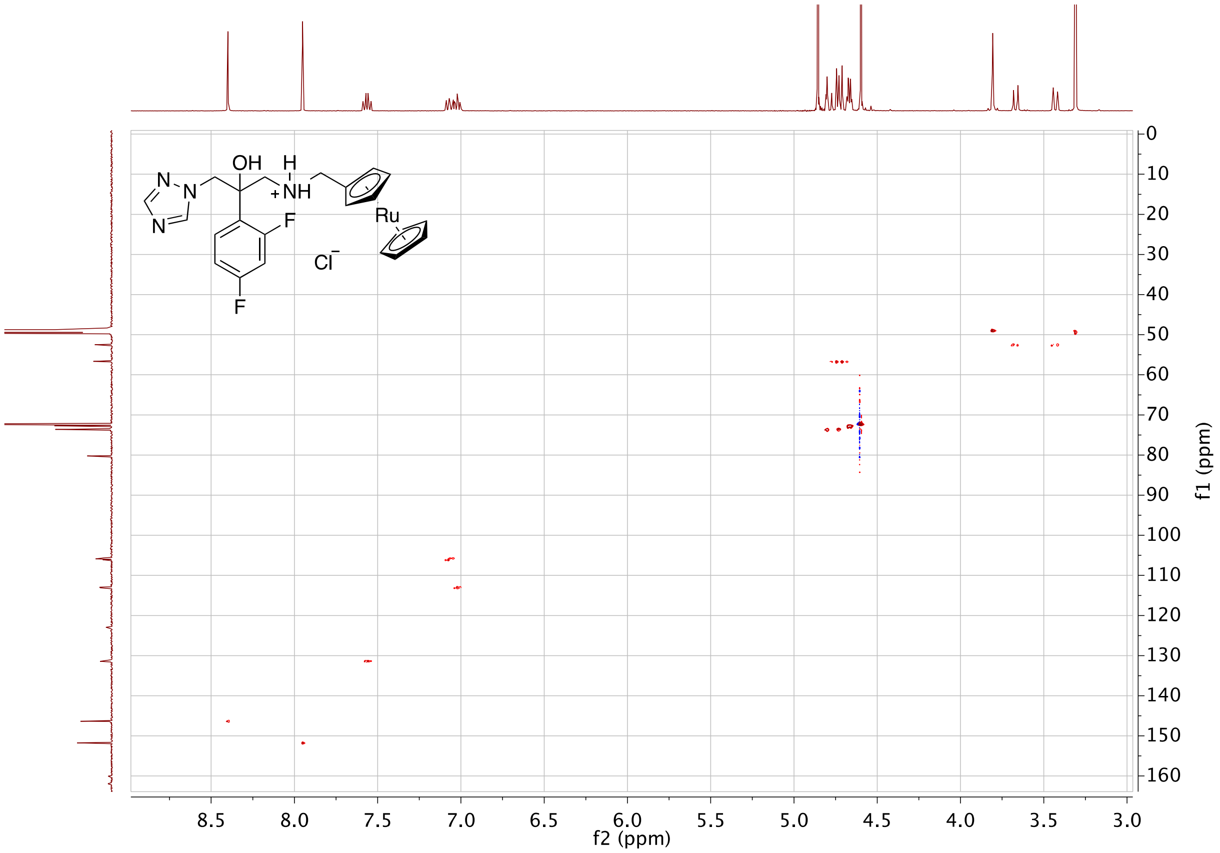

**Fig S88.** HSQC of **5a** in CD3OD

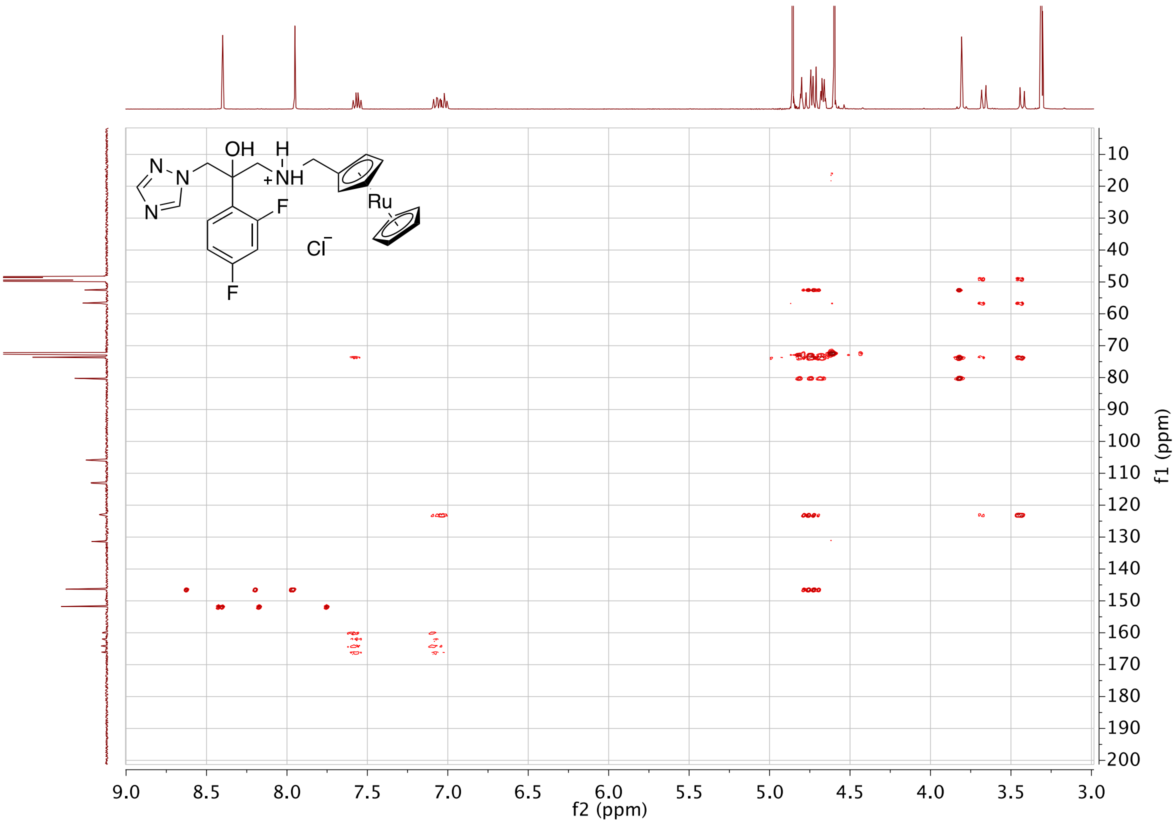

**Fig S89.** HMBC of **5a** in CD3OD

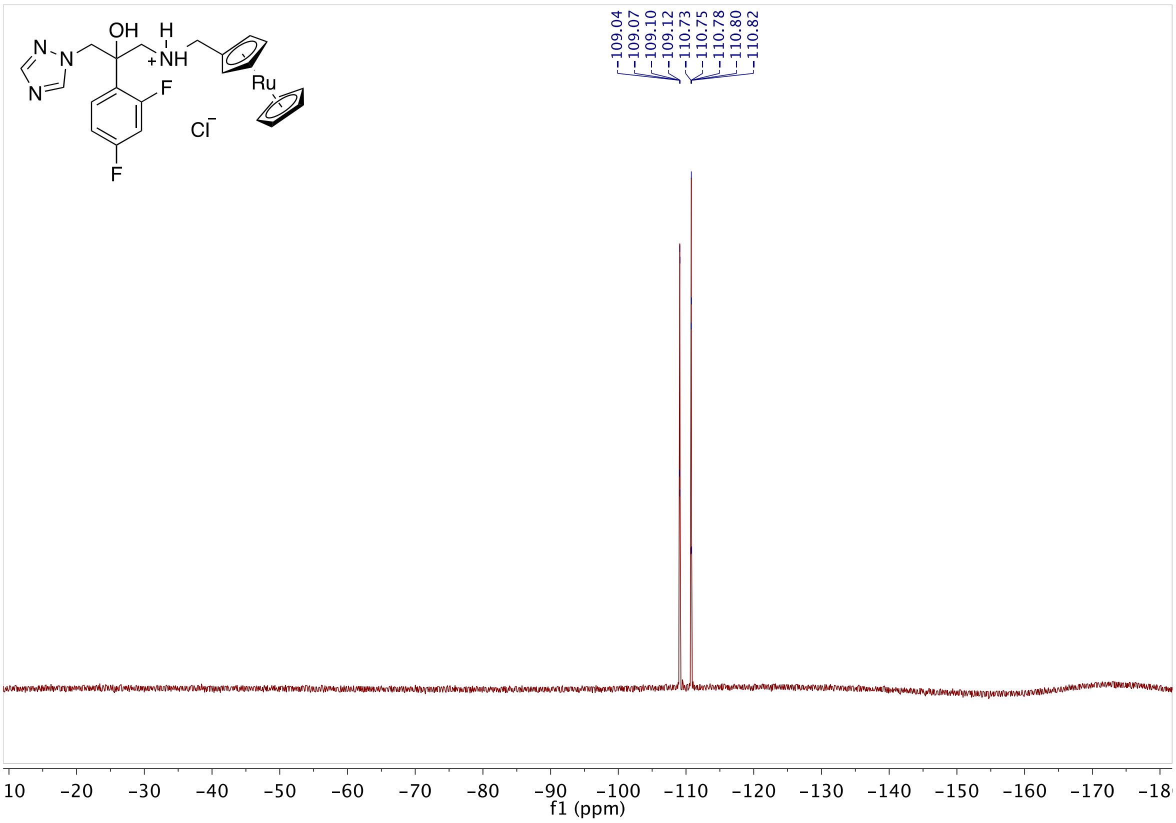

**Fig S90.** 19F NMR of **5a** in CD3OD

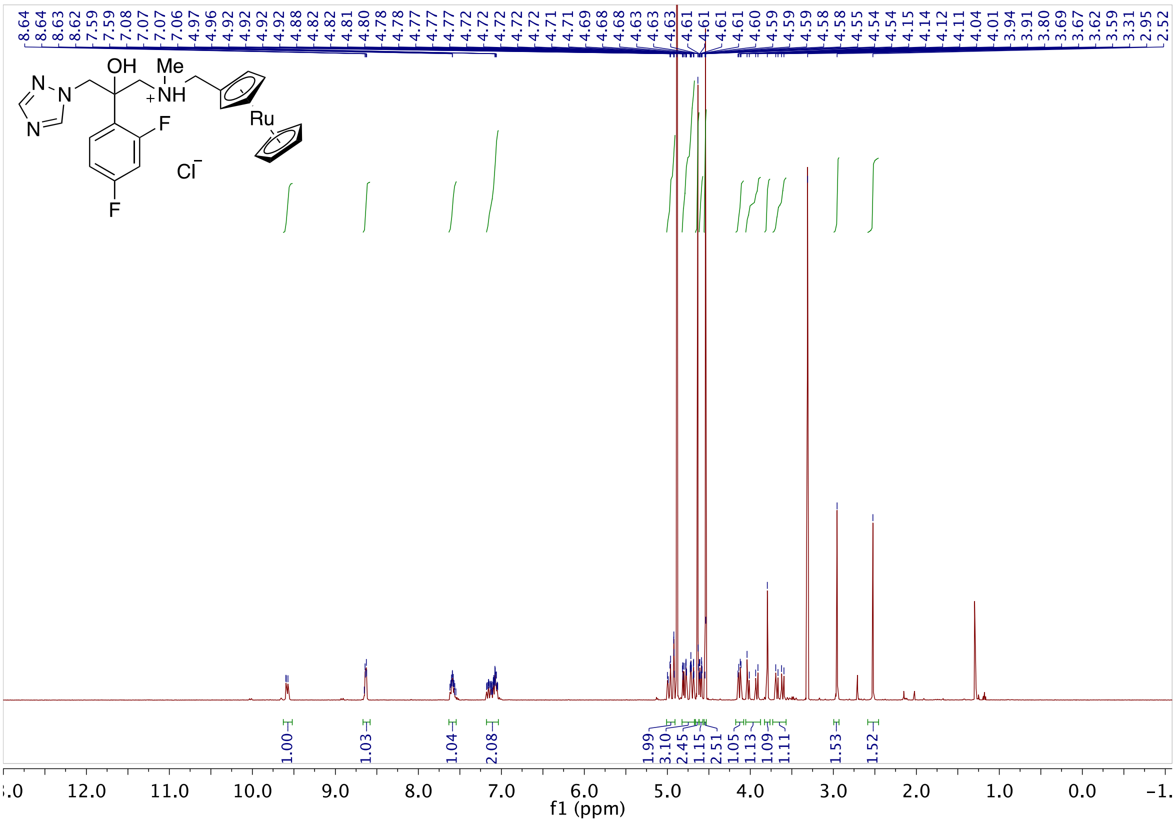

**Fig S91.** 1H NMR of **6a** in CD3OD

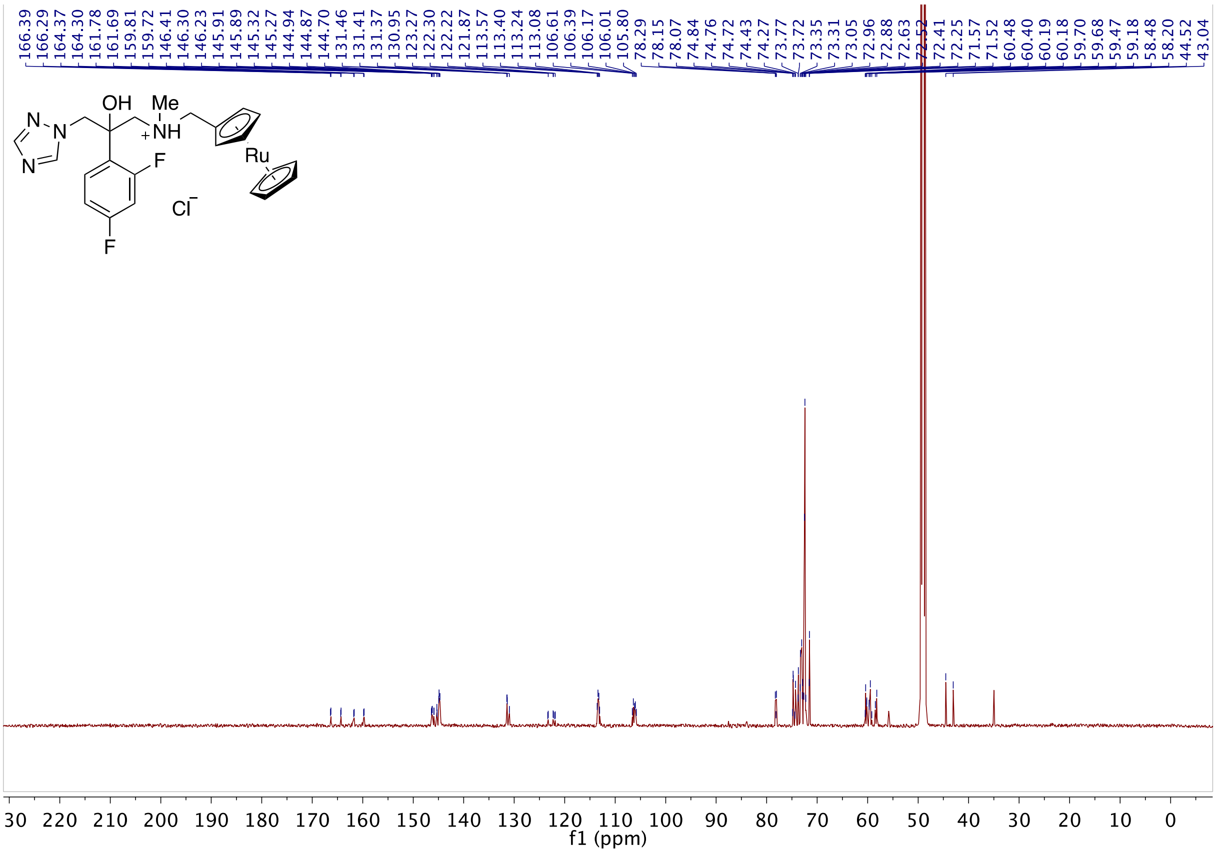

**Fig S92.** 13C NMR of **6a** in CD3OD

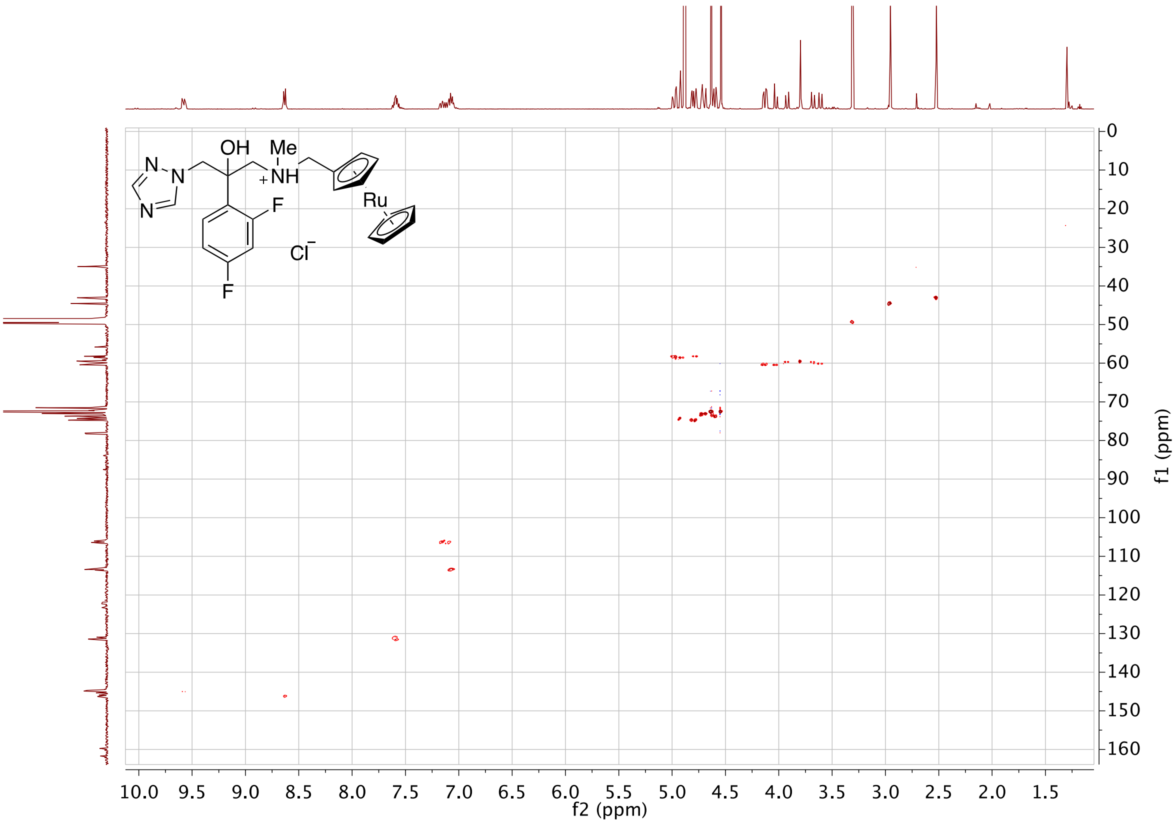

**Fig S93.** HSQC of **6a** in CD3OD

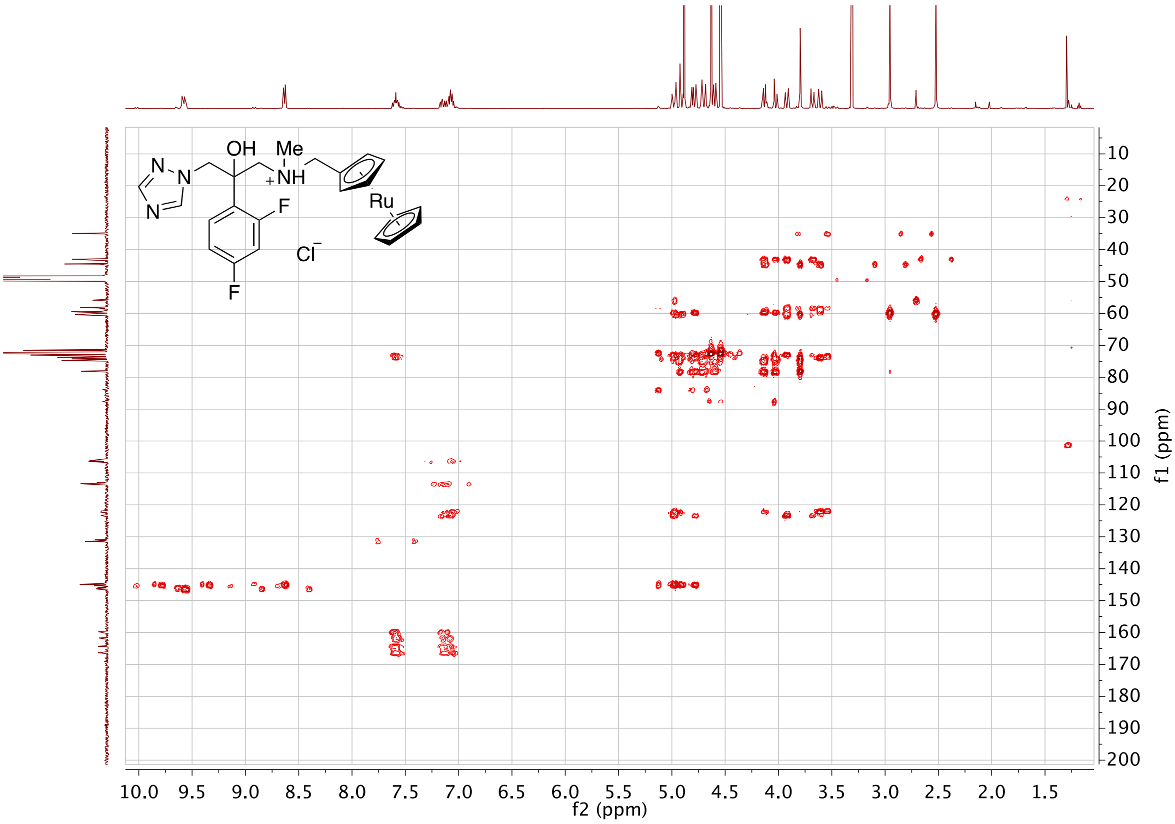

**Fig S94.** HMBC of **6a** in CD3OD

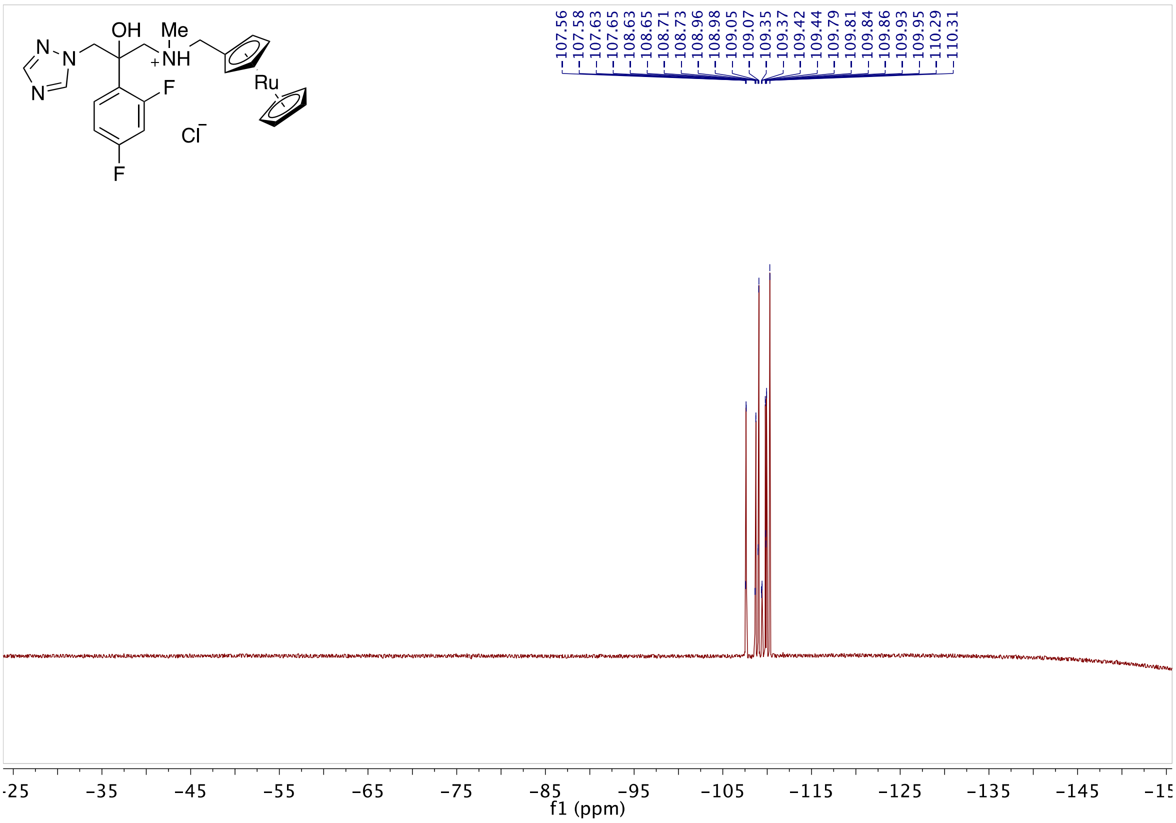

**Fig S95.** 19 F NMR of **6a** in CD3OD

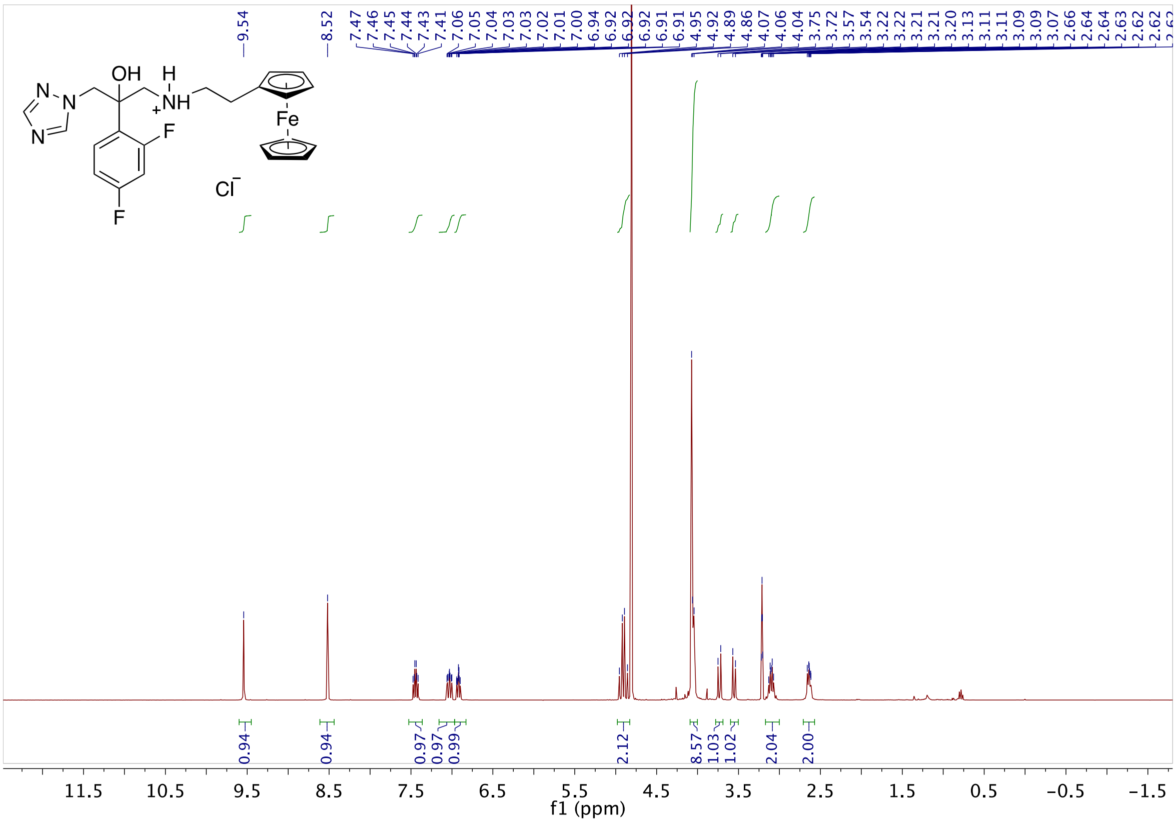

**Fig S96.** 1H NMR of **7a** in DMSO-d$_6$

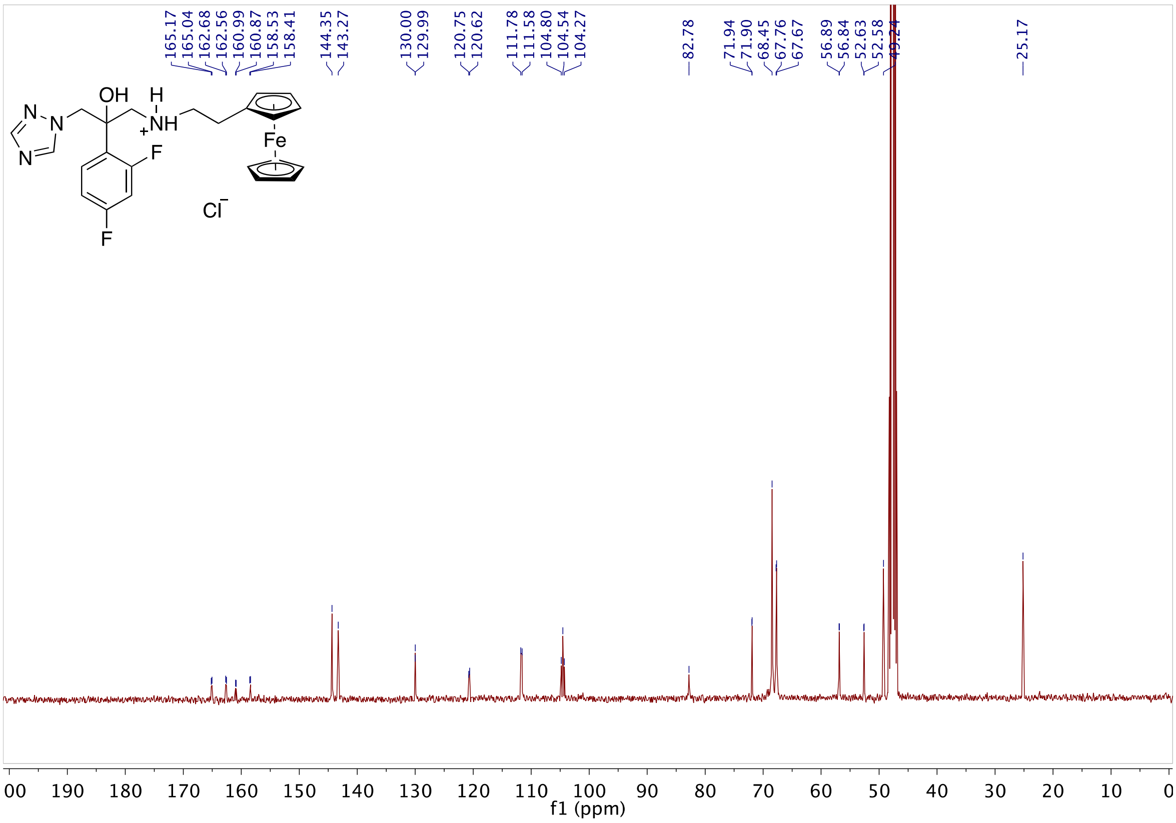

**Fig S97.** 13C NMR of **7a** in DMSO-d6

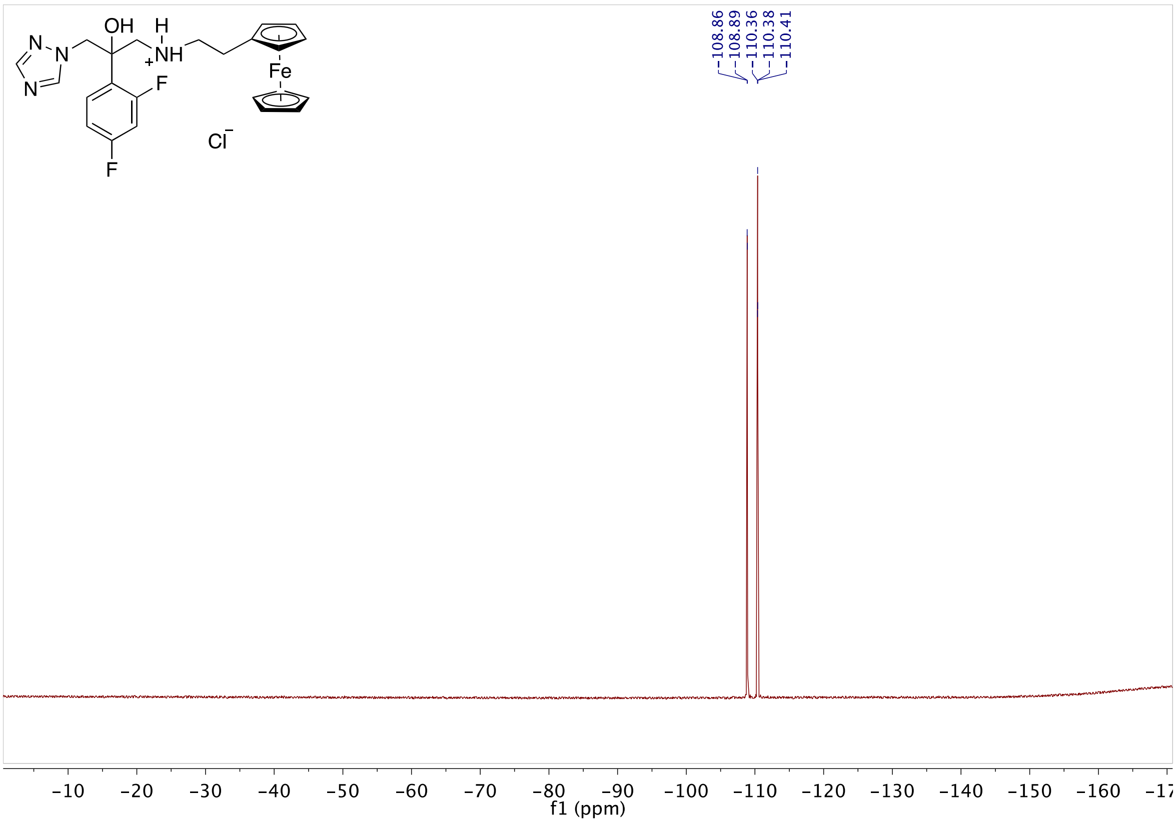

**Fig S98.** 19F NMR of **7a** in DMSO-d6

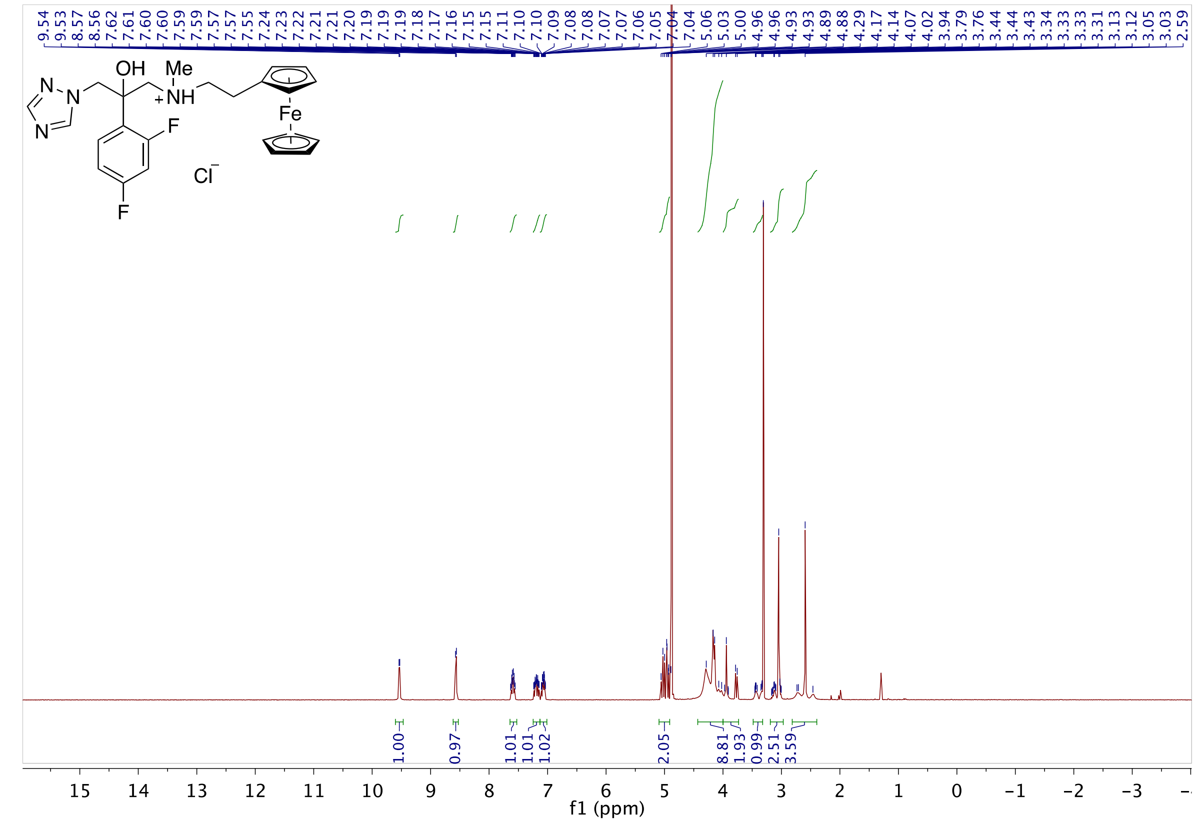

**Fig S99.** 1H NMR of **8a** in $CD_3OD$

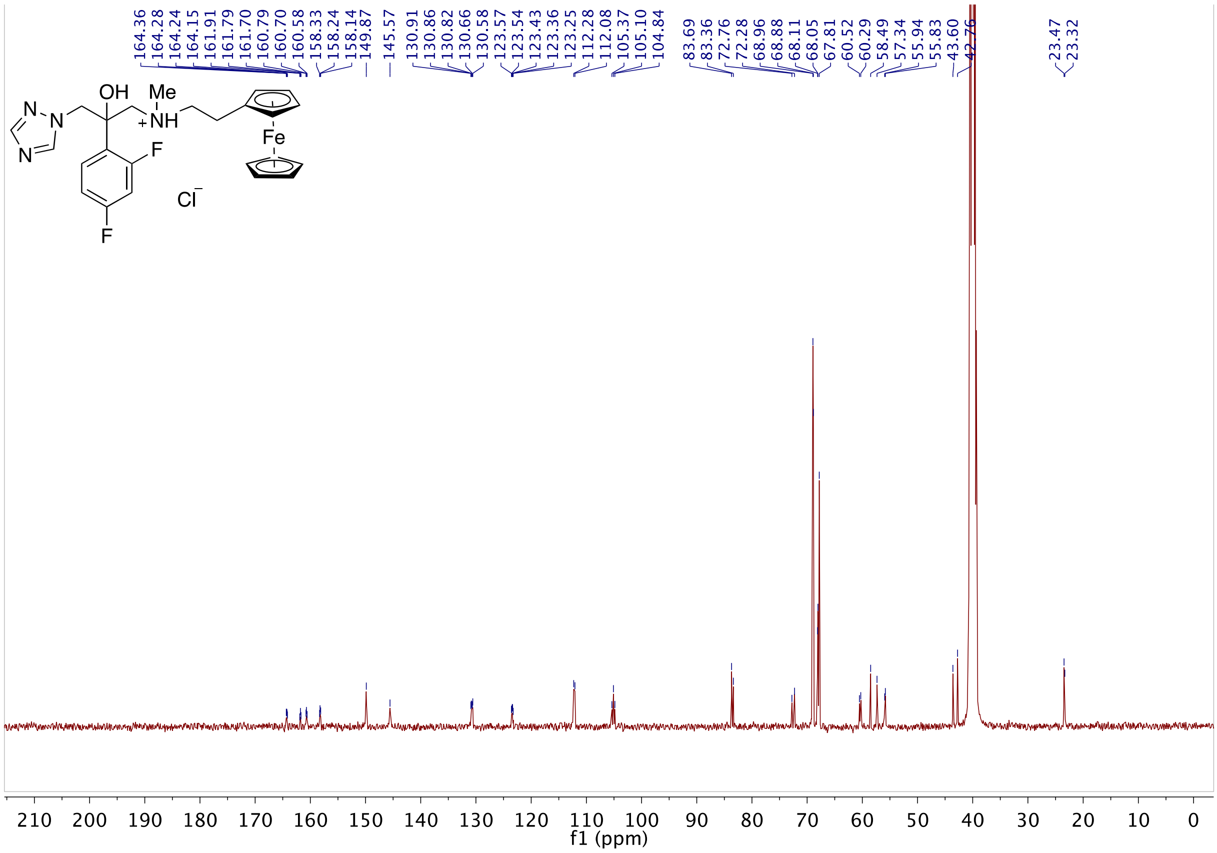

**Fig S100.** 13C NMR of **8a** in $CD_3OD$

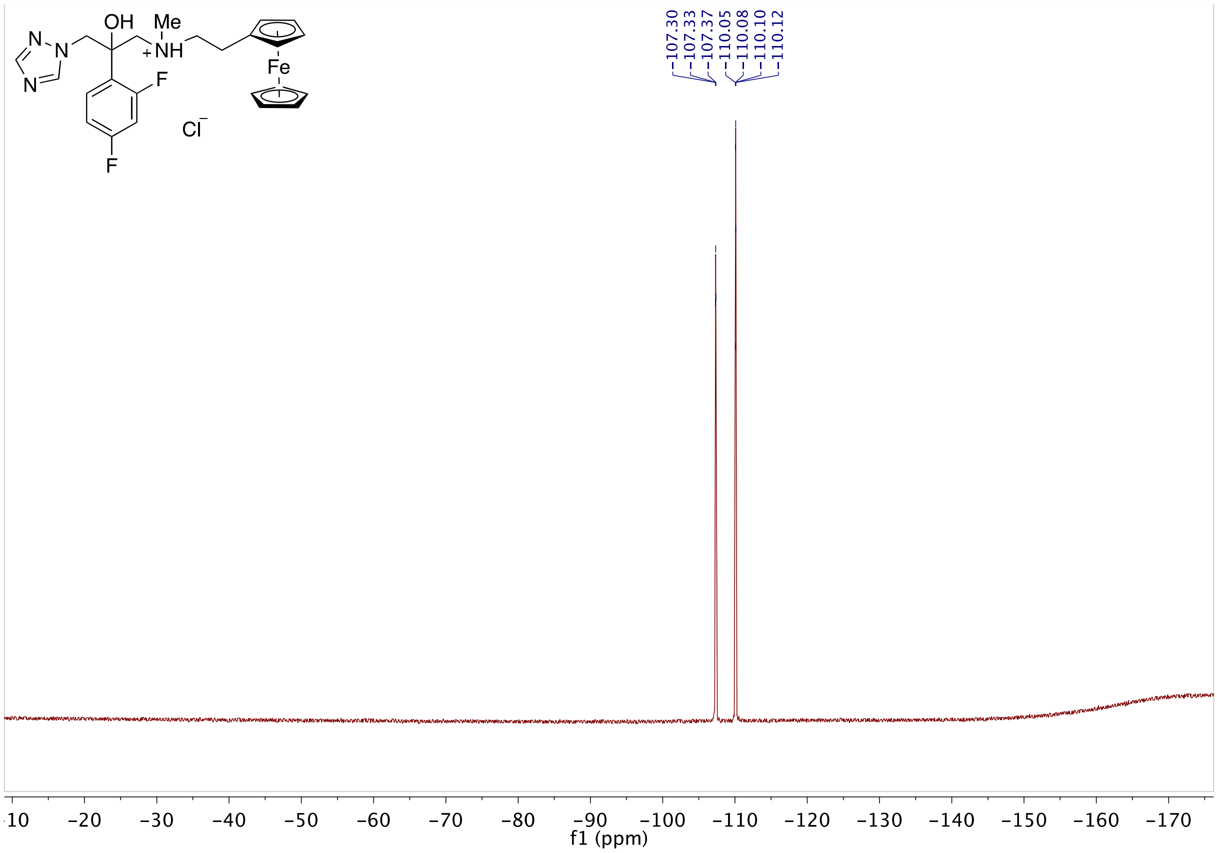

**Fig S101.** 19F NMR of **8a** in $CD_3OD$

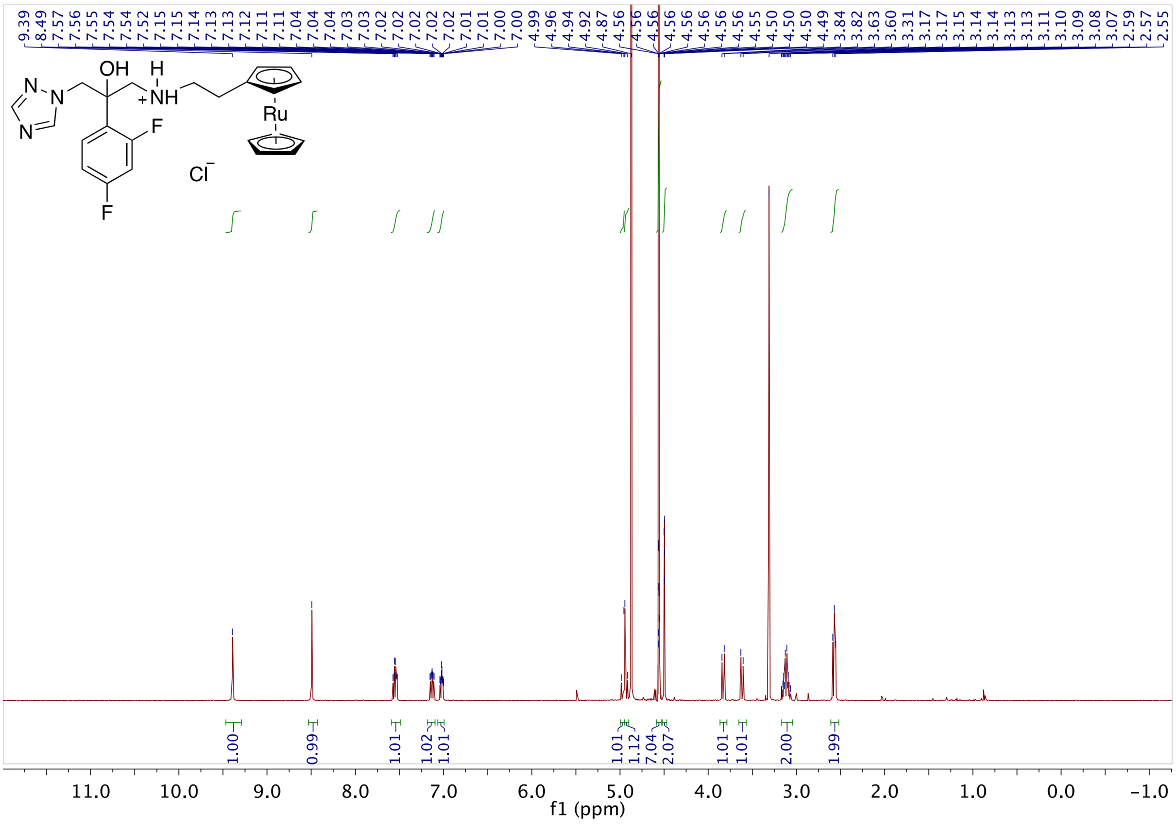

**Fig S102.** 1H NMR of **9a** in $CD_3OD$

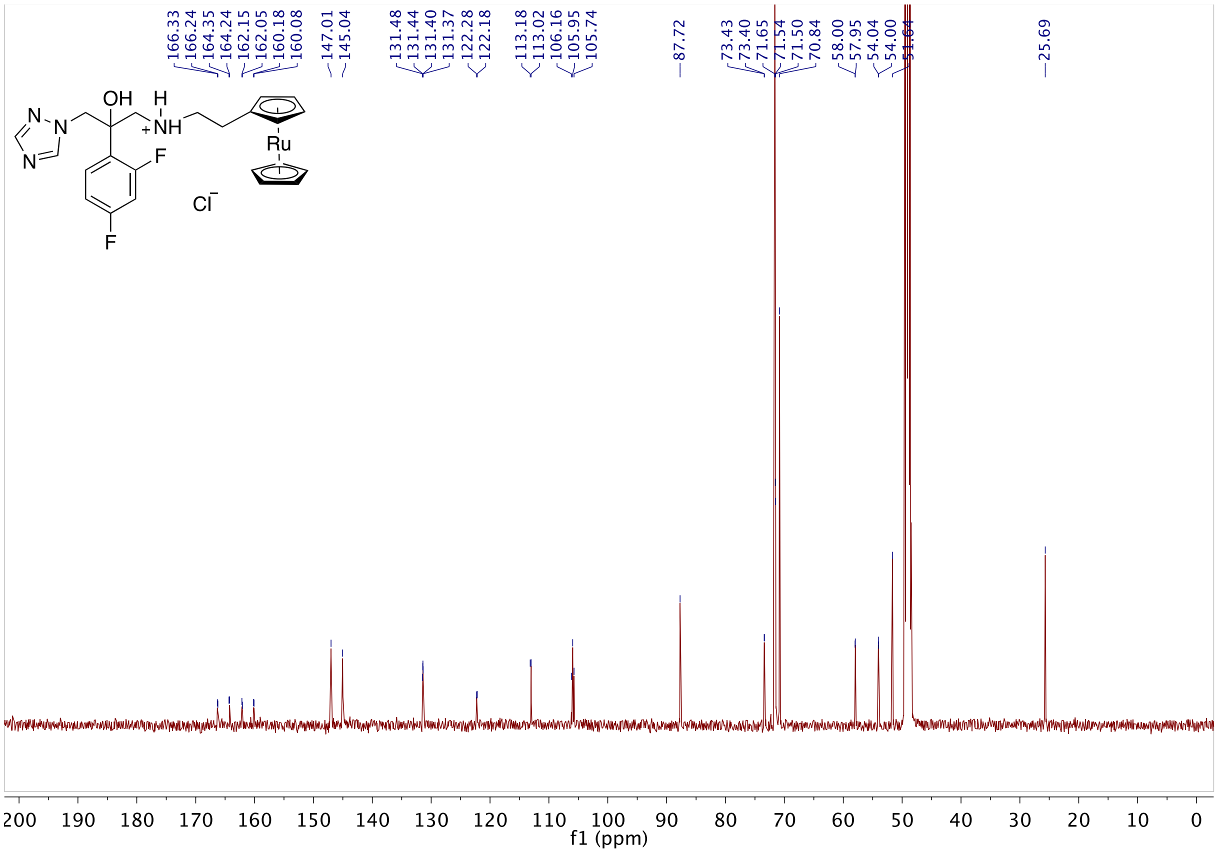

**Fig S103.** 13C NMR of **9a** in $CD_3OD$

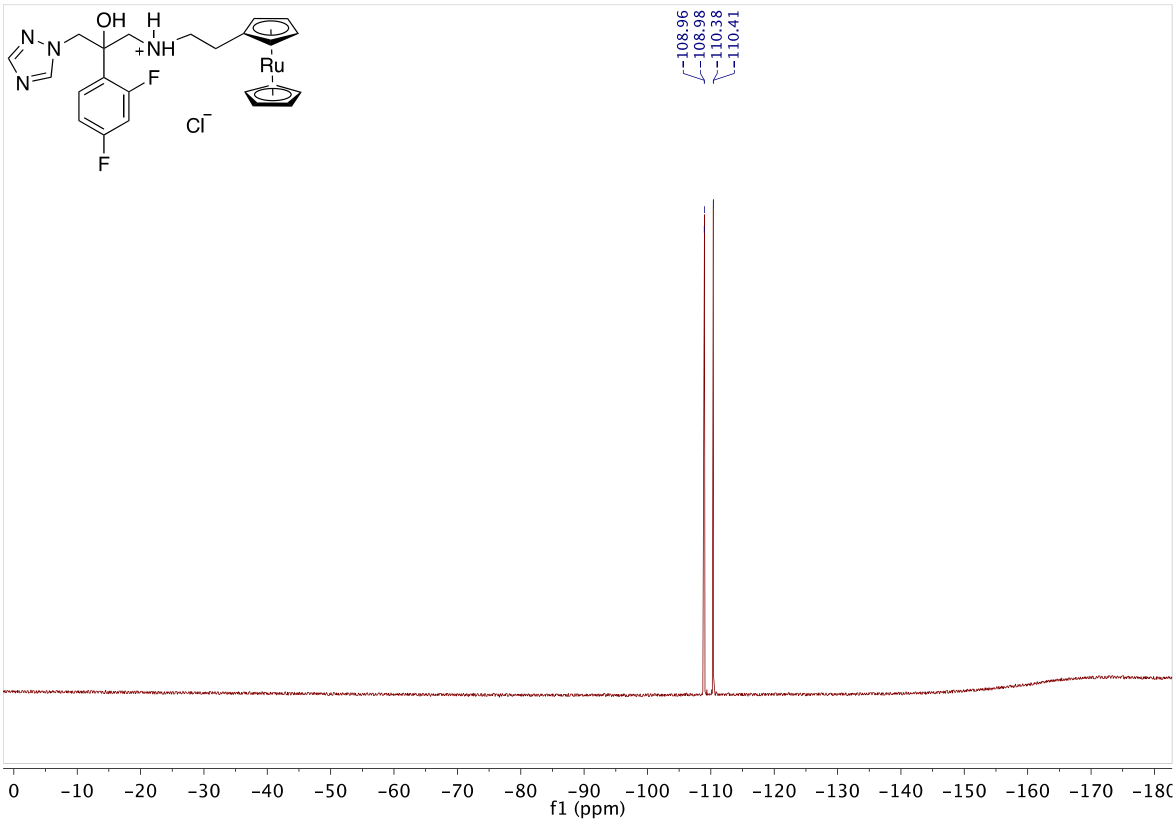

**Fig S104.** 19F NMR of **9a** in $CD_3OD$

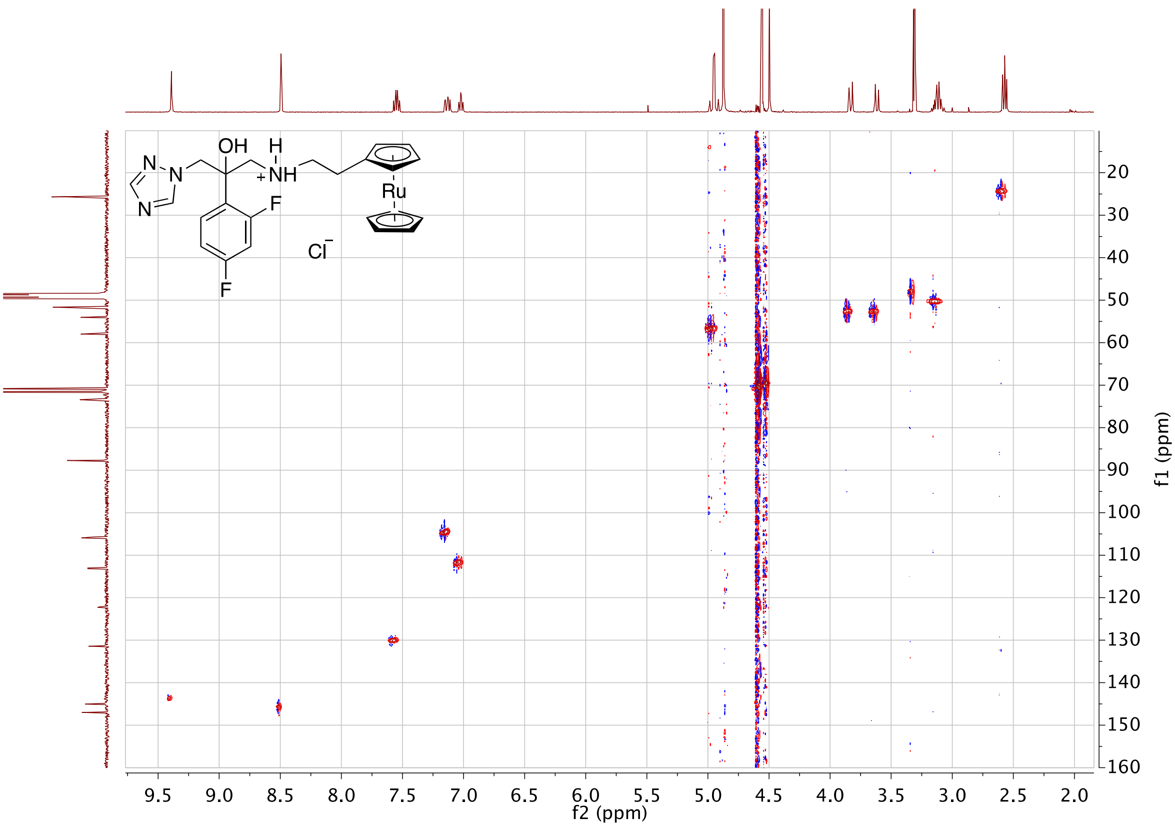

**Fig S105.** HSQC of **9a** in $CD_3OD$

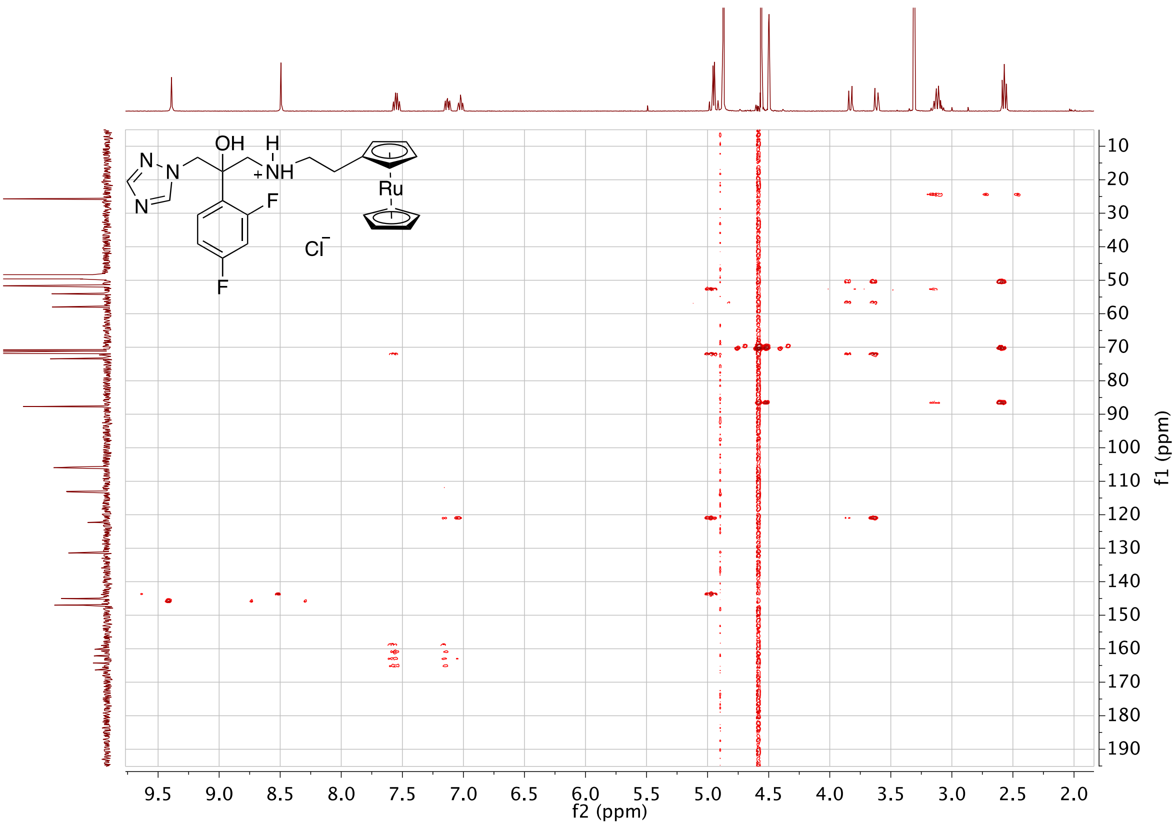

**Fig S106.** HMBC of **9a** in $CD_3OD$

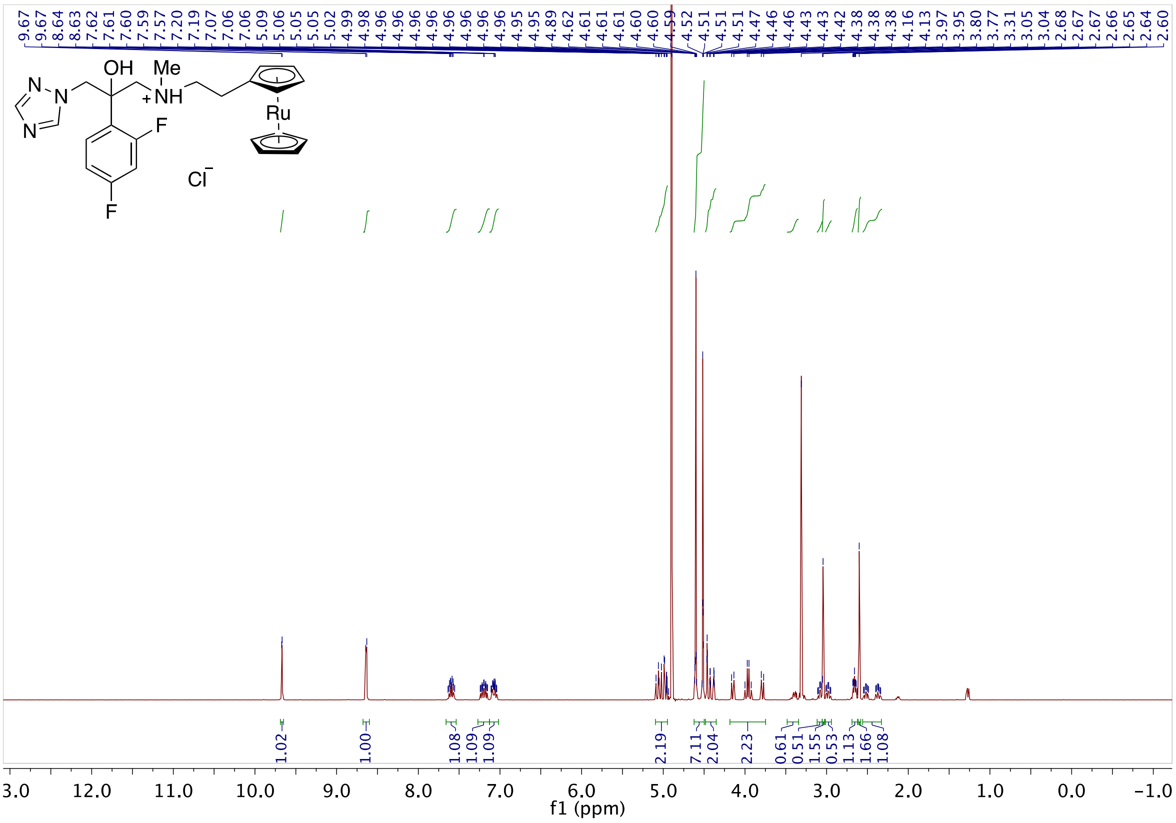

**Fig S107.** 1H NMR of **10a** in $CD_3OD$

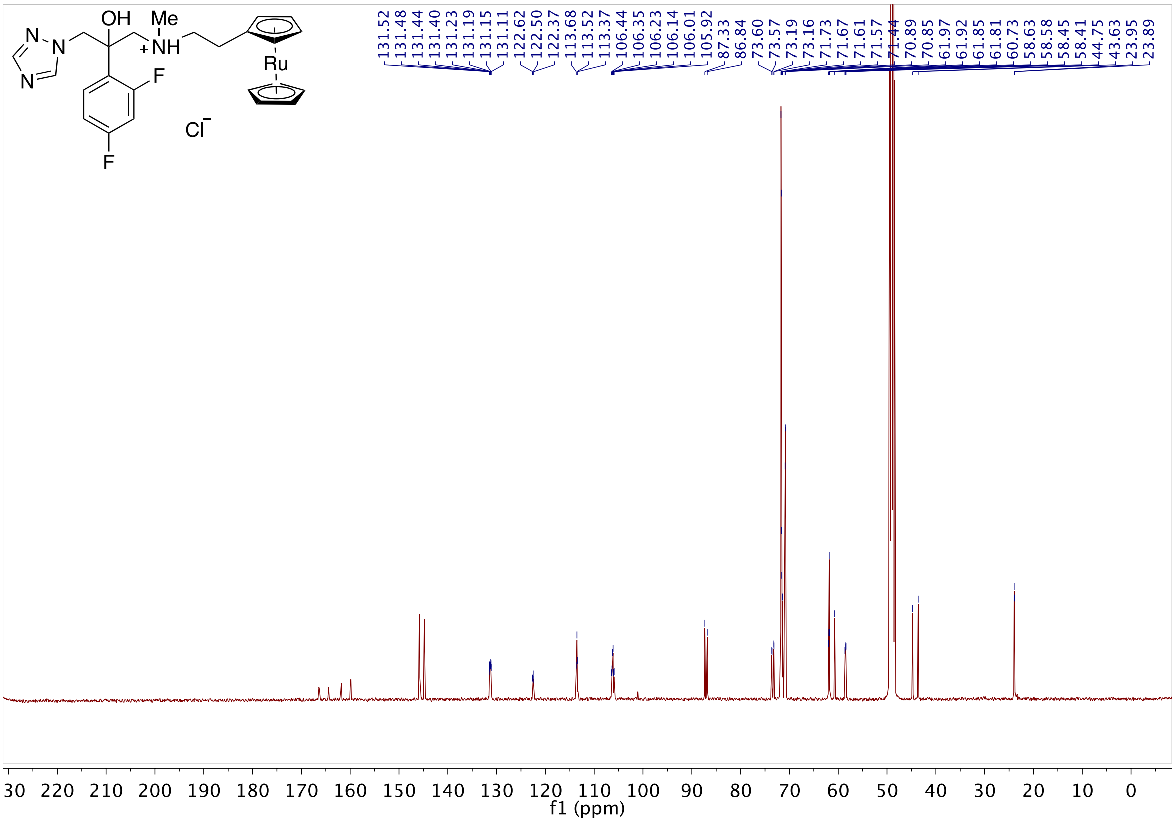

**Fig S108.** 13C NMR of **10a** in $CD_3OD$

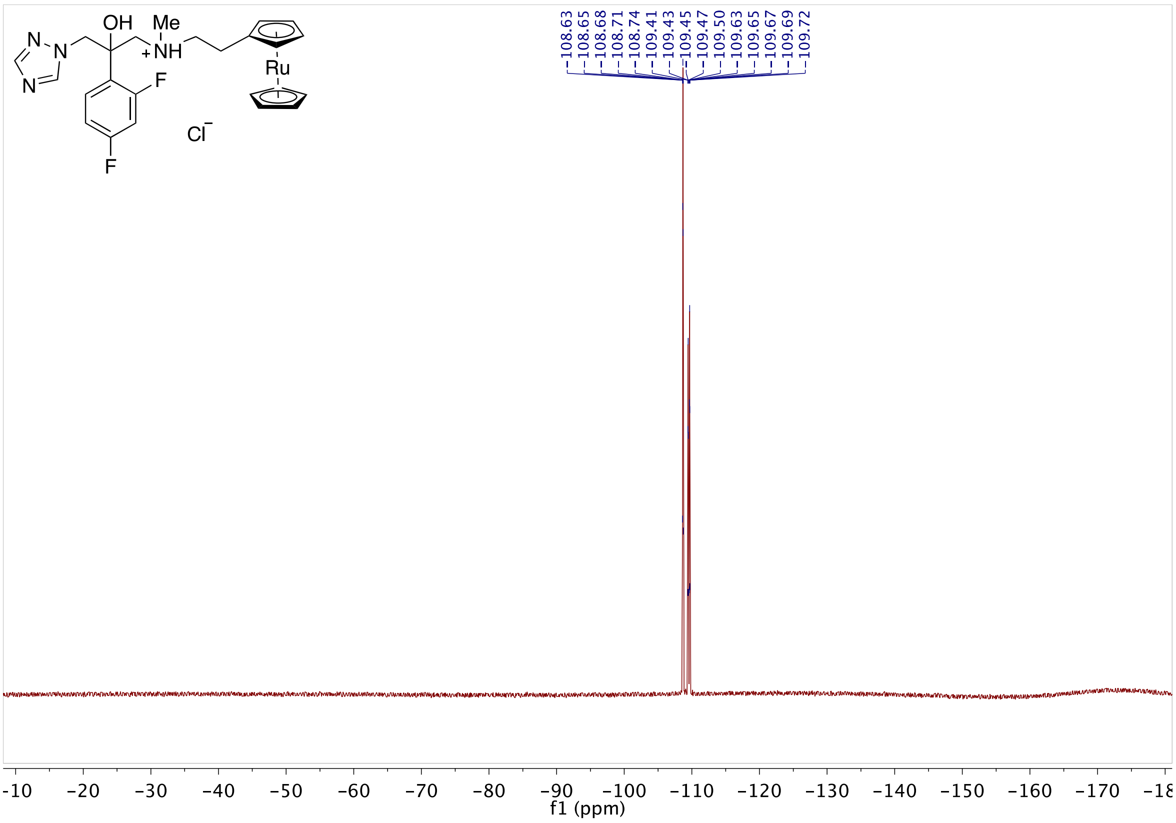

**Fig S109.** 19F NMR of **10a** in $CD_3OD$

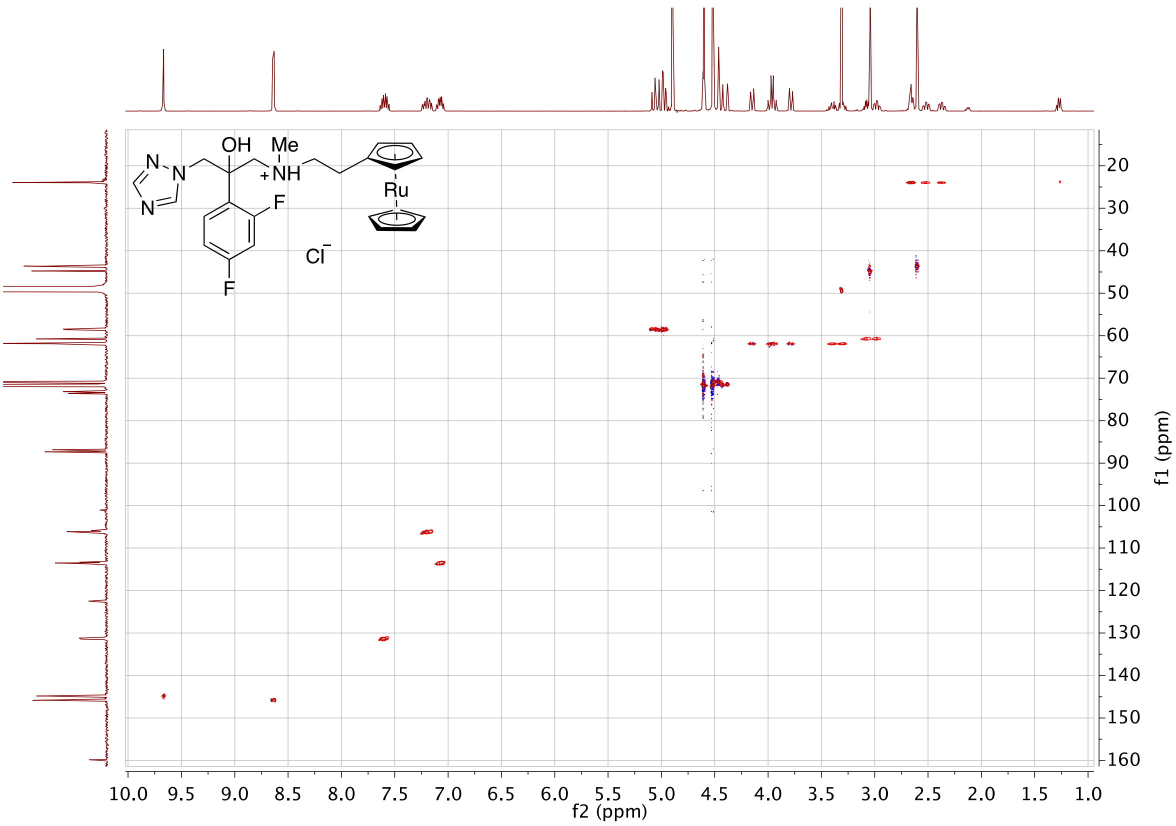

**Fig S110.** HSQC of **10a** in $CD_3OD$

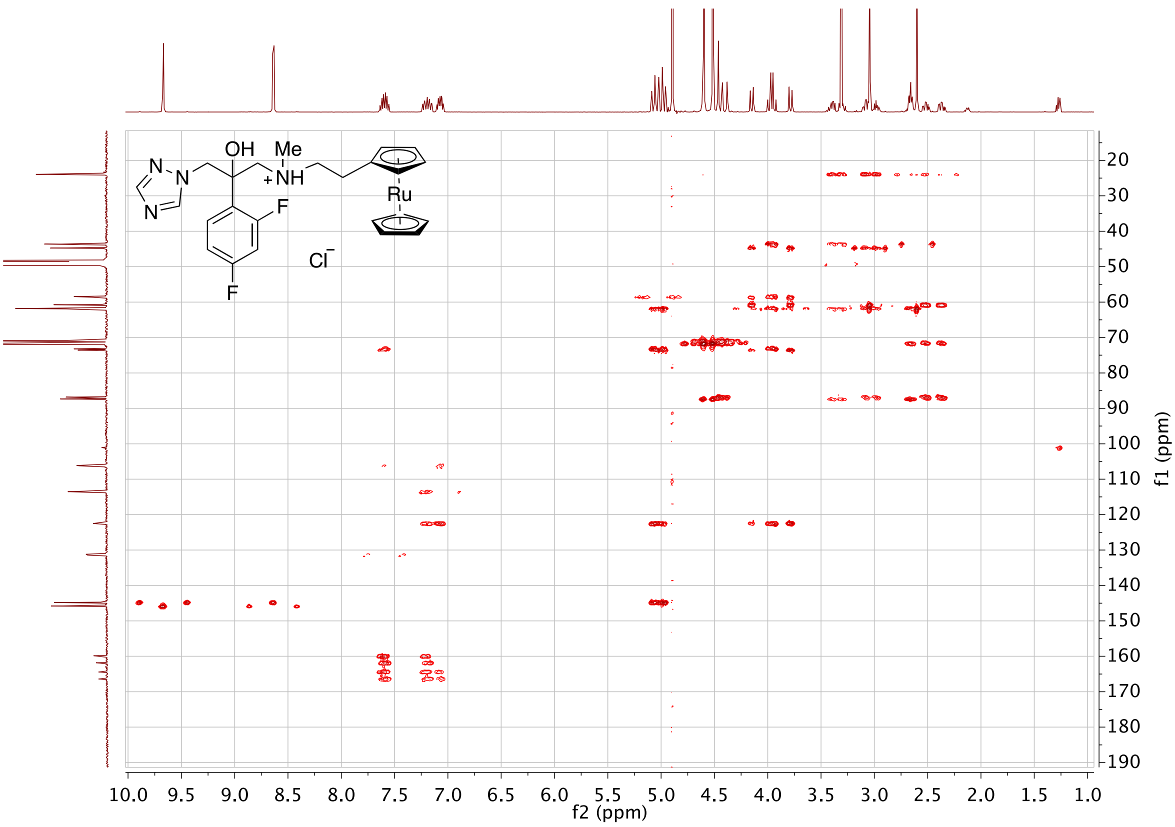

**Fig S111.** HMBC of **10a** in CD3OD

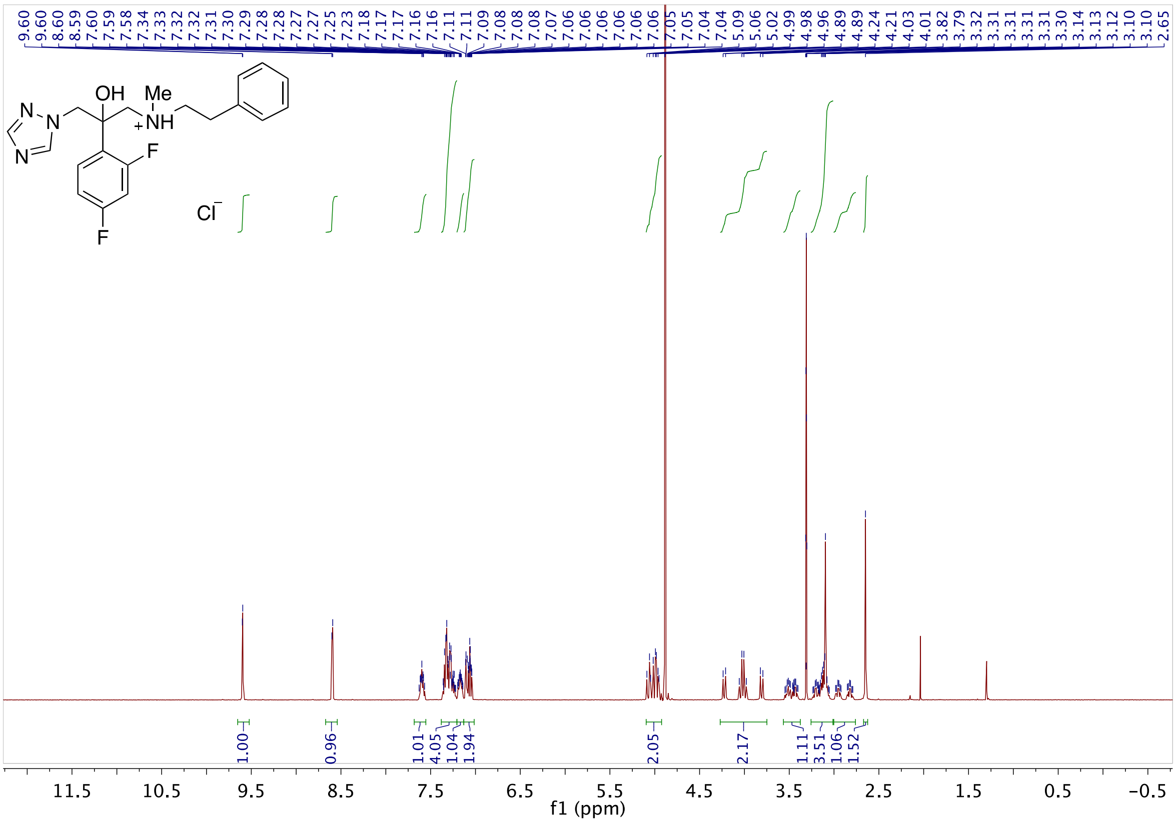

**Fig S112.** 1H NMR of **11a** in CD3OD

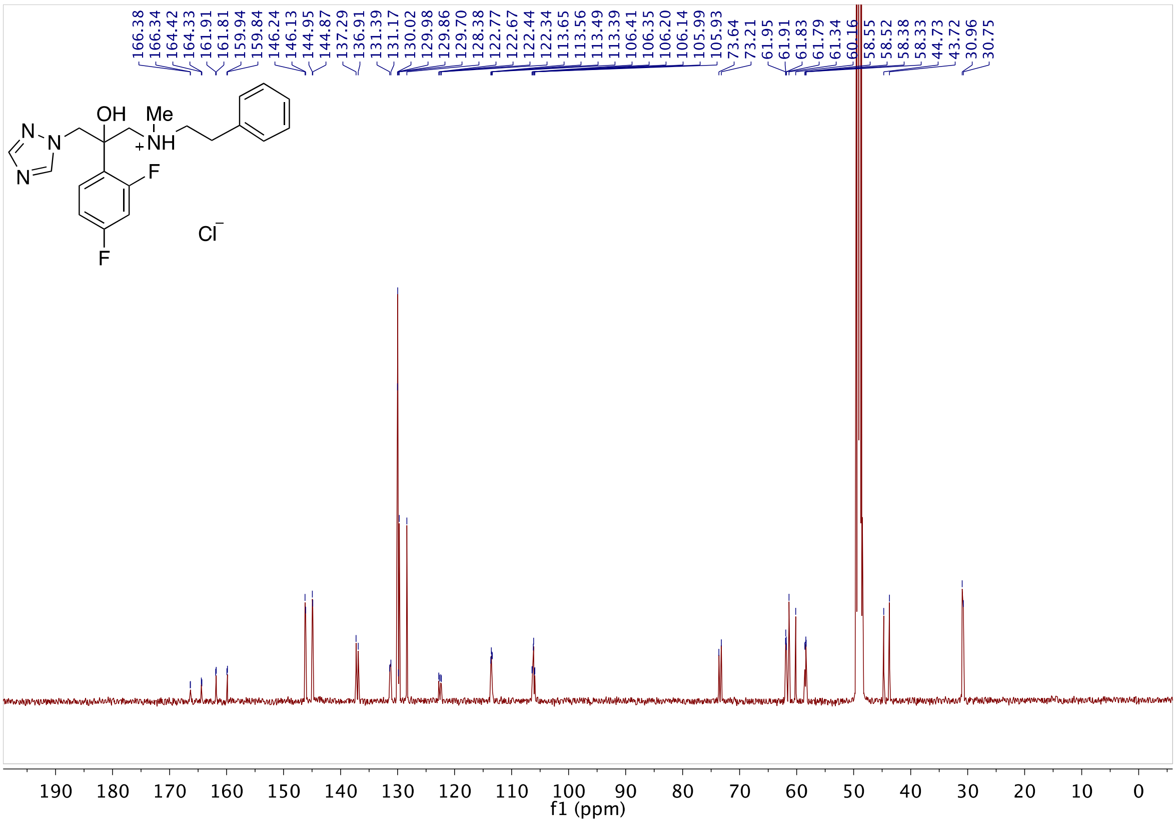

**Fig S113.** 13C NMR of **11a** in $CD_3OD$

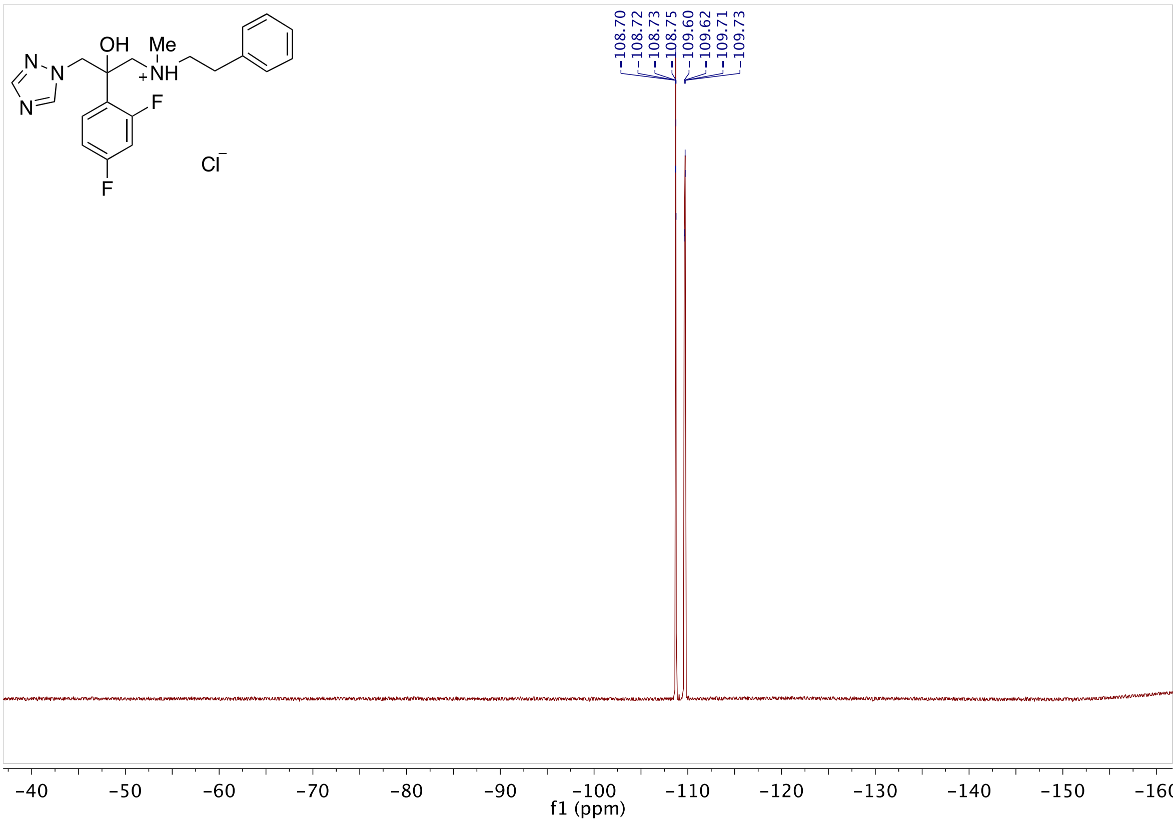

**Fig S114.** 19F NMR of **11a** in $CD_3OD$

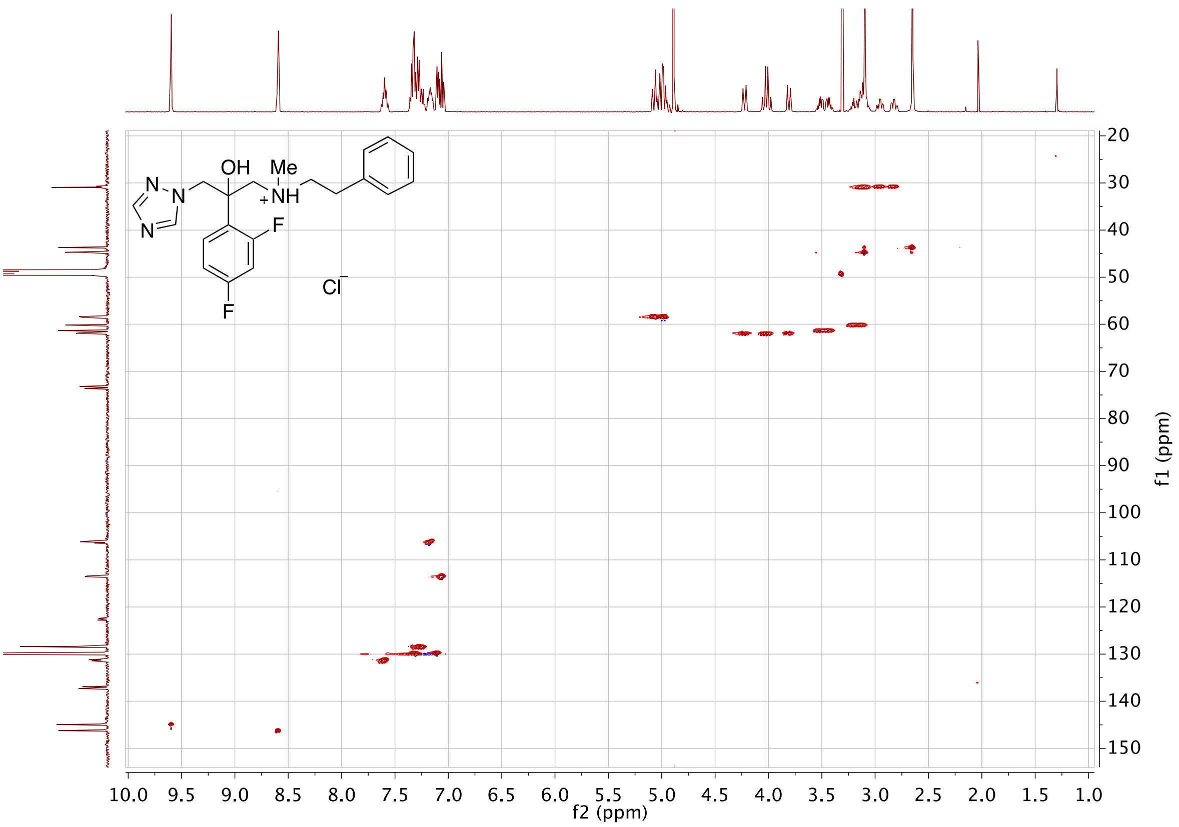

**Fig S115.** HSQC of **11a** in $CD_3OD$

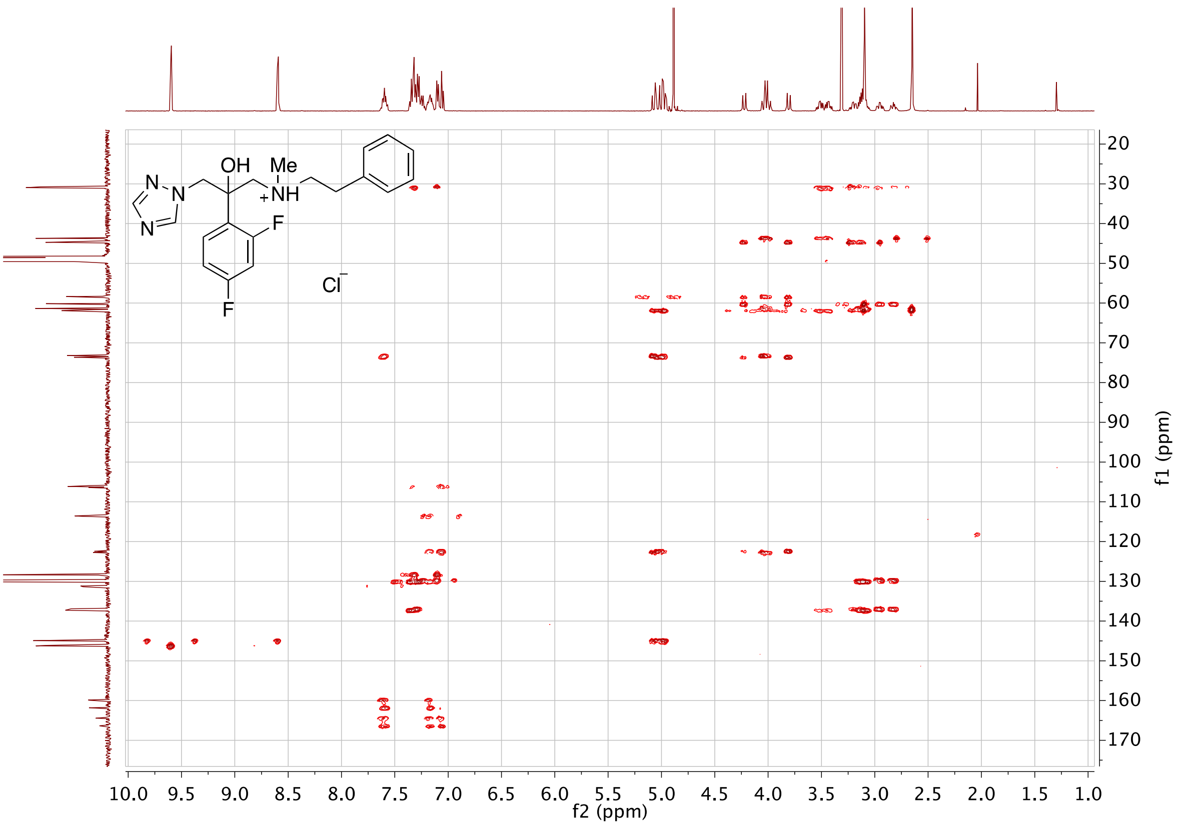

**Fig S116.** HMBC of **11a** in $CD_3OD$

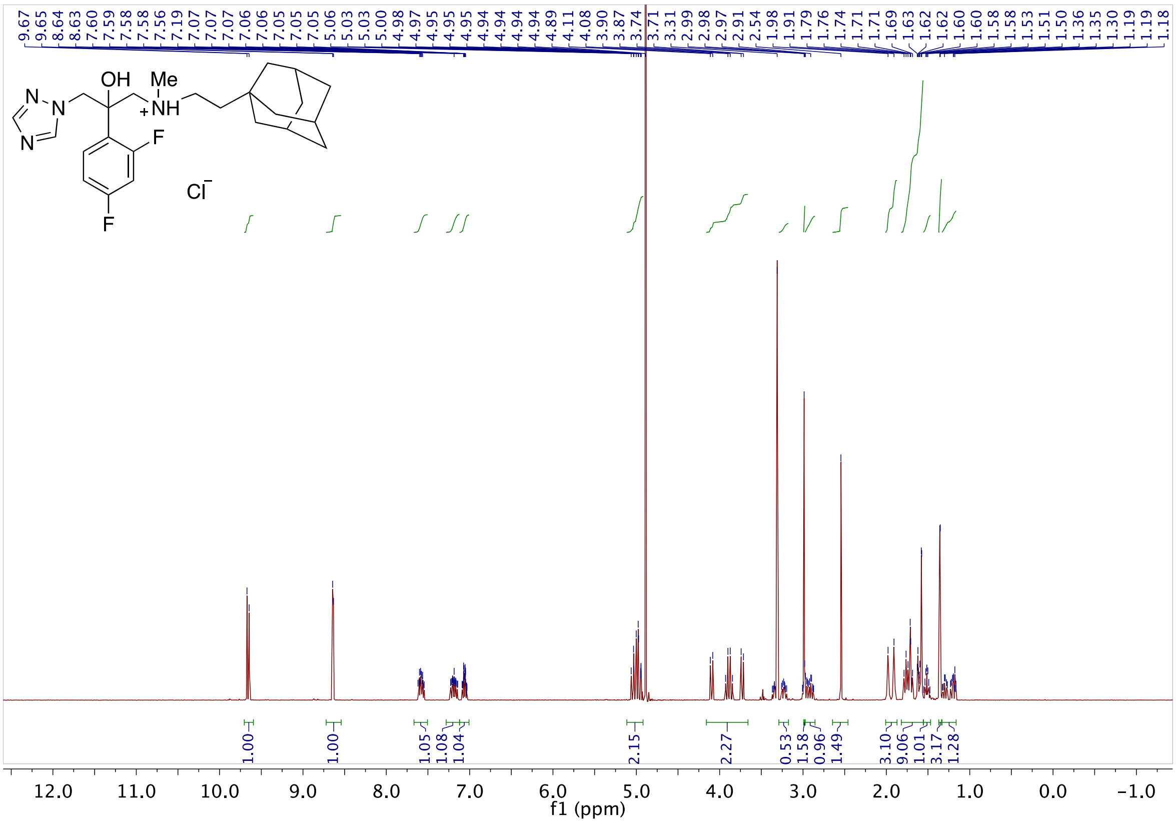

**Fig S117.** 1H NMR of **12a** in $CD_3OD$

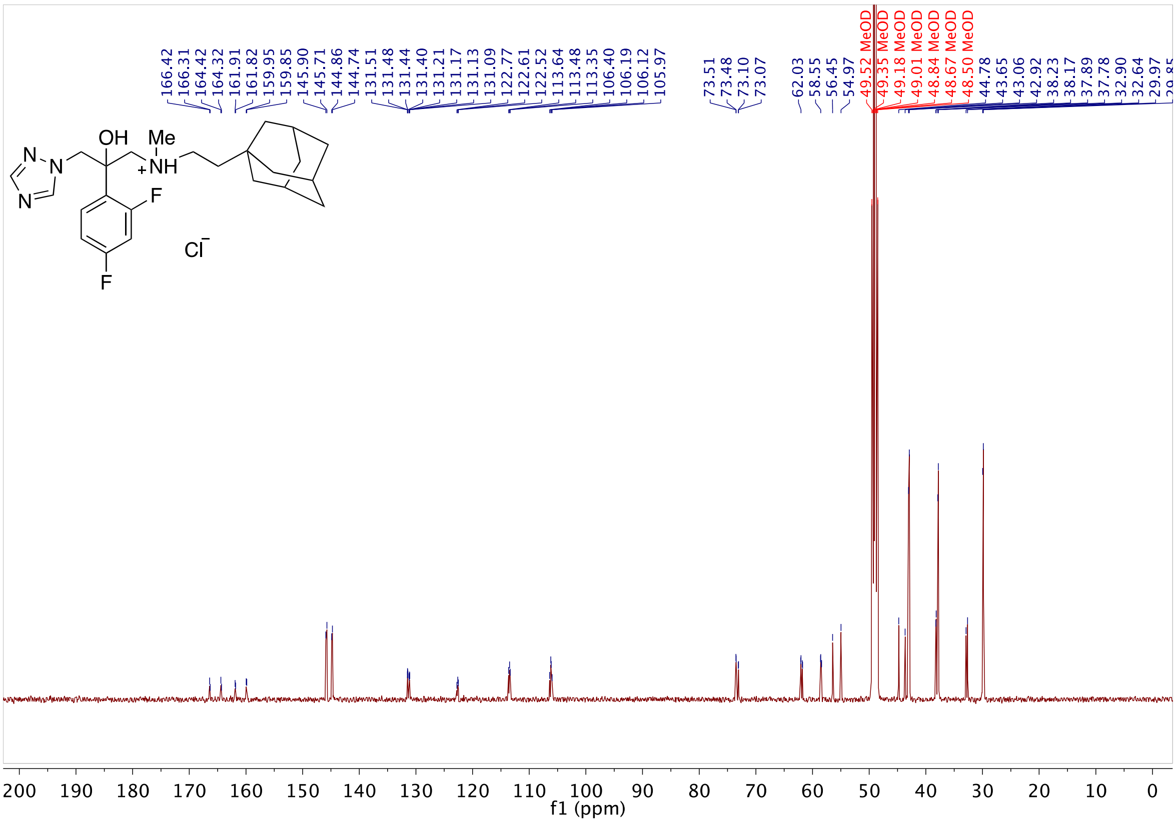

**Fig S118.** 13C NMR of **12a** in $CD_3OD$

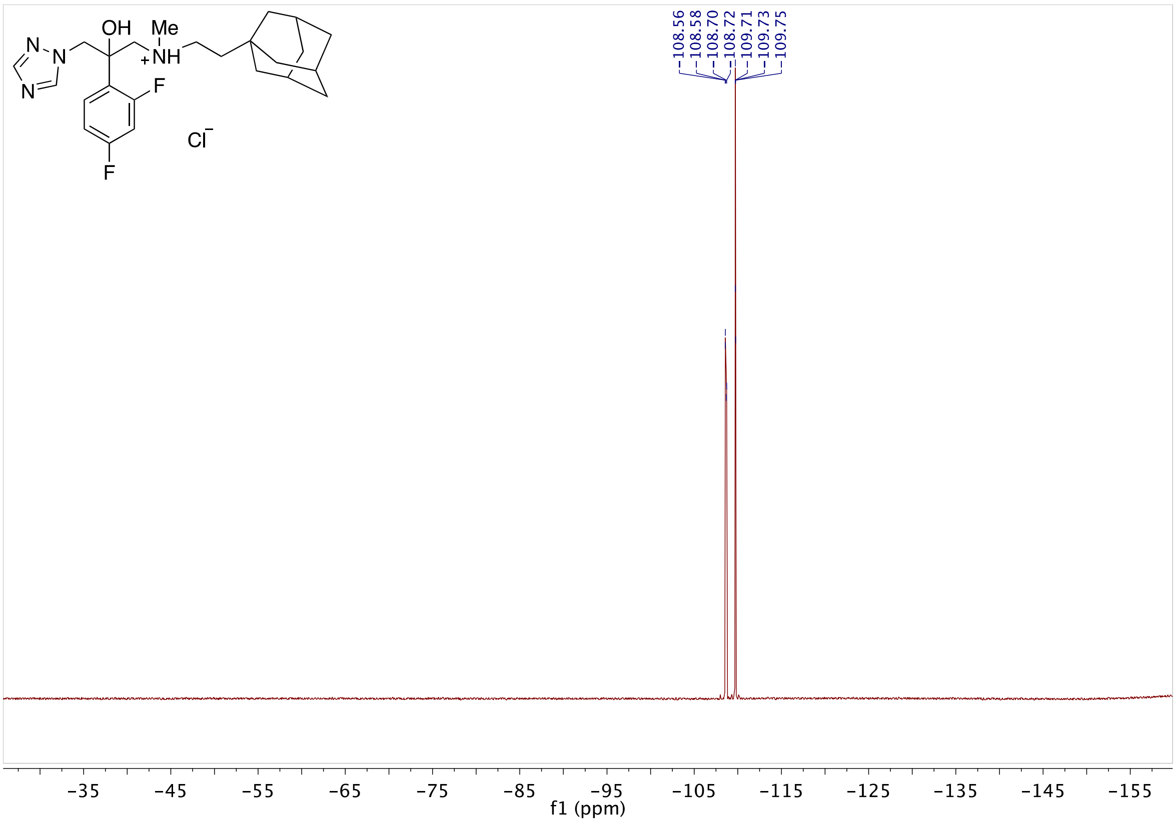

**Fig S119.** 19F NMR of **12a** in $CD_3OD$

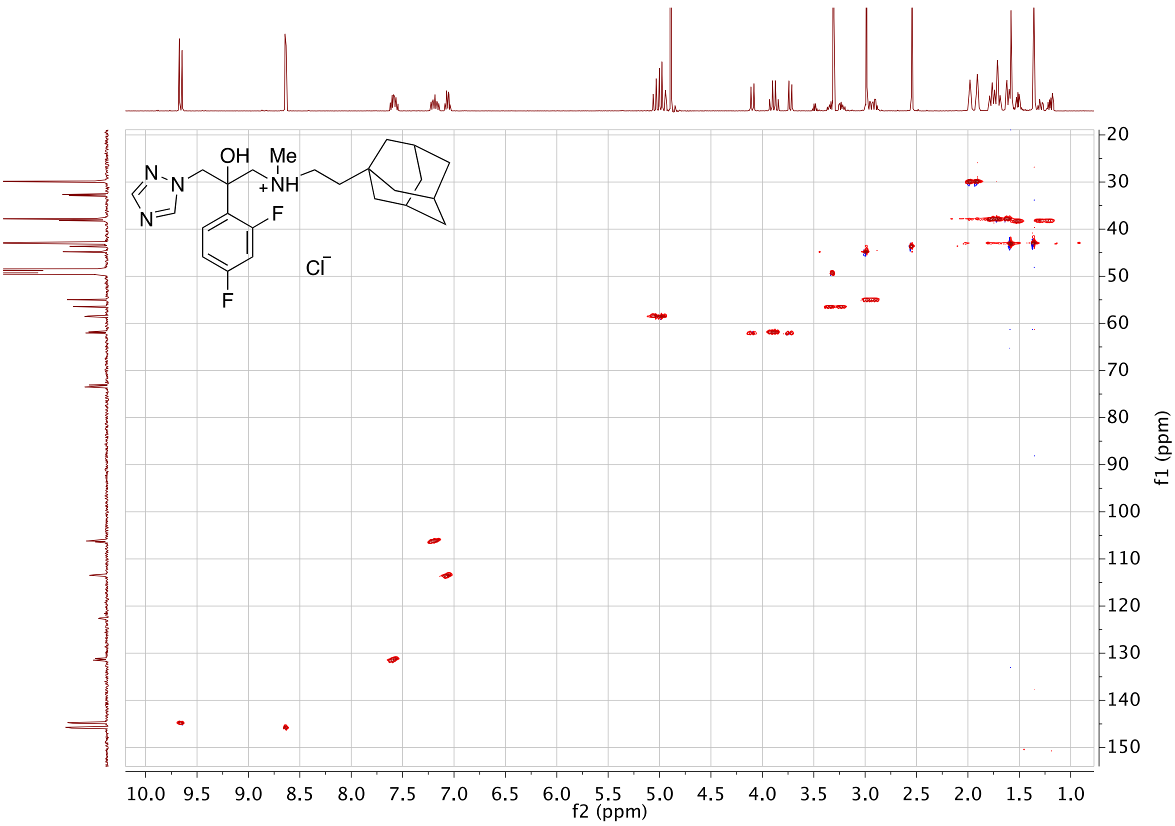

**Fig S120.** HSQC of **12a** in $CD_3OD$

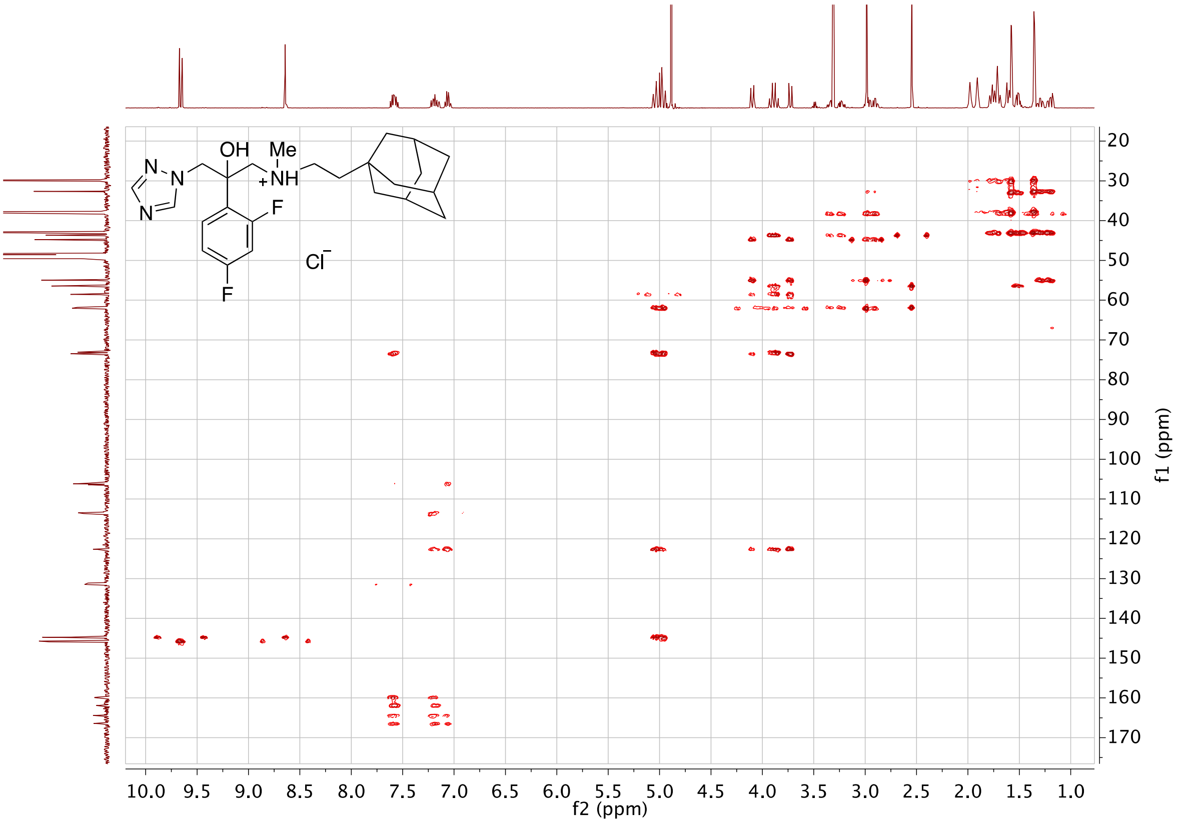

**Fig S121.** HMBC of **12a** in $CD_3OD$
